# Human Parental Relatedness through Time - Detecting Runs of Homozygosity in Ancient DNA

**DOI:** 10.1101/2020.05.31.126912

**Authors:** Harald Ringbauer, John Novembre, Matthias Steinrücken

## Abstract

At present day, human parental relatedness varies substantially across the globe, but little is known about the past. Here we use ancient DNA to provide new insights, leveraging that parental relatedness leaves traces in the offspring’s genome in the form of runs of homozygosity. We present a method to identify such runs in low-coverage ancient DNA data using linkage information from a reference panel of modern haplotypes. As a result, the method facilitates analysis of a much larger fraction of the global ancient DNA record than previously possible. Simulation and experiments show that this new method has power to detect runs of homozygosity longer than 4 centimorgan for ancient individuals with at least 0.3× coverage. We used this new method to analyze sequence data from 1,785 humans from the last 45,000 years. Generally, we detect very low rates of first cousin or closer unions across most ancient populations. Moreover, our results evidence a substantial impact of the adoption of agricultural lifestyles: We find a marked decay in background parental relatedness, co-occurring with or shortly after the advent of sedentary agriculture. We observe this signal, likely linked to increasing local population sizes, across several geographic regions worldwide.

## Introduction

An individual’s parents can be related to each other to varying degrees. For present-day humans much intriguing geographic variation in parental relatedness has been observed. On one end of the spectrum, globally more than 700 million living humans are the offspring of second cousins or closer relatives, and in some regions, the rate of such unions reaches 20-60% (Bittles and Black, 2010). Parents can also be more distantly related to each other, often via many deeper connections in their pedigree, as a common consequence of small population sizes (e.g. Ceballos et al., 2018; Henn et al., 2011; Mondal et al., 2016), or as a consequence of founder effects in tight-knit groups (e.g. Broman and Weber, 1999; Goldschmidt et al., 1960). At the other end of the spectrum, in large populations where cousin marriages are not common, many parents have no recent connections in their pedigree at all (Ceballos et al., 2018). Going back in time, sporadic matings of close kin are documented in royal families of Europe, ancient Egypt, Inca and pre-contact Hawaii (Bixler, 1982; Ceballos and Álvarez, 2013), but comparably little is known about broader patterns of parental relatedness, because archaeological evidence alone is often not informative about mating preferences, especially for prehistoric societies.

The genetic sequence of an individual contains information about the relatedness of their parents, since co-inheritance of identical haplotypes from both parents results in stretches of DNA that lack genetic variation, so called runs of homozygosity (ROH) (Fig. 1A, Broman and Weber, 1999). The more recent a parental relationship, the more frequent and longer the resulting ROH tend to be (Ceballos et al., 2018). ROH can be identified in genome-wide data (e.g. Purcell et al., 2007; Narasimhan et al., 2016a), and this signal has been analyzed for a wide range of purposes in medical, conservation, and population genetics (Ceballos et al., 2018).

**Figure 1:**
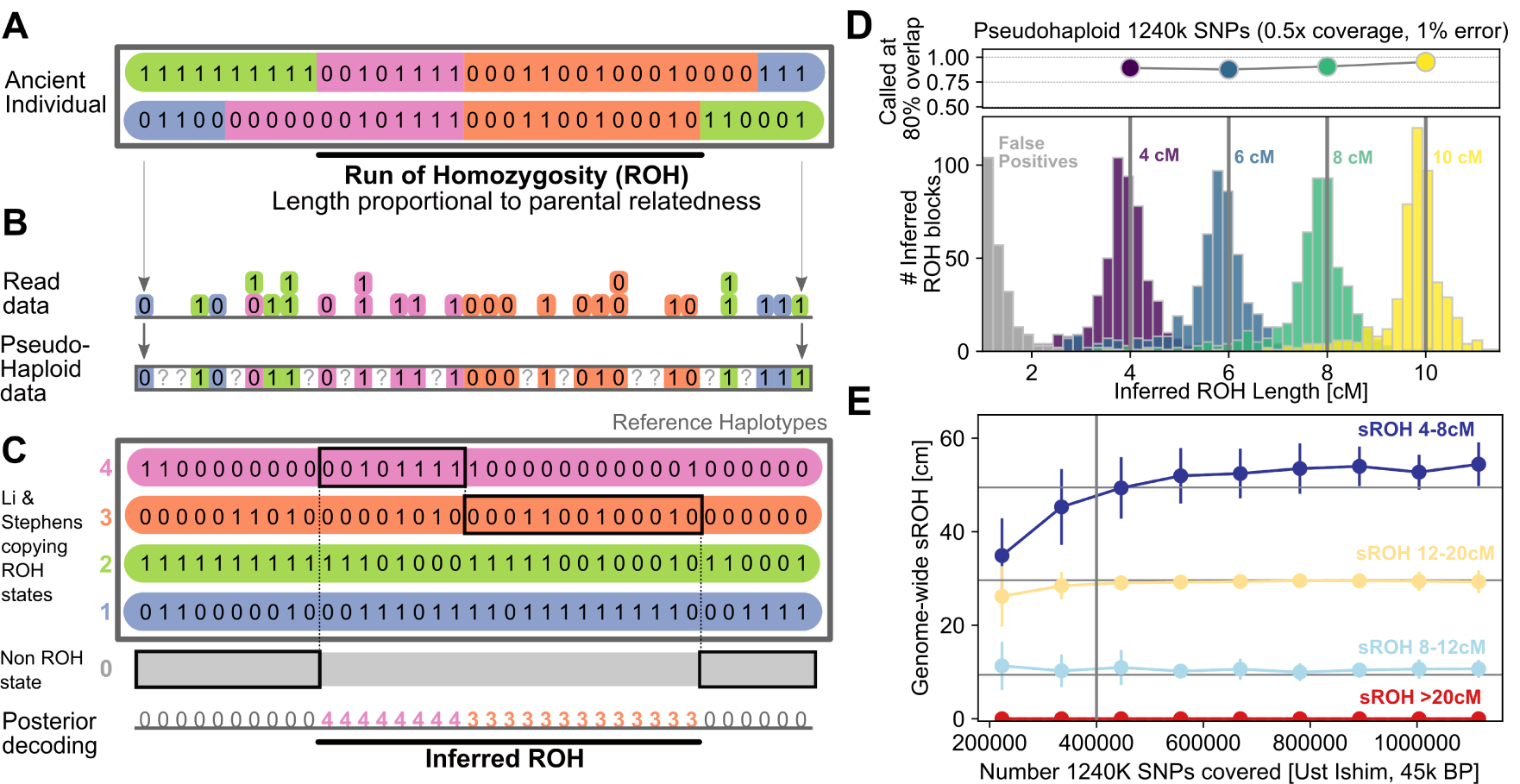
Detecting runs of homozygosity using a reference panel. **Panel A,B**: Illustration of genotype data for a diploid individual (Panel A). Mapping sequencing reads to a biallelic SNP produces counts of reads for each allele (Panel B), from which in turn pseudo-haplotype genotypes, i.e. single reads per site, are sampled (Panel B). **Panel C**: Schematic of Method. A target individual’s genotype data is modelled as mosaic copied from haplotypes in a reference panel (ROH states, colored) and one additional background state (non-ROH, gray). **Panel D**: We applied our method to simulated data with known ROH copied in (see Supp. S1.7 for details). We copied 500 ROH of exactly 4, 6, 8, and 10 cM into 100 artificial chromosomes, and depict histograms of inferred ROH lengths (in color) as well as false positives (in gray) after downsampling and adding errors (0.5 of all 1240K SNPs, with 1% error added). **Panel E**: Down-sampling experiment of a high coverage ancient who lived 45,000 years ago (Ust Ishim man) using a modern reference panel (1000 Genomes dataset). We down-sampled pseudohaploid data to random subsets of the 1240K SNPs, running 100 replicates for each target coverage. We depict mean and standard deviation of the inferred ROH in four length bins (4-8, 8-12, 12-20, and >20 cM). The horizontal lines indicate high confidence estimates using diploid genotype calls from all available data.

Recently, ROH have been identified in ancient DNA (aDNA, e.g. Broushaki et al., 2016; Sikora et al., 2017; Schroeder et al., 2018; Racimo et al., 2020; Mafessoni et al., 2020; Cassidy et al., 2020), that is, genetic material extracted from ancient human remains. This advance is especially promising, as large datasets of aDNA have been generated in the last decade (Skoglund and Mathieson, 2018). However, major challenges persist, coverage for aDNA is often around or less than ∼1× per site (see Fig. S17), and contamination and DNA degradation introduce genotyping errors (Furtwängler et al., 2018). As a consequence, ROH detection is currently only possible for ancient individuals of exceptional high coverage. Recent methodological advances enable identifying ROH in data with at least 5× coverage (Renaud et al., 2019), but this threshold precludes analysis on all but a small fraction of the currently available ancient DNA record.

Here, we present an approach to detect ROH that can identify ROH longer than 4 centimorgan (cM) in individuals with coverage as low as 0.3×. It is designed to perform well for a common type of human ancient DNA data: Pseudo-haploid genotypes (Fig. 1B), which consist of a single allele call for each diploid site. Such data do not convey homozygote versus heterozygote genotype states directly; however, as we show, one can still extract ROH from such data by leveraging linkage disequilibrium information provided by a panel of reference haplotypes. Using this method, we analyze 1,785 ancient individuals from the last 45,000 years, a substantial fraction of the published global human ancient DNA record. We focused on quantifying two domains of parental relatedness: 1) Close-kin unions, measured by the sum of all ROH >20 cM, denoted throughout as sROH_>20_; and 2) Background relatedness as measured by the sum of all ROH 4-8 cM (sROH_[4,8]_). First, we find that matings among first-cousins or closer relatives, though widely practiced today in numerous societies, were generally infrequent in data from the pre-historic societies. Second, we observe decreasing levels of short ROH across many regions coinciding with or shortly after the local Neolithic transition from foraging to agricultural subsistence strategies. This genomic evidence of reduced background relatedness supports long-held evidence of the Neolithic transition involving a major demographic shift towards increased local population sizes.

### Detecting ROH using a haplotype reference panel

Our approach to detect ROH employs a phased reference panel to leverage haplotype data (full details are described in Supp. S1). Briefly, our method utilizes the fact that sequencing reads in a region of ROH are effectively sampled from a single haplotype only, because the maternal and paternal haplotypes of the diploid individual are identical. In contrast, outside an ROH, two distinct haplotypes are carried by the target individual, and thus, the sequencing reads originate from both. As a result, modeling sequencing reads as mosaic of long stretches copied from single reference haplotypes works substantially better within ROH regions (see Fig. 1). To detect this signal, we developed a Hidden Markov Model (HMM) with hidden copying states, one for each reference haplotype, to model copying long stretches from the panel (similar to the copying model of Li and Stephens, 2003), and an additional single non-ROH state as in Narasimhan et al. (2016a). We implemented this algorithm in the software package `hapROH`, available at https://pypi.org/project/hapROH. The default parameters of the current implementation are tuned for pseudo-haploid genotype calls from a widely used capture technology that targets ca. 1.24 million SNPs (hereafter the “1240K” SNP panel, Fu et al., 2015) when using a reference panel of 5,008 haplotypes of present-day human genetic variation (1000 Genomes, Consortium et al., 2015).

### Validating the ROH inference

We tested the method in four scenarios: 1) Spiking ROH segments of various length (4-10 cM) into data (Supp. S1.11), 2) Down-sampling high-coverage ancient individuals (Supp. S1.13), 3) Down-sampling present-day individuals (Supp. S1.14), and 4) Testing different divergence times between the reference panel and the target individual (Supp. S1.12).

For the spike-in experiments, we observe that the power to detect at least 80% of an inserted ROH block was above 85% in all simulated scenarios (Fig. 1D, Supp. S1.11). Bias in the estimated length of the longest overlapping inferred ROH was consistently below 0.5 cM, and we observed no false positives for ROH > 4 cM. In the down-sampling experiments, we tested the ability to recover the sum of ROH segments falling into four length ranges (4-8 cM, 8-12 cM, 12-20 cM, and > 20 cM). When down-sampling the oldest modern human genome sequenced to high depth (Ust-Ishim, 45,000 years old, Fu et al., 2014), the method produces little bias in estimating the respective sROH statistics from pseudo-haploid data with as few as 400,000 of the 1240K sites covered (see Fig. 1E, Supp. S1.13). In the experiments where we down-sampled 599 present-day individuals from a global sample (Supp. S1.14), the ROH inference from pseudo-haploid data performs generally well (sROH_[4,8]_: *r =* 0.925 between diploid ROH calls and pseudo-haploid data, Sroh_>20_: *r =* 0.988, Fig. S9), except for sub-Saharan forager populations. When assessing the impact of different divergence times between the test population and the haplotype reference panel in individuals with a simulated coverage of 1×, we find that, using a European-only reference panel, the method can detect ROH in low coverage test individuals from East Asia and South America but showed much less power for test individuals from West Africa (see Fig. S5).

Together, our tests show that the new method can infer ROH segments longer than 4 cM for individuals with more than 400,000 of the 1240K sites covered at least once, while tolerating sequencing error rates up to 3%. Additionally, our experiments demonstrate that the method can analyze target individuals from populations that diverged from the reference panel up to several ten thousand years ago. Therefore, all present-day and ancient humans that share the out of Africa bottleneck (20,000 – 40,000 BP, Speidel et al., 2019) fall into the range of applicability of our method when using the 1240K marker set and the full 1000 Genomes dataset as haplotype reference panel.

## Application to aDNA data

We then applied our method to a large publicly available dataset of aDNA data (“1240K”, v42.4, released on March 1, 2020, https://reich.hms.harvard.edu) using the 1000 Genomes dataset as haplotype reference panel (see `Methods`). Only 134 individuals in this dataset have sufficient coverage for previous methods that detect ROH (e.g. > 5 × coverage, Renaud et al., 2019) (Fig. S17). Using our new method, 1,833 of the 3,723 individuals can be analyzed with high confidence (400,000 of the 1240K sites covered at least once, Fig. 1, Supp. S1.13). We also integrated a dataset of modern individuals genotyped at the Human Origins SNPs (HO, Lazaridis et al., 2014). After quality control and filtering (see `Methods`), we arrived at a combined dataset of 1,785 ancient and 1,941 present-day individuals, in which we inferred ROH longer than 4 cM.

## Low abundance of long ROH

We first identified individuals with Sroh_>20_ greater than 50 cM. We chose this threshold based on calculations (Supp. S3) and simulations (Supp. S4), which show that in large populations, 88% and 20% of the offspring of first and second cousins, respectively, have sROH_>20_ >50 cM, but less than 1% of offspring of third or more distant cousins do. The 50 cM threshold for sROH_>20_ can also be surpassed in very small isolated populations, specifically, 34% of individuals in populations of size 250 and 8% for size 500 (Fig. S14). Hereafter we refer to this as the “long ROH” threshold, and individuals crossing it as having “long ROH”.

Overall, we find that only 54 out of the 1,785 ancient individuals (3.0%, CI: 2.3-3.9%) have sROH_>20_ above 50 cM (all 54 depicted in Fig. 3A). Generally, these individuals with long ROH do not concentrate in any particular region or time period (Fig. 2B and Fig. 3). The only archaeological cluster (defined in annotations from the source dataset, modified for readability) with more than two individuals is ‘Iron Age Republican Rome’, where 3 of 11 samples reported in Antonio et al. (2019) fall above the long ROH threshold. In the Pontic-Caspian Steppe region, 3 of 54 individuals who lived between 2,600-1,500 BP (5.6%, CI: 1.2-15.4%) exceed the threshold (Fig. 2F), but this signal is not significantly different from the rate in the full dataset. Three individuals with long ROH appear in the late pre-contact Andes region (Fig. 2D). A follow-up study investigates this signal with a larger sample size [Ringbauer et al. 2020, in press]. Notably, 11 of the 54 individuals with long ROH are located on islands: Ordered by time and using the cluster annotations from the publicly available dataset (modified for readability) these are: ‘Sardinia Early Copper Age’ (1 of 1, Supp. Fig. 4), ‘Sweden Megalithic’ (1 of 5, all on Gotland), ‘England Neolithic’ (1 of 16), ‘Chilean Western Archipelago’ (1 of 3), ‘England C-EBA’ (2 of 14, Fig. 2), ‘Russia Bolshoy’ (2 of 6), ‘Vanuatu 1,100 BP’ (1 of 3), ‘Argentina Tierra del Fuego’ (1 of 1), and ‘Indian Great Andaman’ (1 of 1).

**Figure 2:**
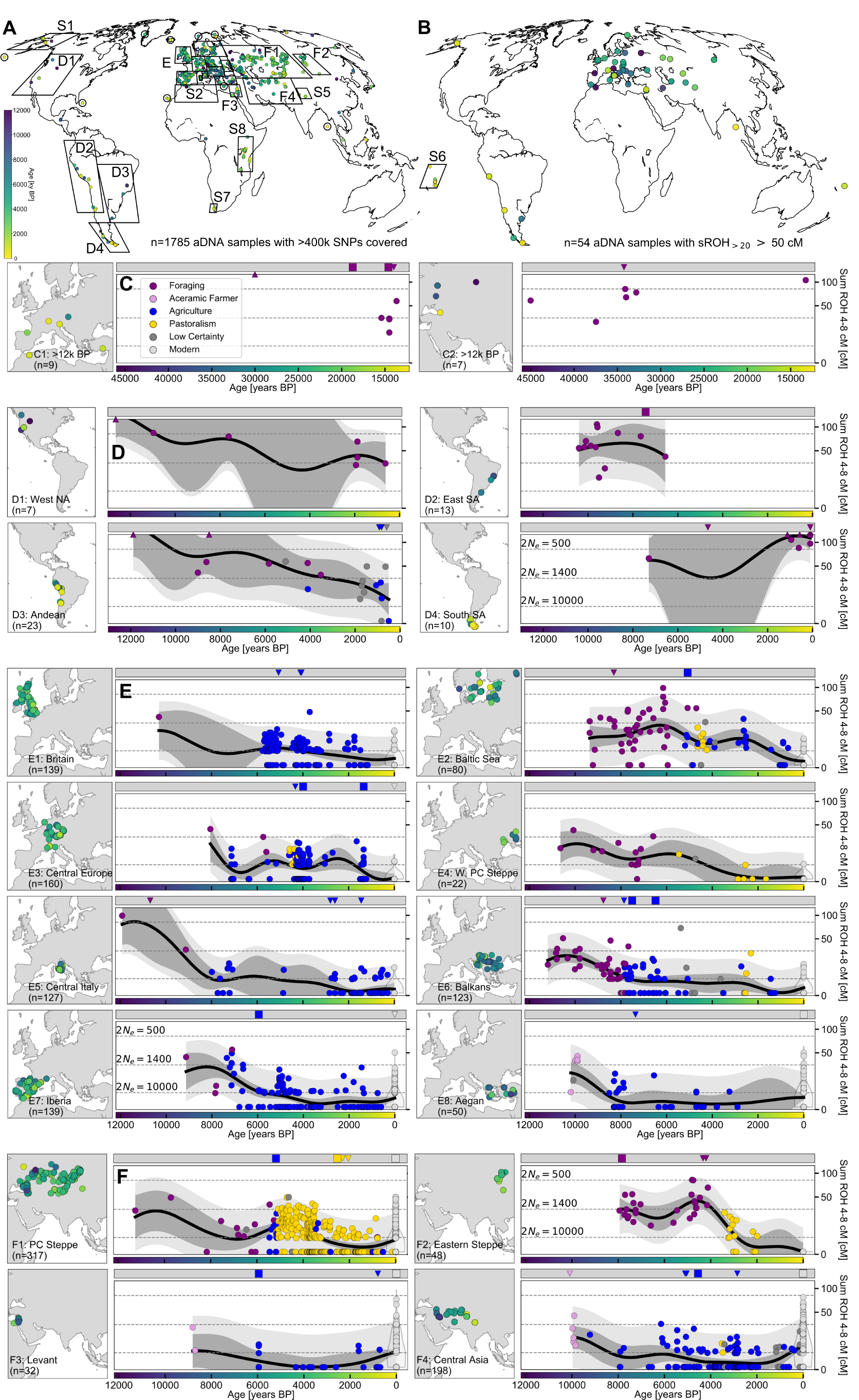
Time transects of major regions. Each circle in the main panels depicts sROH _[4,8]_ for one ancient Individual. We plot individuals that fall into certain regions (See Table S1 for definition). We also show mean estimates calculated from a Gaussian Process model (solid black line, see `Methods`), and 95% empirical confidence intervals for individuals (light gray) and for the estimated mean (dark gray). Note the square root scale (a transform chosen to facilitate Gaussian Process modeling, see `Methods`). Horizontal lines depict the theoretical expectations for sum of ROH blocks 4-8 cM for panmictic population sizes calculated using analytical formulas (Supp. S3). In the gray bar at the top of each panel, we indicate individuals with sROH_>20_ more than 100 (squares) and 50 cM (downward triangles), plausible candidates for offspring of close kin (which have been held out when fitting the GP). Where available, we also show ROH in present-day individuals (light-gray points for each individual, violin plot for density estimate).

**Figure 3:**
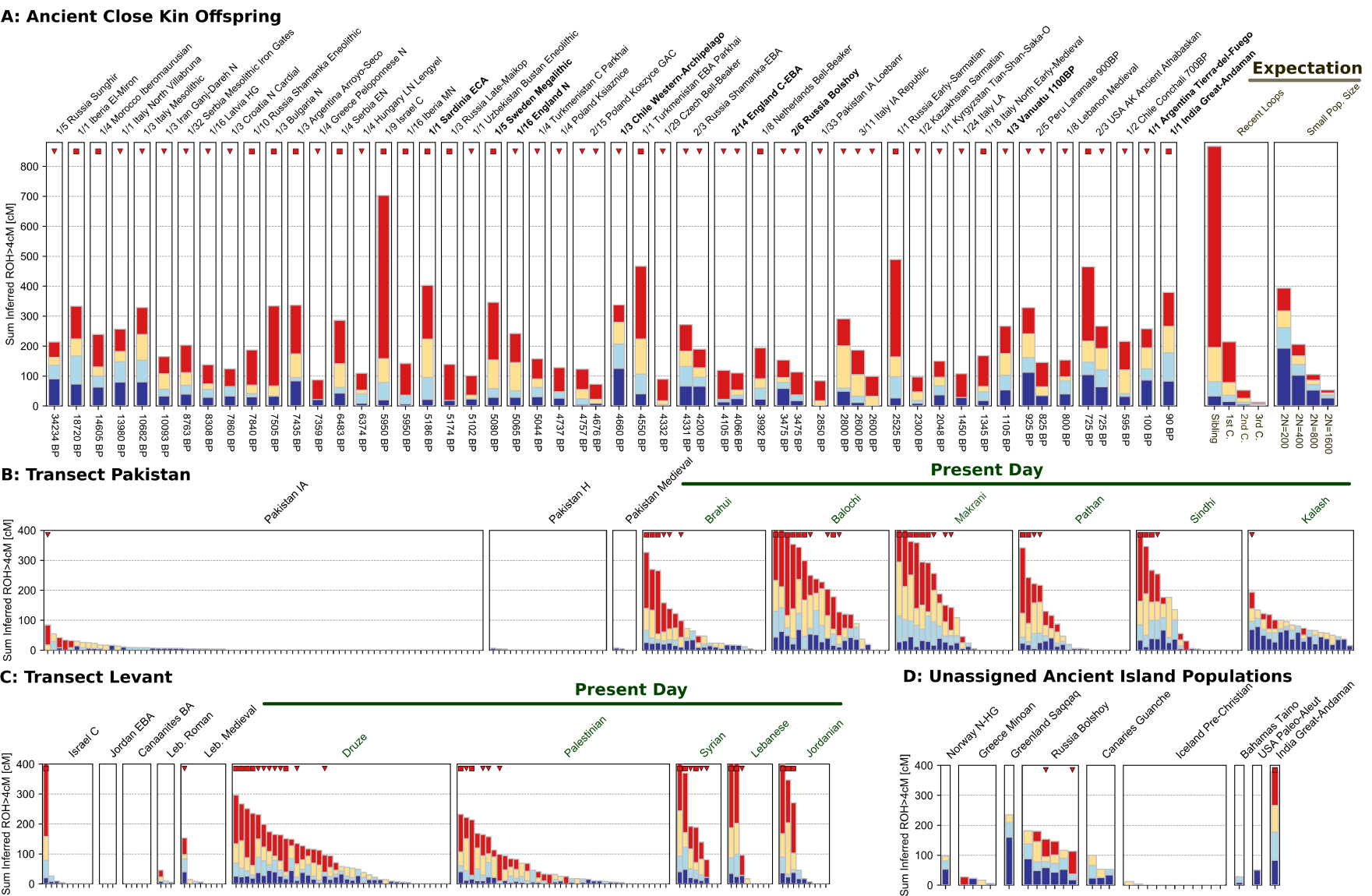
ROH in a subset of ancient and present-day populations. Each individual is represented by stacked vertical bars, where the length of each bar is determined by the ROH of this individuals falling into four length classes (4-8, 8-12, 12-20, and >20 cM, color-coded). **Panel A**: All 54 ancient individuals (out of 1,785) with at least 50 cM sROH_>20_ (x/n indicates number of individuals x exceeding threshold in cluster of size n defined by archaeological label). The labels of the 11 individuals from island populations are printed in bold. We also show a legend (top right) of expected ROH for offspring of close kin or in small populations, based on analytical calculations (Supp. S3). For simulations exhibiting individual variation around the mean see Supp. S3 and Supp. S4. **Panel B,C**: Time transects for regions covering present-day Pakistan and Levant, respectively. Modern individuals are indicated by the green horizontal bars. **Panel D**: Ancient individuals from island populations not assigned to geographic regions (circles in Fig. 2A).

**Figure 4:**
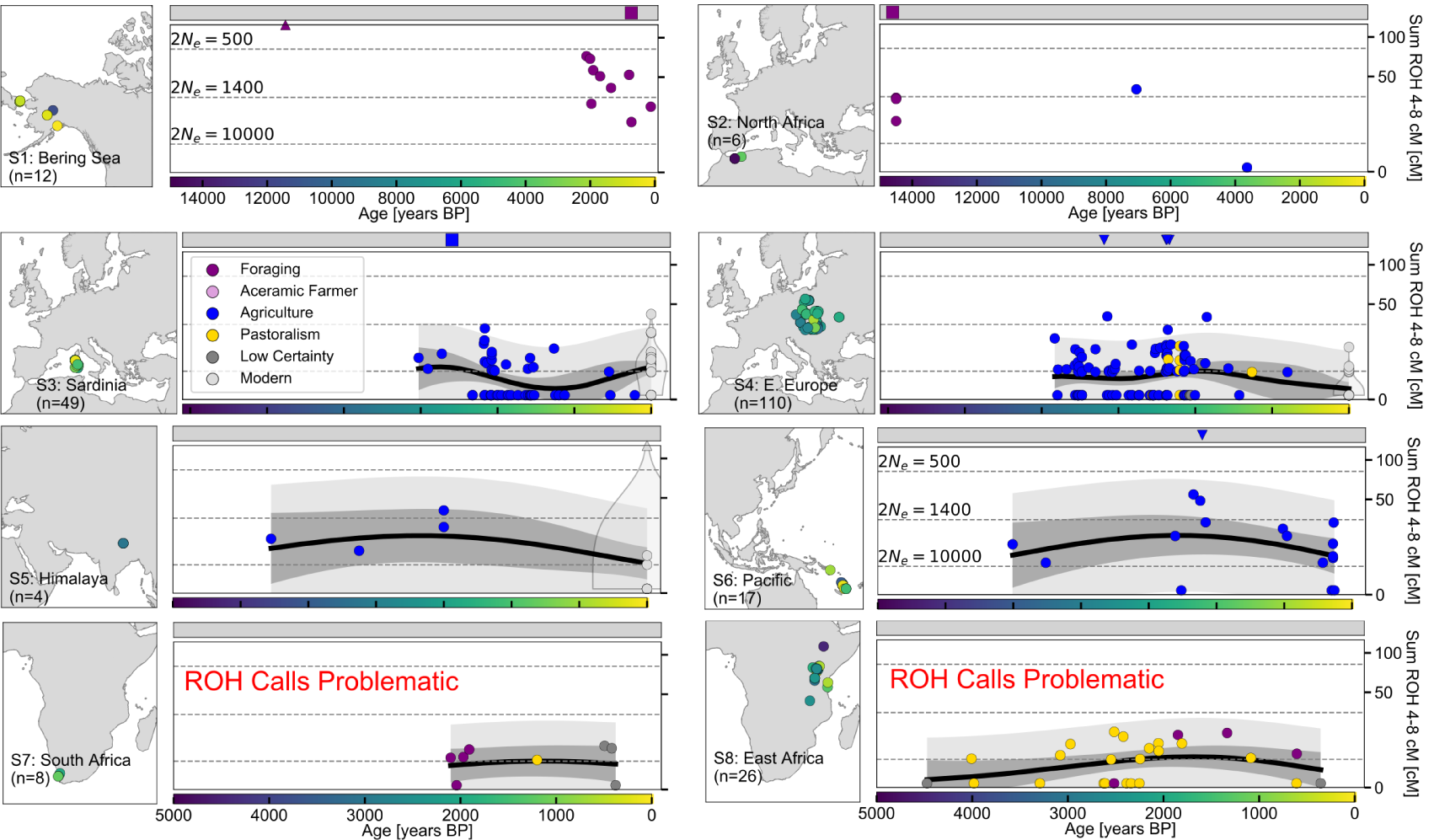
Time transects of eight additional regions. As in Fig. 2: Each circle in the main panels depicts sROH _[4,8]_ for one ancient Individual. We plot individuals grouped by geography (Fig. 2A). We also show mean estimates calculated from a Gaussian Process model (solid black lines, see `Methods`), and 95% empirical confidence intervals for individuals (light gray) and for the estimated mean (dark gray). Horizontal lines depict the theoretical expectations for sum of ROH blocks 4-8 cM for panmictic population of different sizes, calculated using analytical formulas (Supp. S3). In the gray bar at the top of each panel, we indicate individuals with sROH_>20_ more than 100 cM (squares) and 50 cM (downward triangles). Where available, we also show ROH in present-day individuals (light-gray points for each individual, violin plot for density estimate). Note that in our tests `hapROH` could not recover a substantial fraction of long ROH for certain Sub-Saharan ancestries (see Supp. 1.14 for details), thus we warn that the power to call ROH is likely also low for ancient Sub-Saharan African individuals.

**Figure 5:**
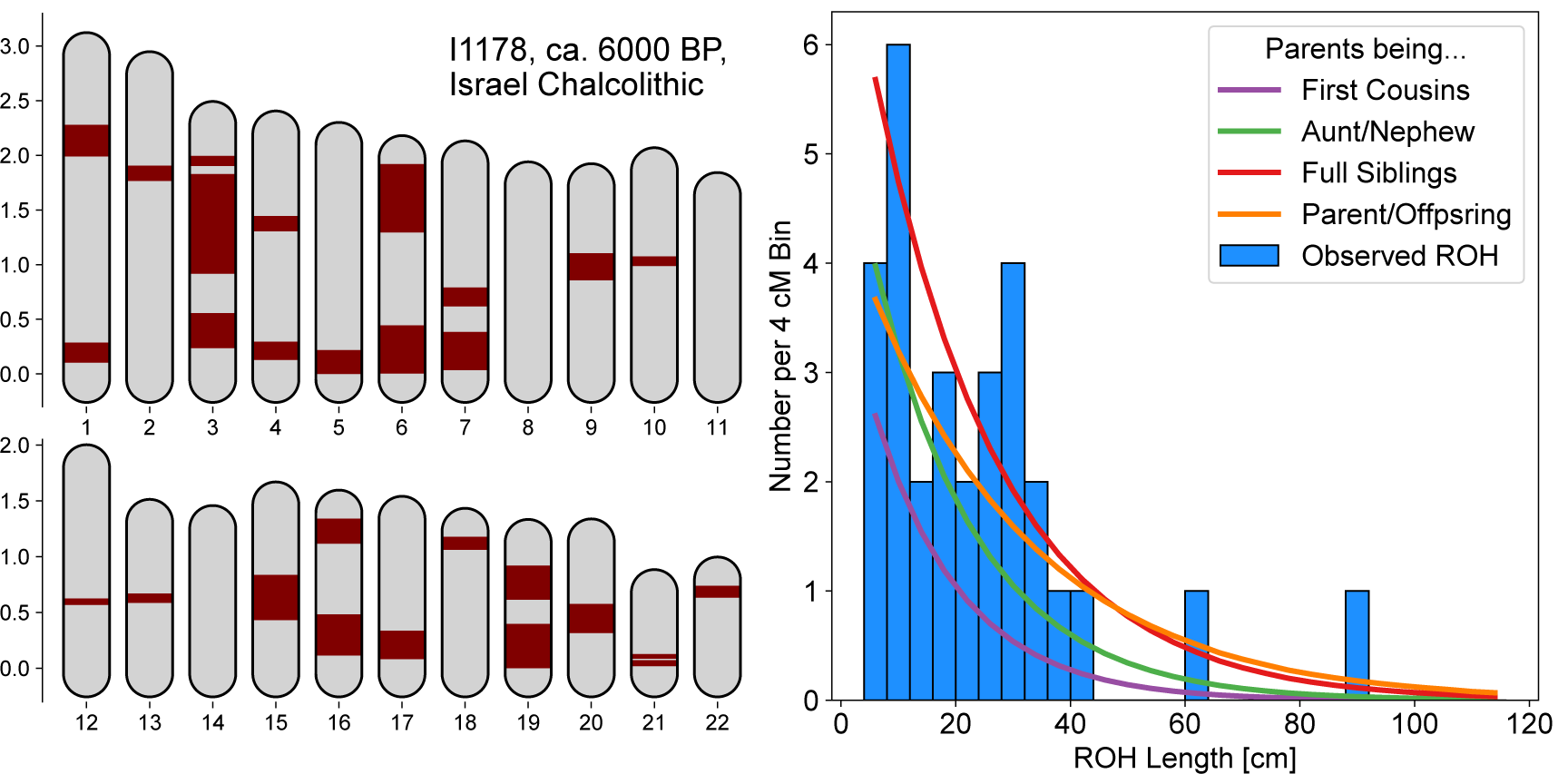
ROH in a 6,000 year old individual. We show ROH of the individual with the highest sum of inferred ROH among all our samples, `I1178`, a male individual context-dated to ca. 4,500–3,900 BCE, reported in Harney et al. (2018). He was inferred to have 703.2 cM of his genome in ROH longer than 4 cM, with the longest ROH spanning 91.1 cM. We depict all these ROH (left panel), with map length annotated in Morgan. We also show a histogram of the ROH lengths, together with expected densities of ROH for certain degrees of parental relationships, calculated as described in Supp. S3 (right panel).

The highest value of sROH_>20_ across the whole dataset (including present-day individuals) is found in a 6,000-year-old Levantine Copper Age individual (`I1178`, Harney et al., 2018) with 545 cM sROH_>20_. The other 8 individuals tested from the same burial site (Peqi’in Cave, Israel Chalcolithic 6000 BP, Harney et al., 2018) had sROH_>20_ values of 0, and very little ROH overall (sROH_>4_ 30 cM). The sum and length distribution of ROH suggest the parents of individual `I1178` were first degree relatives (Supp. Fig. 5), i.e. parent-offspring or full siblings whose offspring will have a quarter of their genome in ROH. We note that the burial context of this male individual was not reported to be exceptional.

The rate of long ROH is substantially higher in the present-day Human Origins dataset; we inferred that 176 of 1,941 modern individuals (9.1%, CI: 7.8-10.4%) have long ROH. In contrast to ancient data, several geographic clusters of long ROH are found, mainly in present-day Near East, North Africa, Central/South Asia, and South America (see ROH grouped per sample label in Supp. Data 1). This signal mirrors previous ROH observations (reviewed in Ceballos et al., 2018) and estimated prevalence of cousin marriages (Bittles and Black, 2010).

In two regions where long ROH are common in the present-day data (Fig. 3, Supp. Data 1) our ancient data contains several ancient individuals, which allowed us to analyze time transects. In the Levant, all five present-day annotated groups in our study (Druze, Palestinian, Syrian, Lebanese, Jordanian) have a high fraction of individuals above the long ROH threshold (30 out of 102 in total, see Fig. 3C). In the ancient sample of this region, only 2 out of 28 analyzed Levant individuals from the Copper Age (n=9), Bronze Age (n=8), Roman times (n=3) to the Middle Age (n=8) fall above this threshold: The first is the Israel Chalcolithic individual with the highest sROH_>20_ in our dataset (see above) (Supp. Fig. 5); the second is a male individual (`SI-38`) excavated from a mass burial in South Lebanon connected to a Medieval Crusader battle, who was found to have local ancestry (Haber et al., 2019). The second region for which we could analyze a time transect is the region of present-day Pakistan. In five out of six modern annotated groups in the dataset (Brahui, Makrani, Balochi, Pathan and Sindhi), many individuals have long ROH (33 out of 98 individuals with sROH_>20_ above 50 cM). In the sixth group, from the Kalash, an isolated valley population, only 1 individual out of 18 exceeds this threshold, despite elevated levels of background ROH being observed (Fig. 3B). In contrast, we infer that only 1 individual out of 75 in this region from the Iron Age (3,200-2,700 BP) has sROH_>20_ above the threshold. None of the 20 individuals from the Historical Period (2,600-1,900 BP) and none of the 4 individuals from the Middle Period (900-400 BP) surpass the long ROH threshold (Fig. 3B).

### Human background relatedness decreased over time

Shorter ROH segments measured by Sroh_[4,8]_ accumulate from parental lineages coalescing on average 10-30 generations ago (Fig. S13); thus, their abundance reflects the size of the ancestral mating pool (“background relatednes”) over approximately the previous half millennium (assuming 30 years per human generation, Fenner, 2005). Because ancestry often spreads out geographically back in time, the probability of recent coalescence and ROH decreases not only with increasing local population size, but also increasing parent-offspring dispersal (Barton et al., 2002; Ringbauer et al., 2017). Assuming that individual mobility is comparable between groups, sROH_[4,8]_ proxies for local population size (Browning et al., 2018). We plotted the values of sROH_[4,8]_ in time transects for 24 major geographic regions that cover 1,763 of the 1,785 ancient individuals (16 regions shown in Fig. 2, 8 regions in Fig. S4, 29 additional individuals from islands are shown in Fig. 3D, and the remaining 22 individuals are depicted in Supp. Data 2). In addition, we tested whether sROH_[4,8]_ differs between subsistence strategies (annotated for most ancient individuals, see `Methods`) in certain regions (PERMANOVA used for Tab. 1 and p-values in text below).

**Table 1:**
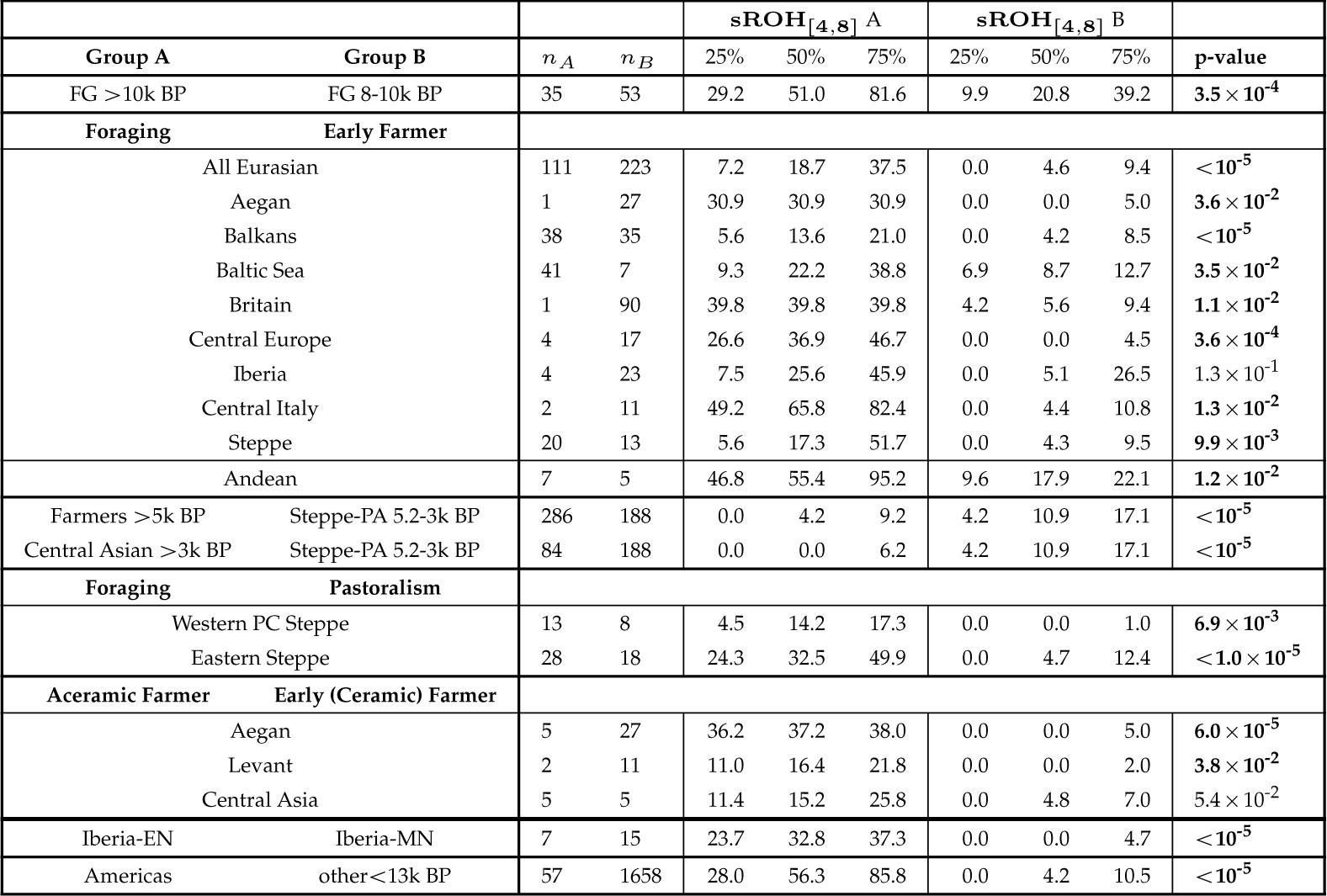
Comparison of Statistics for sROH_[4,8]_ for pairs of groups. For each of two groups (A and B), we calculate the *sROH* 4,8 statistic with individuals having sROH_>20_ > 50 cM removed. We report the total sample size per group (*n*_*A*_ and *n*_*B*_), quantiles of the sROH _[4,8]_ statistic (25%, 50%, and 75%), and a p-value for the pairwise comparison (`PERMANOVA` with 99,999 Permutations, see `Methods`). Early Farmer groups are defined as 2,000 years after the first individual with an “Agriculture” annotation per region (Fig. 2). Other population abbreviations are FG: Foraging, PA: Pastoralism, EN: Early Neolithic, MN: Middle Neolithic.

We find that sROH_[4,8]_ is highest among the most ancient individuals in the dataset and then generally decreases going forward in time. Each of 43 ancient individuals in the global sample dated to before 10,000 BP was inferred to have sROH_[4,8]_ >0, with a median value of 54.5 (39 individuals shown in Fig. 2). We then observe a substantial decline in sROH_[4,8]_ coinciding with the Neolithic transition to sedentary, agricultural lifestyles (Fig. 2). In Western Eurasia, we contrasted individuals from forager cultures to those from early farming cultures (i.e., farming cultures within the first 2,000 years after the first annotated “Agriculture” individual per region). We found that sROH_[4,8]_ decreases substantially in all 8 regional transects which contain both annotated foragers and farmers (p-value < 0.05 in 7 of these 8 transects), and median sROH_[4,8]_ values drop from 13-66 cM per foraging group to 0-9 cM per early farming group (Tab. 1). In the Andes, where agriculture gradually increased in intensity starting around 5,000 BP in a heterogeneous process lasting thousands of years (Piperno and Fritz, 1994), the median sROH_[4,8]_ decreases from 55.4 for foragers to 17.9 for agriculturalists (*p* 1.2 × 10^−2^, Tab. 1).

Detailed inspection of the transitions from foraging to farming reveals interesting finer-scale dynamics. First, for the earliest western Eurasian farmers not using ceramics yet, who lived ∼10,000 years ago and predate the Neolithic expansions into Europe, we still observe short ROH values, with a median sROH_[4,8]_ of 36.7, 16.4, and 15.2 in Aegean, Levant, and Central Asian aceramic farmers, respectively, which is comparable to values observed for western Eurasian foragers (sROH_[4,8]_ ranging from 13-66 cM, Table 1). In all three regions there is a subsequent marked drop to ceramic early farmers, with median sROH_[4,8]_ decreasing substantially to 0, 0, and 4.8, respectively (*p* 6.0 ×10^−5^, 3.8 × 10^−2^, and 5.4 × 10^−2^, Tab. 1).

Furthermore, one ceramic early farming group in our sample stands out: Individuals annotated in the original dataset as Iberian Early Neolithic (7,400-7,000 BP, Olalde et al., 2019) have median sROH_[4,8]_ of 32.8 cM, which is substantially higher than in other early Eurasian farmers (median sROH_[4,8]_: 0-8.7 cM, Tab. 1). However, in Iberian Middle Neolithic farmers (6,800-4,600 BP) ROH decreases (median sROH_[4,8]_=0, *p <* 10^−5^, Tab. 1) and becomes more typical of other early European farmers. As the early Iberian individuals have exceptionally high early farmer ancestry (> 90% Olalde et al., 2019), this signal cannot be explained by forager (hunter gatherer) ancestry. However, archaeological evidence of a rapid maritime spread (Cardial Ware expansion) within a few hundred years around 7,500 BP (Zilhão, 2001) provides one plausible explanation of this increased abundance of short ROH in the Early Neolithic, as a rapid spread could have caused an initial bottleneck. Moreover, an initially small population of early farmers would explain why forager admixture substantially increased in Middle Neolithic Iberians and remained one of the highest of European Neolithic populations (∼25%, Olalde et al., 2019).

In the ancient Americas, elevated sROH_[4,8]_ values evidence sustained high levels of background relatedness. This signal is found across all American regions: Western North America (West NA) (Fig. 2D1, median sROH_[4,8]_=67.1 cM excluding long ROH individuals, n=7), Eastern South America (East SA) (Fig. 2D2, 59.0 cM, n=13), Andean (Fig. 2D3, 31.5 cM, n=20), Southern SA (Fig. 2D4, 112.5 cM, n=8) and Beringia (Fig. 4S1, 52.4 cM, n=9). This abundance of ROH (overall median sROH_[4,8]_ = 56.3) is higher than the rest of the global sample in the same broad time period (a13k years ago, median 4.2 cM, *p*<10^−5^, Tab. 1). Since sROH_[4,8]_ is driven by co-ancestry within the last few dozen generations (Supp. S3), this elevated sROH_[4,8]_ cM cannot be explained by bottlenecks during early migrations into the Americas, but one needs to invoke more recent, sustained small effective population sizes. Overall we observe little temporal variation (Fig. 2), with one exception in the dataset being Andean populations around the time of the shift to agriculture (see above; also note the dataset does not include individuals from other early centers of agriculture in the Americas, e.g. Central Mexico, eastern North America).

Another observation of elevated ROH on a large geographical scale is found in the Eurasian Steppe, where early pastoralist groups all have substantial amounts of sROH_[4,8]_ (Steppe-PA 5.2-3k BP, median sROH_[4,8]_ = 10.9, Tab. 1), including the Yamnaya (median 17.5 cM, n=17), Afanasievo (18.1 cM, n=22), Sintashta (5.7 cM, n=21), Okunevo (24.5 cM, n=12) and Srubnaya (4.8 cM, n=19). These sROH_[4,8]_ levels are significantly higher than in Western Eurasian farmer populations before 5,000 BP (median 4.2 cM, *p <* 10^−5^, Tab. 1), and, notably, also significantly higher than their southern contemporaneous neighbors, sedentary farmers from Central Asia (median 0, *p <* 10^−5^, Tab. 1). In samples from the Western Pontic-Caspian Steppe (present-day Ukraine and Moldavia), at the transition from foragers to pastoralists, we observe a substantial decrease of sROH_[4,8]_ from median 14.2 to 0 (*p* 6.9 × 10^−3^, Tab. 1). Similarly in the Eastern Steppe (around Lake Baikal and present-day Mongolia), a shift from foragers to pastoralism coincides with a significant reduction in sROH_[4,8]_ (median 32.5 to 4.7, *p <* 10^−5^). We note that in both the Western and Eastern Steppe many of the pastoralists in our sample are from 3,000-2,000 BP (Scythian and Xiongnu, respectively), substantially later than the early pastoralists mentioned above.

## Discussion

We developed a method for measuring ROH in low coverage ancient DNA. Analyzing aDNA data from 1,785 individuals, this tool enabled us to generate new evidence for two key aspects of the human past: Identifying long ROH (>20 cM) provided insight into past prevalence of close kin unions, whereas short ROH (4-8 cM) revealed patterns of past background relatedness that proxies for local population sizes.

Analyzing long ROH, we found that only 1 out of 1,785 ancient individuals had ROH typical for the offspring of first-degree relatives (e.g. brother-sister or parent-offspring). Historically, matings of first degree relatives are only documented in royal families of ancient Egypt, Inca and pre-contact Hawaii, where they were sporadic occurrences (Bixler, 1982). The only other example of an offspring of first-degree relatives found using aDNA to date is the recently reported case from an elite grave in Neolithic Ireland (Cassidy et al., 2020). Our findings are in agreement that first-degree unions were generally extremely rare in the past.

Further, we find that only 54 out of 1,785 ancient individuals (3.0%, CI: 2.3-3.9%) have long ROH typical for the offspring of first cousins (88%) and less commonly observed for second cousins (20%). Such long ROH can also arise as a consequence of small mating pools (e.g. 8% in randomly mating populations of size 500, which may explain the long ROH we observed on certain island populations). Therefore, the rate of long ROH is an upper bound for the rate of first cousin unions. On the other hand, because of incomplete power some long ROH may be missed in our empirical analysis; however, even if the method would fail to detect half of all ROH >20 cM, well below the power that we observed in our simulations, we would still detect 60% of first cousins (see Supp. S5). We conclude that in our ancient sample substantially less than 10% of all parental unions occurred on the level of first cousins.

In two specific regions with high levels of long ROH in the present-day (Ceballos et al., 2018), the dataset contained a sufficient number of ancient individuals to allow analyzing time transects and for both transects (the Levant and present-day Pakistan), we observe a substantial shift in the levels of long ROH. In contrast to the high abundance of long ROH typical of close kin unions in the present-day individuals, long ROH were uncommon in the ancient individuals, including up to the Middle Ages. Additional data from these regions and others with high levels of long ROH today, such as North Africa as well as Central, South, and West Asia (Ceballos et al., 2018), will help resolve with more precision the origin and spread of these well-studied kinship-based mating systems (King-Irani, 2004; Korotayev, 2000). Overall, our results show how an ROH-based method can be used to inform understanding of shifts in cultural marriage/mating practices.

As a second major finding, we observed that human background relatedness as measured by short ROH (4-8 cM) decreased markedly over time in many regions, with a significant drop occurring during or shortly after the transition to sedentary farming (“Neolithic Transition”). Assuming that early farmers had no increased individual mobility compared to foragers, which would agree with observations in present-day forager populations (MacDonald and Hewlett, 1999), the substantial decrease of short ROH evidences markedly increasing local population sizes. As such, our results provide a new genetic line of evidence for the inference that local population sizes increased following the Neolithic transition (Diamond and Bellwood, 2003; Ammerman and Cavalli-Sforza, 2014; Bacci, 2017). Future application of our methodology at finer-scales may give additional insights regarding the timing and variance in how the adoption of farming and related changes impacted population growth.

For individuals from early Eurasian Steppe pastoralist groups, we observe an intermediate level of short ROH. These early cultures (e.g. the Yamnaya) have drawn much attention in archaeological and ancient DNA studies to date, as they are a strong candidate for the origin of Indo-European languages and of several population expansions (Anthony, 2010; Allentoft et al., 2015; Haak et al., 2015; de Barros Damgaard et al., 2018; Narasimhan et al., 2019). The observed elevated rate of short ROH provides evidence that many matings occurred within and among small, related groups. An alternative interpretation for the abundance of short ROH could be that burial sites (“Kurgans”) represent a biased sample of societal classes with more short ROH than the general population (Anthony, 2010). However, as short ROH probe parental ancestry up to several dozen generations into the past, this signal would require reproductive isolation between societal strata maintained over many generations. Therefore, it is likely that at least part of the signal is due to Steppe populations having comparably low population densities or a recent bottleneck.

Our analysis is limited by several caveats. Importantly, skeletal remains accessible by archaeological means often do not constitute a random cross-section of past populations. While levels of background relatedness are expected to be broadly similar within a mixing population, rates of close kin unions can vary substantially because of social structure; e.g. elite dynasties may practice close kin unions despite them being uncommon in the general population. Another limitation is the incomplete sampling of the current aDNA record and that for much of the world, we necessarily make inferences from small numbers and sparse sampling. Future work analyzing the rapidly growing aDNA record will help to resolve additional details of other social and cultural factors operating at finer scales (e.g., leveraging more precise timings of shifts and more subtle shifts in ROH patterns). In particular, future studies focusing on specific localized questions can combine archaeological and genetic evidence such as the one provided by the new methodology presented here (Racimo et al., 2020).

In addition to denser sampling, there are several ways how our analysis can be improved upon by future work. Here we analyzed long ROH (>20 cM) and short ROH (4-8 cM) only. While this dichotomy helped us to disentangle more clearly recent and distant parental relatedness, we expect that future work refining the downstream analysis of ROH will be able to extract more subtle signatures by looking across all ROH scales. Furthermore, we note that our application focused on a set of SNPs widely used for human ancient DNA (1240K SNPs). For whole-genome sequencing data (available for a subset of the data analyzed here), using all genome-wide variants would likely lower the requirements for coverage below the current limit of 400,000 of the 1240K SNPs covered at least once (corresponding to ca. 0.3× whole-genome sequencing coverage).

Identifying ROH can also be a starting point for other powerful applications: ROH consist of only a single haplotype (the main signal of our method), which is therefore perfectly phased, a prerequisite for powerful methods relying on haplotype copying (e.g Lawson et al., 2012) or tree reconstruction (e.g. Kelleher et al., 2019; Speidel et al., 2019). Moreover, long ROH could be used to estimate contamination and error rates, an important task in ancient DNA studies (Furtwängler et al., 2018). ROH lack heterozygotes, allowing one to identify heterozygous reads within ROH that must originate from contamination or genotyping error, similar to estimating contamination from the hemizgyous X chromosomes in males (Korneliussen et al., 2014). Another promising future direction is the development of a method to identify long shared sequence blocks in ancient DNA not only within (ROH), but also between individuals, called identity-by-descent (IBD). Calling IBD between individuals would substantially increase power for measuring background relatedness, since signal from every pair of individuals could be used. Moreover, a geographic IBD block signal is highly informative about patterns of recent migration (Ralph and Coop, 2013; Palamara and Pe’er, 2013; Ringbauer et al., 2017; Al-Asadi et al., 2019). Extending our method to similarly use linkage information from reference haplotypes when detecting IBD could enable such analyses in low coverage ancients individuals.

Finally, the analysis of ROH has additional implications beyond human demography and kinship-based mating systems. In many plants and animal species, ROH are more prevalent (due to different mating systems, small population sizes, or domestication), and the study of ROH may be particularly interesting for understanding early plant and animal breeding, as actively controlled mating among domesticates would be expected to alter ROH (Frantz et al., 2020). For aDNA from extinct or endangered species, ROH can shed light on the extinction and inbreeding processes, as is observed for example in aDNA from high-coverage Neanderthal individuals (Prüfer et al., 2014; Kuhlwilm et al., 2016; Prüfer et al., 2017; Mafessoni et al., 2020), or modern DNA from Isle Royal wolves (Robinson et al., 2019). Finally, as ROH expose rare deleterious recessive alleles (Narasimhan et al., 2016b), the temporal dynamics of ROH are relevant for understanding the evolutionary dynamics of deleterious variants and health outcomes (Szpiech et al., 2019; Robinson et al., 2019; Clark et al., 2019; Walters et al., 2019). Overall, we hope new methods building upon the core ideas of our approach will contribute useful tools for studying low-coverage data from a wide range of natural populations.

## Supplemental Figures

## Methods

### Calling ROH in a global dataset

To detect ROH, we developed a method, `hapROH`, which is based on a hidden Markov model (HMM) with ROH and non-ROH states and uses a panel of reference haplotypes. The detailed method description, parameter settings, and evaluation via simulation and downsampling are provided in Supp. S1, Supp. S1.7, Supp. S1.11 and Supp. S1.14. The software is available at https://pypi.org/project/hapROH/.

### Empirical dataset

The global ancient DNA dataset we analyze originates from a curated dataset of published ancient DNA (“1240K”, v42.4, released on March 1, 2020, https://reich.hms.harvard.edu). This release provides ancient DNA data in pseudo-haploid format for 1.24 million SNPs (The “1240K” SNP panel). We added an additional 40 ancient Sardinian individuals in pseudo-haploid format from Marcus et al. (2020) that have not yet been compiled into the global reference dataset. We filtered to individuals that contained `PASS` in the `ASSESSMENT` column of the meta-data table in order to remove individuals with possible contamination. We furthermore removed individuals with insufficient permissions (see `Ethics Statement`). For ancient individuals that had multiple genotypes listed, we kept the record with the highest coverage. Furthermore, we removed all Neanderthal and Denisovan individuals, as well as the individual `tem003`, for which initial analysis showed that it has all of chromosome 2 in ROH, but no other long ROH. Finally, we kept only individuals with at least 400,000 SNPs of the 1.24 million covered, the approximate cutoff above which our method can provide robust ROH inference (Fig. S1). For present-day data, we downloaded the Human Origins dataset with diploid genotype calls for ca. 550,000 autosomal SNPs (Lazaridis et al., 2014), that are a subset of the 1240k SNPs.

We applied `hapROH` to the pseudo-haploid data for the 1,785 ancient individuals and the diploid data for the 1,941 modern individuals that pass our quality thresholds. We used all SNPs with available data for which the reference and the alternative allele matched the information in the reference panel, set the respective emission probabilities to values designed for these two types of data (Supp. 1.3), and used the default parameters of `hapROH` that were optimized for the 1240K SNPs (Supp. 1.8). For the haplotype reference panel, we used the full global set of 5,008 phased haplotypes of the 1000 Genomes Project dataset (Phase 3, release 20130502) accessible via ftp://ftp.1000genomes.ebi.ac.uk/ (Consortium et al., 2015), down-sampled to the biallelic 1240K SNP markers with `bcftools` (version 1.9). We report the detailed ROH calls for all individuals in Supplemental Information 1.

### Annotation of food economy

For each ancient individual, we annotated the primary subsistence strategy into standard broad categories of food production (Haviland et al., 2013), using descriptions of the archaeological sites and cultural affiliations. We used three main labels: We denoted 1) hunter gatherer and horticulture lifestyles based on collecting wild plants, hunting, or fishing with the label “Forager”; groups that practiced substantial amounts of sedentary farming (e.g. cereals and domesticates observed in the archaeological record) as “Agricultural”, and groups with nomadic and semi-nomadic mobile lifestyles based on herding and breeding of domestic animals (e.g. cattle) as “Pastoralist”. Groups that had intermediate and transitory lifestyles were annotated using the plausible dominant food economy of the associated archaeological culture. To better resolve the transition to agricultural food production, we denoted early groups that practiced agriculture, but lack ceramics in the archaeological record as “Aceramic Farmers”. Individuals and groups for which the archaeological record does not contain sufficient information to annotate a subsistence strategy were labelled as “Uncertain”. We stress that archaeological evidence is often sparse and assignments are frequently interpretations of various lines of evidence, therefore assessments might change with updates to the archaeological record. Here we tolerate some error, since we address questions regarding very broad temporal and geographic patterns, but we advise against using our subsistence assignments as a reference for questions on a finer scale.

### Detecting offspring of close relatives from ROH

We screened all individuals for ROH longer than 20 cM to identify potential offspring of close relatives. Previous work (Chiang et al., 2016) had demonstrated that pairwise identity-by-descent (IBD) > 20 cM, which translate to ROH in the offspring, are very unlikely to be a concatenation of multiple shorter IBD blocks. Moreover, recombination quickly breaks up long ancestry segments of the genome and thus most long ROH originate from co-ancestry within only a small number of generations back. Therefore, if the fraction of the genome in ROH longer than 20 cM in an individual is large, this provides strong evidence for a close relationship of its parents. We report individuals where the sum of all such ROH exceeds 50 cM as potential offspring of closely related parents (i.e. sROH_>20_ > 50). This cutoff is motivated by analytical calculations and simulations, see Supp. 3 and Supp. 4 for details. Briefly, this threshold detects a large fraction of close kin offspring (parents being first cousin or closer) while also being insensitive to background relatedness unless a population has a very small size (< 500).

### Gaussian Process Modeling of short ROH

To visualize the trend of the abundance of ROH in the individuals in certain regions over time, while still conveying the levels of uncertainty due to varying sample sizes, we fit a Gaussian Process (GP) model (Williams and Rasmussen, 2006) using the Python package `scikit-learn` (Pedregosa et al., 2011). As input, we used the square root of the sROH_[4,8]_ statistic to stabilize its variance (McCullagh and Nelder, 1989), since sROH_[4,8]_ corresponds closely to count data, which can be approximated by a Poisson distribution. Furthermore, since we use this statistic as a proxy for background relatedness (which in turn proxies for local population size), we removed all individuals with sROH_>20_ above 50 cM when fitting the GP model, to minimize the impact of putative offspring of close kin on this analysis (Supp. Fig. S12).

For the variance model of the GP, we used a standard squared-exponential covariance kernel summed with a residual white noise kernel. In preliminary analyses, we estimated all parameters of the model via maximum likelihood, but we found that these estimates appeared to over-fit the data for several time transects. Thus, we set custom length scales for the covariance kernel for each transect (1,500 for all non-American populations and 2,000 for American populations, because they had larger temporal sampling gaps) and only fit the two coefficients of the squared-exponential and white noise kernel. To visualize the final output, we estimated the variance of the predicted mean across a dense set of time points (Williams and Rasmussen, 2006). We estimated the uncertainty of the predicted mean and the uncertainty of each individual point and plotted both as 95% confidence interval bands (± 1.96 standard deviations) on a dense grid.

### Analytical expectations of ROH

To aid interpretation of ROH, we visualize expectations of sROH using formulas describing ROH of closely related parents in otherwise outbred populations and finite populations without substructure, which we derived in a unified framework (Supp. S3). These formulas have been derived previously (e.g., Carmi et al., 2014; Browning and Browning, 2015). In addition, we verified these formulas by simulating ROH for these demographic scenarios and comparing expected sROH values to empirical averages (Supp. S3 and Supp. S4).

### Comparing ROH between groups

To test significant differences in the distributions of the sROH statistics between two groups, we applied the Permutational multivariate analysis of variance method (PERMANOVA Anderson, 2001), which calculates a pseudo-F statistic and assesses its significance via permutation tests. We used the `permanova` function implemented in the Python package `skbio`, and based the distance matrices on absolute differences of individual’s sROH_[4,8]_. For each test, we ran 99,999 permutations (minimal p-Value: p=10^−5^). As with the GP modeling, we removed all individuals with sROH_>20_ above 50 cM when comparing distributions of sROH_[4,8]_ between groups.

## Supporting information

Supp. Data 1: Bar plots of ROH in present-day sample

Supp. Data 2: Bar plots ofROH in ancient sample

## Author Contributions

We annotate author contributions using the CRediT Taxonomy labels (https://casrai.org/credit/). Where multiple individuals serve in the same role, the degree of contribution is specified as ‘lead’, ‘equal’, or ‘support’.

- Conceptualization (Design of study) – lead: HR; support: JN, MS
- Software – lead: HR; support: MS
- Formal Analysis – HR
- Data Curation – HR; support: JN
- Writing (original draft preparation) – lead: HR; support: JN, MS
- Writing (review and editing) – input from all authors
- Supervision – equal: JN, MS
- Project Administration – equal: JN, MS
- Funding Acquisition – JN

## Code Availability

A Python package implementing the method is available on the Python Package Index (https://pypi.org/project/hapROH/) and can be installed via *pip*. The documentation provides example use cases as blueprints for custom applications. Code developed for the analysis and data visualization presented here is available at the github repository https://github.com/hringbauer/hapROH.

## Data Availability

The modern dataset (Human Origins Lazaridis et al., 2014) as well as the ancient dataset we analyzed (Version V42.4) can be downloaded from https://reich.hms.harvard.edu. The processed reference panel that we used in our tests and analysis (haplotypes from the 1000 Genomes dataset downsampled to biallelic SNPs at 1240k sites) is available upon request.

## Ethics Statement

We only analyzed previously generated, publicly available genetic data. For all data, we contacted the corresponding authors of each original study regarding our project and publication plan. We included in our final analysis the data from all studies for which we received a response confirming the use is consistent with the original permits.

## Acknowledgements

We thank the original study authors for sharing their data publicly, and David Reich and his lab for compiling and making publicly accessible a normalized pseudohaploid compilation of those data. We thank David Anthony and Alissa Mittnik for reviewing parts of the subsistence strategy annotations and for helpful discussions. We thank Arjun Biddanda, Shai Carmi, David Schloen, Lars Fehren-Schmitz, Montgomery Slatkin and Mashaal Sohail for their comments on the manuscript. Funding for HR and JN was provided by NIH grant R01HG007089 and R01GM132383 to JN.

