## Supplementary Information for "Human Parental Relatedness through Time - Detecting Runs of Homozygosity in Ancient DNA"

July 2020

#### 1 The hidden Markov model

We first describe how to model diploid genotype data $y$ from a focal individual and a reference panel of $n$ phased haplotypes $x_1, \dots, x_n$ at a set of $L$ loci, assuming biallelic markers. Thus, $y \in \{0, 1, 2\}^L$ and $x_i \in \{0, 1\}^L$. In Section 1.3, we describe how types of data $y$ relevant to applications using low-coverage sequencing data, like ancient DNA, can be modeled by treating the unobserved diploid genotypes as latent variables and using appropriate emission probabilities.

Throughout, we measure the distance between loci along haplotypes in genetic map units (i.e. Morgans) $\mathbf{r} = r_1, \dots, r_{L-1}$, where $r_l$ denotes the distance between locus $l+1$ and $l$. We assume that a genetic map is available, which is the typical case for humans and model organisms. If no genetic map is available, the map distances can be approximated using the average recombination rate, but we note that here we only tested scenarios where a map is available.

##### 1.1 State Space

The Hidden Markov model (HMM) can assume any of $n+1$ hidden states $0, \dots, n$ at every marker $l$, where $n$ is the number of haplotypes in the reference panel. As we outline below, the $0$-th state represents that the focal individual is not in a ROH at the respective marker, and has emission probabilities according to Hardy-Weinberg proportions, while the states $1, \dots, n$ are the classical copying states (Fig. 1). In each of these copying states (denoted here as the ROH states), we model the copying as in the original Li & Stephens model, with one important modification: We assume that the genotype of the focal individual $y$

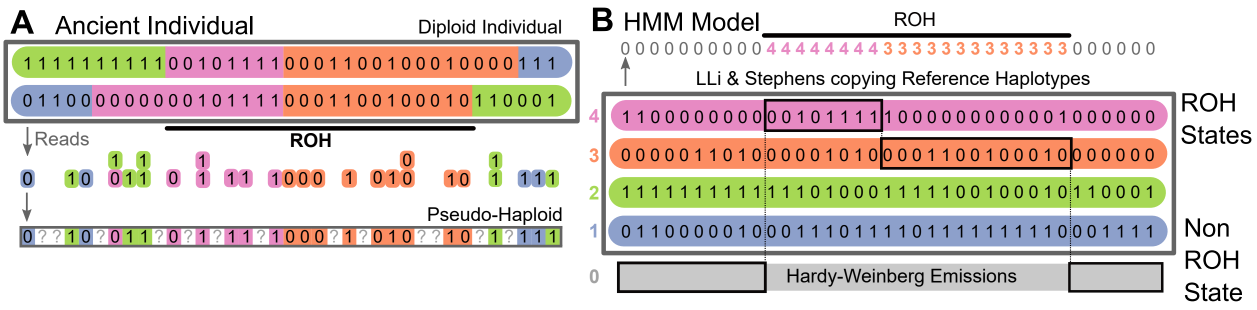


**Supplementary Figure 1: Detecting runs of homozygosity using a reference panel.** Panel A: Illustration of genotype data from a diploid individual. Sequencing reads mapping to a biallelic SNP produces counts of reads for each allele, from which in turn pseudo-haplotype genotypes, i.e. single reads per site, are sampled (at random). Panel B: Schematic of Method. A target individuals genotype data is modelled as being copied from a reference panel (colored) and one additional non-ROH state, where copying probabilities are given by Hardy-Weinberg proportions.

is homozygous for the allele of the reference haplotype that it copies from. The emission probabilities are specific to the exact kind of data that is analyzed, and can include various types of error models, which we discuss in more detail in Section 1.3.

In the Hardy-Weinberg state $0$, the probabilities of observing a diploid genotype reflect the probabilities of an underlying genotype in Hardy-Weinberg equilibrium, with probabilities of the alleles according to the underlying allele frequency in the reference panel at this locus. We note this state is identical to the non-ROH state used in a previously developed HMM to call ROH (Narasimhan et al., 2016).

#### 1.2 Infinitesimal Transition Rates

To define a hidden Markov model, one needs to specify the transition probabilities between the hidden states for each pair of successive loci $l$ and $l+1$. In our model, we do so by using an infinitesimal rate matrix $Q$ of dimension $(n+1)\times(n+1)$, from which the transition probability matrix $A_{l\to l+1}$ can be obtained via exponentiation: $A_{l\to l+1} = \exp(Q\cdot r_l)$, where $r_l$ is the genetic distance between the respective loci.

Following Li & Stephens, the copying states $i = 1,\ldots,n$ are symmetric in our model. We can thus specify the infinitesimal rate matrix by three parameters: A single rate for the transition from the non-copying into a copying state $Q_{0j}$ for all $j > 0$, a single rate for leaving a copying state $Q_{j0}$ for all $j > 0$ and a third rate for transitioning from one copying state to another $\phi_{\text{ROH}} = Q_{jk}$ for all $j,k > 0, j \neq k$. The diagonal entries of the rate matrix $Q$ are determined by the rate matrix condition $Q_{ii} = -\sum_{j\neq i} Q_{ij}$.

We point out that in the limit of infinite jumping rates within ROH ($\phi_{\text{ROH}} \to \infty$), our model converges to the full model of Narasimhan et al. (2016), as the probabilities of being in one of the allelic states (the sum of probabilities of copying from all reference haplotypes that have this allelic state) will then reflect its frequency, as in this limit jumps occur between any two consecutive markers.

#### 1.3 Emission Probabilities

In our model, the emission probabilities that specify the probability of observing the data at locus $l$ given some hidden state $i$, $e_i(y_l)$ depend on the type of data. We implemented two emission models: diploid genotype data and pseudo-haploid genotype data, with all three of them incorporating a model for genotype error. Throughout, we always disregard markers with missing data by removing them both from the reference panel as well as the target and adjusting the transition rates accordingly.

We implemented the emission model for diploid genotypes as follows. In the non-ROH state (i=0), the Hardy-Weinberg emission probabilities for the genotypes are $(1-p_l)^2$, $2p_l(1-p_l)$, and $p_l^2$, for observing homozygosity for the ancestral allele, heterozygosity, and homozygosity for the derived allele, respectively, where $p_l$ is the frequency of the derived allele in the reference panel at locus $l$. For the ROH-states ($i = 1, \ldots, n$), the genotype probabilities are 1 to be homozygous for the allelic type of the source haplotype in the reference panel, and 0 for the two other possible diploid genotypes. We extend these genotype probabilities to model possibly erroneous genotypes by assuming that with probability $\epsilon$ a genotype is flipped to one of the two other genotypes at random. This simplified error model has the advantage of having only a single parameter while broadly modeling a wide range of possible errors, including genotyping error in the reference as well as in the target, or new mutations that are private to the target individual. We note that for ancient DNA data, where genotyping error rates (including errors due to contamination) are typically on the order of $10^{-2} - 10^{-3}$ (Racimo et al., 2016), the genotyping error rate will be the main driver of $\epsilon$, as for modern human populations the reference panel is almost always separated no more than $10^5$ generations from the target. The per base-pair mutation rate is on the order of $10^{-8}$ per generation, which results in an upper bound for the substitution rate of order $10^{-3}$.

The second emission model we implemented is for pseudo-haploid genotype data, a common data type for human ancient DNA. For the copying states ($i = 1, \ldots, n$), the allele on haplotype $i$ is emitted with probability $1 - \epsilon$, and the alternative allele is emitted with probability $\epsilon$. For the non-ROH state (i=0), the emission probabilities model sampling one read from an underlying genotype in Hardy-Weinberg equilibrium under the allele frequencies in the reference panel: A derived pseudo-haploid marker is observed with probability $p_l$, and an ancestral marker with probability $1 - p_l$. To account for errors, with probability $\epsilon$ the observed read actually reflects the opposite allelic state. As in the case of diploid genotypes, this error rate $\epsilon$ models both the disagreement rate due to new mutations occurring on the genealogical lineage between the reference haplotype and the target, as well as the rate of genotyping errors.

We note that extensions for more complex error models that include position and context-specific effects or leverage base quality scores from the sequencing, and models for other kind of data could be incorporated by adjusting the emission probabilities linking the unobserved diploid genotypes to the data. Importantly, such extensions can be naturally modelled using a genotype likelihood framework that describes the likelihood of the observed data under each of the three possible latent diploid genotype states.

We implemented a third emission model for read count data based on genotype likelihoods. Here, the data for a specific locus consists of $n$ reads, with $k$ of them mapping to the derived allele and $n-k$ to the reference allele. Given the underlying genotype, modelled probabilistically as in the diploid genotype case described above, we add a second layer that describes the sampling of the $n$ reads. We use a binomial likelihood model, where the probability of observing $k$ out $n$ marker to be derived is binomial with probability $p=0$, $p=0.5$, and $p=1$ given the heterozygous ancestral, homozygous, and heterozygous derived genotype, respectively. We add two levels of error: One at the read level, where each read is flipped to the opposite allele with probability $\epsilon$, which can be absorbed into the binomial probabilities. We add an additional level of error at the genotype level, corresponding to the error model of erroneous diploid genotypes described above, where a diploid genotype is flipped to one of the other two possibilities with probability $\epsilon_{ref}$. This is to account for rare errors in the reference panel that would induce mismatches between the target individual's genotype. Applying the Binomial read count model to real data would require extensive testing of this likelihood model, which we leave for future work, as the assumption of ancient DNA data being modelled well by a Binomial likelihood of read counts is likely often violated. Importantly, potential biases could depend on the type of data (e.g. whole genome sequencing or 1240K enrichment data), which could introduce unwanted batch effects in ROH analysis.

#### 1.4 Posterior Decoding

We use standard Hidden Markov model algorithms to calculate the posterior probability $P(\pi_l = i|y)$ of the hidden state $i$ at locus $l$ observing the data $y_1, \dots y_L$ (Durbin et al., 1998). Specifically, we compute the forward probabilities,

$$f_i(l) := P(y_1, \dots, y_l, \pi_l = i) = e_i(y_l) \sum_k f_k(l-1) A_{ki}, \tag{1}$$

as well as the backward probabilities,

$$b_k(l) := P(y_{l+1}, \dots, y_L | \pi_l = k) = \sum_i A_{ki} e_i(x_{l+1}) b_i(l+1), \tag{2}$$

using dynamic programming, where $A$ denotes the transition matrix $A_{l-1 \to l}$. Together, these are combined to obtain the posterior:

$$P(\pi_l = i|y) = \frac{f_i(l) b_i(l)}{P(y)}, \tag{3}$$

where $P(y)$ denotes the full probability of the data, which can be computed as $P(y) = \sum_k f_k(L)$.

To complete the posterior decoding and thereby call ROH segments, we use posterior thresholding. We return consecutive regions where the posterior probability of the non-ROH state remains below the threshold $1-T$, or equivalently the sum of the posteriors of the copy states is above $T$. In Section 1.8 we describe the procedure for how we set the default value of $T$ for our implementation of the method.

#### 1.5 Computational Speedup

The run-time (and memory requirement) of the algorithm for the posterior decoding of the HMM scales linearly with the number of loci $L$ that are analyzed. In the naive implementation, the scaling with the number of hidden states $K$ (the number of reference haplotypes plus one here) is quadratic, since the full transition matrix has to be computed and each entry employed in Equation (1) and (2). Thus, the run-time of the naive implementation is $\mathcal{O}(LK^2)$.

However, as is standard for these models, we can reduce this run-time to linear in the number of hidden states, to $\mathcal{O}(LK)$, by using the symmetry of the copying states: For hidden state $i > 0$, the sum in Equation (1) can be split up into three parts (we suppress dependencies on $l-1$ here):

$$\sum_k f_k A_{ki} = \underbrace{f_0 A_{0i}}_{I} + \underbrace{\sum_{k>0} f_k A_{12}}_{II} + \underbrace{f_i(A_{ii} - A_{12})}_{III}, \tag{4}$$

where we used that $A_{ki} = A_{12}$ for all $k, i > 0$, which follows from the symmetry of the transition rate matrix $Q$. Similarly, for $k$=0 we get:

$$\sum_k f_k A_{k0} = \underbrace{f_0 A_{00}}_{I} + \underbrace{\sum_{k>0} f_k A_{10}}_{II}, \tag{5}$$

because $A_{k0} = A_{10}$ for all $k > 0$.

The quadratic dependence of the run-time on the number of states is caused by the sum in $II$ in Equation (4), and similarly in Equation (5). However, when updating the forward probabilities $f_i(l)$ for all states $i$, we only need to pre-compute $\sum_{k>0} f_k$ once for every locus. Doing so achieves the reduction to linear run-time. The backward algorithm can be modified analogously, with first splitting the sum in Equation (2) and then pre-computing $\sum_{i>0} A_{ki} e_i b_i$ only once when updating $b_k(l)$ for all states $k$.

#### 1.6 Efficient computation of the transition matrices

In the naive implementation of our algorithm, the infinitesimal rate matrix $Q$ has to be exponentiated at every locus $l$, which would be computationally costly (depending on the implementation scaling quadratic or worse with number of states). However, due to the speed-up described in Section 1.5, we only require a small subset of the entries of the full transition matrix, namely $A_{00}$, $A_{11}$, $A_{12}$, $A_{01}$ and $A_{10}$. We note that a truly symmetric model (such as the original Li & Stephens copying model) could be reduced even further into a single transition rate (the probability of staying in a copy state, Price et al., 2009). However, due to the additional non-ROH state here, one has to keep track of at least three rates, and these can be efficiently pre-compute as follows.

Using the symmetry of the copying states $1, \ldots, n$, we can collapse these states into state 1 and a single surrogate state for $2, \ldots, n$. We then only need to consider the states 0,1, and the surrogate state, thus arriving at a $3 \times 3$ transition rate matrix $\tilde{Q}$, where

$\tilde{Q}_{ij} = Q_{ij}$ for $i \leqslant 2$, $j < 2$ and $\tilde{Q}_{i2} = \sum_{j>1} Q_{ij} = (n-1)Q_{i2}$ for $i < 2$. Importantly, by exponentiation of $\tilde{Q}$ the three relevant entries of $A$ can be recovered by first computing $\tilde{A} = \exp(\tilde{Q})$ and then using $A_{ij} = \tilde{A}_{ij}$ for $i, j < 2$ and $A_{12} = \tilde{A}_{12}/(n-1)$.

To efficiently incorporate variable recombination distances between loci, we first diagonalize the common collapsed rate matrix: $\tilde{Q} = P^{-1}\tilde{D}P$. For each locus $l$, we can then exponentiate using $\exp(\tilde{Q} \cdot r) = P^{-1} \exp(\tilde{D} \cdot r_l)P$, which only requires exponentiation of a diagonal matrix, and recover the corresponding entries of $\tilde{A}$ and consequently $A$ required for calculating the full posterior. In Section 1.8 we describe the procedure for how we set the default rates of $Q$ for our implementation.

#### 1.7 Simulating genetic data with ROH

To test the performance of our method, we simulated genetic data with known ROH. We use this data below to carry out experiments where we down-sample to lower coverage and add genotyping errors to 1) help determining robust HMM parameters (Section 1.8) and to 2) test the performance (Section 1.8). First, we describe the method we used to generate these simulated datasets with known ROH.

We used a copying approach inspired by Ralph and Coop (2013) to generate ground-truth ROH block sharing data for testing methods. A synthetic mosaic individual without long ROH >1 cM is first generated by concatenating stretches of diploid genotypes in 0.25 cM tracts from randomly chosen individuals of the reference set. The intuition is that the probability of long ROH blocks (>1 cM) arising inadvertently is very low (as multiple ROH blocks would have to be concatenated), while still mostly retaining local LD structure typical for diploid human individuals. In our simulations, we used the positions of a widely used set of 1.24 million SNPs widely used for human ancient DNA studies (1240K capture technology for Fu et al., 2015), and we focused on chromosome 3, a human chromosome with a typical density of these sites per map unit (Morgan).

We then copied in five ROH blocks of a given length uniformly at random, enforcing that ROH blocks do not overlap by placing them at random in 5 evenly split up sectors of the chromosome. The copied-in stretch originates from one haplotype of the source population (chosen uniformly), and both alleles of the synthetic individual are set to the allele of the copied-in stretch. The source population for the simulations is then excluded from the reference panel. These synthetic mosaic individuals, with known diploid genotypes, serve as test cases for the method. These data were down-sampled and error added to it to simulate data of varying quality (Fig. 2C,D).

#### 1.8 Parameter Choice

The model has several parameters that have to be set when analyzing data. Here we describe how we set the parameters we used throughout our empirical analysis and our simulation experiments. We set the infinitesimal transition rates based on the typical tracts we are interested to find. Our target use case here is to detect ROH blocks that are of length 5 cM that occur once every 100 cM. Accordingly, we chose the infinitesimal rate parameters (per Morgan) as 1 (jump from non-ROH into ROH) and 20 (jump from

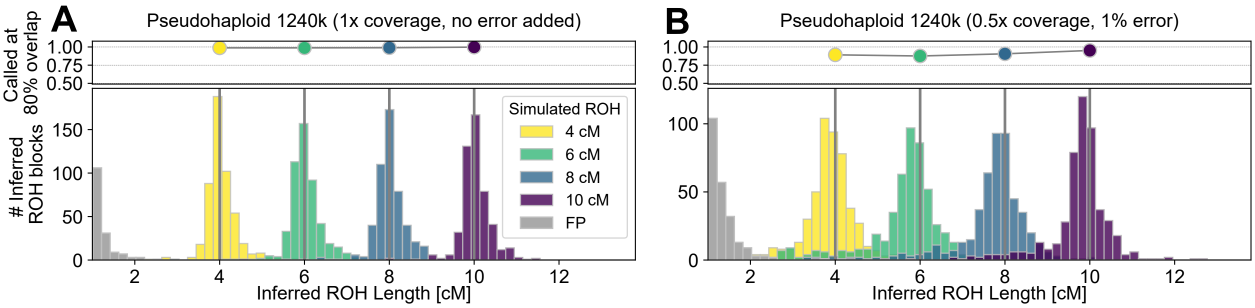


**Supplementary Figure 2: Detecting simulated ROH.** Panel A: We applied our method to simulated data with known ROH copied. We copied in ROH of either 4, 6, 8, and 10 cM length (5 of every length class into each of 100 simulated chromosomes, 1.7), and depict histograms of inferred ROH lengths (in color) as well as false positives (in gray). Panel B: Same as panel A, but a simulation with erroneous and missing data typical for lower quality ancient DNA data.

a ROH state into non-ROH). We fixed the transition rate between ROH states (i.e. the haplotype copying model switch rate) to 300 per Morgan, corresponding to an average copy tract length of ca. 0.3 cM. This value was chosen based on performance of ROH calling in pilot simulations and a likelihood profile of a Li & Stephens model of Tuscany haplotypes from all non-Tuscany Europeans in the 1000 Genomes dataset. We fix this set of parameters throughout our analysis.

#### 1.9 Choice of Posterior Threshold

To determine a robust posterior threshold, we ran simulation experiments with data typical for our use case, which is analysis of 1240K pseudo-haploid data with the full 1000 Genomes dataset set as a reference panel. As test cases, we simulated mosaics of chromosome 3 with pseudo-haploid data, i.e. one allele chosen at random, and then downsampled (at random) to 50% of all 1240K SNPs covered (and the rest set as missing data). We then flipped the allele with probability 0.01 to the other allele to simulate data with low quality. This is a representative use case for our method: As described below (Section 1.11) we apply our method to individuals in real datasets with more than 400,000 SNPs covered, for which estimated error rates are below 5% . We point out that error rates cover both sequencing error and contamination, and that not all contamination results in erroneous reads. The reason for choosing the cutoff based on low quality data is that we want the cutoff to be robust in these cases. We tradeoff maximum specificity for high quality data (where more aggressive cutoff settings would be possible) to allow our method being applicable to a wide range of use cases with default parameters.

Using the TSI (Tuscany, Italy) samples from the 1000 Genomes dataset, we simulated 100 replicates of mosaics of chromosome 3 for two scenarios: 1) with 4 cM ROH blocks copied in (to determine power and bias of inferred ROH length) 2) no blocks copied in as well (to assess false positives). We then ran the method using the 1000 Genomes dataset and only TSI individuals removed as reference panel, tested various posterior cutoffs, and monitored false positive rate, power, length bias, and standard deviation of the longest block overlapping the true ROH blocks, with blocks of length 4 cM as the test case. When analyzing 100 replicates with various posterior cutoffs, we found that a cutoff of 0.998 lead to a good performance in terms of the magnitude of bias for ROH, as well as standard deviation of inferred length of ROH (Table 1). As our overall goal is to call ROH with little bias and also with little variation in length, we chose this value of 0.998 as posterior cutoff in our implementation.

| Posterior Cutoff | Rep. | STD 4cM | FP ROH>1cM | FP ROH>2cM | Avg. Bias 4 cM [cM] | Frac. 80% of 4 cM called |
|---|---|---|---|---|---|---|
| 0.996 | 100 | 0.61 | 5.39 | 0.47 | 0.06 | 0.958 |
| 0.997 | 100 | 0.59 | 4.70 | 0.35 | 0.02 | 0.950 |
| 0.998 | 100 | 0.57 | 3.78 | 0.21 | -0.03 | 0.930 |
| 0.999 | 100 | 0.60 | 2.34 | 0.11 | -0.15 | 0.892 |

**Supplementary Table 1: Varying the posterior cutoff on various performance metrics.** We varied the posterior cutoff used for calling ROH, calculated several summary statistics when calling ROH for mosaic individuals (TSI). For each line, 100 replicates for chromosome 3 with five 4 cM ROH copied or no ROH copied in were simulated to calculate the performance statistics. False positive rates (FP) are calculated as the average number of falsely inferred blocks per replicate chromosome.

For applications on 1240K pseudo-haploid SNPs with at least 400,000 autosomal SNPs covered and using the 1000 Genomes data as the reference panel, this set of parameters can be readily applied, and we provide these parameters as the default settings in our software package that implements the method. For users who wish to apply our method to another set of SNPs, a different reference panel, or non-human data, we strongly recommend to repeat a similar strategy to find a suitable threshold in the respective scenario.

#### 1.10 Merging of Gaps between ROH

Motivated by the observation that the vast majority of false positive ROH are shorter than 2 cM (Fig. 2), we only record ROH blocks >2 cM. We observed that long ROH are sometimes broken up by spurious gaps (Fig. 3 and manual inspection of blocks where the length was substantially underestimated), as similarly seen in methods that call long IBD blocks between individuals (Browning and Browning, 2015). Such gaps may arise due to genotyping error, structural variation or very low SNP density. Following a standard procedure of IBD block calling (Ralph and Coop, 2013) and of genomic feature annotation with HMMs (Durbin et al., 1998), we decided to merge gaps, as experiments with lowering the posterior threshold or with decreasing the jump rate introduced a large surplus of additional false positives. To ensure that we do not merge two false positives (the false positive rate >2 cM is non-zero), we additionally require at least one of the merged blocks to be longer than 4 cM, and the gaps to be less than 0.5 cM in length. Fig. 3 shows that this procedure improves the performance substantially.

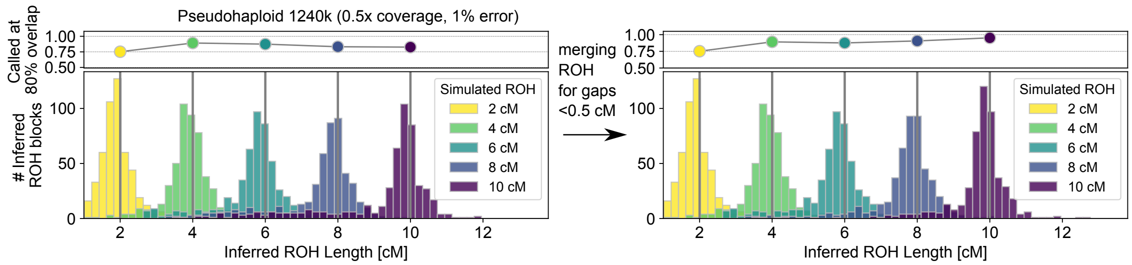


**Supplementary Figure 3: Improving power for long ROH blocks by merging gaps between ROH stretches** We depict the effect of merging ROH gaps for the "worst case" simulation scenario where we expect our method to have the least power to detect uninterrupted segments of ROH. Merging gaps <0.5 cM for between blocks where the longer block >4 cM markedly improves performance for long ROH blocks (>8 cM), without changing the distribution of shorter ROH blocks (4 cM).

#### 1.11 Performance on simulated data

To test its performance, we applied our implementation of the method with default parameters chosen as described in Section 1.8 to mosaic individuals with copied in ROH blocks as detailed in Section 1.7. When applying the method to pseudo-haploid data down-sampled to varying degree, we found that it has high power (>95%) to detect ROH blocks >4 cM while having simultaneously a low false positive rate (Fig. 4A) down to ca. 0.3× covered 1240K sites. Moreover, we find that, when first applying random genotype errors, the method can tolerate genotype error rates up to 5% (Fig. 4B).

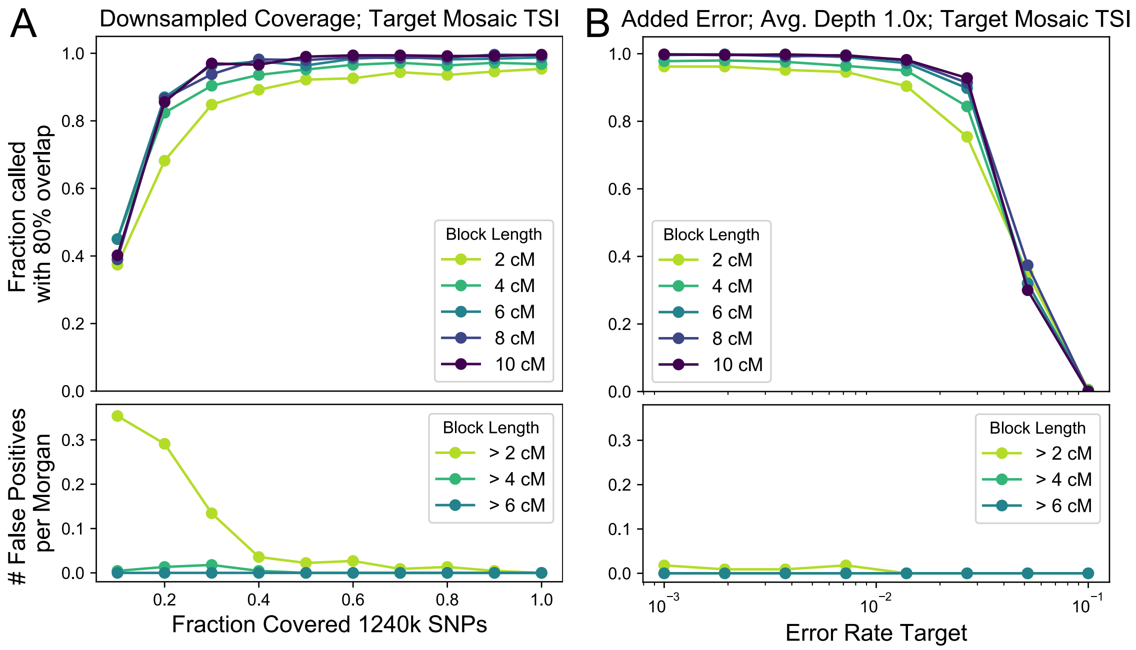


**Supplementary Figure 4: Performance of the method to detect ROH within mosaic individuals** We analyzed 100 individual chromosomes 3 which have been copied together as mosaics from 0.25 cM stretches from TSI individuals (Tuscany) of the 1000 genomes dataset (Section 1.7) on the 1240K sites. For each site, we then sampled one read from the diploid genotype at random, creating pseudo-haploid data. We further down-sampled to varying degrees (0.1-1.0×, Panel A), or introduced random genotype genotype errors at different rates (0.001-0.1) and applied the method with a copying error rate set to 1% (Panel B), using the 1000 genome data with the TSI haplotypes removed as reference panel (4794 haplotypes).

#### 1.12 Reference panels with varying genetic distance

To test the impact of different coalescence time distributions to the reference panel, we tested the method on simulated mosaic individuals from various global populations when using a reference panel consisting of European haplotypes. We note that under a simple model of a clean population split, the divergence time between the target and the reference population introduces a minimum boundary for coalescence times of the reference haplotype with the reference panel, similar to a temporal separation of an ancient target from the reference panel.

We tested how well the method works when using a European reference panel (with TSI removed, 792 out of 1,006 haplotypes remaining) for mosaic individuals generated from several target populations of the 1000 Genomes dataset. We tested four target populations, chosen to cover a wide range of population genetic distances. We tested with pseudo-haploid data on 1240K SNPs, picking one allele at random at each 1240K site (Tab. 2 and Fig. 5).

With divergence occurring tens of thousands of years ago, such as target for CHB (Han Chinese) with European reference haplotypes, 95.0% of copied-in blocks are identified with at least 80% overlap with the true ROH block. However, this behavior does not continue across all pairs of populations, we observe little power to infer ROH in mosaic individuals constructed from YRI haplotypes when using European haplotypes as reference. In this case, while some ROH blocks are still identified, only less than 10% of copied in ROH blocks are inferred with at least with 80% overlap.

| Target | Panel | Power at 80% overlap [4cM] | Bias in Length [4cM] | Standard Deviation Length [4cM] |
|---|---|---|---|---|
| **TSI** | **EUR*** | 0.986 | 0.151 | 0.46 |
| **CHB** | **EUR*** | 0.950 | 0.138 | 0.54 |
| **CLM** | **EUR*** | 0.882 | -0.10 | 0.69 |
| **YRI** | **EUR*** | 0.096 | -2.01 | 0.90 |

**Supplementary Table 2: Effect of varying distance from reference panel to target.** We tested the performance with mosaic individuals from Tuscany, Italy (TSI); Han Chinese from Beijing (CHB); Colombians from Medellin (CLM) and Yoruba from Ibadan (YRI), and tested the power to call ROH blocks of length 4 cM. As before, we define a successful inference when at least 80% of the original ROH block are inferred to be within a single inferred ROH. EUR*: European reference haplotypes with TSI (Tuscany) removed.

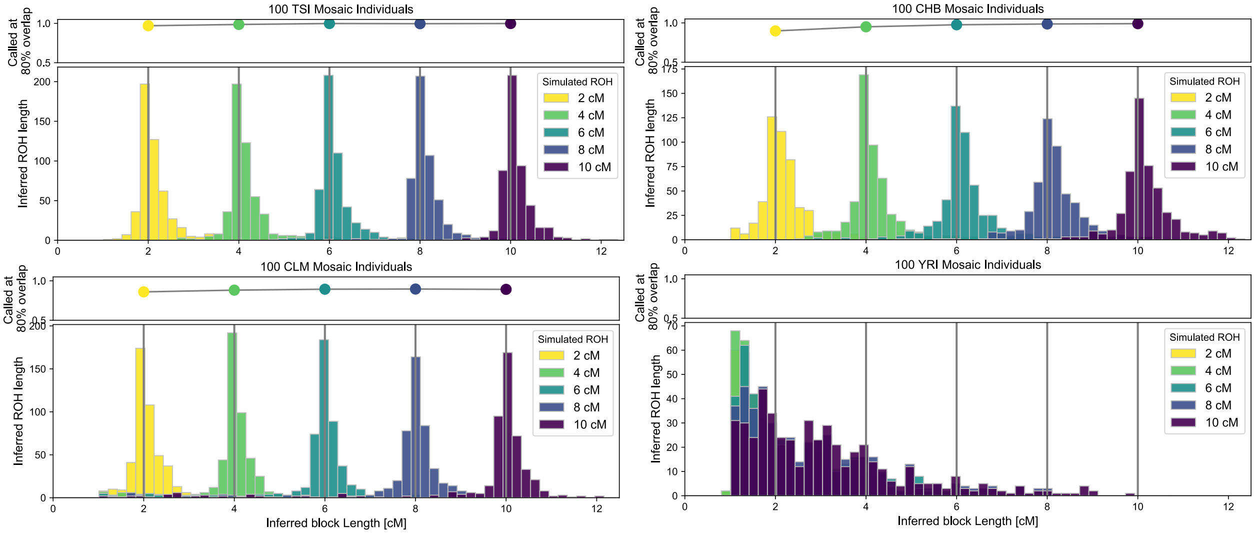


**Supplementary Figure 5: Effect of varying distance from reference panel to target.** We tested the performance using European reference haplotypes (without TSI haplotypes) for target individuals that were simulated as mosaics of haplotypes from Tuscany, Italy (TSI); Han Chinese from Beijing (CHB); Colombians from Medellin (CLM) and Yoruba from Ibadan (YRI).

#### 1.13 Performance on down-sampled Ust Ishim man

High-coverage ancient DNA data provides a useful test case to assess ROH inference. Here we analyzed a Western Siberian individual radio carbon dated to about 45,000 years before present, called "Ust Ishim man". His complete genome has been sequenced to remarkable depth (ca. 40×) from a femur bone (Fu et al., 2014), allowing for robust diploid genotype calls.

Importantly, high-coverage data allows one to call ROH with high reliability by simply identifying stretches that lack sites where many reads indicate heterozygosity (Fig. 6). Moreover, as "Ust Ishim man" is the oldest anatomically modern human sequenced to high coverage to date, it provides us an opportunity to examine an extreme case in terms of how much temporal distance from the reference panel our method can tolerate.

We analyzed read count data for the 1240K SNPs from Ust Ishim man (40× read depth on the target) - using the post-processed publicly available data from Marcus et al. (2020). We then down-sampled these reads to lower coverage (0.2-40×) at random. Furthermore, we created pseudo-haploid data for all SNPs covered (1,115,315 of the 1240K variants were covered) by choosing one read at random per site, and then created artificial data down-sampled to subsets (0.3-1.0× smaller) of the 1240K sites. We analyzed both read-count data and pseudo-haploid data and summed up all ROH blocks longer than a given threshold.

Our results show that we can consistently infer ROH blocks >4 cM when down-sampling to low coverage (0.5× mean depth) of the 1240K markers (Fig. 6). Importantly, even for pseudo-haploid data, which effectively only uses LD information as signal, inference seems to work reliably with as low as 0.3× coverage, with little observable bias for blocks >8 cM and a small false positive rate for blocks >4 cM (Fig. 6). We hypothesize that this is at least in part caused by the extension of a large number of shorter ROH that then get pushed beyond the 4 cM detection threshold.

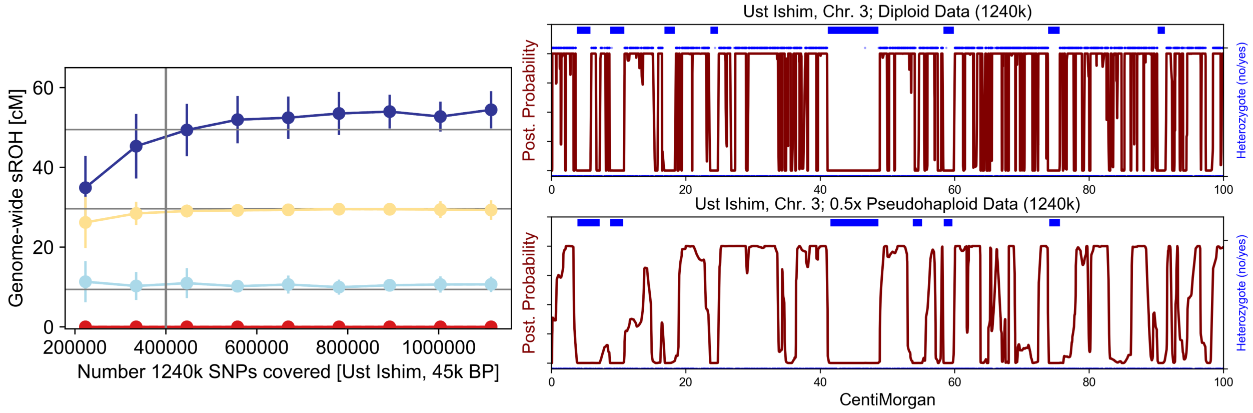


**Supplementary Figure 6: Properties of inferred ROH when downsampling from high coverage data on the Ust Ishim man.** Left: We down-sampled pseudohaploid data of Ust Ishim man to random subsets of 1240K SNPs to nine target coverages, and inferred ROH for each of 100 replicates. We depict mean and standard deviation of the inferred ROH in four length bins (4-8, 8-12, 12-20, and >20 cM). Right: Posterior and inferred ROH for a region of Chromosome 3, when using the full diploid genotype data (top) and pseudo-haploid data down-sampled to 0.5× coverage (bottom). We depict inferred ROH greater than 1 cM before gaps are merged as blue lines above the posterior. For the diploid genotype data, we indicate heterozygous genotypes (blue dots above the posterior) and homozygous genotypes (blue dots below the posterior). Long gaps of heterozygosity align well with the inferred ROH segments (blue lines).

#### 1.14 Performance on present-day populations

We applied our method to the Human Origins dataset of 1,941 present-day humans originating from 162 global populations genotyped at autosomal SNPs (Lazaridis et al., 2014). These SNPs constitute a subset of the 1240K enrichment targets (≈0.6 of ≈1.24 million SNPs). Because this dataset provides diploid genotype calls, we ran our method with the diploid mode and called ROH >4 cM in all 1,941 individuals, using 5,008 global haplotypes from the 1000 Genomes reference panel. We manually checked several called ROH, and confirmed that ROH calls correctly identify regions with almost no heterozygous markers.

To test the pseudo-haploid mode of our method on a global panel of variation, we used all HO individuals with at least one ROH longer than 12 cM identified (599 individuals) as a test set. In addition to the high quality diploid ROH calls, we ran the pseudo-haploid mode on these individuals, choosing one allele at random for each diploid genotype call (ca. 550,000 SNPs per individual, ranging from individuals with 537,000 to 556,000 called gentoypes). Our tests confirmed that the ROH calls from the haploid and the diploid data closely agree for the majority of individuals (Fig. S7), with a correlation between datasets of $r$ = 0.984 when comparing ROH >8 cM (Fig. 7). A notable exception are certain Sub Saharan populations, in particular South and East African hunter gatherers, for which a substantial fraction of long ROH are not identified in the haploid data (Tab. 7).

When investigating these African Hunter gatherers, we noticed that the typical pattern in the inference from pseudo-haploid data is many gaps dispersed throughout ROH identified in the diploid data (e.g. Fig. 8). This pattern mirrors the one we observed when analyzing mosaic targets created from Yoruba haplotypes using an European only reference panel (Section 1.7), pointing toward some haplotype segments not captured well by the reference panel. Indeed, it has been observed previously that hunter gatherer populations in Sub Saharan Africa possess deeply diverged ancestry (Schlebusch et al., 2012), which together with the fact that the African reference haplotypes from the the 1000 Genomes data only include a single population from Central, Southern and Eastern Africa (i.e. the Luhya), yields a plausible explanation for the limited power of a method based on copying of long haplotypes.

After removing Sub Saharan African populations from Central, South and Eastern Africa, the correlation increases to $r$= 0.997, and the average difference between the sum of ROH $> 8$ cM inferred from pseudo-haploid and diploid genotype data is $-0.53$ cM (the mean of the sum of ROH inferred from diploid data is 98.03 cM). Upon inspecting specific length categories, ROH calls from all length classes are highly correlated, ranging from $r$ = 0.925 for ROH 4-8 cM to $r$ = 0.988 for ROH longer than 20 cM (Fig. 9). No population other the Sub Saharan African exhibits a substantial bias, which provides evidence that the reference panel is suitable for all other groups in the HO panel.

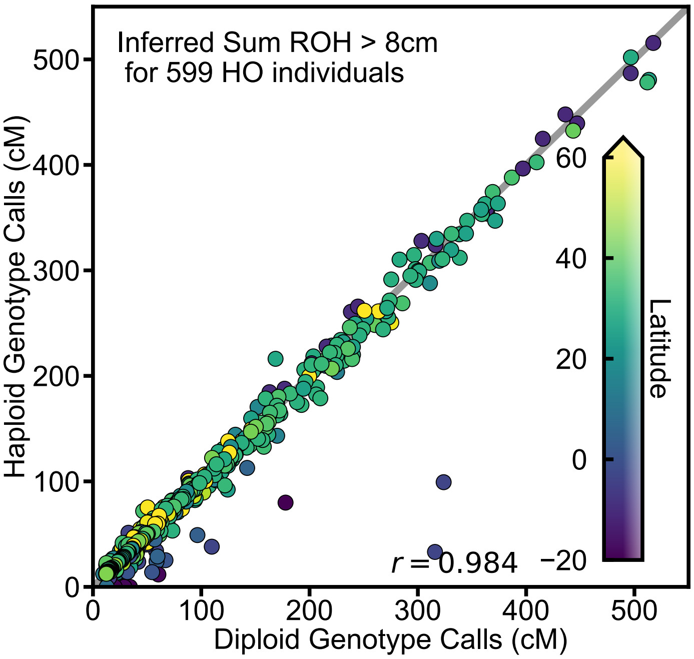


| HO Population | Failed | Total |
|---|---|---|
| Ju_hoan_North | 4 | 4 |
| Hadza | 3 | 3 |
| Mbuti | 3 | 4 |
| Khomani | 3 | 3 |
| Biaka | 2 | 3 |
| Ethiopian_Jew | 1 | 2 |
| Somali | 1 | 6 |

**Supplementary Figure 7 & Supplementary Table 3: Comparison of diploid and pseudo-haploid ROH calls for HO individuals.** Left: Comparison of ROH calls >8 cM for pseudo-haploid and diploid data for each HO individual with at least one ROH >12 cM (599 individuals). The scatter plot compares the total sum of all ROH blocks >8 cM. Right: Table summarizing individuals where more than 50% of sum ROH >8 cM are not called with pseudo-haploid data. These individuals correspond to the individuals that deviate substantially downwards from the diagonal line.

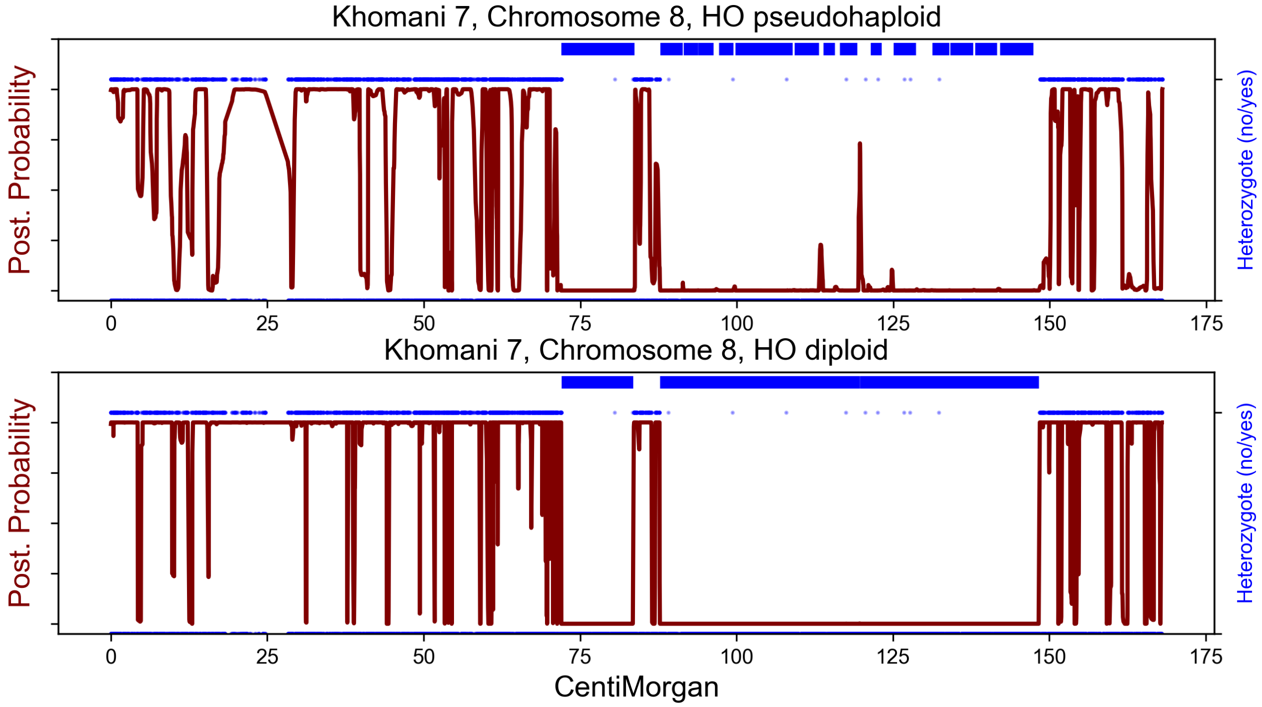


**Supplementary Figure 8: Comparison of diploid and pseudo-haploid ROH calls for a present-day Southern African Hunter gatherer individual.** We compare the ROH calls from pseudo-haploid data (top) and diploid genotype data (bottom) from a HO African hunter gatherer in the HO origin dataset (Khomani 7). We show chromosome 8, as this individual has two long ROH on this chromosome that can be identified with high confidence in diploid genotype calls (blue dots above posterior depict heterozygous sites). The diploid mode correctly identifies these regions, whereas the pseudo-haploid mode breaks them up.

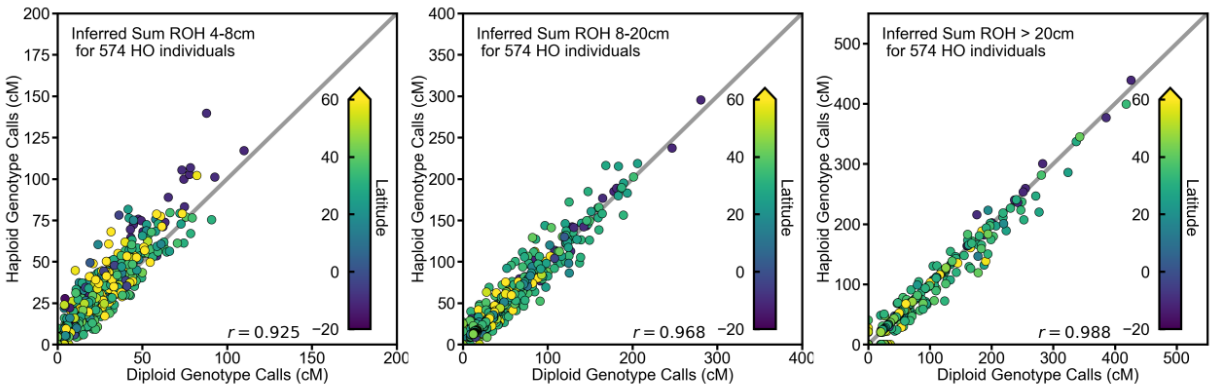


**Supplementary Figure 9: Comparison of diploid and pseudo-haploid ROH calls for HO individuals without Sub Saharan populations.** As in Fig. 7 we compare ROH calls for HO populations, with ROH calls from diploid genotype data (x-axis) compared to ROH calls from pseudo-haploid data (y-axis). Here we have removed the Sub Saharan populations from the panel, and show comparison for three length classes: 4-8 cM (left), 8-20 cM (middle) and >20 cM (right).

### 2 Comparison to existing Methods

Two programs are currently widely used to identify ROH from high quality present-day data (Ceballos et al., 2018). The software `PLINK` scans for windows of genotypes that lack heterozygous markers (Purcell et al., 2007). This simple but robust method uses diploid genotype calls. The second common method, `bcftools/ROH` (Narasimhan et al., 2016) uses a HMM with two hidden states, the non-ROH state emitting homozygotes and heterozygotes, and the ROH state emitting only homozygotes, with Hardy-Weinberg proportions according to the population allele frequency at each site. It can take genotype likelihood data as input, and therefore can in principle also operate on data where coverage is too low for accurate diploid genotype calls.

We compared the performance of these methods to our method on simulated data for the 1240K array with ROH blocks spiked in, generated using the procedure detailed in Section 1.7. We applied all three methods in two scenarios. First, we applied all three methods to diploid genotype data, typical for SNP array data from present-day individuals. We find that all three methods have excellent power and little bias when using diploid data (Supp. Table 4 and Supp. Fig. 10). However, we note that for some long ROH, `PLINK` (when using default settings) can break up long ROH, which we did not observe when using `bcftools/ROH` or `hapROH`.

The second scenario (see Supp. Fig. 11) is designed to test performance on typical ancient data for which diploid genotype calls are not possible. We created down-sampled mosaic individuals and compared the performance of our method to `bcftools/ROH`. In this comparison we did not include `PLINK`, as it can only operate on diploid genotype data and it has been previously shown to perform very poorly for down-sampled data, where the maximum likelihood genotype is used (Renaud et al., 2019).

To be able to apply `bcftools` in this scenario, we calculated genotype likelihoods from read count data: We assume that the probability of observing $k$ derived out of a total of $n$ reads for a singular SNP site given genotypes G=00, 01, or 11 is given by a binomial likelihood:

$$\Pr(\text{RC}|\text{G}) = \Pr(k, n|\text{G}) = \binom{k}{n} p^k (1-p)^{n-k} \tag{6}$$

where $p$ denotes the probability to observe a derived read ($p = 0, 0.5,$ and $1.0$ for genotypes 00,01, and 11, respectively). In these likelihood, we included a read error of $0.001$ (similar to the default setting of our method) by modifying the read count probabilities to $p = 0.001, 0.5, 0.999$. These likelihoods were normalized and encoded in SHRED-scale in the PL field in custom output .vcfs, as required for the input of `bcftools/ROH`.

We also tested the performance on read count data. To this end, we generated total reads per site according to a Poisson model. To mimic realistic read count distributions, which can be highly heterogeneous for 1240K data, we first calculated the ratio of total read depth per site and genome-wide read depth from a subset of 1240K data (Marcus et al., 2020) per site (calling these ratios $\lambda_i$). We then sampled at each SNP from a Poisson distribution with mean weighted by $\lambda_i$ times the genome-wide coverage we wish to simulate. We then sample derived reads according to a binomial model with $p$ =0, 0.5, or 1.0, depending on the underlying diploid genotype. In this simulation scenario (which we term "$\lambda$ read count"), the likelihood from Eq. (6) provides the exact likelihood, therefore we compare the performance of our method and `bcftools/ROH` under ideal conditions. We find that using default settings, `hapROH` works to much lower coverage than `bcftools`, in particular for ROH a few cM in length (Fig. 11). We note that the performance of `bcftools` can be improved by fine-tuning the transition parameters (which is currently implemented only in a experimental setting), however throughout all tested parameter ranges we could not find a setting where performance was comparably to `hapROH`, indicating that using linkage information provides a crucial advantage. We note that the read count model of `hapROH` is experimental (see above). We chose this data type for comparison, as one cannot apply `bcftools` to pseudo-haploid data, since no genotype likelihoods can be calculated for only a single read per site.

| Method | Power [4cM] | Bias [4cM] | SD [4cM] | FP Rate >1cM | FP Rate > 2cM |
|---|---|---|---|---|---|
| **hapROH** | 0.994 | -0.0022 | 0.160 | 0.00 | 0.00 |
| **bcftools/ROH** | 1.000 | 0.0844 | 0.155 | 0.17 | 0.00 |
| **PLINK** | 0.986 | 0.1000 | 0.229 | 0.15 | 0.00 |

**Supplementary Table 4: Comparison of the three methods on diploid genotype data (1240K SNPS)** We show performance metrics on 100 simulated Mosaic Individuals with five stretches of 4 cM, non-overlapping positions. Power is defined as ability to detect at least 80% overlap. False positive rate is calculated for 100 Chromosomes, with no ROH copied in (rate is per chromosome).

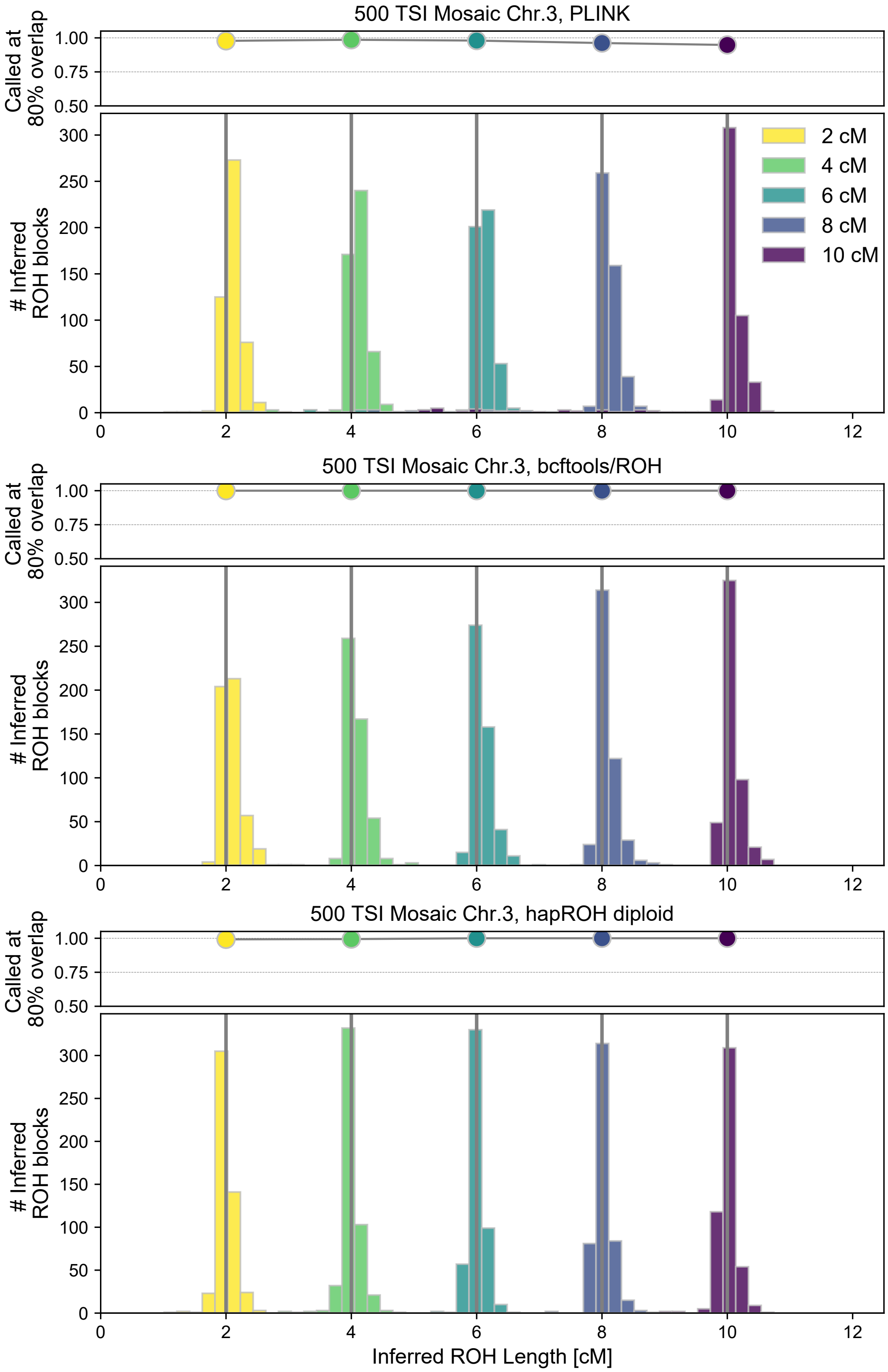


**Supplementary Figure 10: Comparison of the three methods to call ROH on diploid genotype data (1240K SNPS).** We show performance metrics on 100 simulated Mosaic Individuals with five stretches of 2, 4, 6, 8, or 10 cM ROH copied in. Power is defined as probability to detect an ROH that overlaps at least 80% of the simulated ROH.

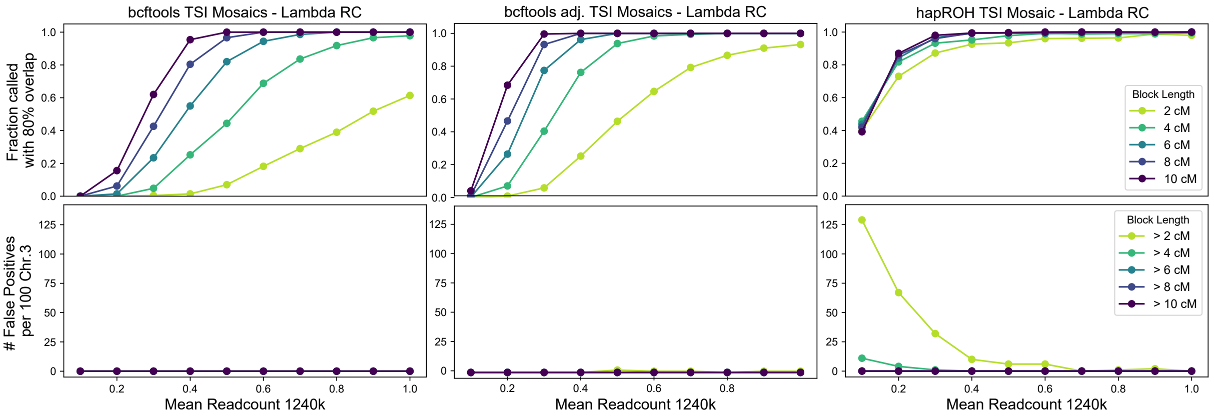


**Supplementary Figure 11: Comparison of `bcftools` and our method (`hapROH`) on low read count data (1240K SNPS).** We show performance metrics on 100 simulated Mosaic Individuals with five stretches of 2, 4, 6, 8, or 10 ROH copied in. We down-sampled these individuals to $(0.1, 0.2, \ldots, 1.0\times)$ coverage. We applied `bcftools` with standard parameters (left panel), adjusted parameters (middle panel) and our method (hapROH) in the experimental lambda read count mode (right panel).

### 3 Expected ROH for close relatives and small population sizes

Here, we calculate the expected number of ROH and the expected sum of lengths for all ROH falling into given length bins, using density functions $f(x)$, i.e. the values have to be integrated $\int_{l_1}^{l_2} f(x)dx$ to give expectations within bins $[l1, l2]$. Denoting $f(x)$ as the density of the expected number of blocks of length $x$, the integral $\int_{l_1}^{l_2} f(x)x\,dx$ yields the expected sum of lengths of the blocks within the length bin $[l1, l2]$.

Throughout, we measure block lengths in Morgans. The first key ingredient in the derivations is the expected number $b(x|t)$ of blocks of length $x$ caused by recombination $t$ generations ago on a chromosome of length $G$ Morgans. Assuming that recombination events are distributed according to a Poisson process with rate $t$, which is a good approximation for all but very close relatives (e.g. 1st and 2nd degree relatives, Caballero et al. (2019)), one gets:

$$b(x|t)\,dx = \underbrace{(G-x)(2t)^2\exp(-2tx)\,dx}_{(i)} + \underbrace{2(2t)\exp(-2tx)\,dx}_{(ii)},$$

where $(i)$ describes blocks in the interior of a chromosome and $(ii)$ from blocks delimited by one of the two chromosome boundaries. One straightforward way to derive this formula is by partitioning over all possible start sites for blocks of length $x$. We note that we ignored blocks extending over the whole chromosome, as in this work we are interested in shorter ROH. Combined with $\psi(t)$, the probability of coalescence $t$ generations ago, one can then express the expected number of ROH as:

$$f(x)\,dx = \int_0^\infty b(x|t)dx\,\psi(t)\,dt. \tag{7}$$

A detailed discussion of these formulas can be found in Ringbauer et al. (2017).

Here, we are interested in two scenarios. First, for the offspring of full $n$-th cousins, where the offspring is separated by $m = 2n+4$ meiosis, and four haplotypes are potential common ancestors:

$$\psi_n(t) = \frac{4}{2^m}\delta(m-t),$$

where $\delta(t)$ denotes the delta distribution. Substituting $\psi_n(t)$ into Eq. 7, we arrive at:

$$f_n(x)dx = \frac{4}{2^m}\big((G-x)m^2\exp(-xm) + 2m\exp(-xm)\big)\,dx. \tag{8}$$

Second, for constant (diploid) panmictic populations with $N$ haploids (often denoted as twice the effective number of diploid individuals $N = 2N_e$):

$$\psi_N(t) = \exp\left(-\frac{t}{N}\right)\frac{1}{N}.$$

Applying the integral Eq. 7, we arrive at:

$$f_N(x)dx = \left(\frac{8(G-x)}{N}\frac{1}{(2x+\frac{1}{N})^3} + \frac{4}{N}\frac{1}{(2x+\frac{1}{N})^2}\right)dx.$$

This formula further simplifies to

$$f_N(x)dx = \frac{4N\,(1+2NG)}{(1+2Nx)^3}dx, \tag{9}$$

which has previously been reported in paragraph 3.1. of Carmi et al. (2014).

As outlined above, the density functions in Eq. (8) and Eq. (9) can be integrated over the interval $[l1, l2]$ to give the expected number of ROH or the expected sum of the length of all ROH falling within this interval. Here we used numerical approximations with a large number of bins (1000), which are sufficiently accurate for all practical purposes, but we note that these integrals can also be solved analytically.

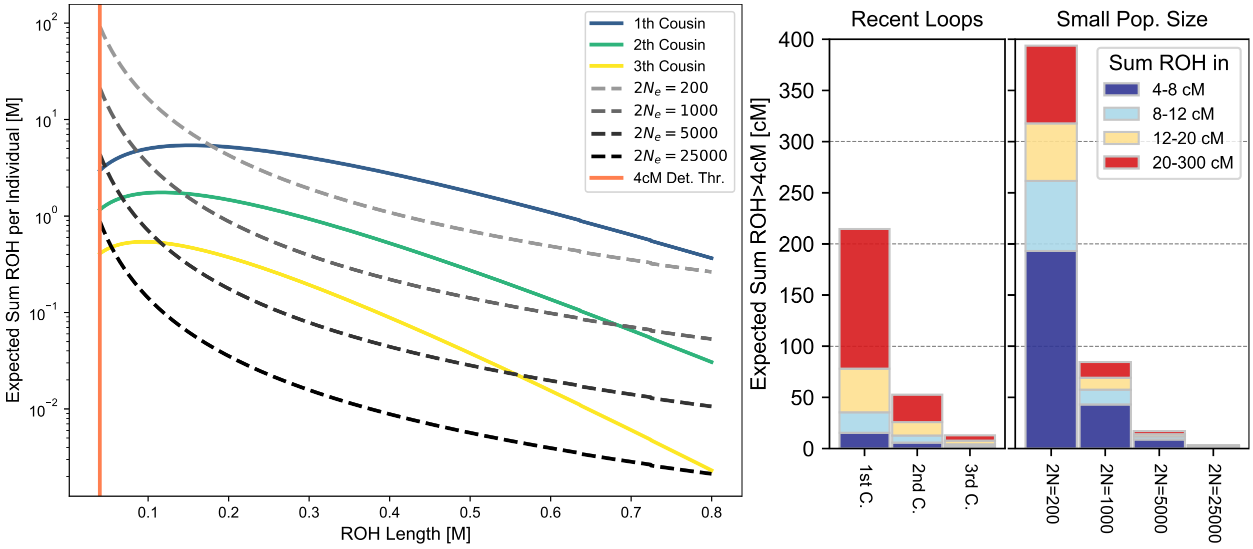


**Supplementary Figure 12: Density of expected sum of ROH per individual** We calculated the density of the sum of ROH per length bin for parents being full cousins of degree $1, 2, 3$ and effective population sizes $200, 1000, 5000$ by multiplying the expected number of ROH blocks $f(x)$ from Eq. (8) and Eq. (9) with the length $x$. Integrating $\int_{l_1}^{l_2} f(x)x\,dx$ would yield the expected sum of block lengths within the length bin $[l_1, l_2]$. Left: Expected densities. The vertical line depicts the detection cut-off we applied in our analysis of the ancient data. Right: Integral of expected densities over bins used in the empirical analysis (4-8, 8-12, 12-20, >20 cM).

We calculated the density of the expected sum of ROH blocks (Supp. Fig. 12). Our results show that offspring of parents that are close relatives, and thus have short circles in their pedigree, has most of its sum of ROH in the upper length category (20-300 cM). In contrast, loops resulting from low population sizes create bottom heavy distributions, where a substantial amount of the sum of ROH >4 cM is concentrated in ROH near the detection threshold.

For the case of a constant population size, one can use the integrand of Eq. (7) to partition the full expected sum of ROH (density) into contributions from each time point, where the integrand can be interpreted as a density of expectations per time interval. Figure 13 depicts this density for the case $2N = 500$. We observe that due to the exponential clock provided by recombination (the $\exp(-2tx)$ term), most of the total ROH of intermediate length classes (depicted for 4, 8, 12, and 20 cM) originate from recent timescales. For short blocks of length 4 cM, there is a substantial contribution from up to 100 generations ago (with less than $1\%$ expected contribution from beyond that), whereas for longer blocks 12 cM substantial contributions from only up to 20 generations ago arise. Also note that blocks from certain generations are more likely to result in the required length, therefore initially the density goes up when going back in time. All these qualitative patterns will in fact hold for all but extreme scenarios of demography (producing exponentially growing coalescent rates back in time) that would counteract the exponential recombination clock, analogously to IBD blocks between individuals (Ringbauer et al., 2017).

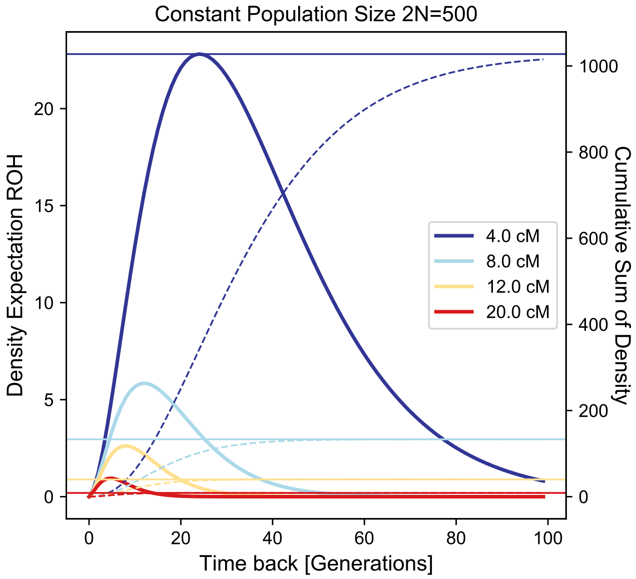


**Supplementary Figure 13: Timescales of ROH sharing for constant population size**. The density of expected ROH blocks with respect to time for a constant panmictic population of size $2N = 500$, depicted for block lengths 4, 8, 12, and 20 cM. We show the cumulative sum of these expectations (dotted curves, y axis labels right axis) when summed over time and also the analytical integral over all times from Eq. (9) (horizontal lines). Calculations were done with chromosome lengths of the human autosomes, and then summing the contribution from each chromosome.

To validate the analytical formulas, we simulated ROH in panmictic populations of sizes $2N = 500, 1,000, 2,000,$ and $4,000$, using the software `msprime` (Kelleher et al., 2016). We simulate ROH on all autosomes, each chromosome in a separate run. For chromosome lengths, we used the map difference between the first and last 1240K SNP on each autosome, both in analytical formulas and the simulations. We defined ROH as regions delimited by two recombination events in the full ARG when simulating two haplotypes. When binning the ROH values into length bins as used in the main paper (4-8, 8-12, 12-20, and >20 cM), the average values over replicate individuals within this bins agree closely with the average of the simulated values (Fig. 14).

Similarly, we simulated the offspring of cousins of various degrees, which are described in detail in Section 4. Again, the simulated values (when averaged over a large

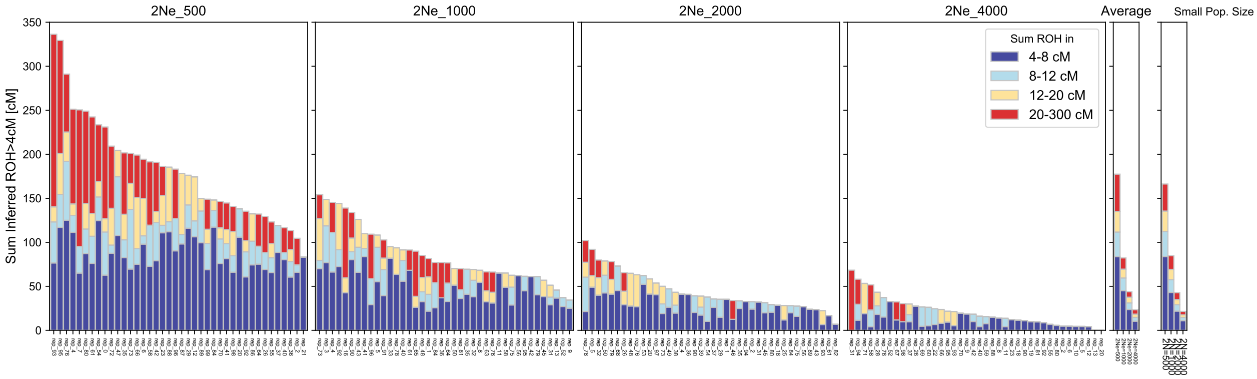


**Supplementary Figure 14: Simulated ROH for four population sizes**. We visualize the simulated ROH distribution on all autosomes for $2N_e = 500, 1,000, 2,000, 4,000$. Each bar represents ROH of one simulated individual (40 independent replicates per population size). The panel denoted "Average" gives the empirical average for each of these groups, and the panel denoted "Small Pop. Size" gives the analytical average calculated from formula, Equation (9).

number of replicates) and analytical values are in close agreement (Fig. 15), validating the formulas derived here.

### 4 Simulations of individual ROH using a detailed recombination model

To gain insight into the length distribution of ROH blocks for a given degree of parental relatedness, one can calculate expected numbers of blocks falling into certain length classes (see Section 3). However, these calculations do not yield insight into the variance of the distribution, and also rely on the assumption that recombination can be modelled as a Poisson process (when genomic distances are measured in Morgan) and do not incorporate the biological process of recombination interference (i.e. recombination events are less clustered than expected) as well as sex-specific recombination maps. For distant relatives beyond second degree these model violations have only minimal impact (Caballero et al., 2019), but this leaves the possibility that this process can significantly influence ROH patterns when an individual's parents are close relatives.

For these reasons, we utilized a recently developed method to simulate shared blocks of genome between close relatives (Caballero et al., 2019) to gain insight into the length distribution of ROH blocks. Importantly, this simulation engine can incorporate both sex-specific recombination maps as well as recombination interference.

We simulated $1000$ full individuals each, using the sex-specific genomic map of Bhérer et al. (2017), and simulating all autosomes. We then cluster individual ROH into bins of various lengths, as done in the empirical analysis. Our simulations demonstrated that ROH sharing among 1st cousin offspring of otherwise outbred individuals ranges from ca. 50-500 cM for $1000$ simulated first-cousin-offspring, with a mean expected value of 1/16th of the autosomal genome, 225 cM. Our results also show that the rate of ROH longer than 20 cM drops quickly with the increasing degree of parental relatedness (Supp. Fig. 15). When simulating $1000$ replicates for each parental relatedness scenario, for offspring of parents who are (full) first cousins, 97.7% have at least one ROH longer than 20 cM (95% binomial CI: 96.6-98.5%), for second cousins, this fraction drops to 57.1% (53.9-60.2%), and for offspring of fifth cousin it is only 0.2% (0.02-0.72%) (Supp. Table 5).

Based on these simulations, we mark individuals as begin potential offspring of very closely related parents if the sum of ROH $>$20 cM exceeds 50 cM. Ca. 88% of all first cousins offspring and 20% of all second cousin offspring pass this threshold. However less than 1% of third and less than 0.1% of offspring of parents fourth or further, fall above the threshold. Even if power to detect long ROH in this length class would be only 50% (a value far below the power estimates from our simulation and down-sampling experiments), one would still expect to detect ca. 60% of all first cousins in the dataset (Supp. Fig. 16).

| Parents being... | Replicates | 4-8 cM | 8-12 cM | 12-20 cM | >20 cM |
|---|---|---|---|---|---|
| 1st_cousin | 1000 | 913 | 848 | 939 | 977 |
| 2nd_cousin | 1000 | 625 | 476 | 557 | 571 |
| 3rd_cousin | 1000 | 289 | 172 | 227 | 142 |
| 4th_cousin | 1000 | 107 | 68 | 65 | 23 |
| 5th_cousin | 1000 | 23 | 12 | 20 | 2 |

| Parents being | sum(ROH >20 cM) >50 | sum(ROH >20 cM) >100 |
|---|---|---|
| 1st_cousin | 883 | 602 |
| 2nd_cousin | 201 | 27 |
| 3rd_cousin | 8 | 0 |
| 4th_cousin | 0 | 0 |
| 5th_cousin | 1 | 0 |

**Supplementary Table 5: Number of simulated individuals with ROH within a given length class.** We simulated 1000 individuals for each class of parental relatedness. The upper table gives the number of individuals which have at least one ROH in a given length class on any of their autosomes (each ROH length class is one column), the lower table the number of individuals with at least a certain amount of ROH longer than $> 20$ (when summing over all such blocks).

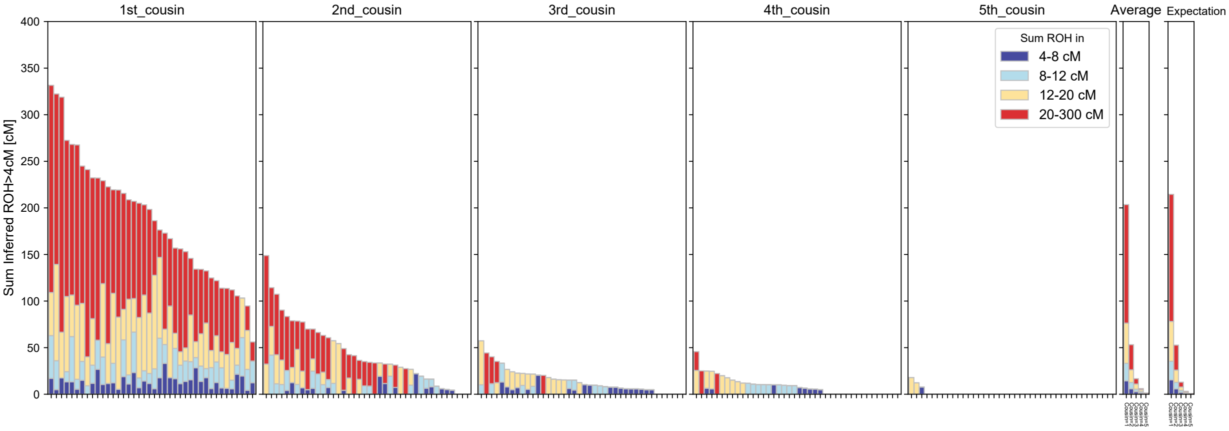


**Supplementary Figure 15: Simulated ROH in offspring of cousins of various degrees of relatedness.** We used the software `pedsim` to simulate ROH given various degrees of parental relatedness on all autosomes. The software modelled both recombination interference and sex-specific genetic maps. Each bar visualizes one individual and we color-code the sum of ROH in distinct length classes. For each parental degree of relatedness (1st to 5th full cousins, i.e. relatedness via both a male and female shared ancestor) we show 40 replicates. The panel denoted "Average" shows the empirical average for each ROH length bin. The panel denoted "Expectation" shows the corresponding expectation calculated from formula Eq. (8).

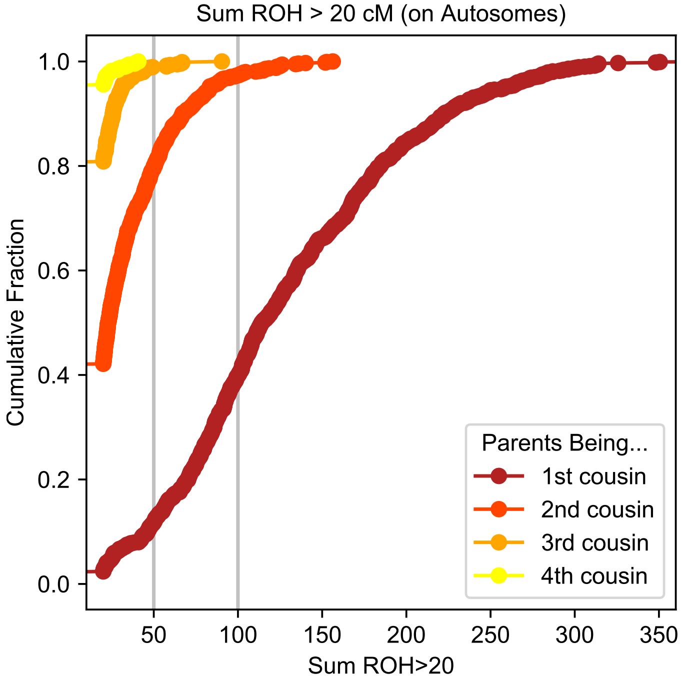


**Supplementary Figure 16: Distribution of ROH > 20cM in offspring of cousins of various degrees of relatedness** We show the cumulative distribution of the sum ROH> 20cM for 1000 simulated individuals each for 1st to 4th degree. The gray vertical bars depict 50 and 100 cM, which are used as threshold in the main manuscript.

### 5 Properties of ancient data set

As outlined in the methods, the bulk of our global ancient DNA dataset originates from a curated dataset of published ancient DNA (released on March 1, 2020, v42), available via https://reich.hms.harvard.edu. This release provides ancient DNA data in pseudo-haploid format with genotypes for the 1240K SNP set. It also contains individuals with whole genome sequenced data available, which had been down-sampled to this set of over a million SNPs. Here, we visualize three key statistics of this data set, as reported in the meta-file: 1) Age Distribution 2) Average Coverage and 3) Estimated autosomal contamination, available for males based on hemizygous X chromosomes (Fig. 17).

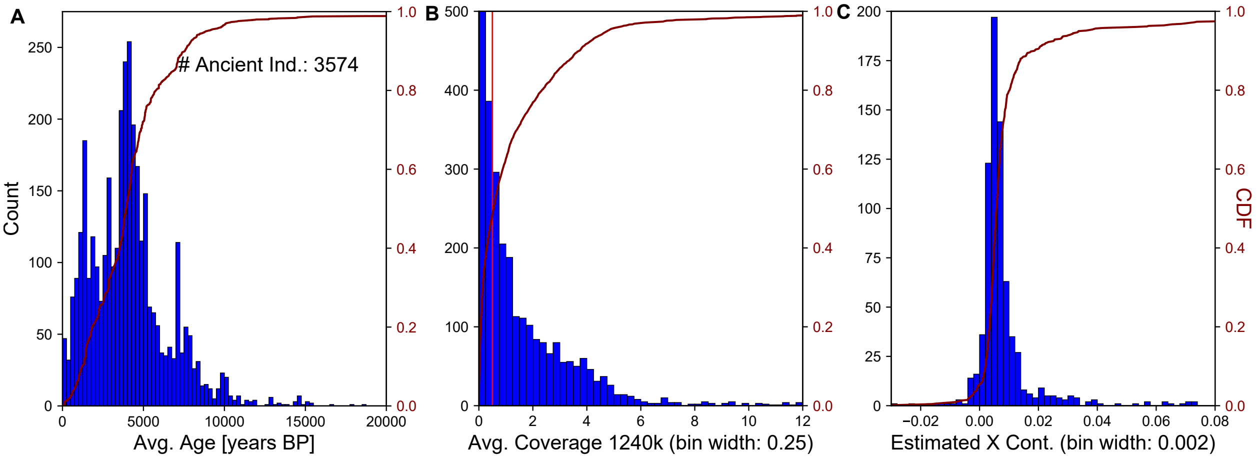


**Supplementary Figure 17: Data Details of ancient Individuals.** We depict key properties of publicly available ancient individuals downloaded from https://reich.hms.harvard.edu (v42). For each ancient individual we kept the record with the highest coverage (several have been genotyped multiple times). We depict histogram to visualize distribution of reported data properties of ancient individuals. Panel A: Age of each individual (mean of radio carbon dates where available, mean of context dates otherwise). Panel B: Mean Coverage on autosomal SNPs (1240K polymorphisms). Panel C: Mean reported error estimates (X contamination estimates ANGSD, MOM point estimator, which can be negative due to estimation uncertainty).

Richard Durbin, Sean R Eddy, Anders Krogh, and Graeme Mitchison. *Biological Sequence Analysis: Probabilistic Models of Proteins and Nucleic Acids*. Cambridge University Press, 1998.

Qiaomei Fu, Heng Li, Priya Moorjani, Flora Jay, Sergey M Slepchenko, Aleksei A Bondarev, Philip LF Johnson, Ayinuer Aximu-Petri, Kay Prüfer, Cesare de Filippo, et al. Genome sequence of a 45,000-year-old modern human from western Siberia. *Nature*, 514(7523):445–449, 2014.

Qiaomei Fu, Mateja Hajdinjak, Oana Teodora Moldovan, Silviu Constantin, Swapan Mallick, Pontus Skoglund, Nick Patterson, Nadin Rohland, Iosif Lazaridis, Birgit Nickel, et al. An early modern human from Romania with a recent Neanderthal ancestor. *Nature*, 524(7564):216, 2015.

Jerome Kelleher, Alison M Etheridge, and Gilean McVean. Efficient coalescent simulation and genealogical analysis for large sample sizes. *PLoS Computational Biology*, 12(5), 2016.

Iosif Lazaridis, Nick Patterson, Alissa Mittnik, Gabriel Renaud, Swapan Mallick, Karola Kirsanow, Peter H Sudmant, Joshua G Schraiber, Sergi Castellano, Mark Lipson, et al. Ancient human genomes suggest three ancestral populations for present-day Europeans. *Nature*, 513(7518):409–413, 2014.

Joseph H Marcus, Cosimo Posth, Harald Ringbauer, Luca Lai, Robin Skeates, Carlo Sidore, Jessica Beckett, Anja Furtwängler, Anna Olivieri, Charleston WK Chiang, et al. Genetic history from the Middle Neolithic to present on the Mediterranean island of Sardinia. *Nature Communications*, 11(1):1–14, 2020.

Vagheesh Narasimhan, Petr Danecek, Aylwyn Scally, Yali Xue, Chris Tyler-Smith, and Richard Durbin. BCFtools/RoH: a hidden Markov model approach for detecting autozygosity from next-generation sequencing data. *Bioinformatics*, 32(11):1749–1751, 2016.

Alkes L Price, Arti Tandon, Nick Patterson, Kathleen C Barnes, Nicholas Rafaels, Ingo Ruczinski, Terri H Beaty, Rasika Mathias, David Reich, and Simon Myers. Sensitive detection of chromosomal segments of distinct ancestry in admixed populations. *PLoS Genetics*, 5(6), 2009.

Shaun Purcell, Benjamin Neale, Kathe Todd-Brown, Lori Thomas, Manuel AR Ferreira, David Bender, Julian Maller, Pamela Sklar, Paul IW De Bakker, Mark J Daly, et al. PLINK: a tool set for whole-genome association and population-based linkage analyses. *American Journal of Human Genetics*, 81(3):559–575, 2007.

Fernando Racimo, Gabriel Renaud, and Montgomery Slatkin. Joint estimation of contamination, error and demography for nuclear dna from ancient humans. *PLoS Genetics*, 12 (4), 2016.

Peter Ralph and Graham Coop. The geography of recent genetic ancestry across europe. *PLoS Biology*, 11(5):e1001555, 2013.

Gabriel Renaud, Kristian Hanghøj, Thorfinn Sand Korneliussen, Eske Willerslev, and Ludovic Orlando. Joint estimates of heterozygosity and runs of homozygosity for modern and ancient samples. *Genetics*, pages genetics–302057, 2019.

Harald Ringbauer, Graham Coop, and Nicholas H Barton. Inferring recent demography from isolation by distance of long shared sequence blocks. *Genetics*, 205(3):1335–1351, 2017.

Carina M Schlebusch, Pontus Skoglund, Per Sjödin, Lucie M Gattepaille, Dena Hernandez, Flora Jay, Sen Li, Michael De Jongh, Andrew Singleton, Michael GB Blum, et al. Genomic variation in seven Khoe-San groups reveals adaptation and complex African history. *Science*, 338(6105):374–379, 2012.
