## Supplementary figures and images for "Human Parental Relatedness through Time - Detecting Runs of Homozygosity in Ancient DNA"

### Aegan.pdf

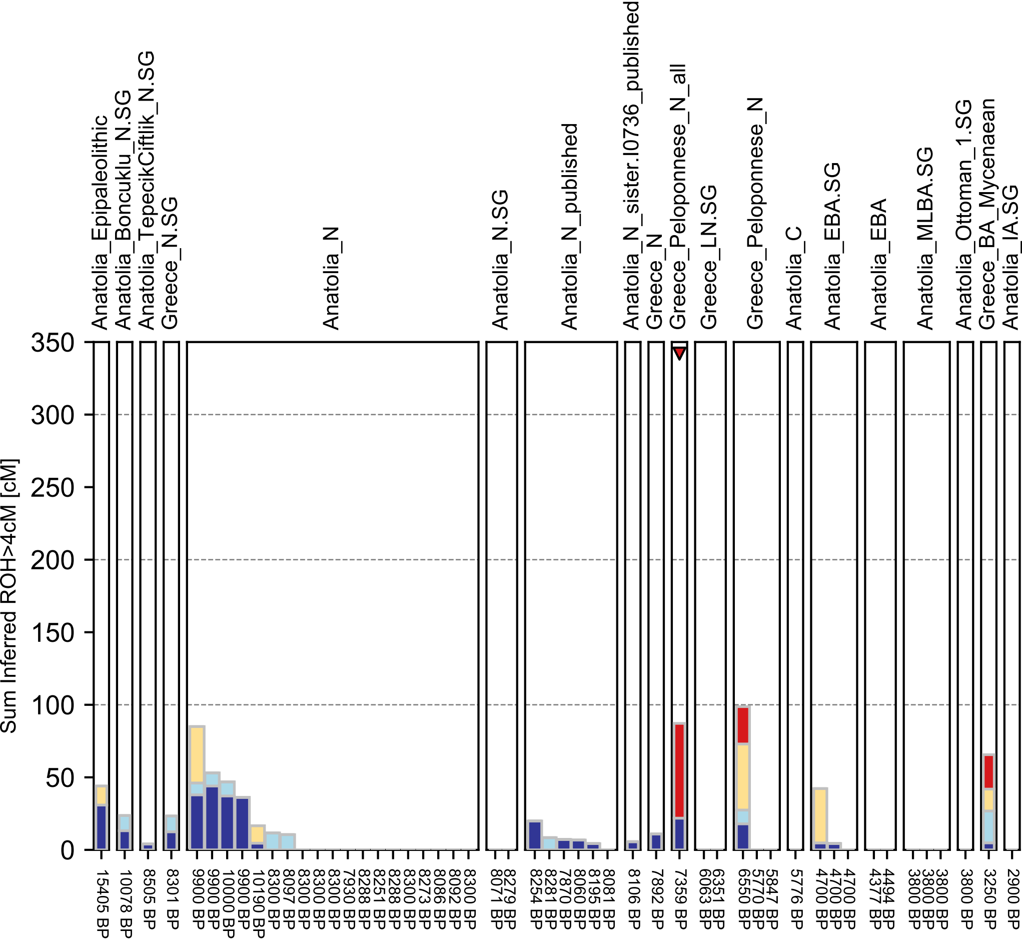

### Andean.pdf

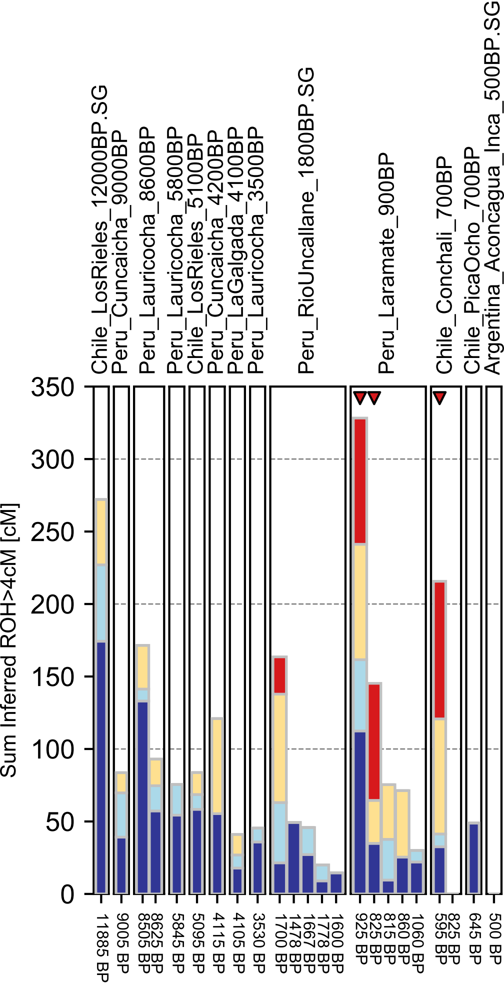

### Atlantic Coast.pdf

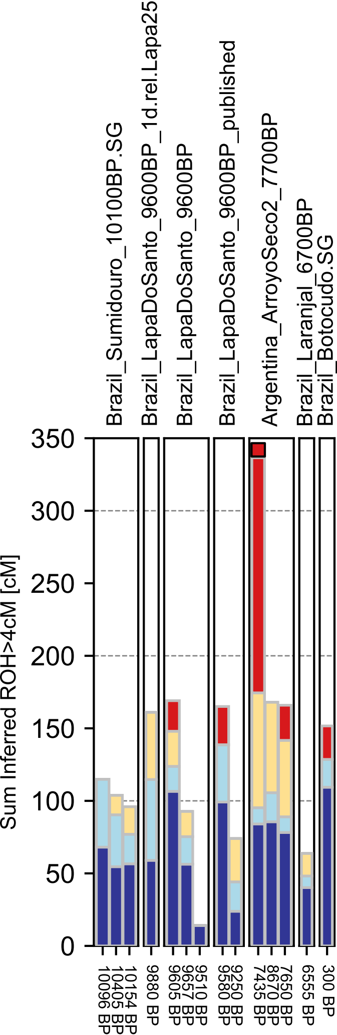

### Balkans.pdf

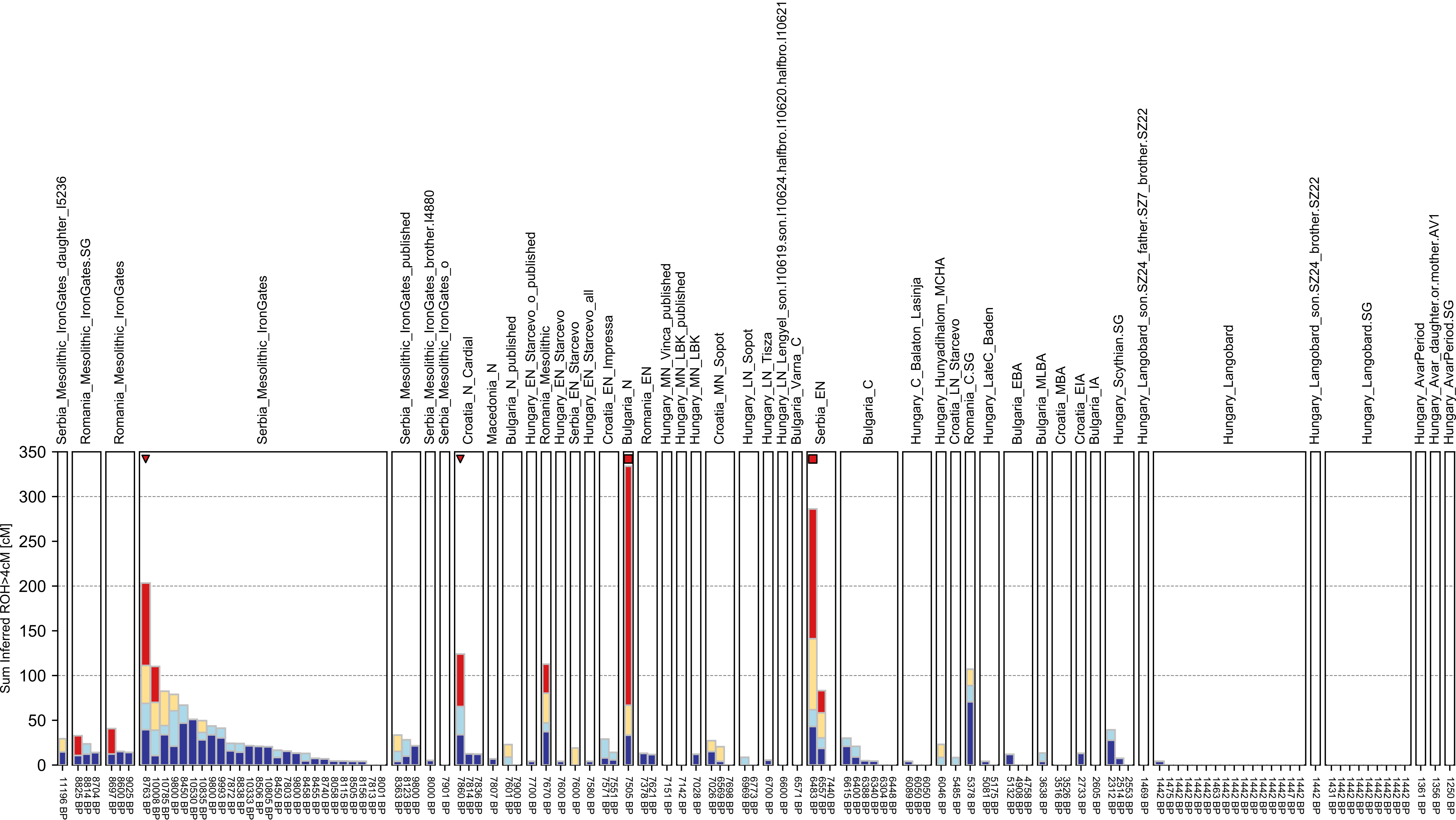

### Baltic Sea.pdf

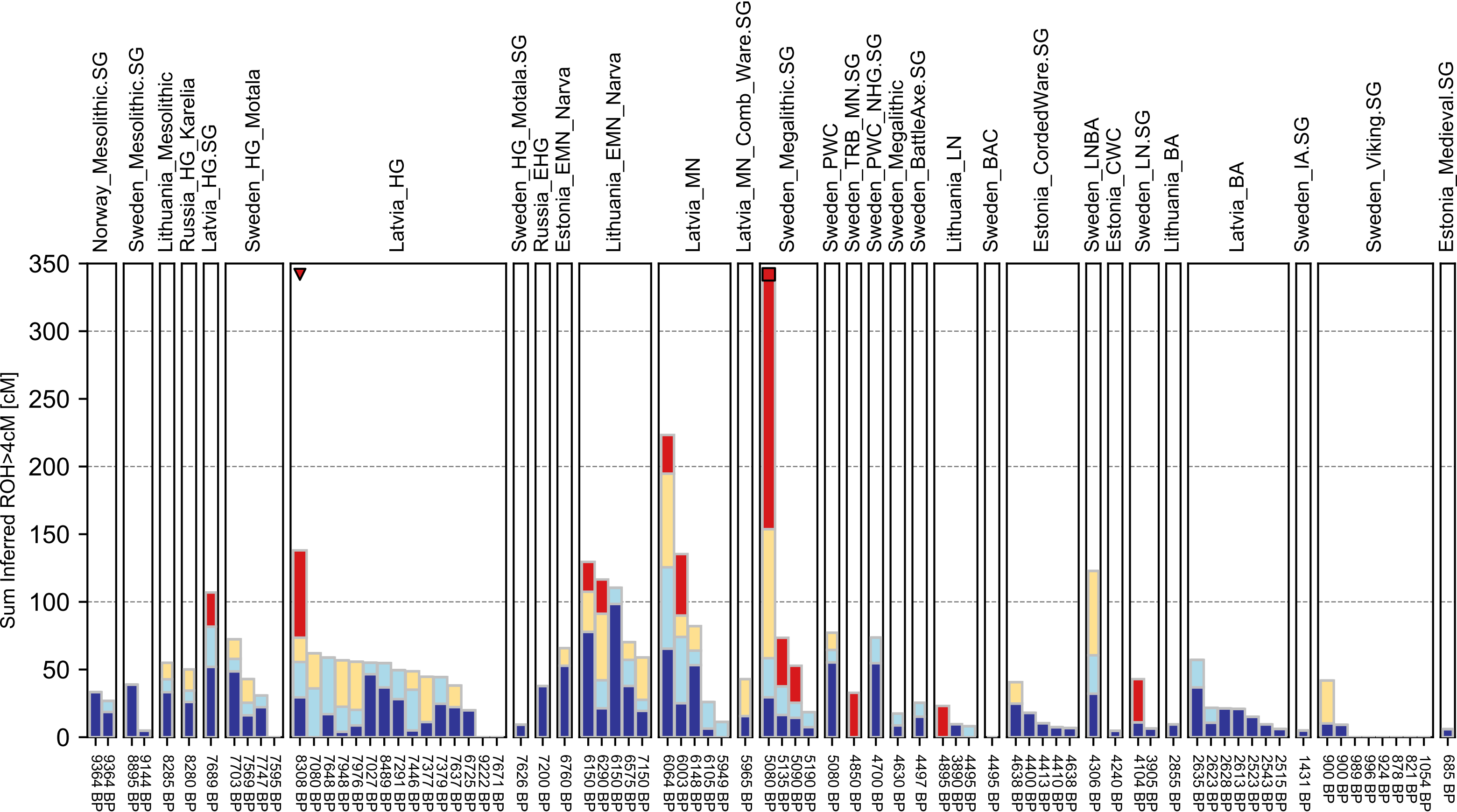

### Bering Sea.pdf

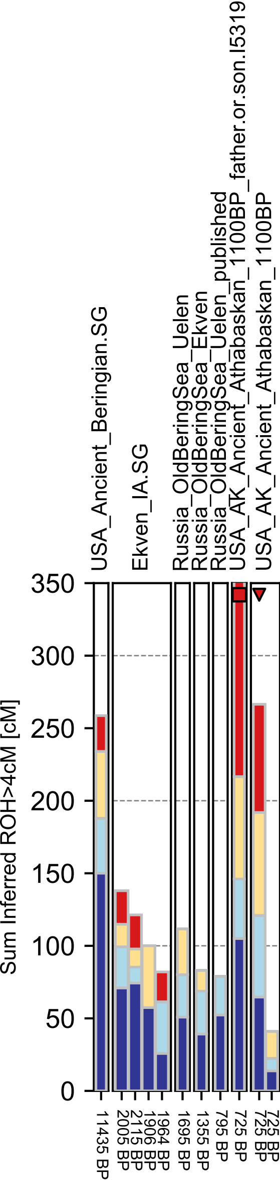

### Black Sea.pdf

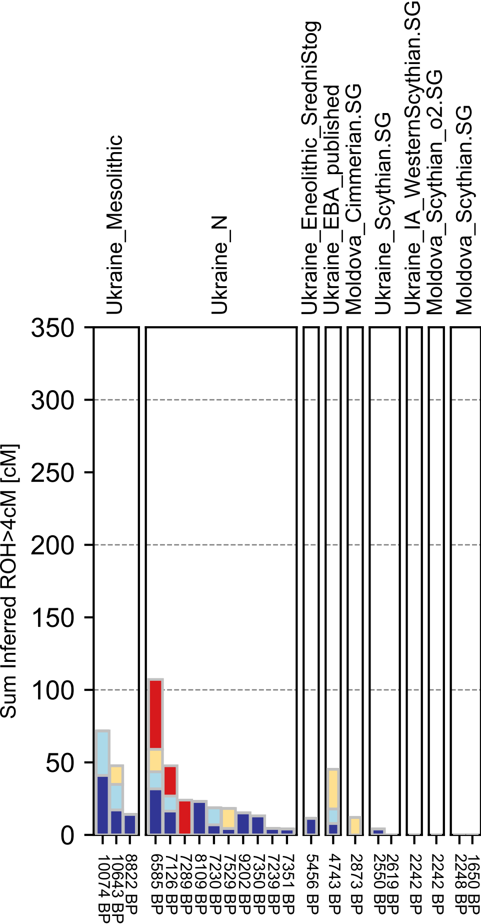

### Britain.pdf

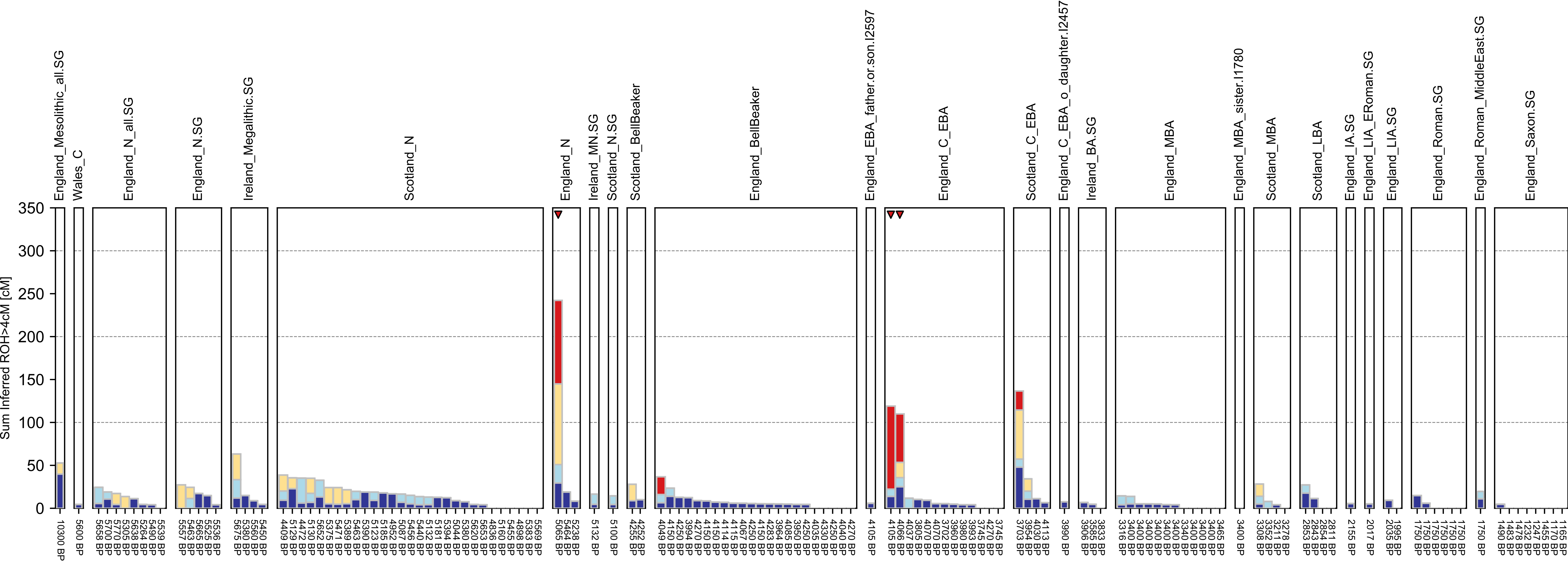

### Central Asia.pdf

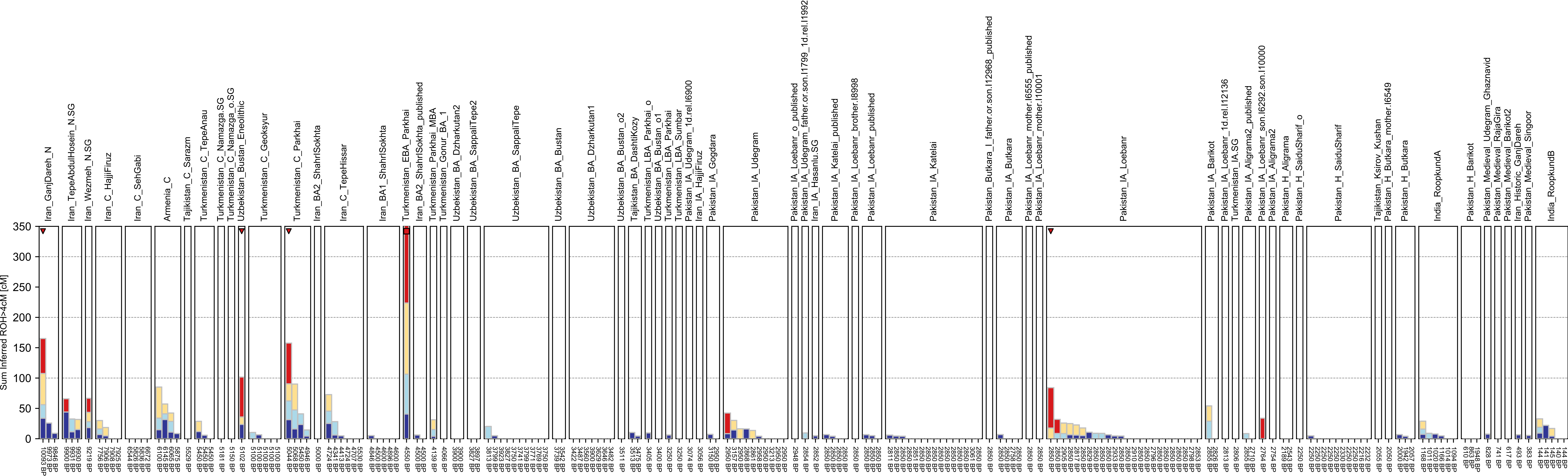

### Central Europe.pdf

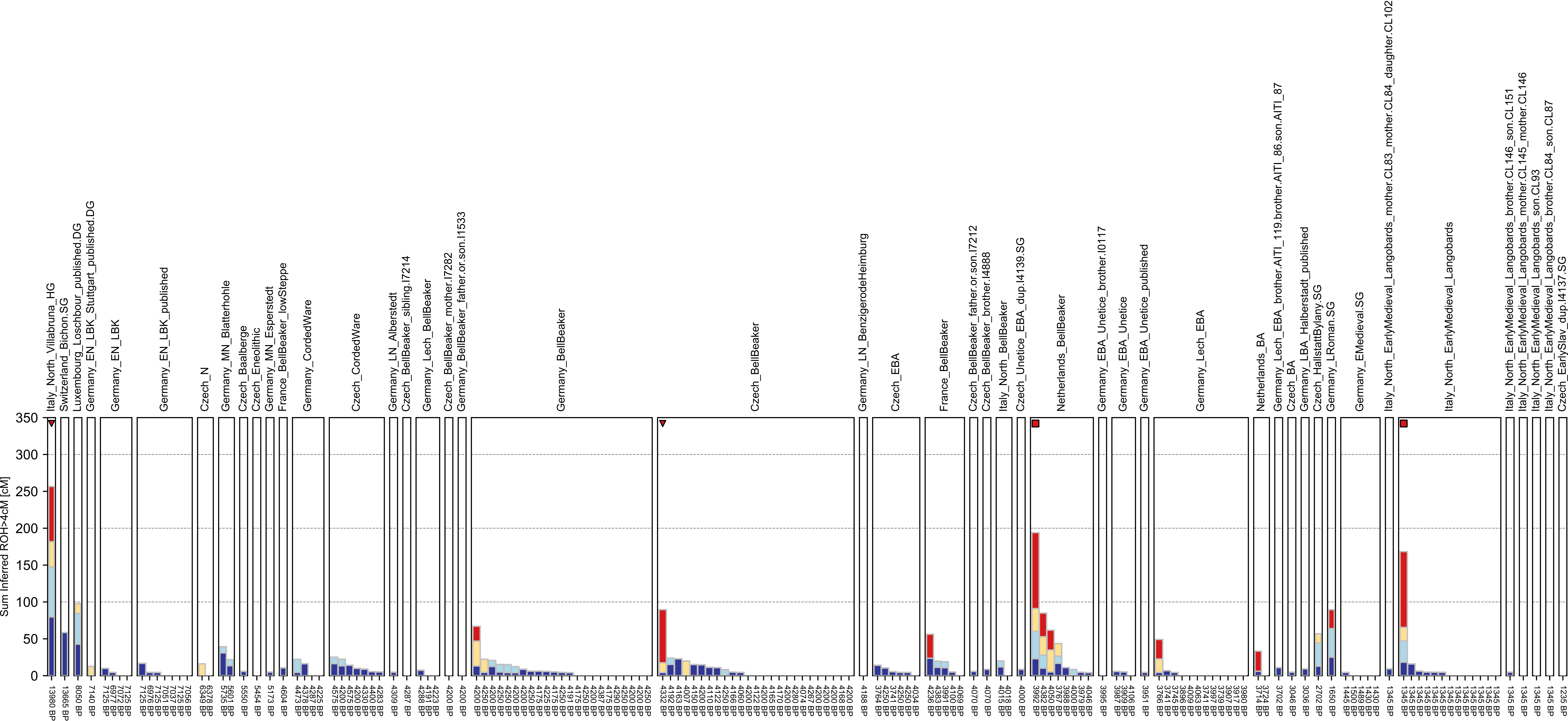

### Central Italy.pdf

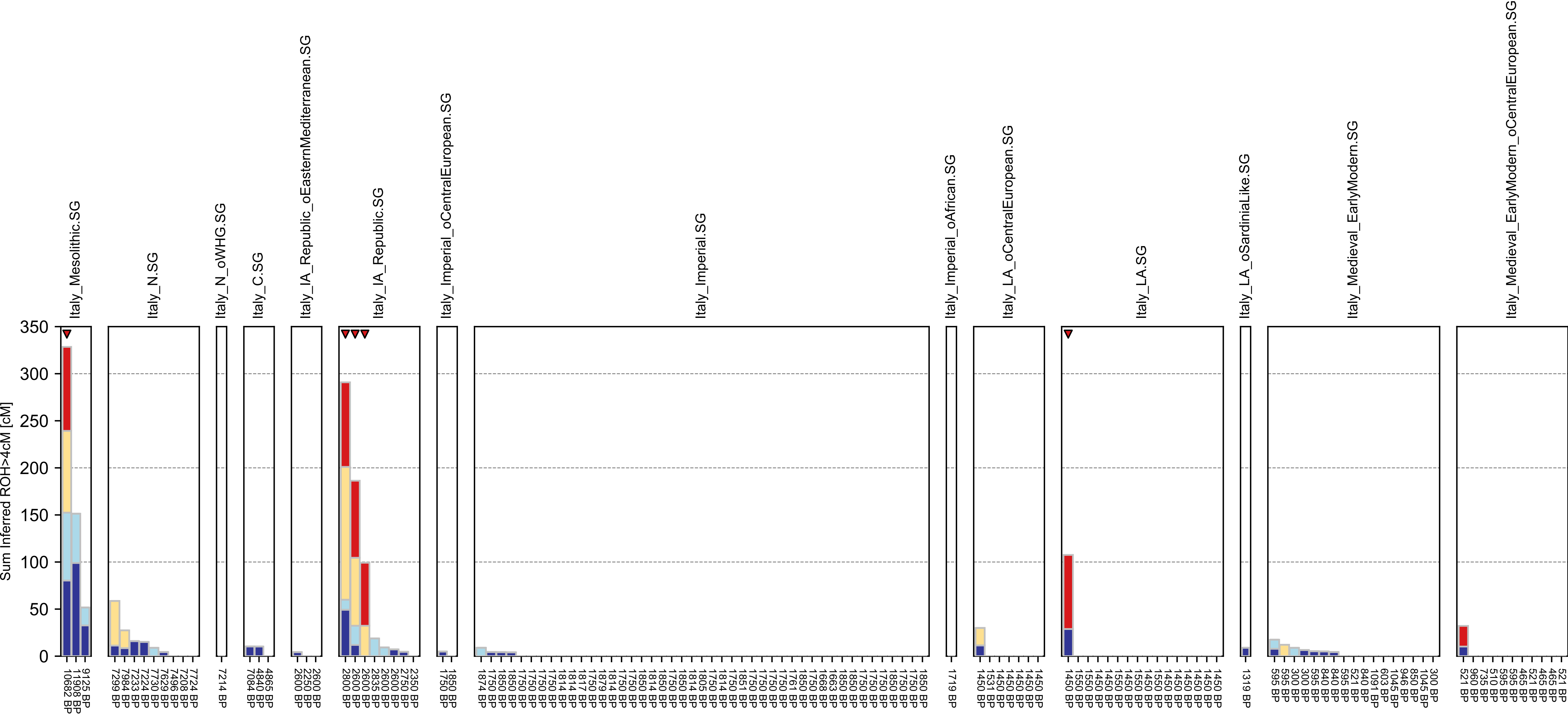

### East Africa.pdf

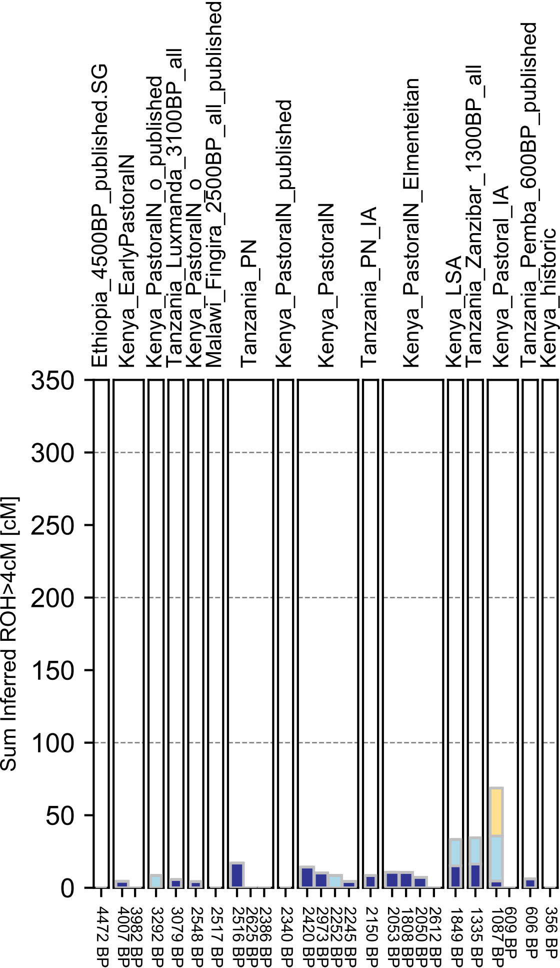

### East Steppe.pdf

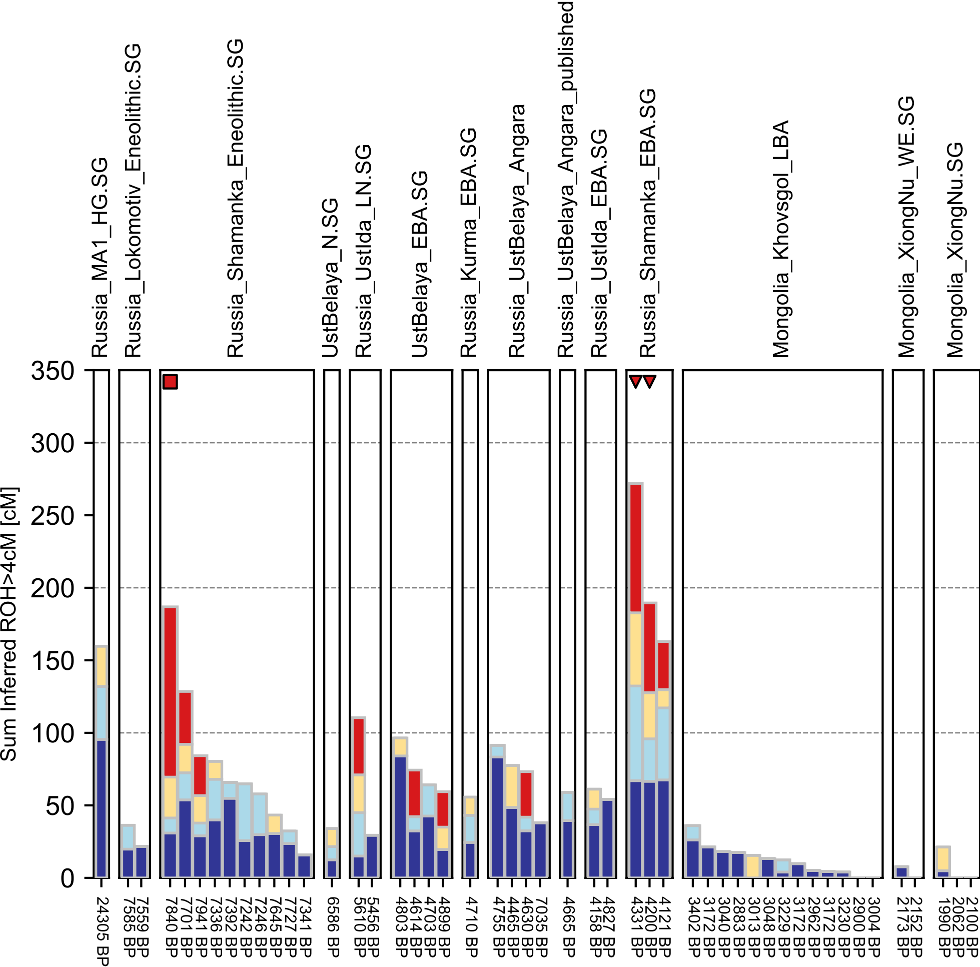

### Eastern Europe.pdf

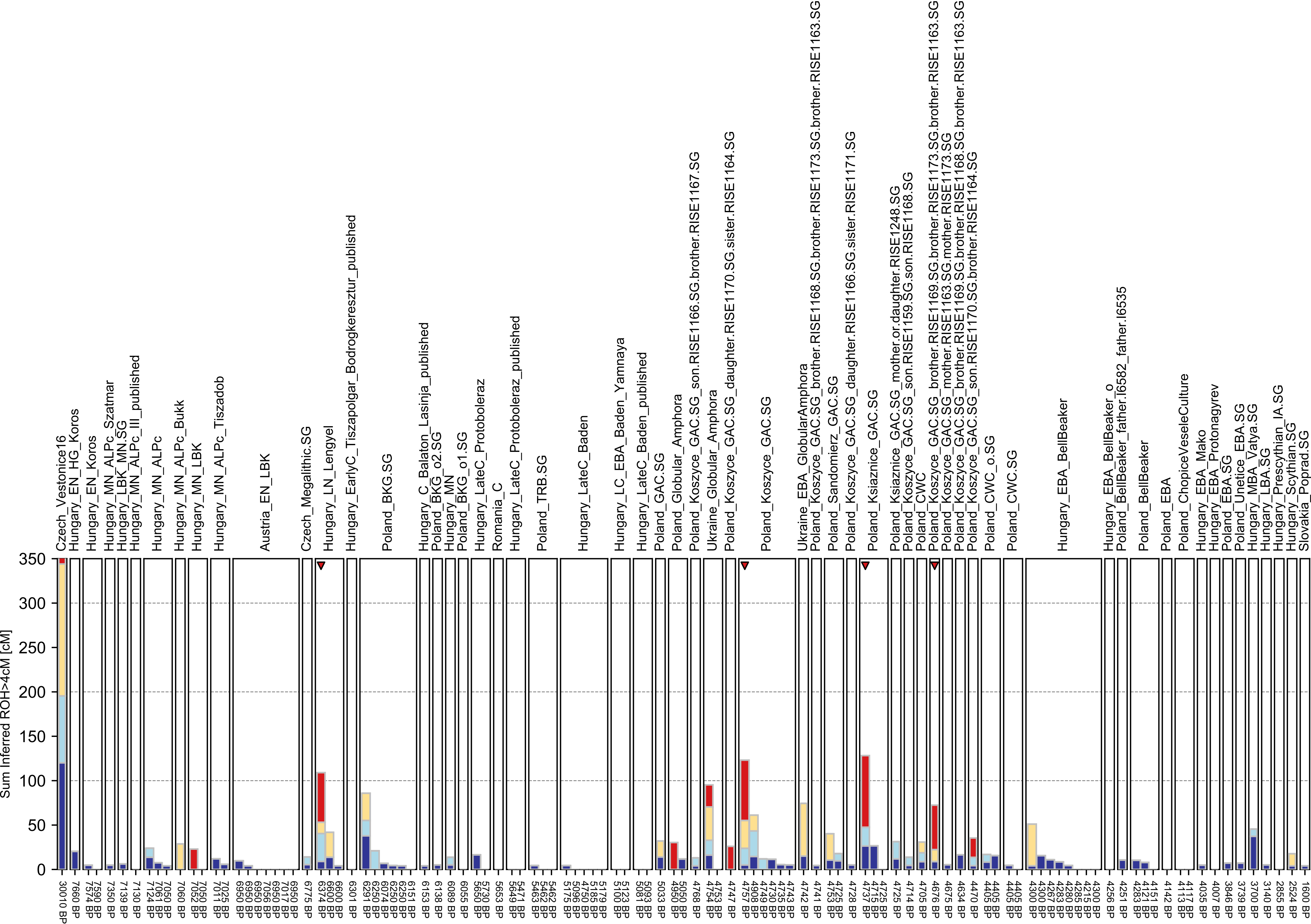

### Himalaya.pdf

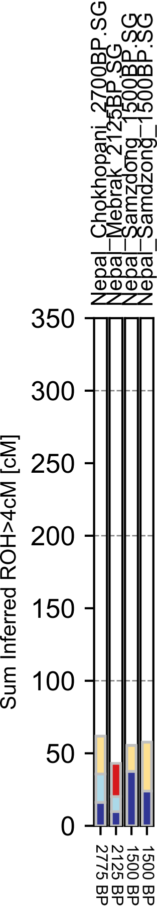

### Iberia.pdf

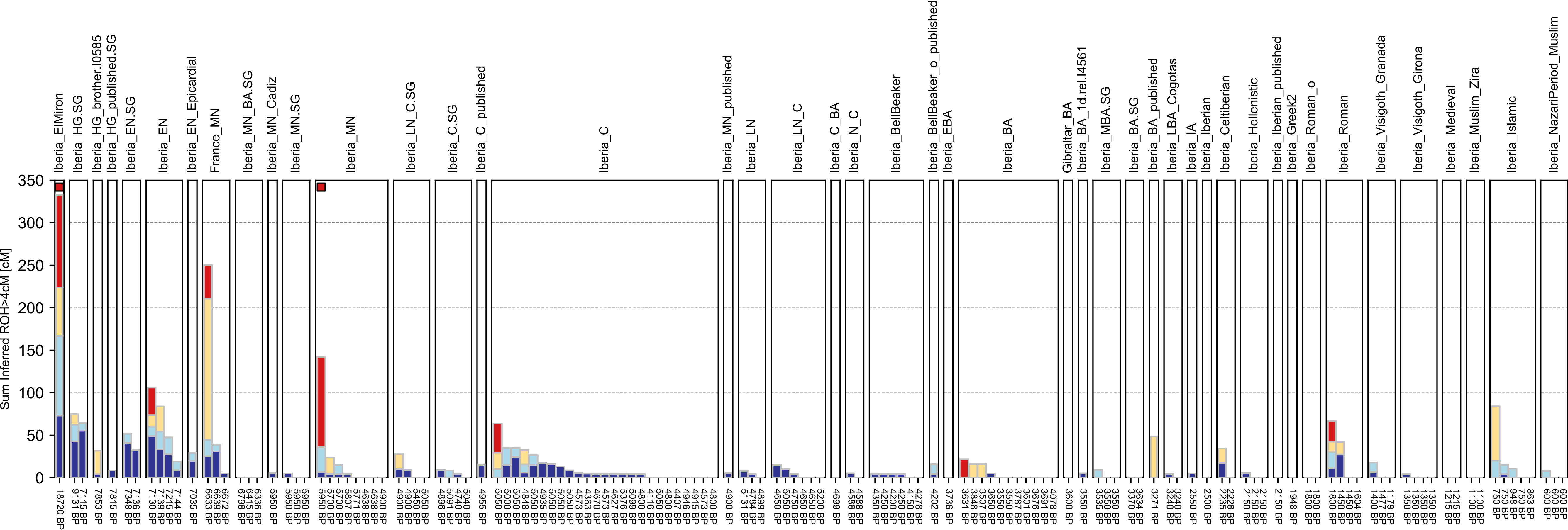

### Islands.pdf

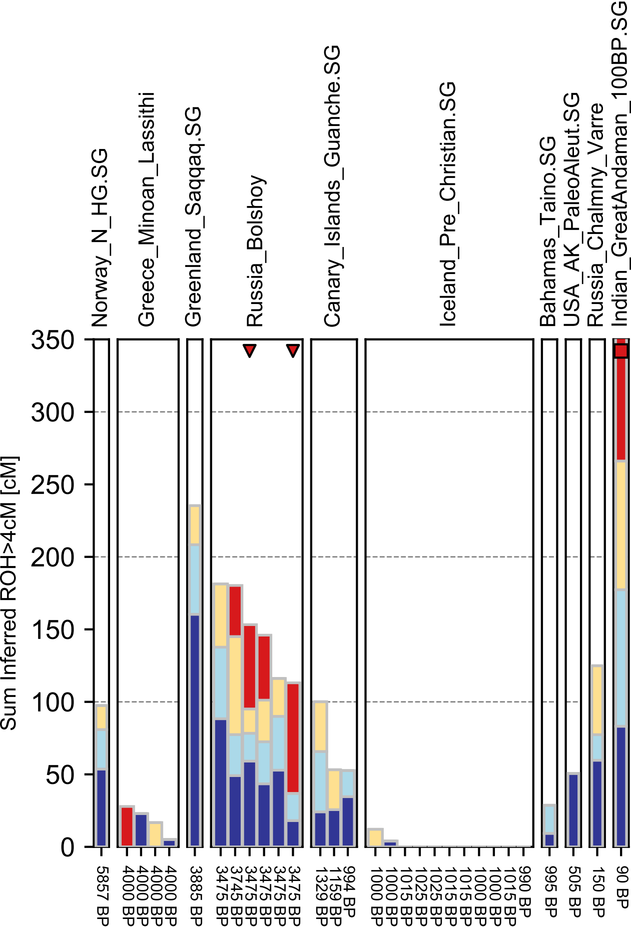

### Levante.pdf

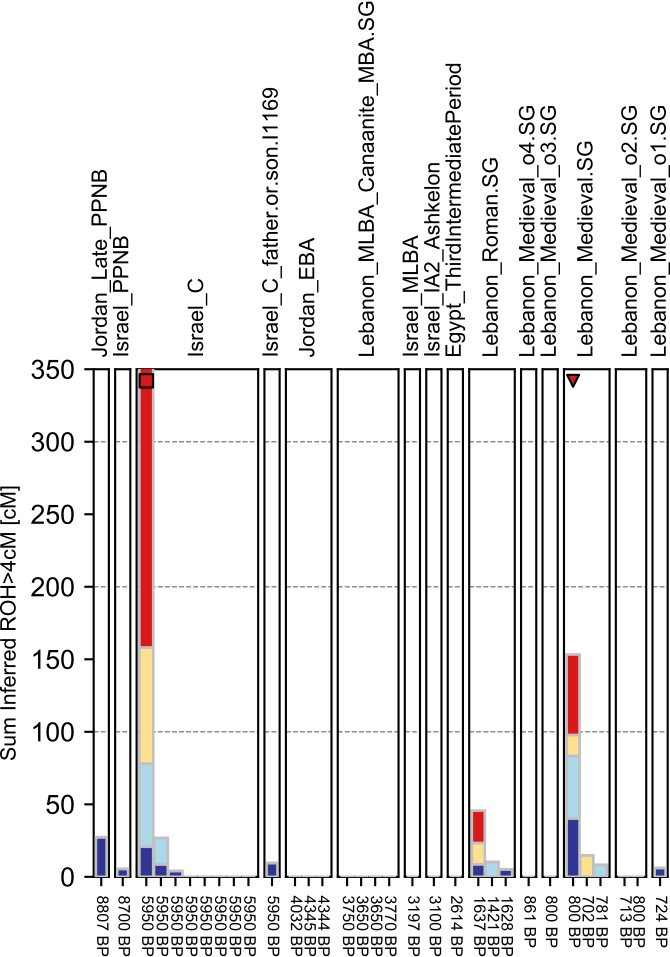

### Mende.pdf

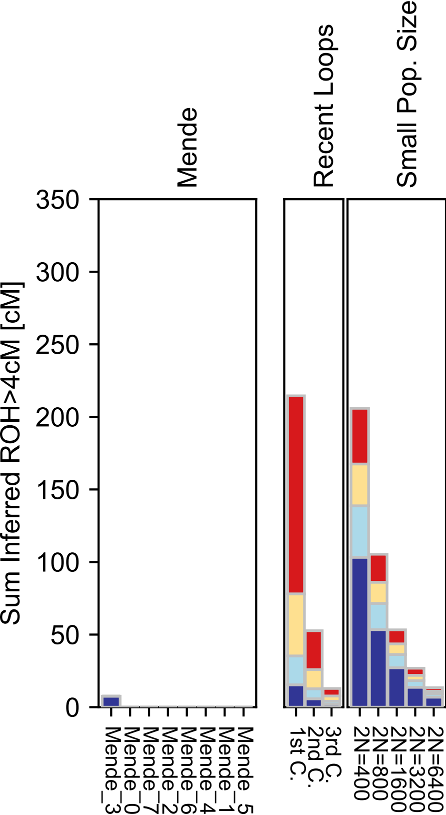

### Miao.pdf

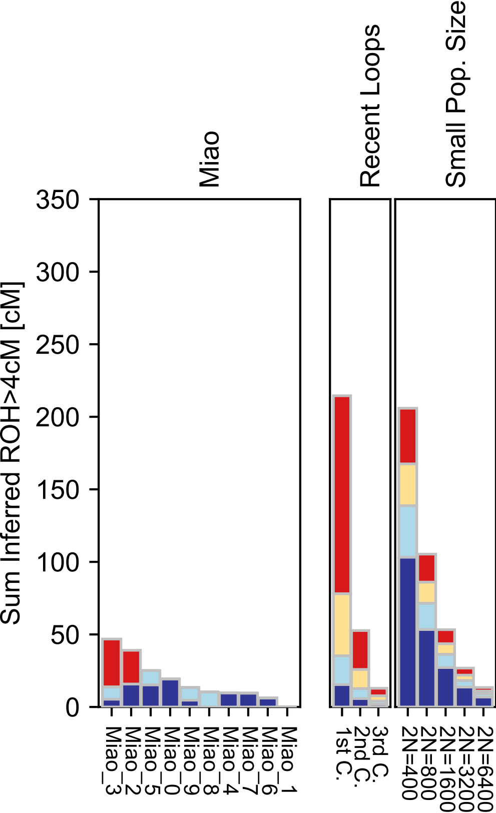

### Mixe-checkpoint.pdf

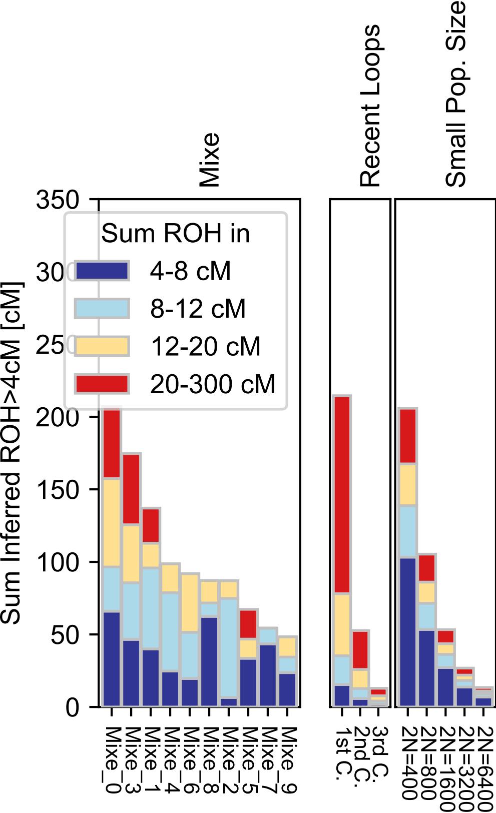

### Mixe.pdf

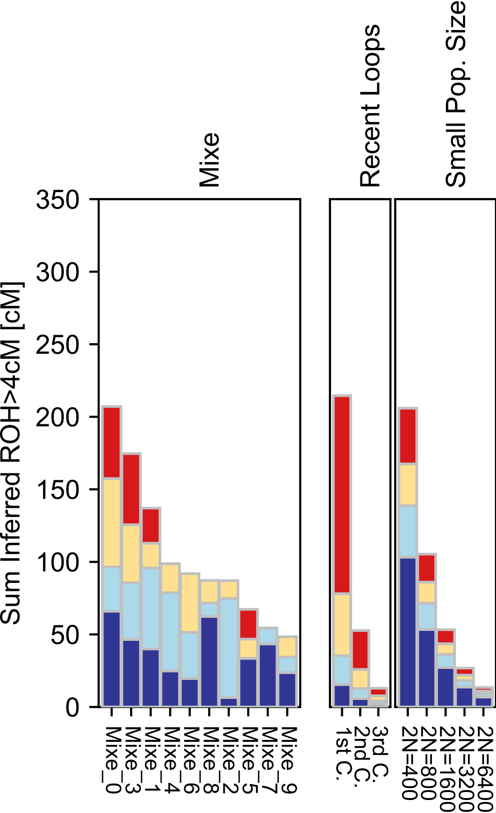

### Mixtec.pdf

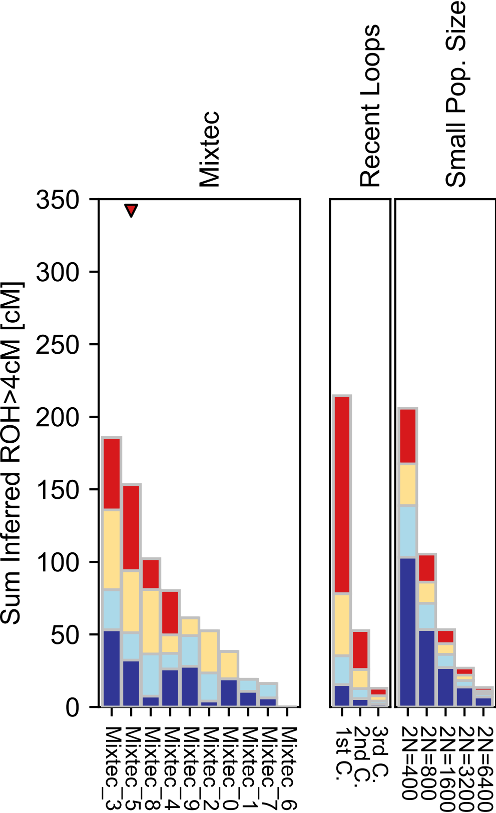

### Mongola.pdf

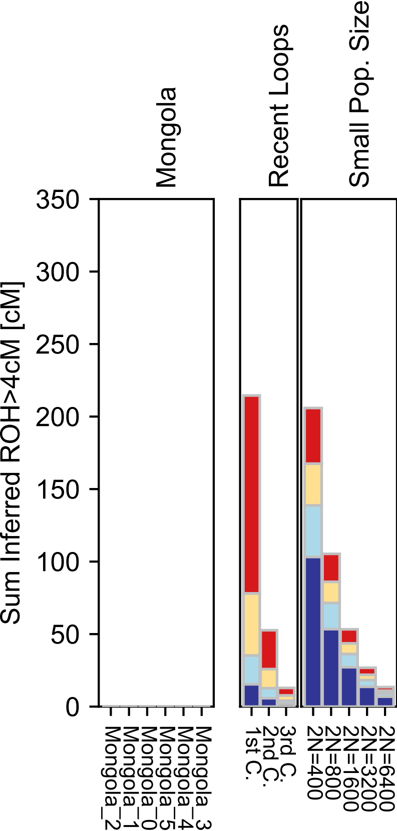

### Mordovian.pdf

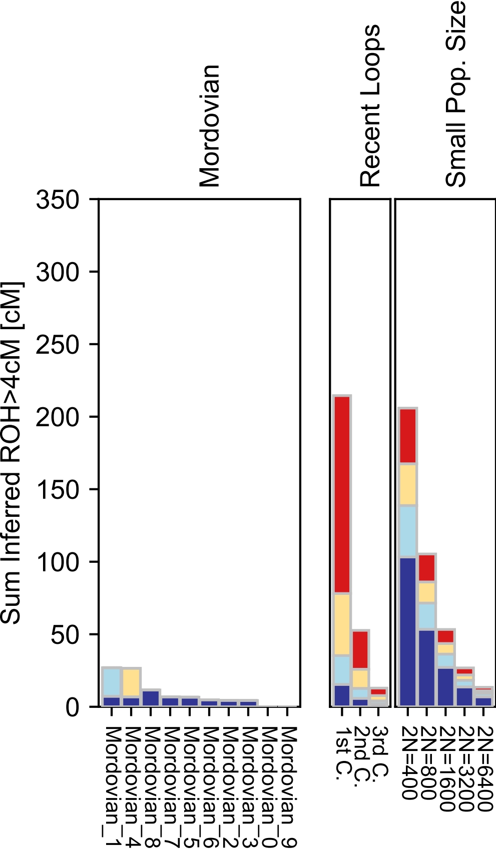

### Moroccan_Jew.pdf

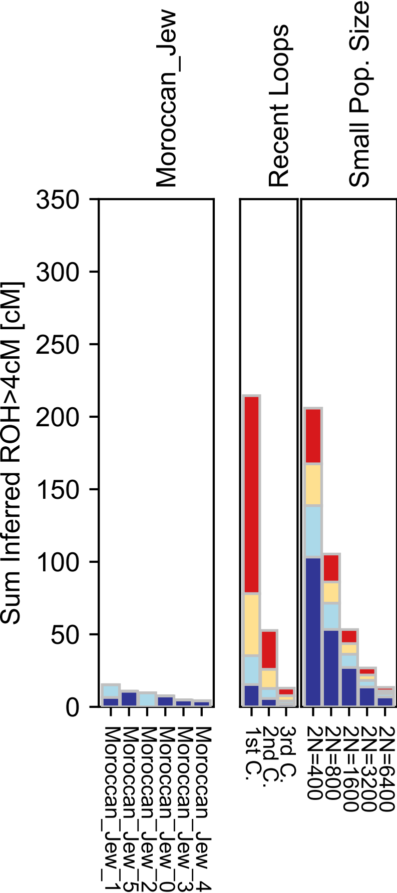

### Mozabite.pdf

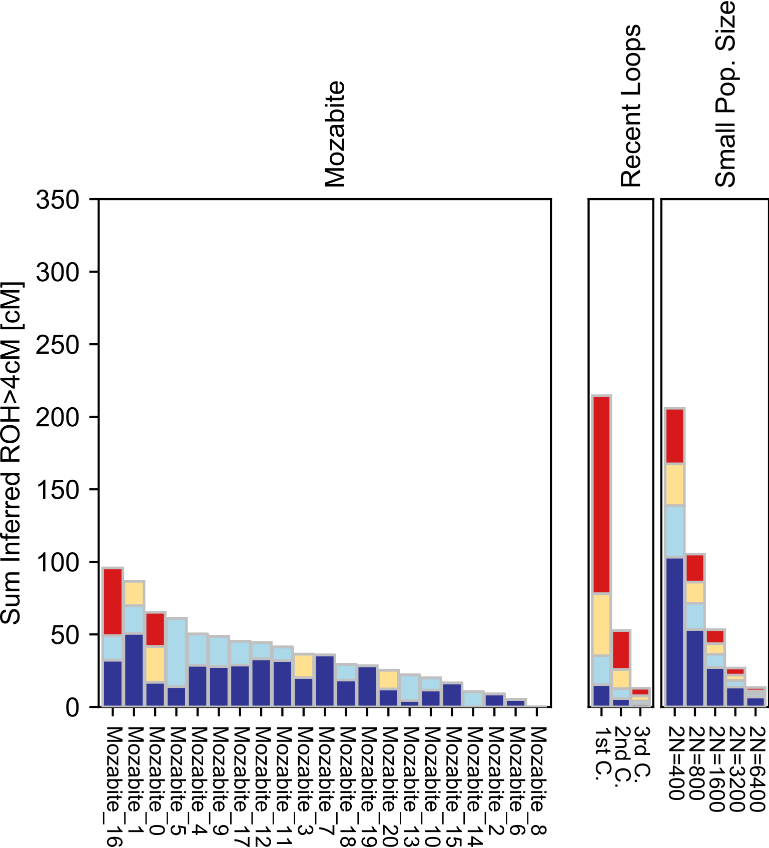

### Naxi.pdf

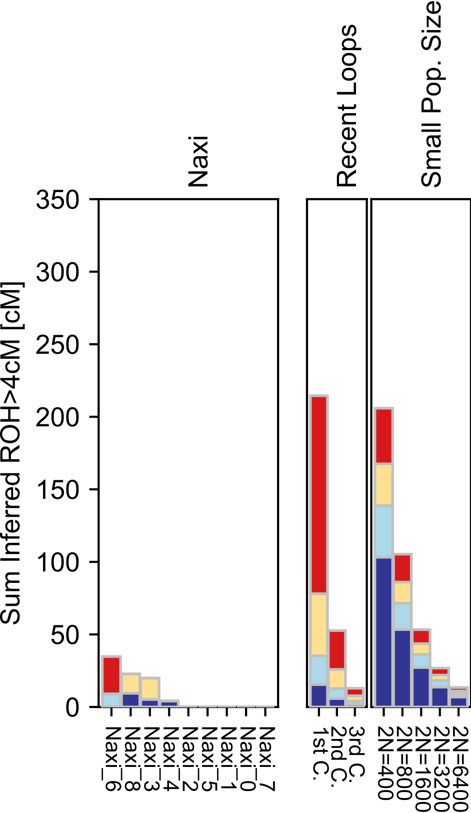

### Nganasan.pdf

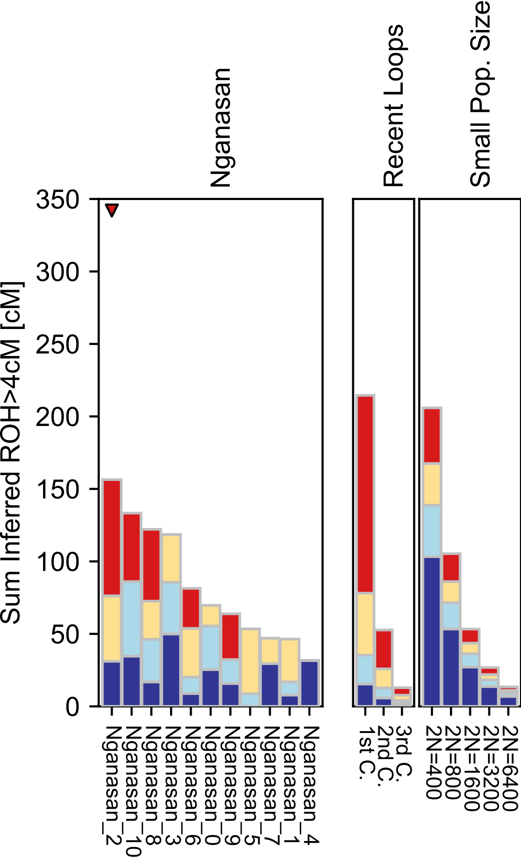

### Nogai.pdf

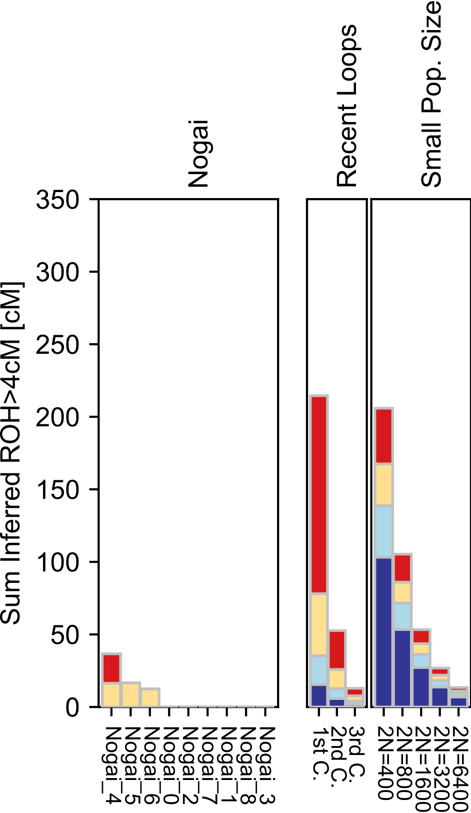

### North Africa.pdf

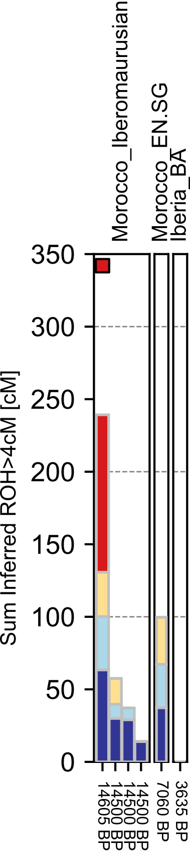

### North_Ossetian.pdf

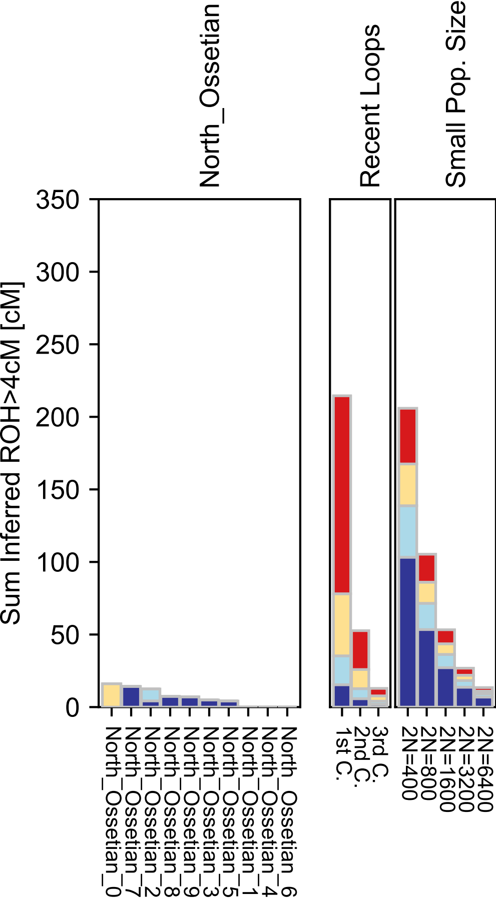

### Norwegian.pdf

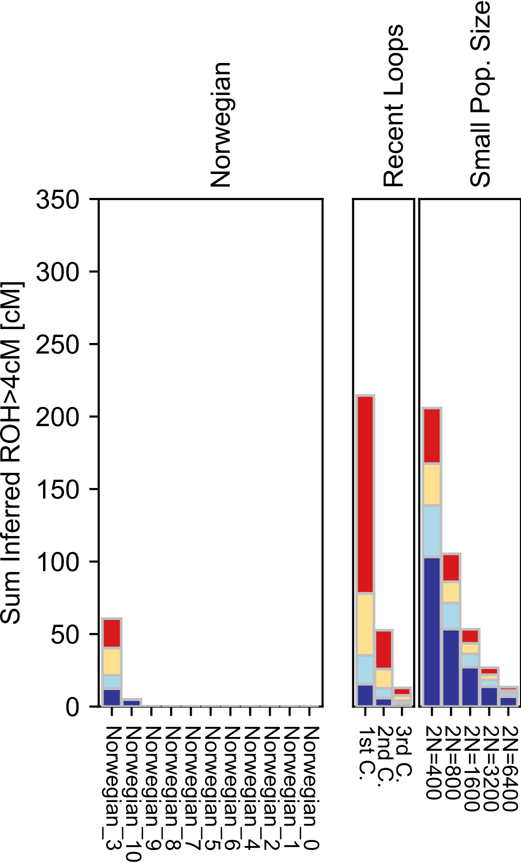

### Not Assigned.pdf

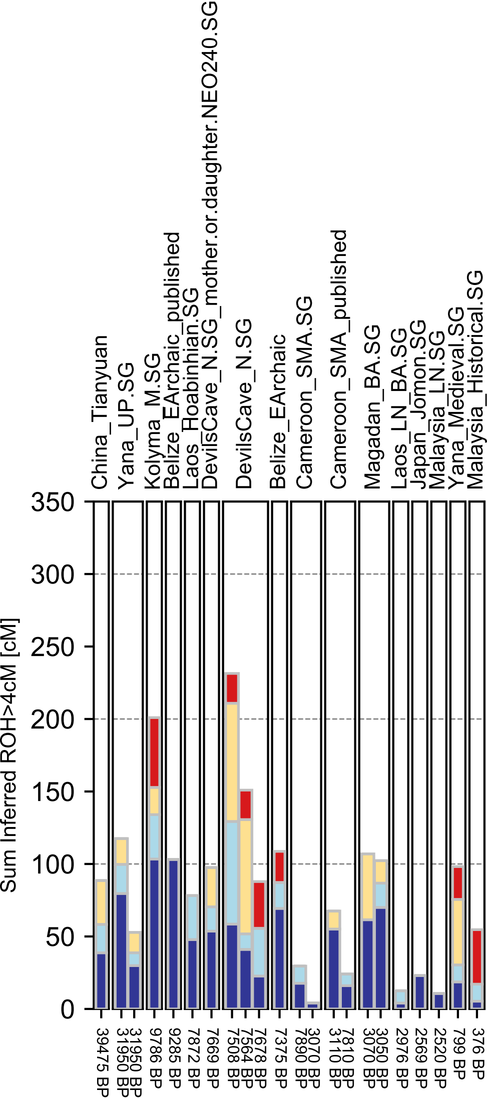

### Orcadian.pdf

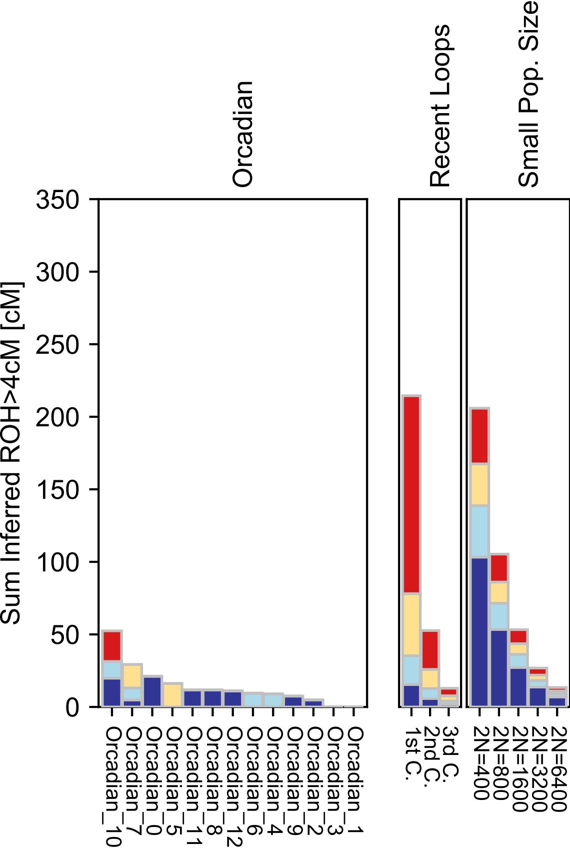

### Oroqen.pdf

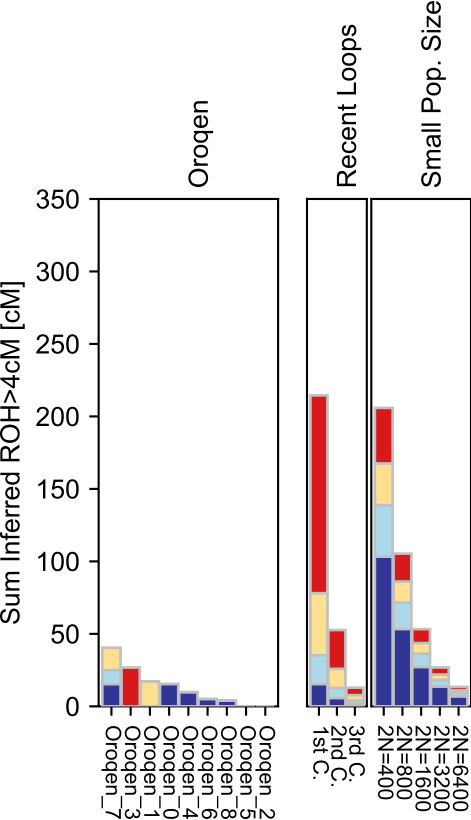

### Pacific NW.pdf

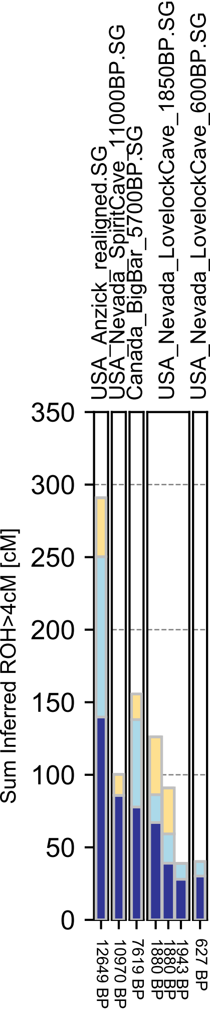

### Palestinian.pdf

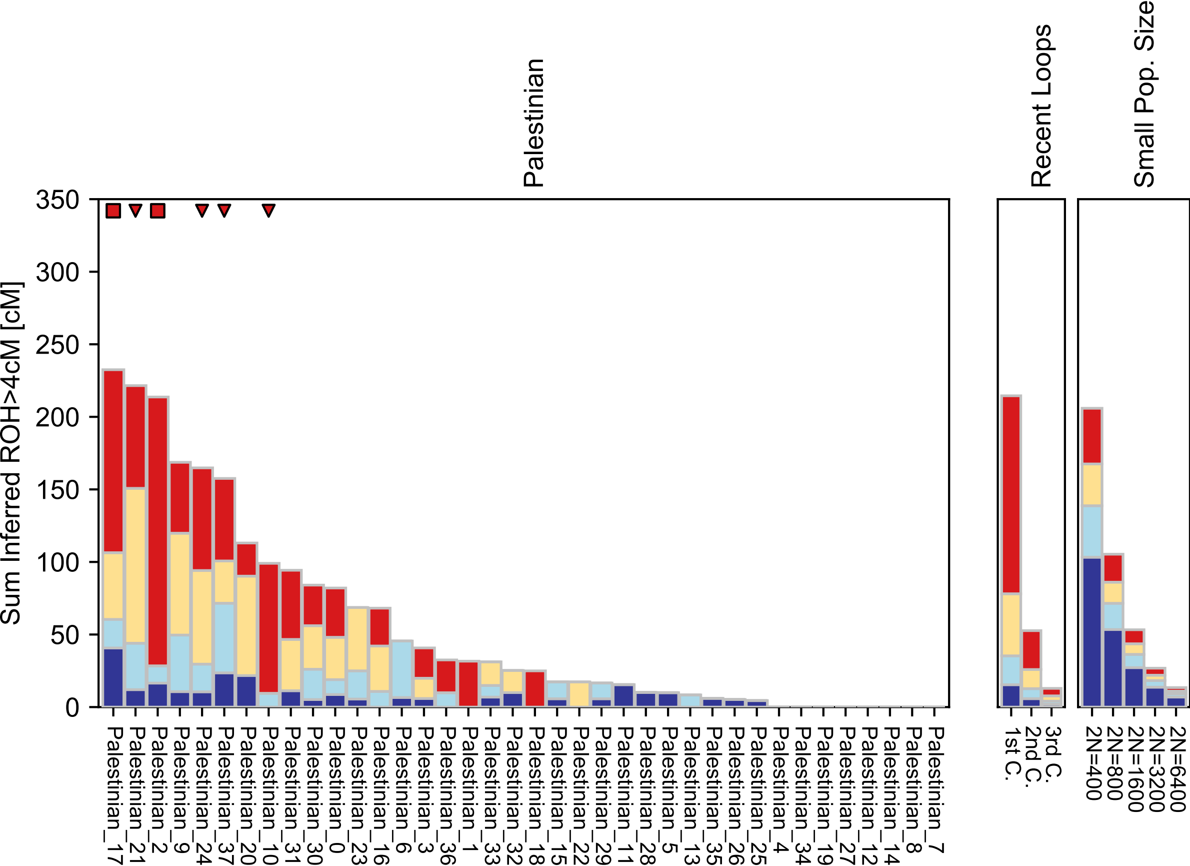

### Papuan.pdf

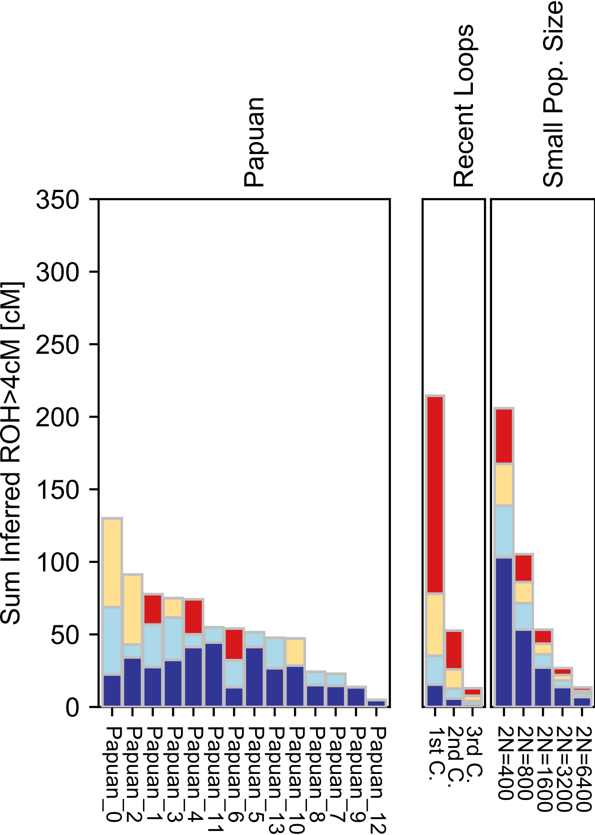

### Patagonia.pdf

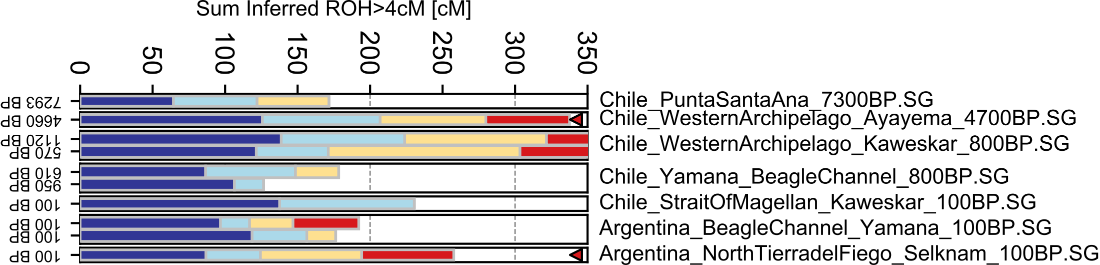

### Pathan.pdf

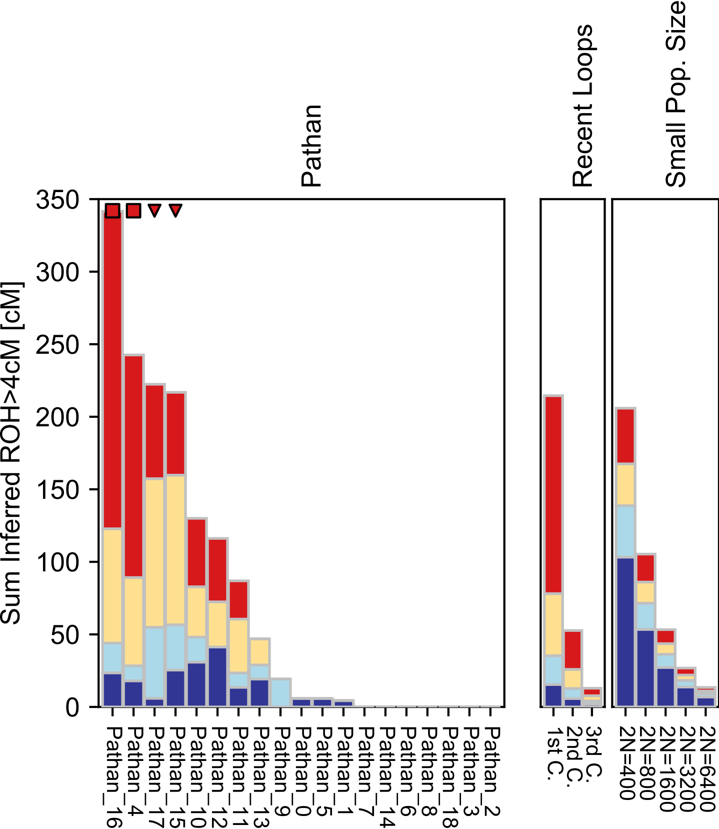

### Piapoco.pdf

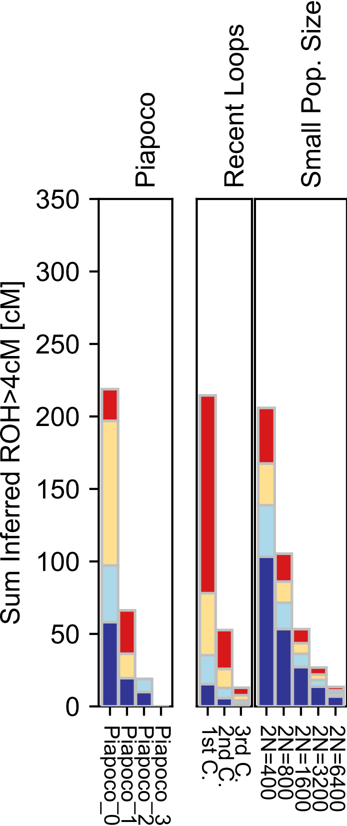

### Pima.pdf

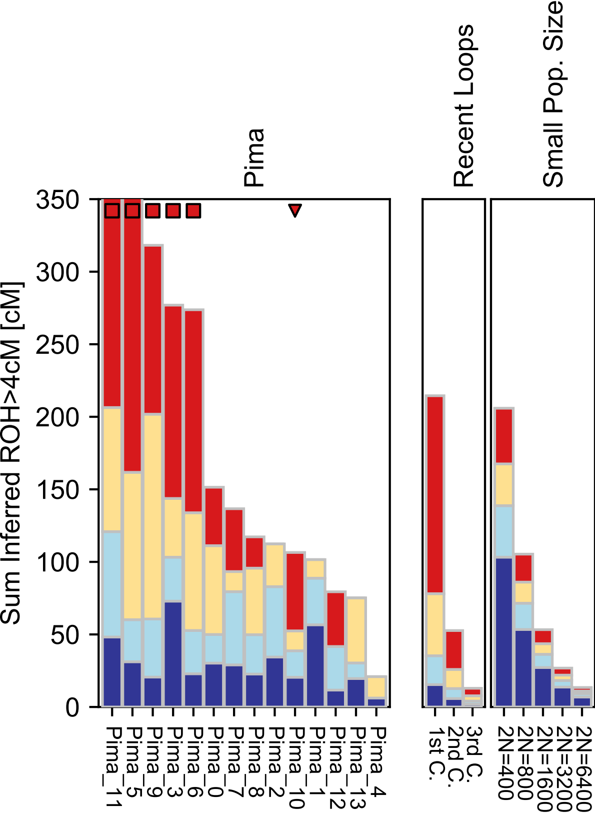

### Punjabi.pdf

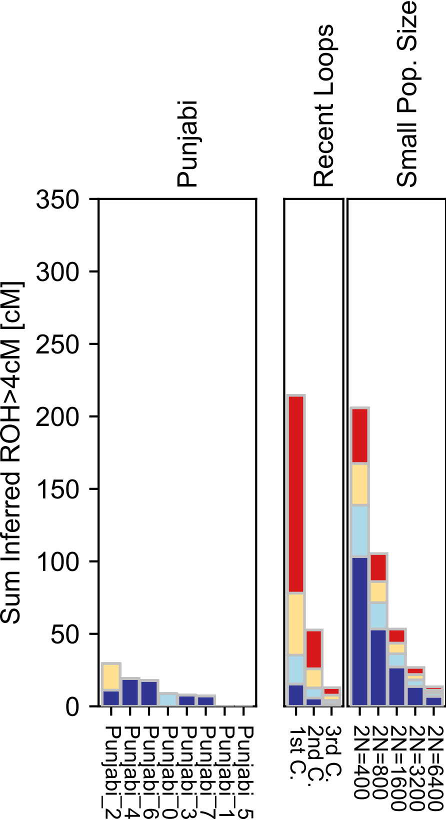

### Quechua.pdf

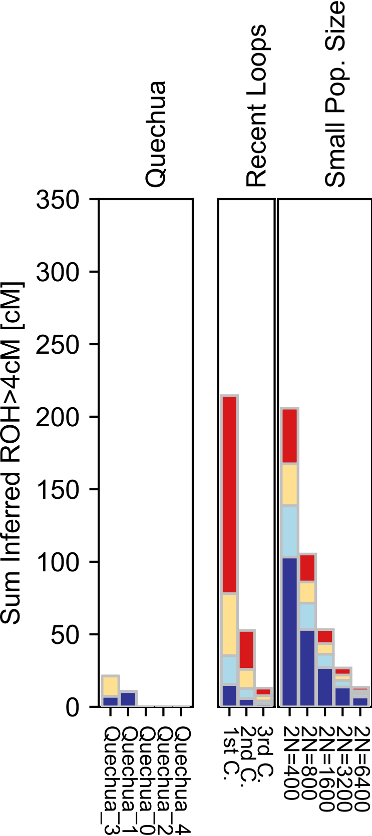

### Russian.pdf

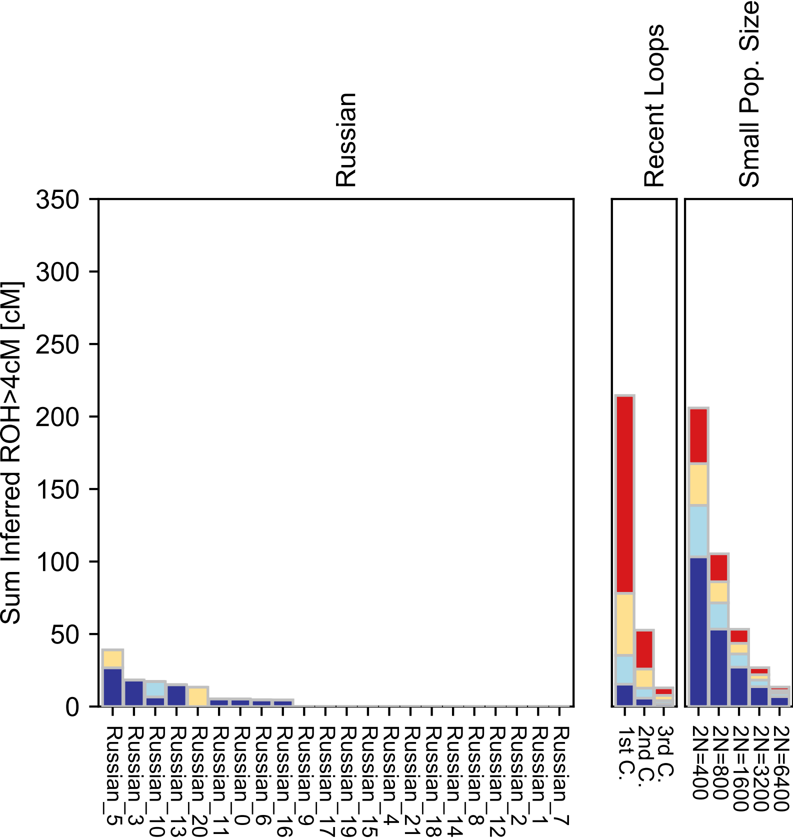

### Saami_WGA.pdf

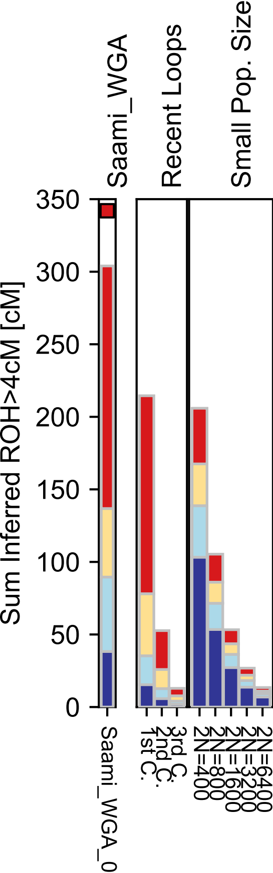

### Saharawi.pdf

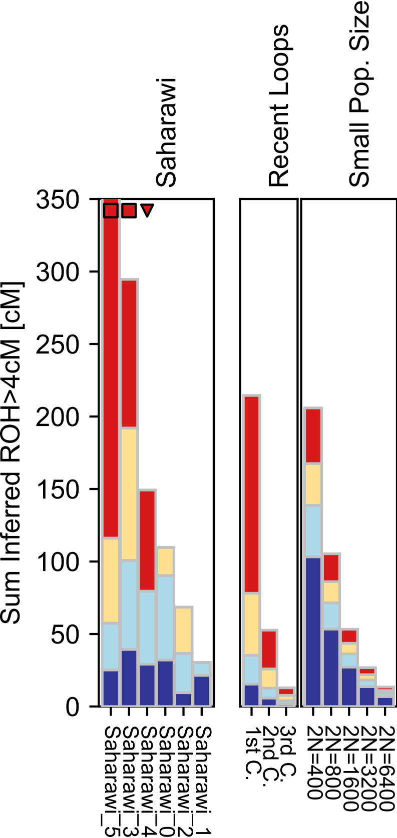

### Sardinia.pdf

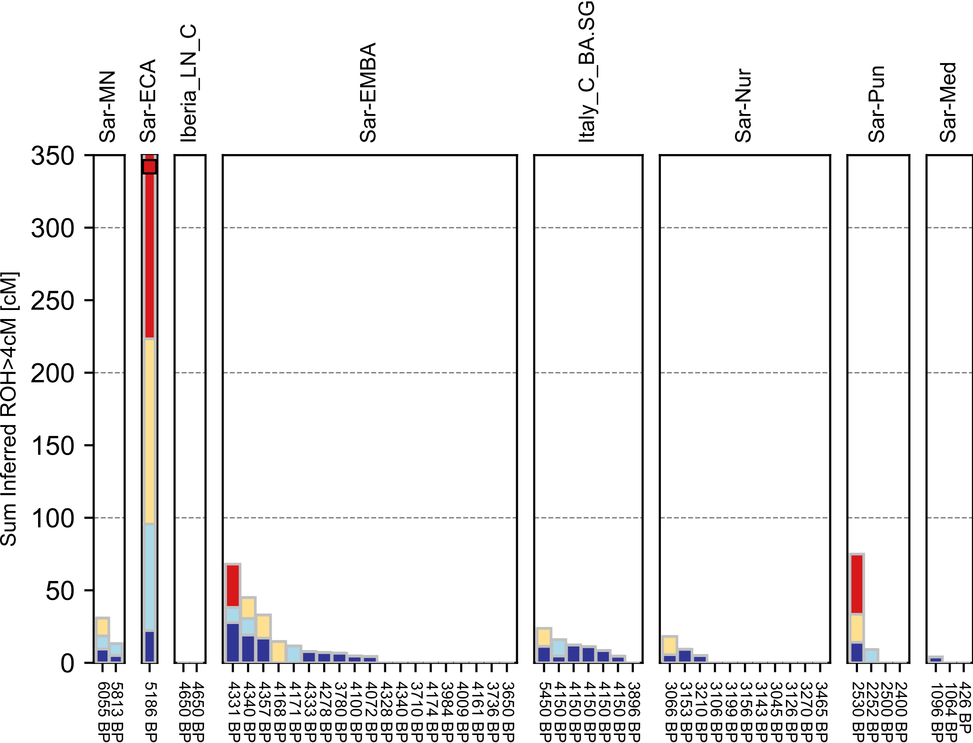

### Sardinian-checkpoint.pdf

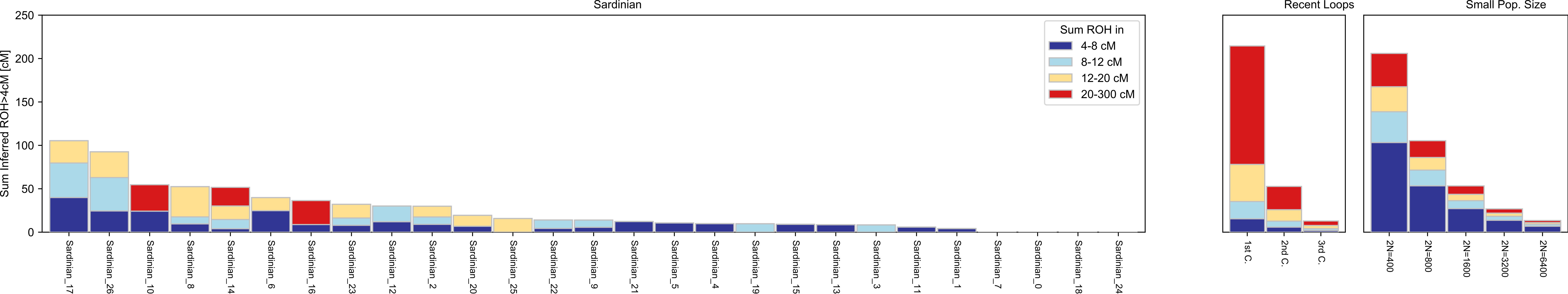

### Sardinian.pdf

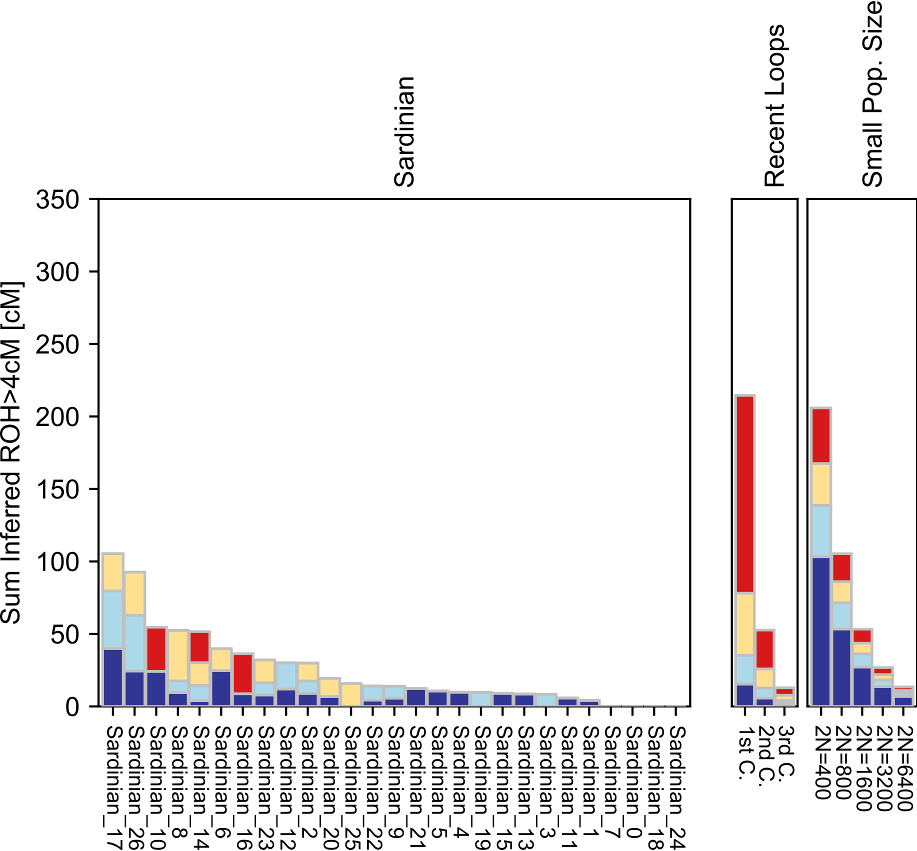

### Saudi.pdf

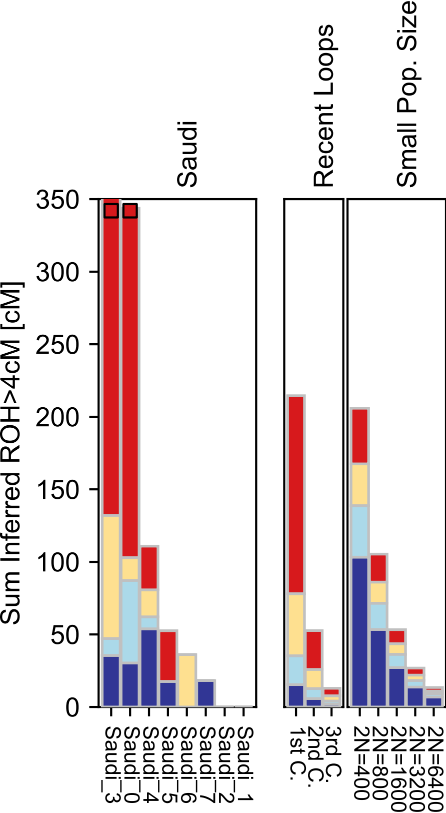

### Scottish.pdf

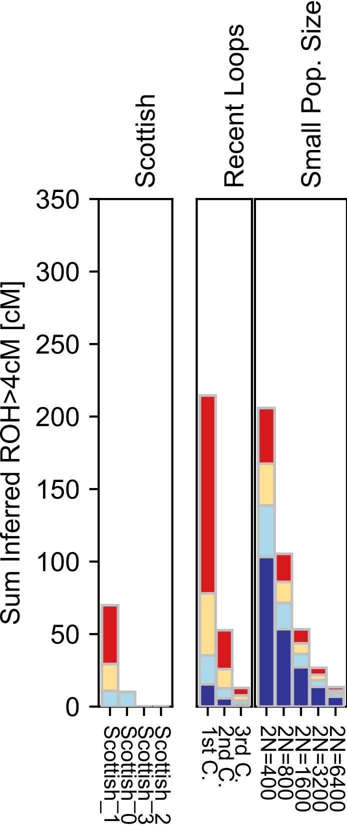

### Selkup.pdf

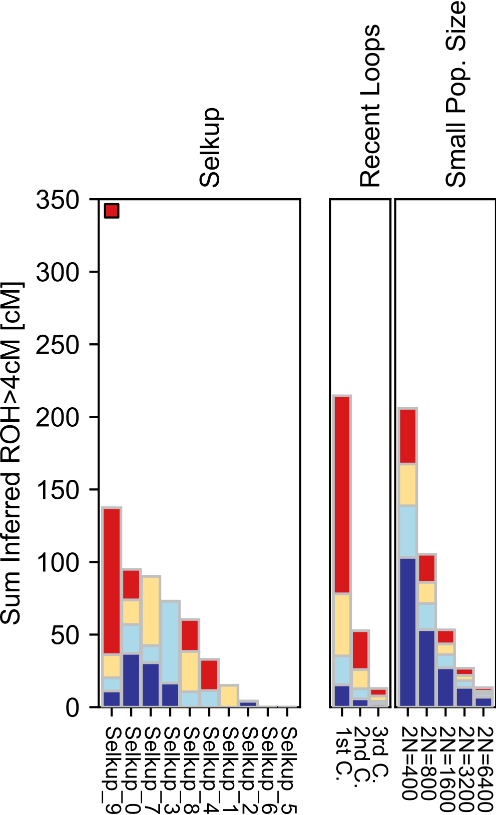

### She.pdf

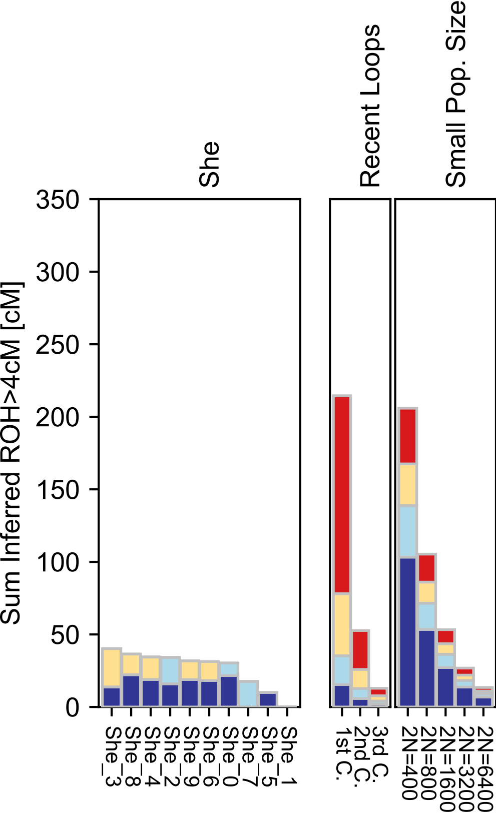

### Sicilian.pdf

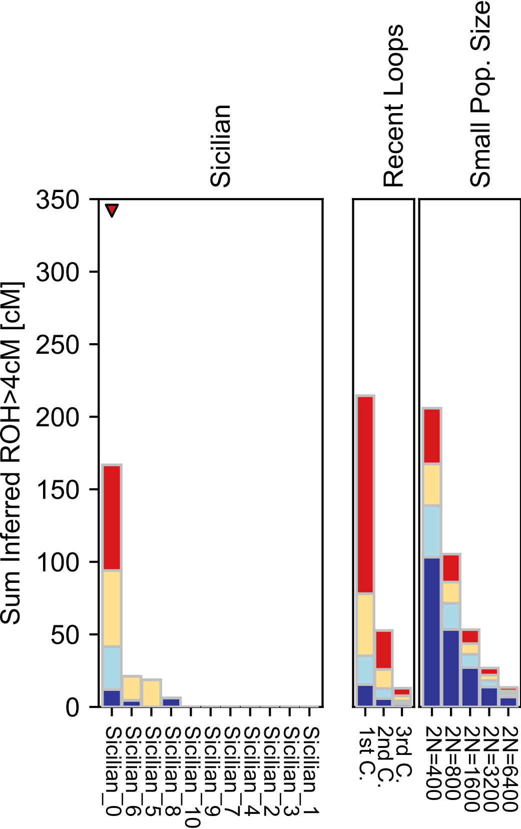

### Sindhi-checkpoint.pdf

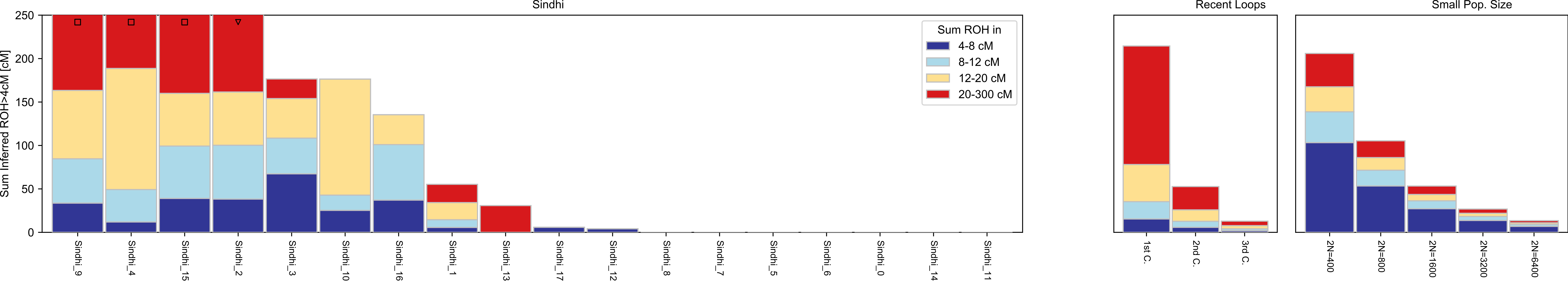

### Sindhi.pdf

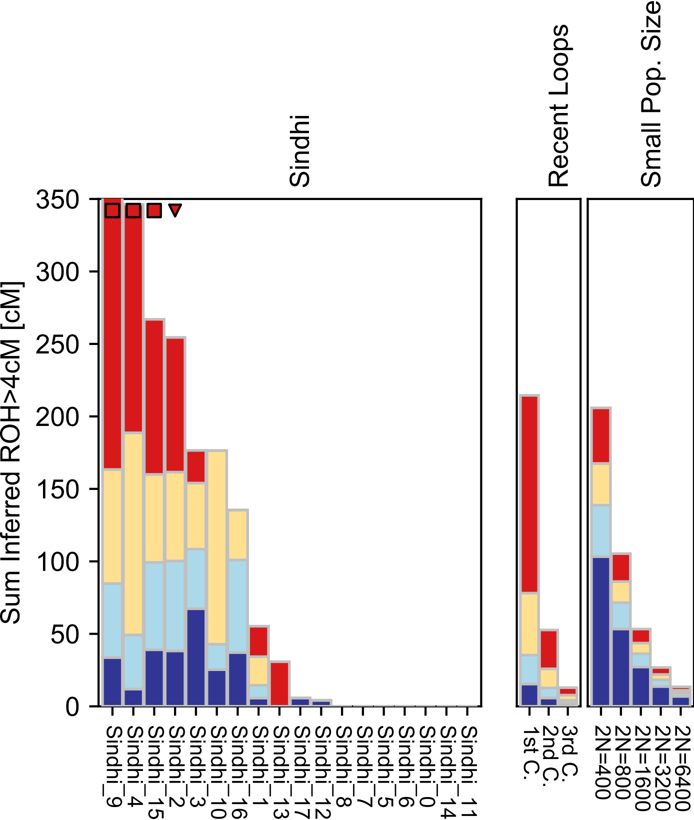

### Somali.pdf

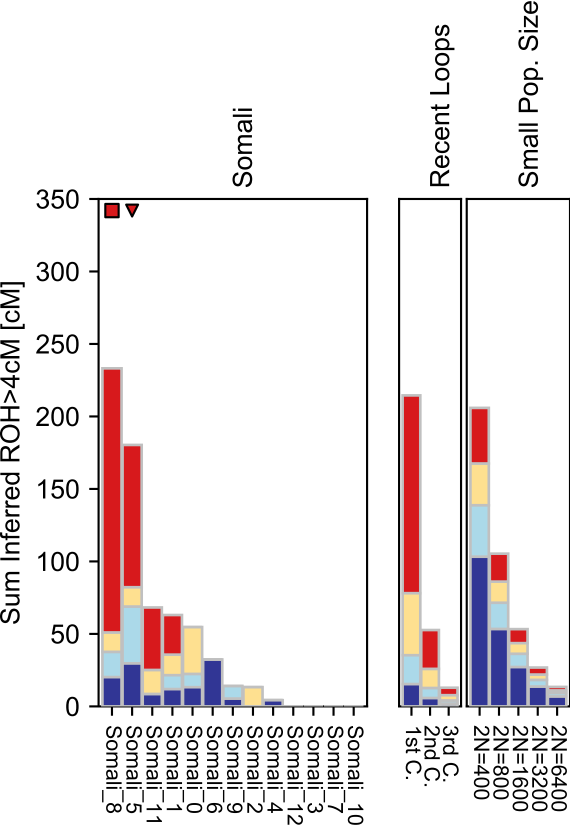

### South Africa.pdf

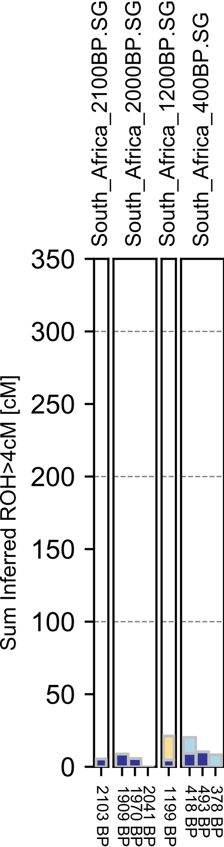

### Spanish.pdf

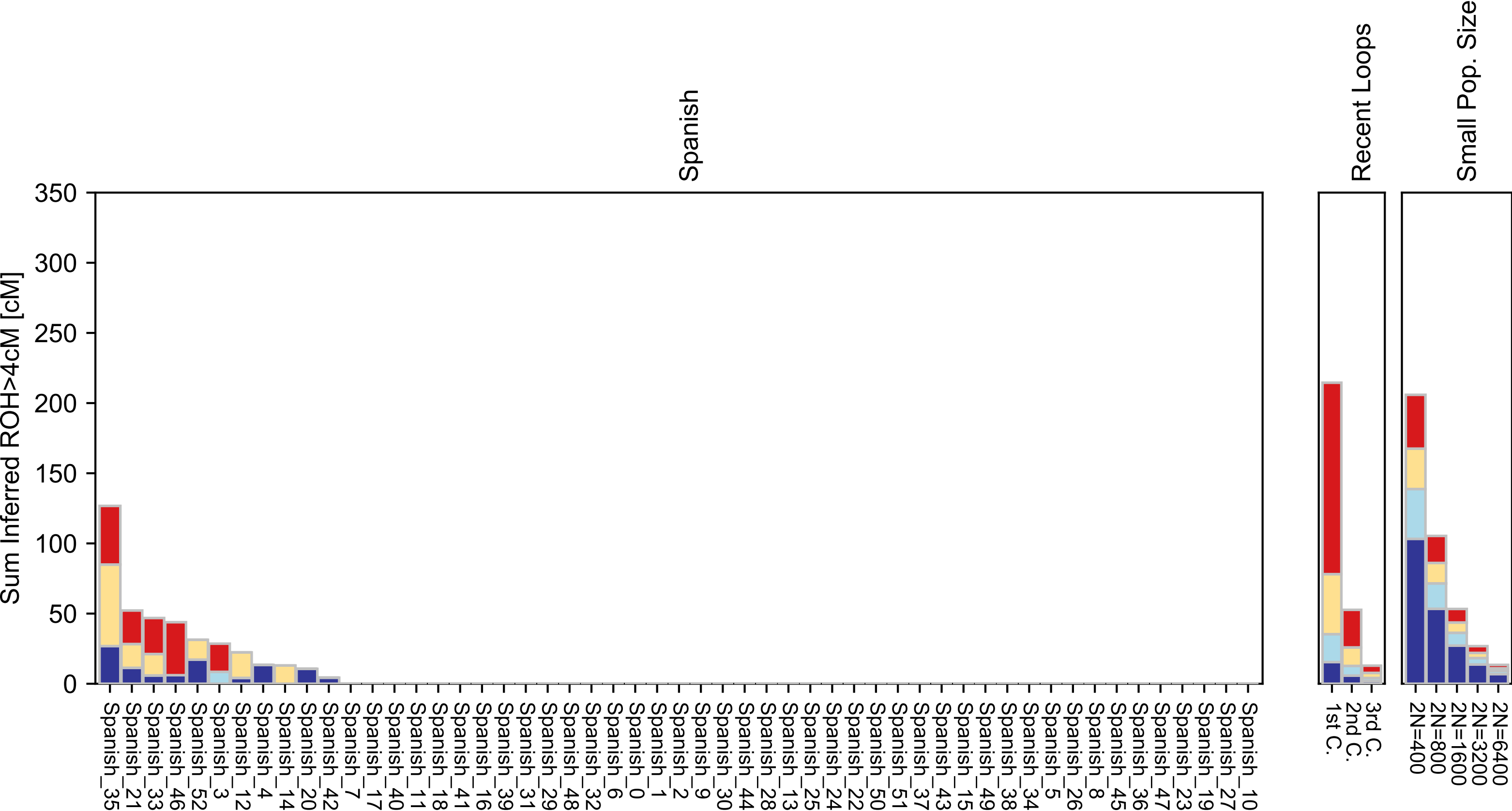

### Spanish_North.pdf

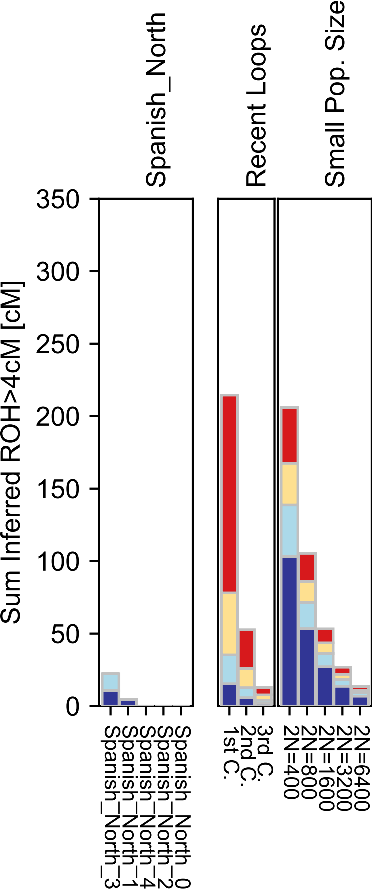

### Steppe.pdf

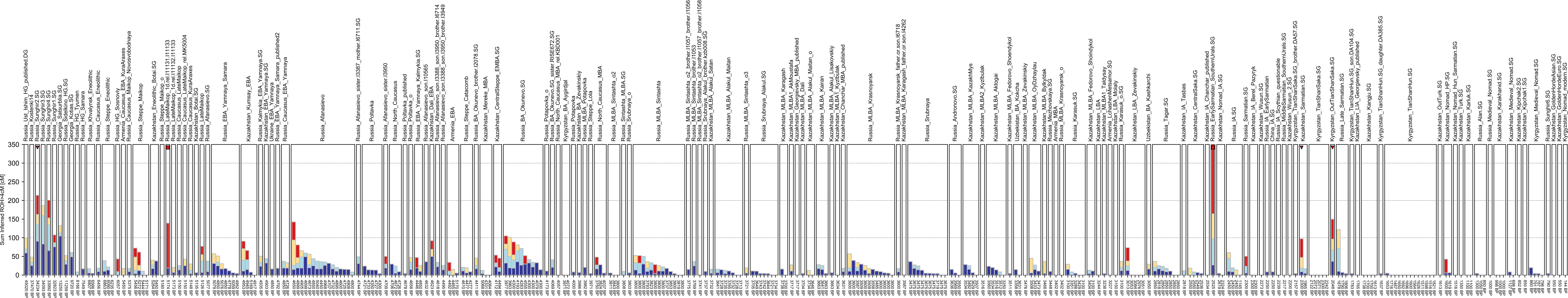

### Surui.pdf

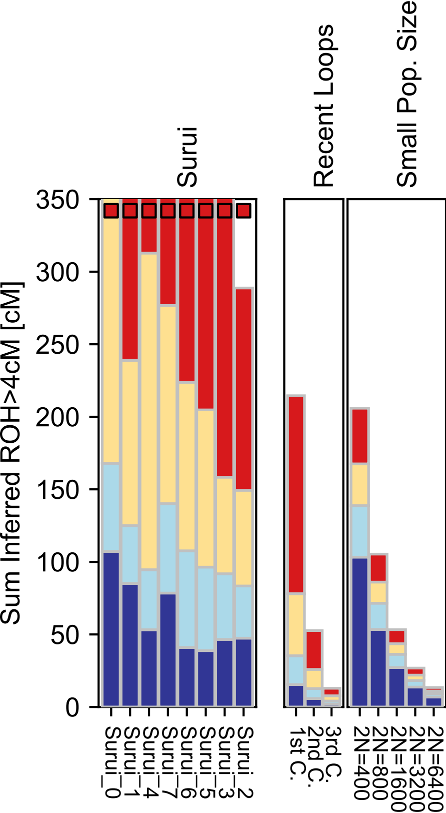

### Syrian.pdf

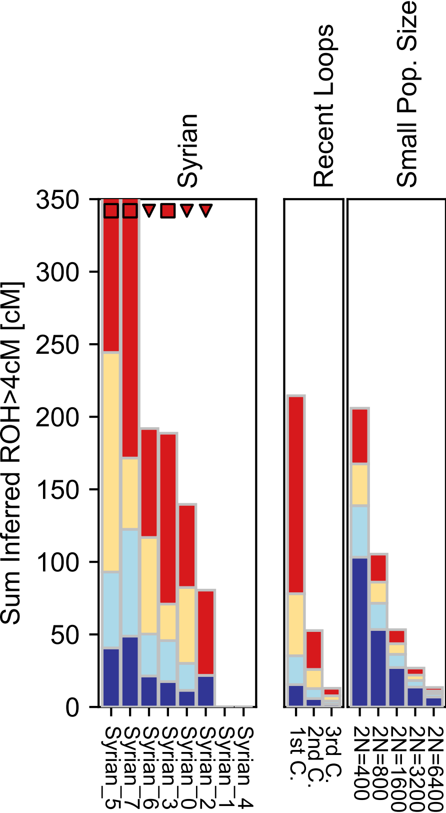

### Tajik_Pomiri.pdf

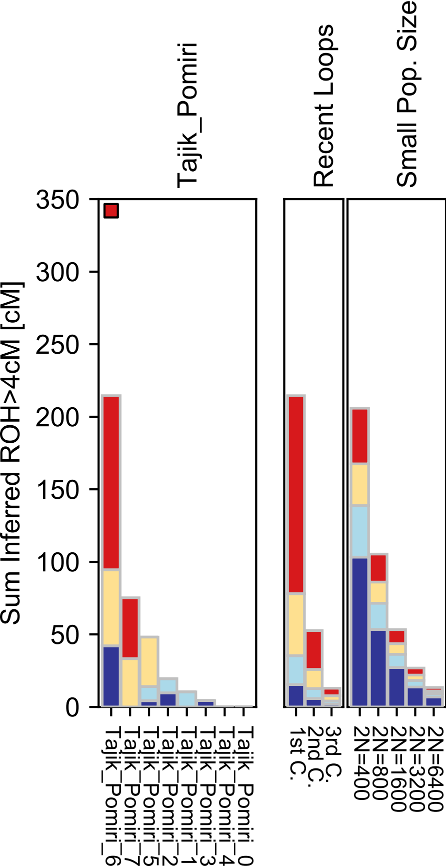

### Thai.pdf

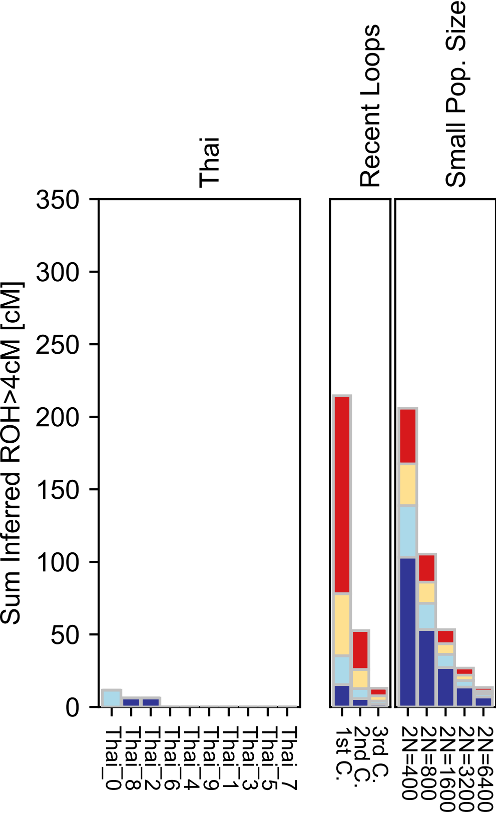

### Tlingit.pdf

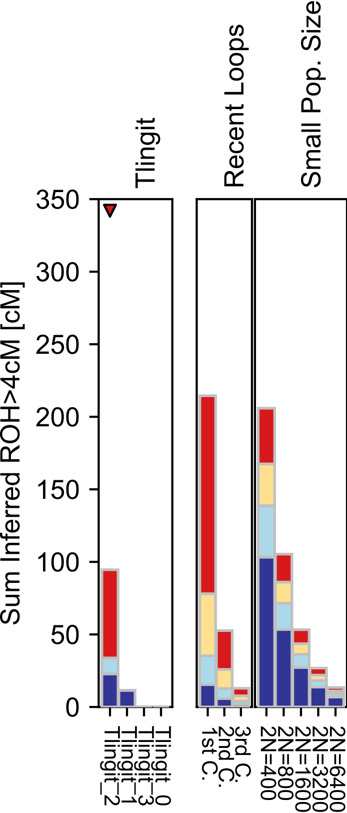

### Tu.pdf

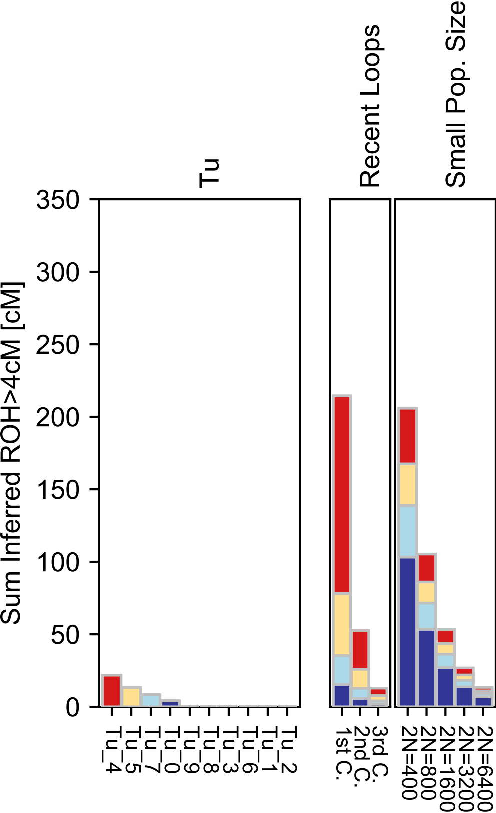

### Tubalar.pdf

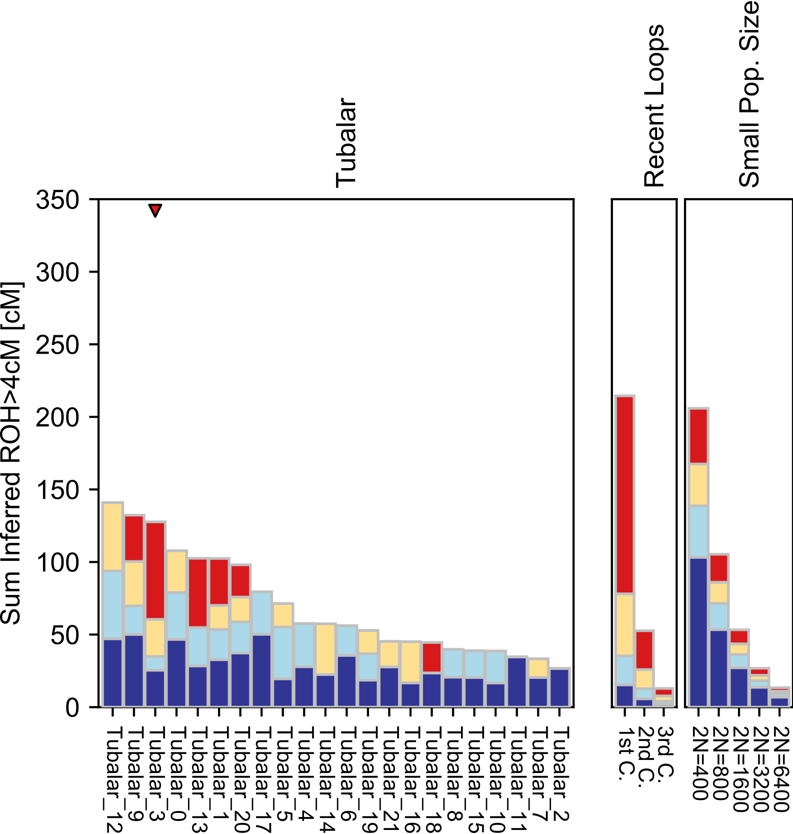

### Tujia.pdf

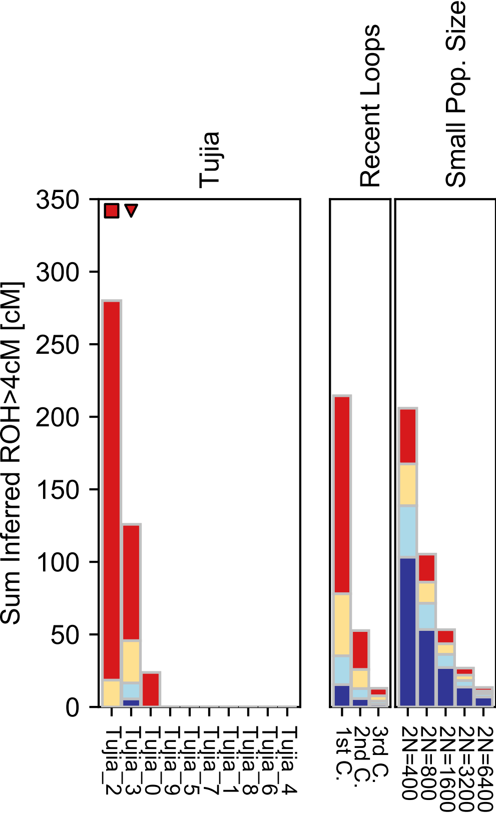

### Tunisian.pdf

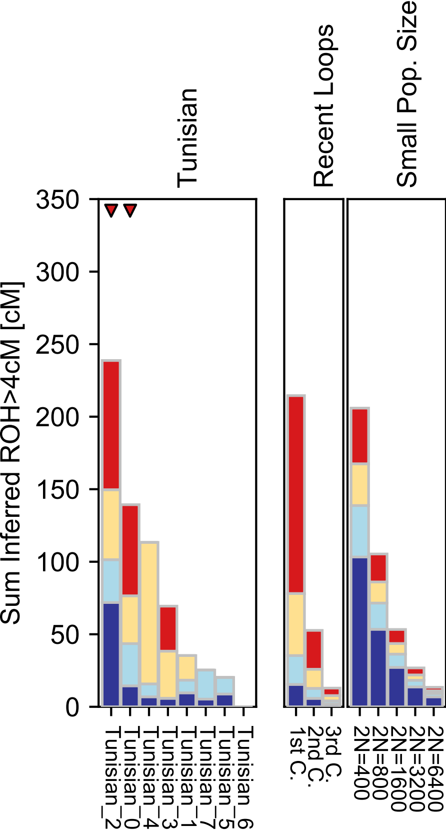

### Tunisian_Jew.pdf

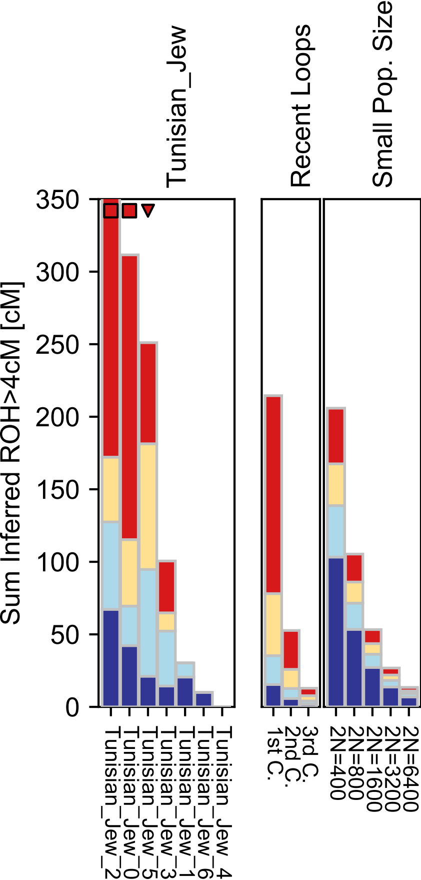

### Turkish-checkpoint.pdf

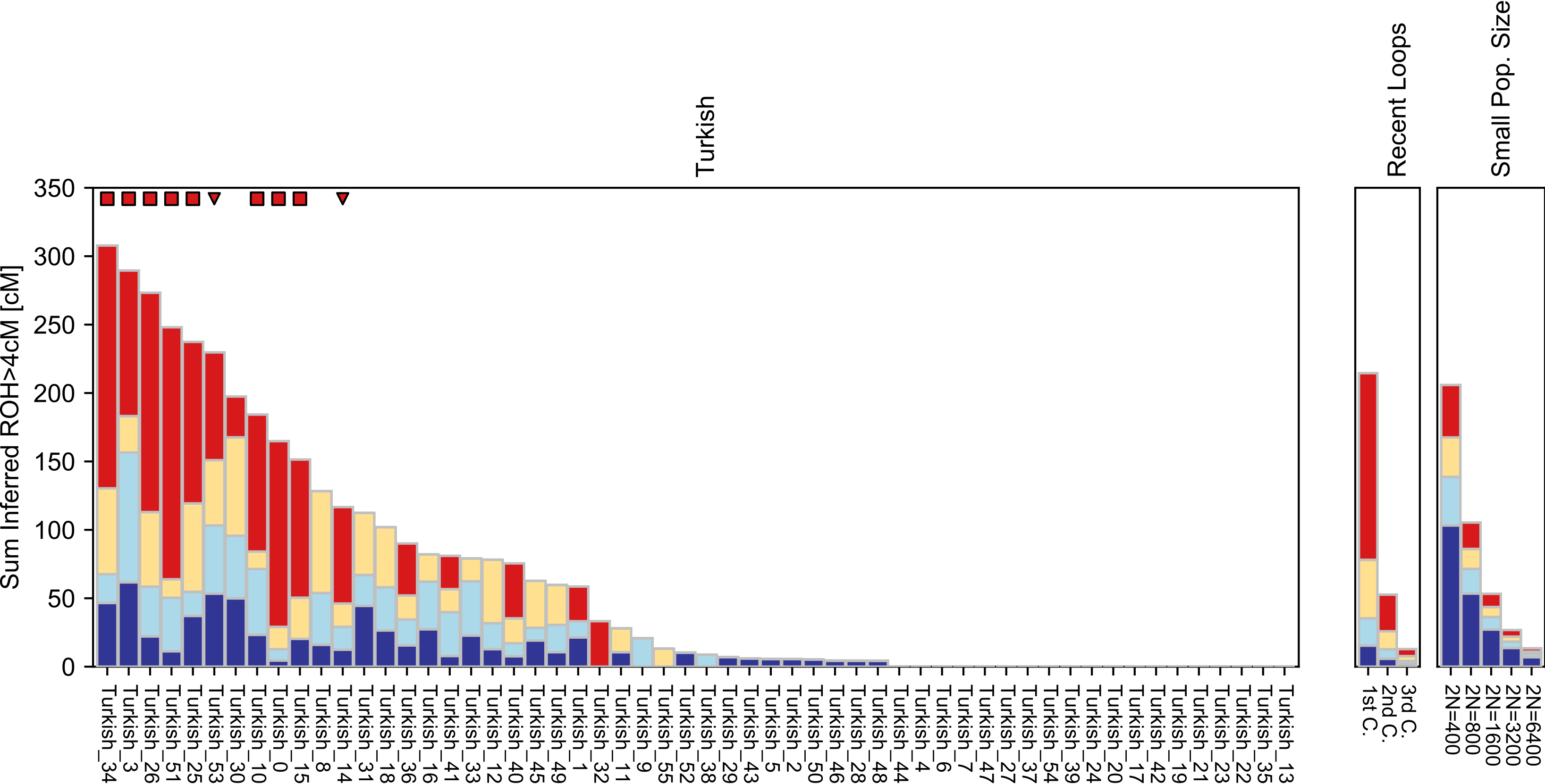

### Turkish.pdf

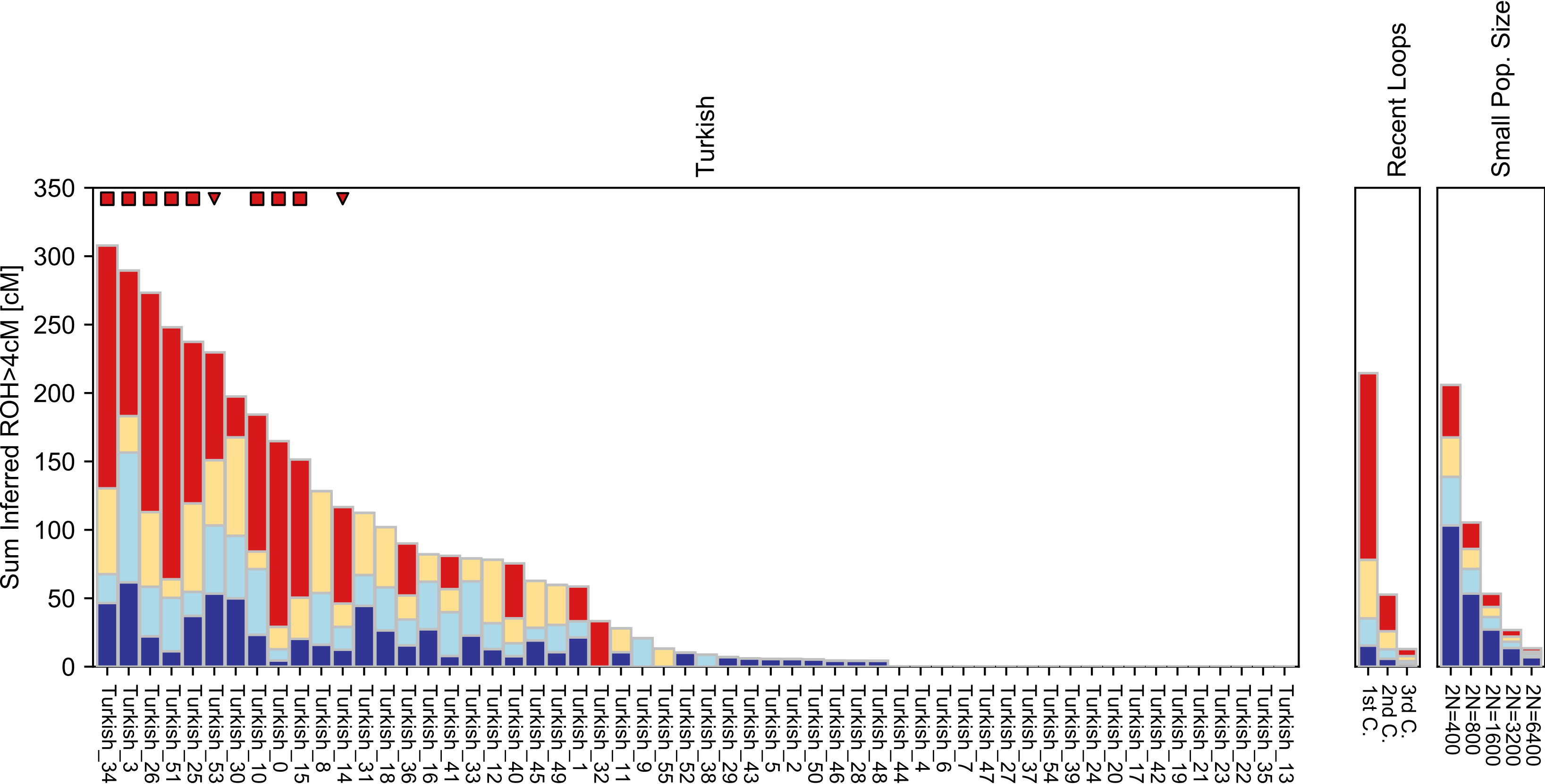

### Turkish_Jew.pdf

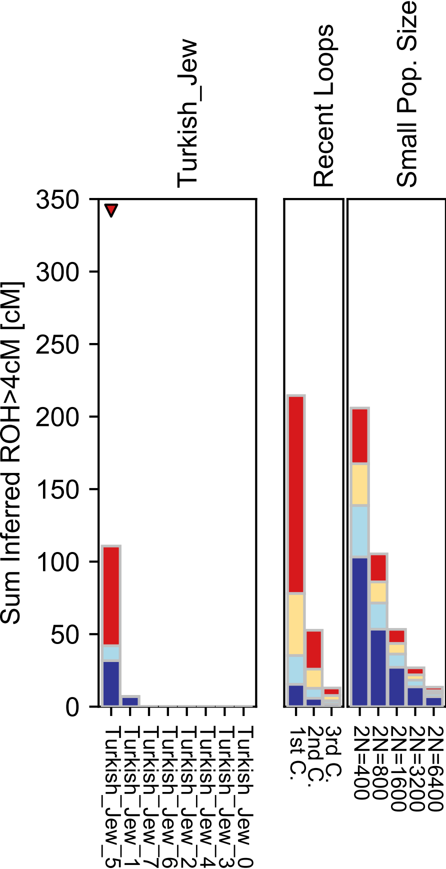

### Turkmen.pdf

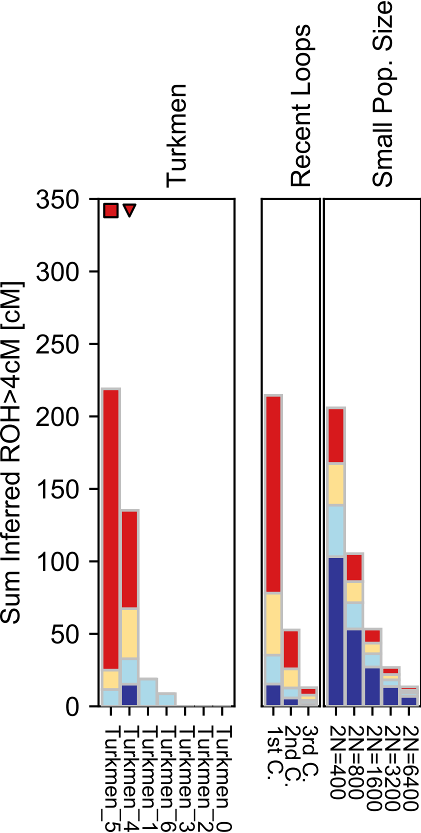

### Tuscan.pdf

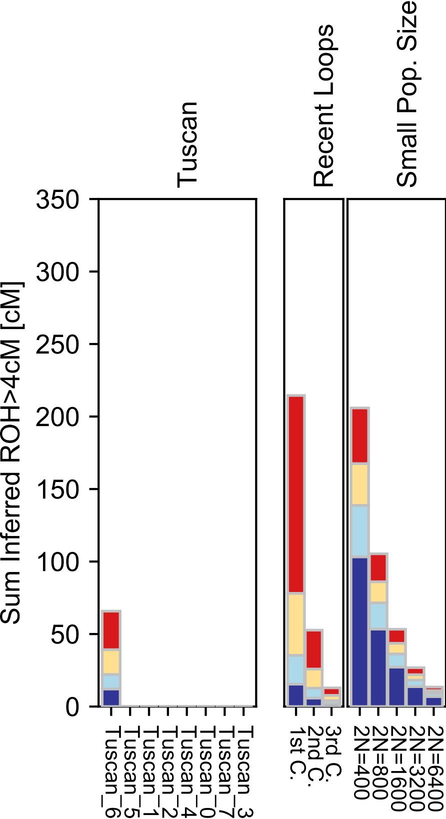

### Tuvinian.pdf

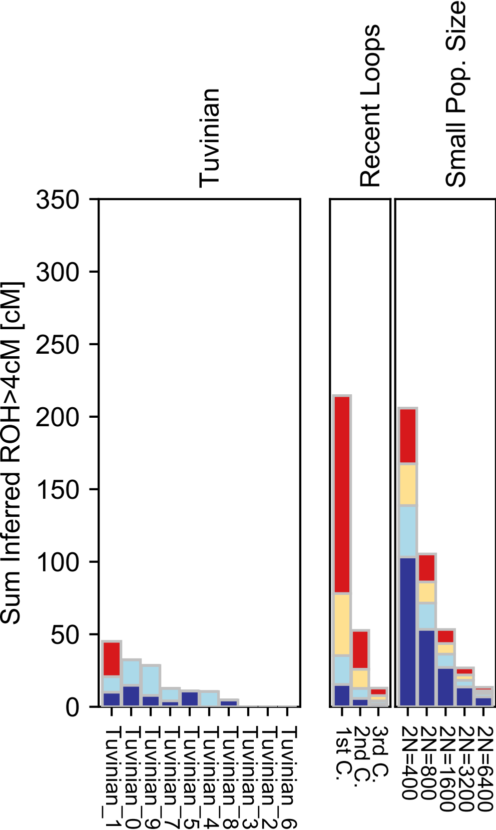

### Ukrainian.pdf

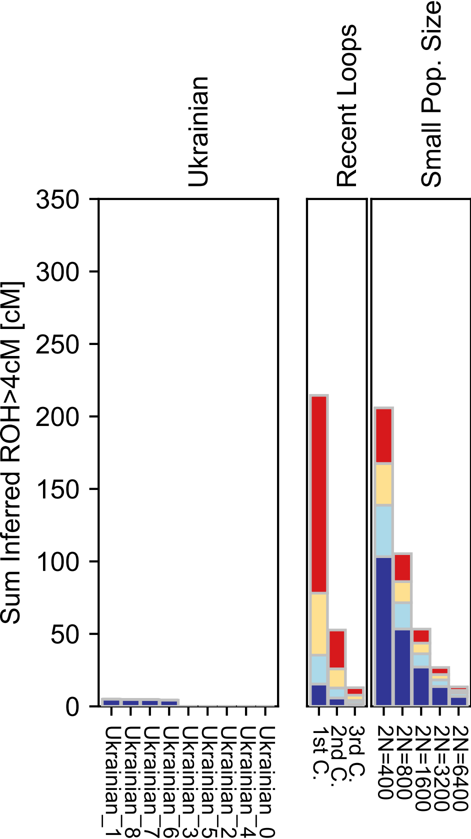

### Ulchi.pdf

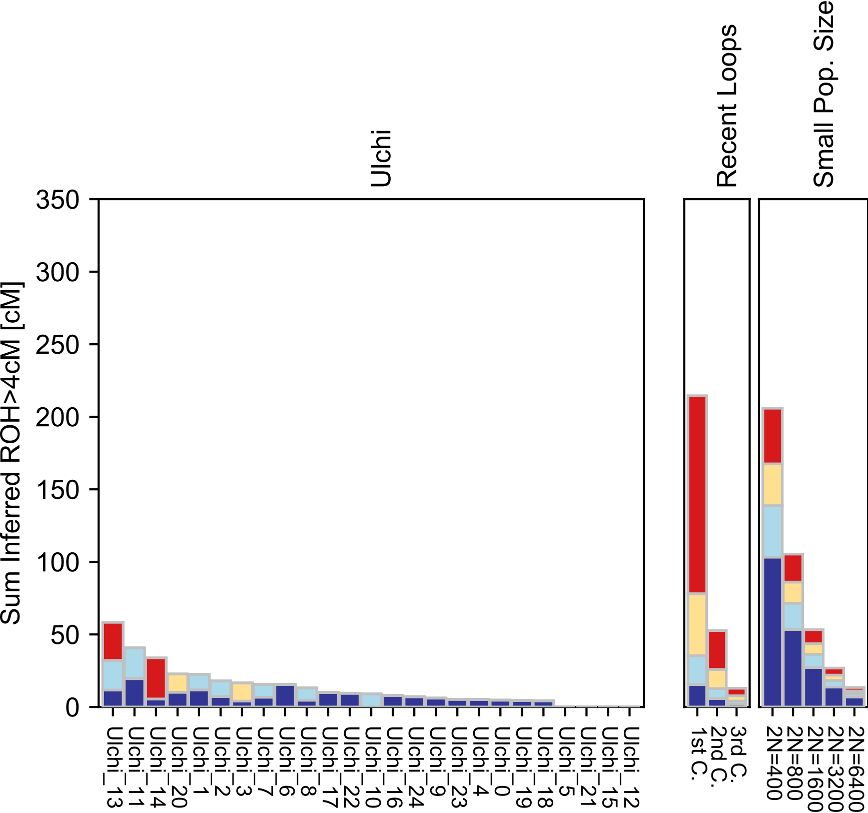

### Uygur.pdf

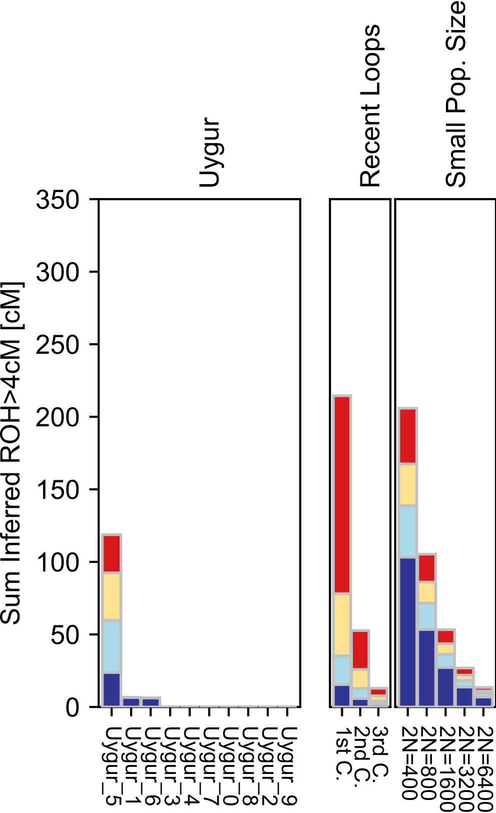

### Uzbek.pdf

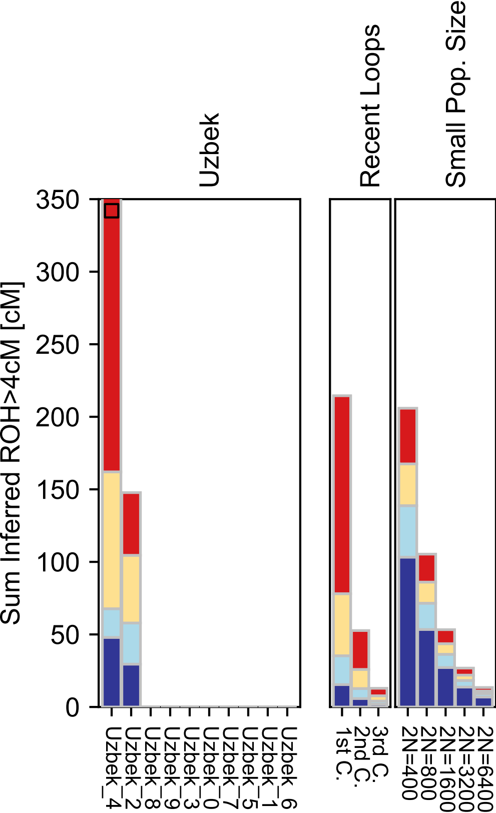

### Vanuatu.pdf

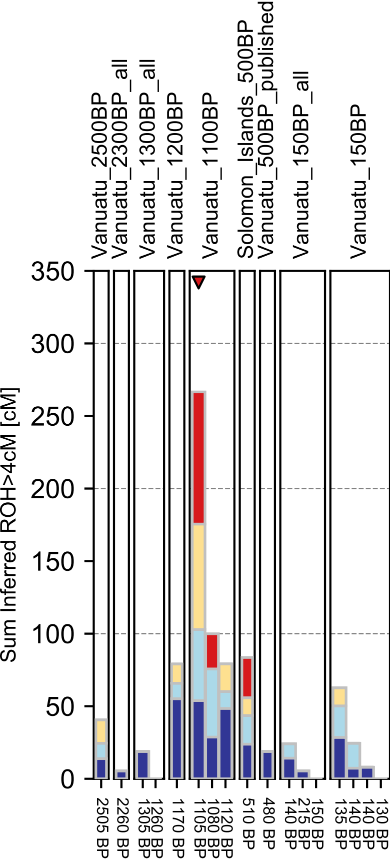

### Xibo.pdf

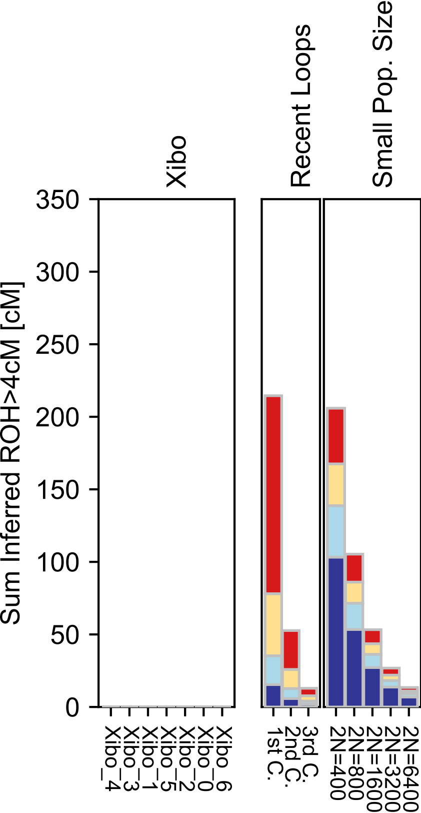

### Yakut.pdf

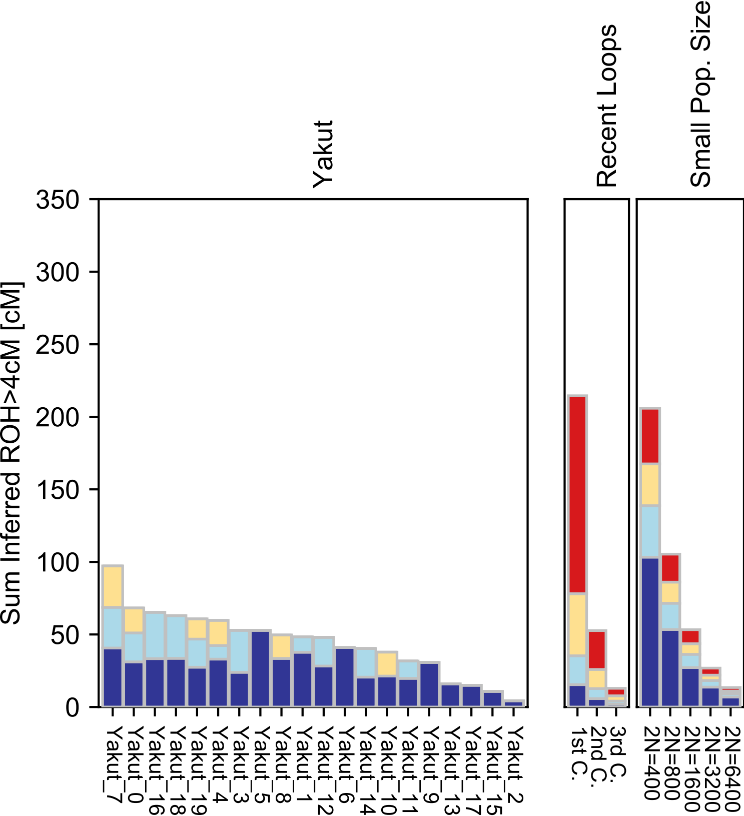

### Yemen.pdf

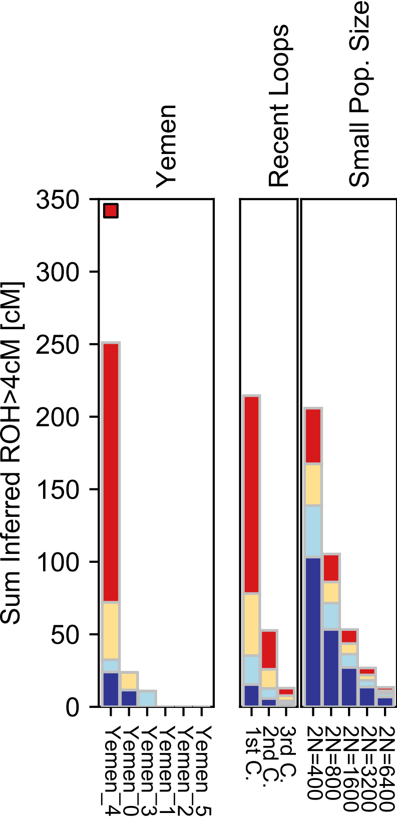

### Yemenite_Jew.pdf

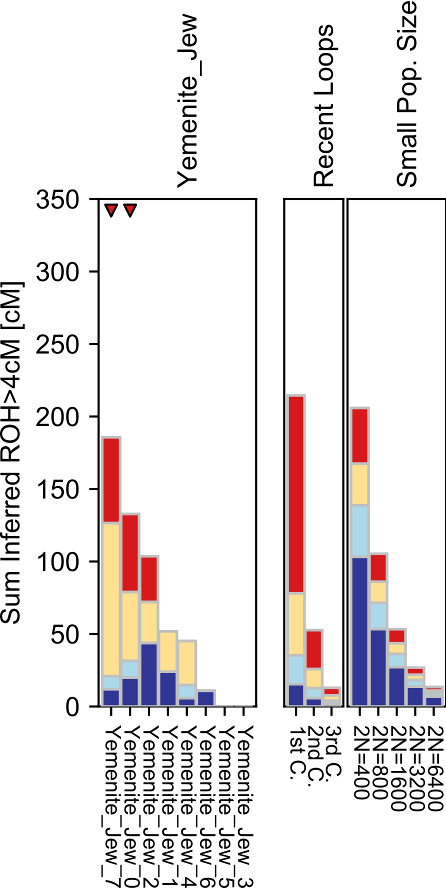

### Yi.pdf

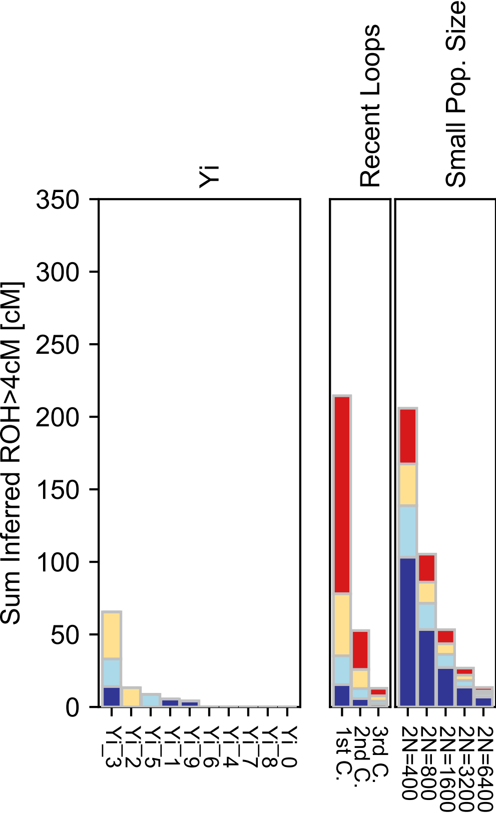

### Yoruba.pdf

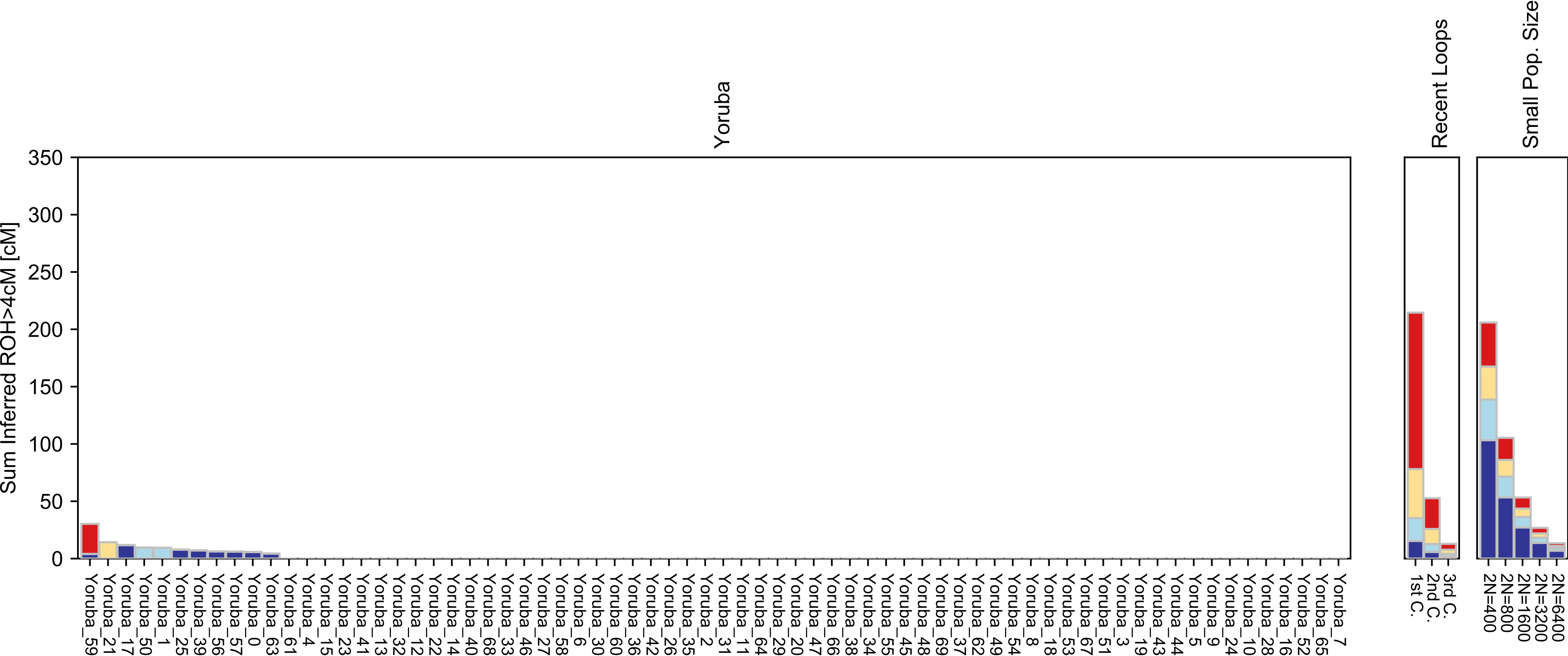

### Yukagir.pdf

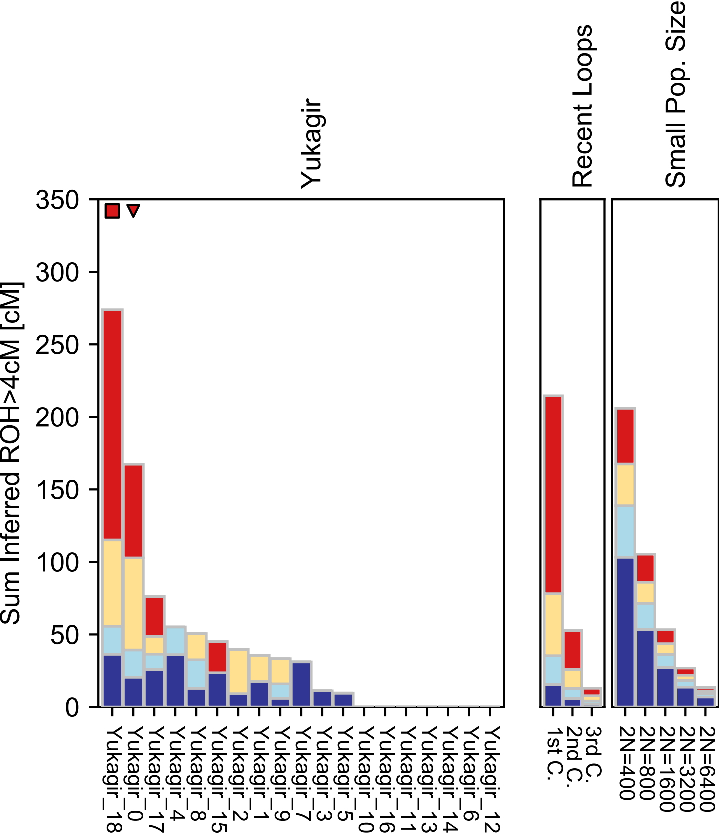

### Zapotec.pdf

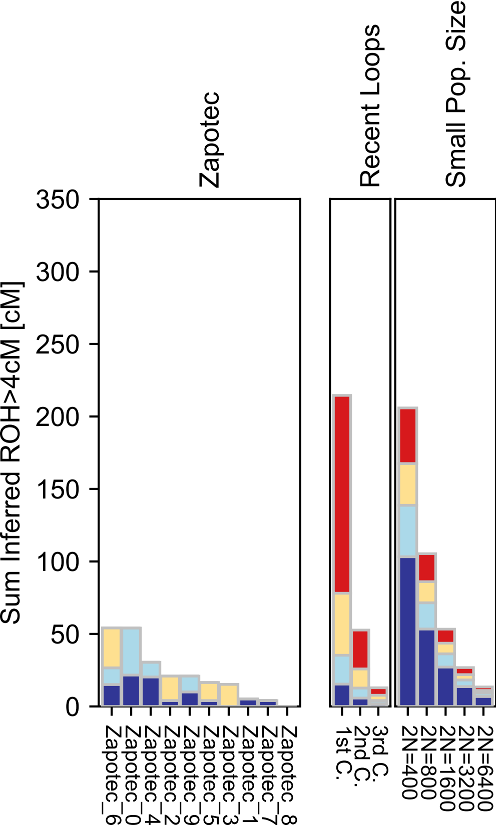
