## Supplementary figures and images for "Human Parental Relatedness through Time - Detecting Runs of Homozygosity in Ancient DNA"

### AA.pdf

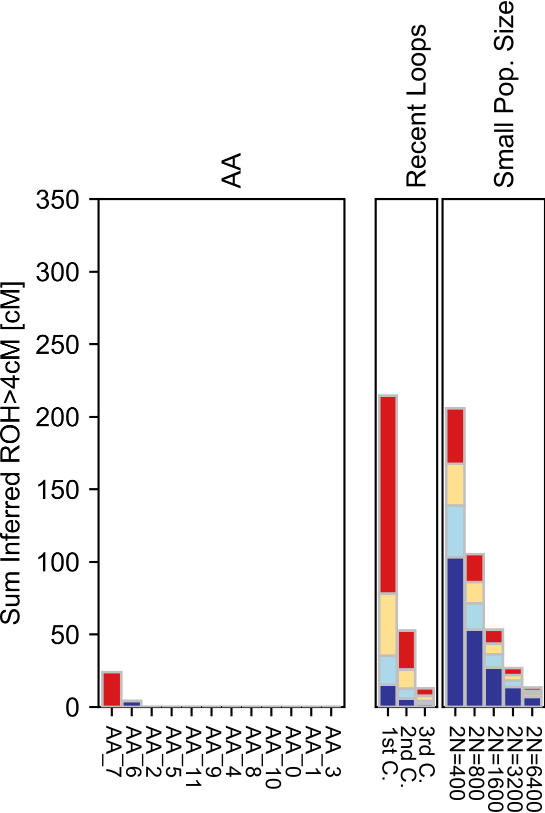

### Abkhasian.pdf

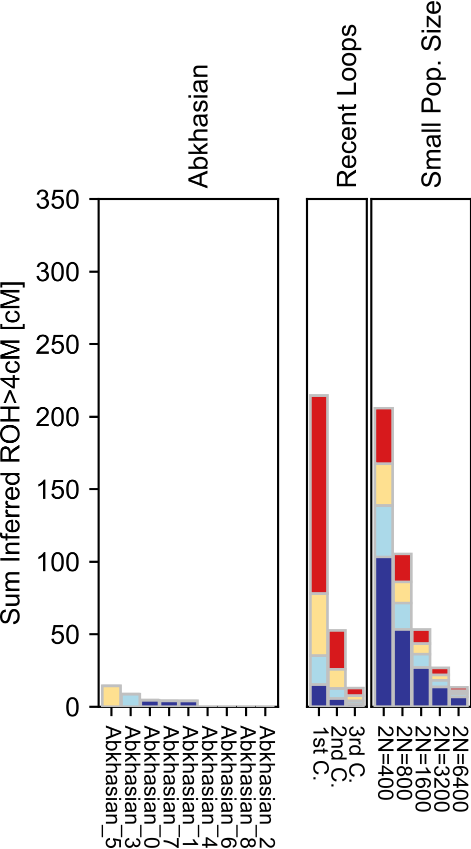

### Adygei.pdf

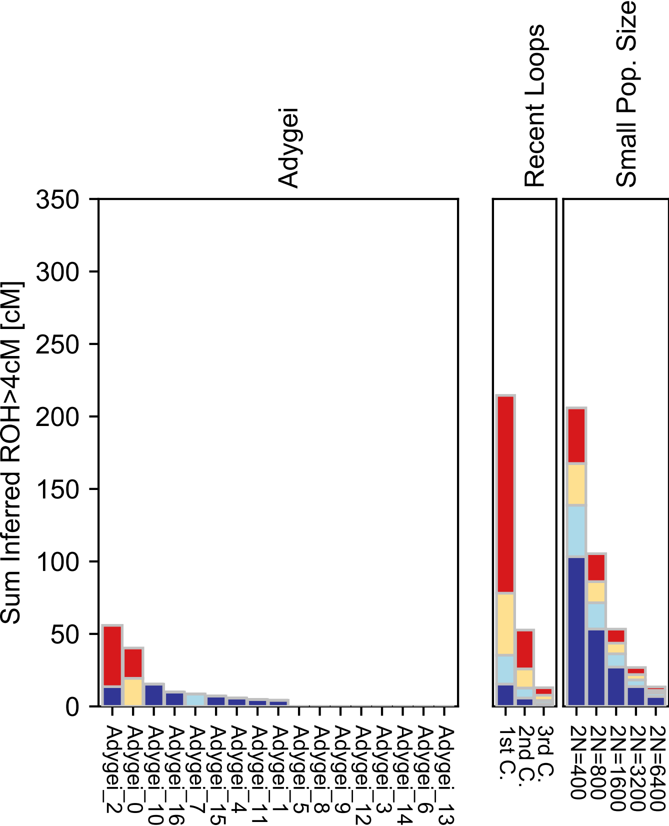

### Albanian.pdf

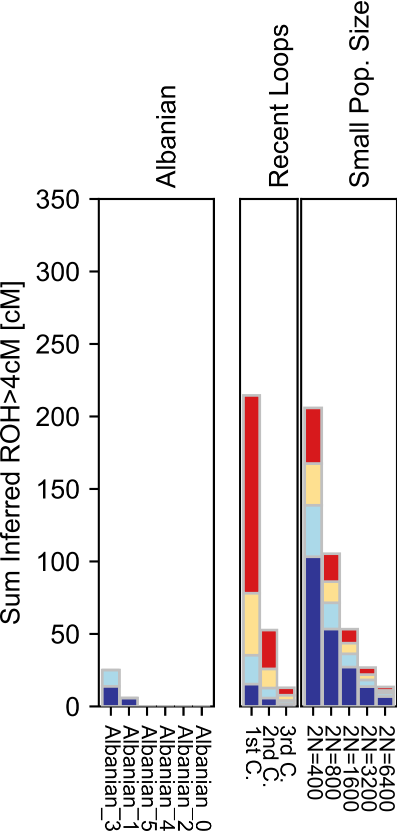

### Aleut.pdf

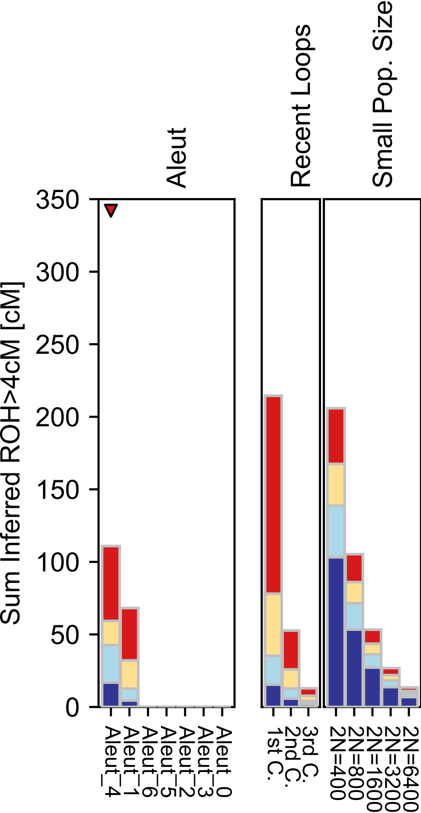

### Algerian.pdf

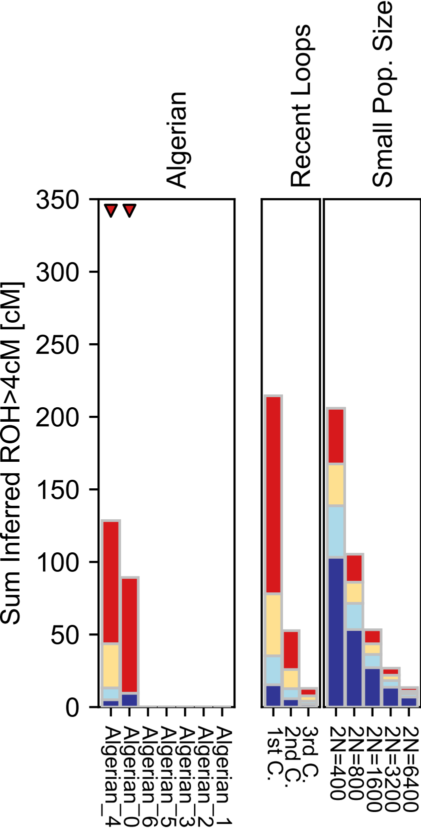

### Altaian.pdf

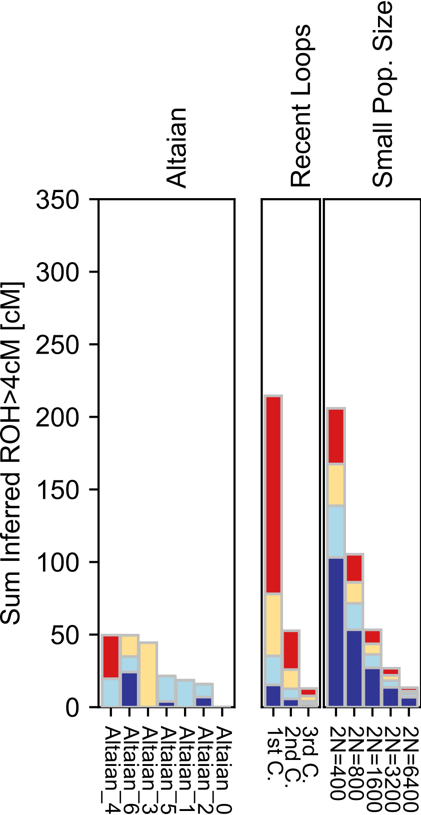

### Ami.pdf

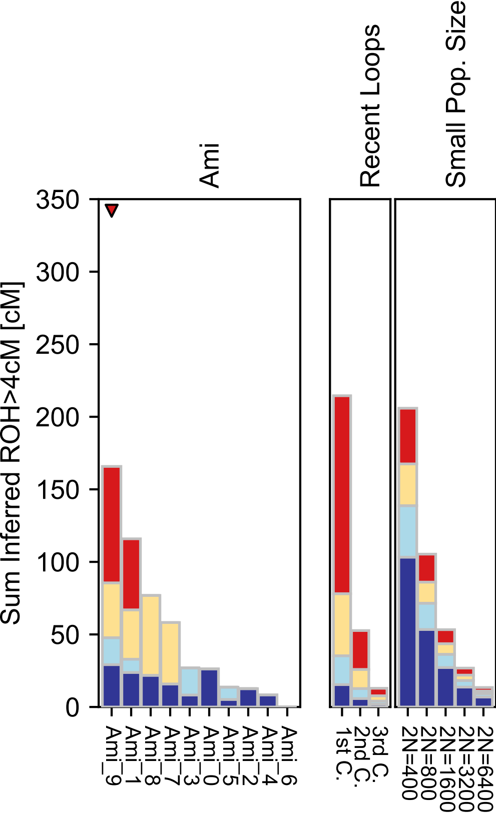

### Armenian.pdf

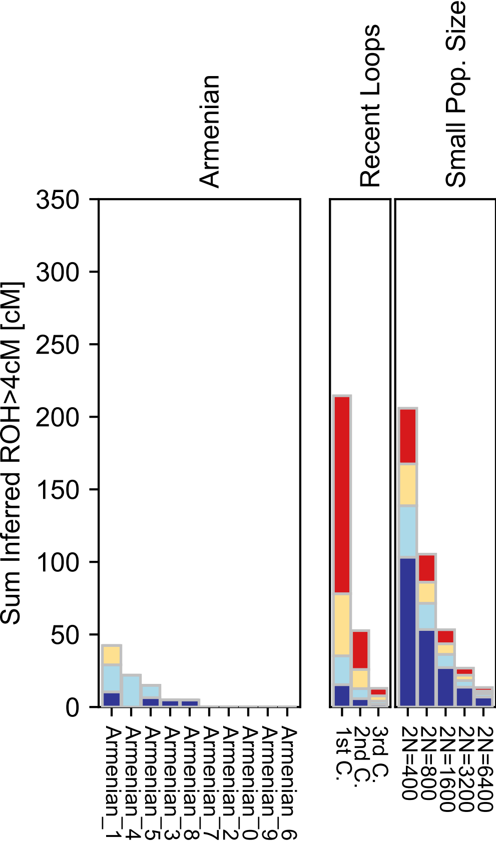

### Ashkenazi_Jew.pdf

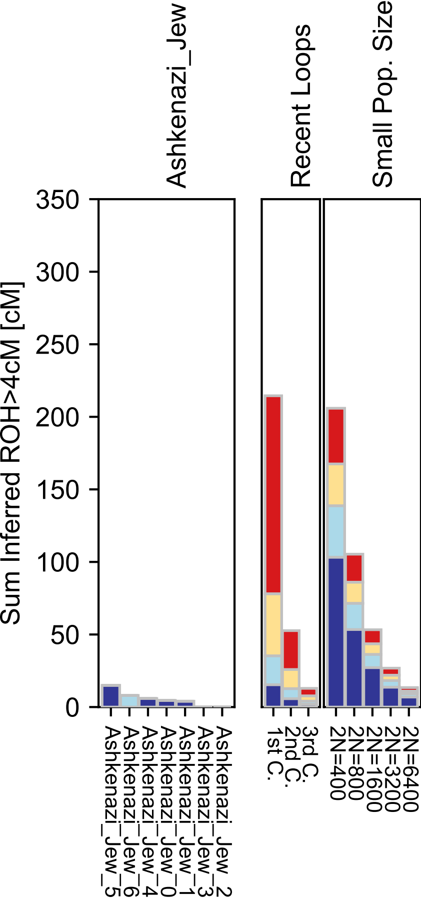

### Atayal.pdf

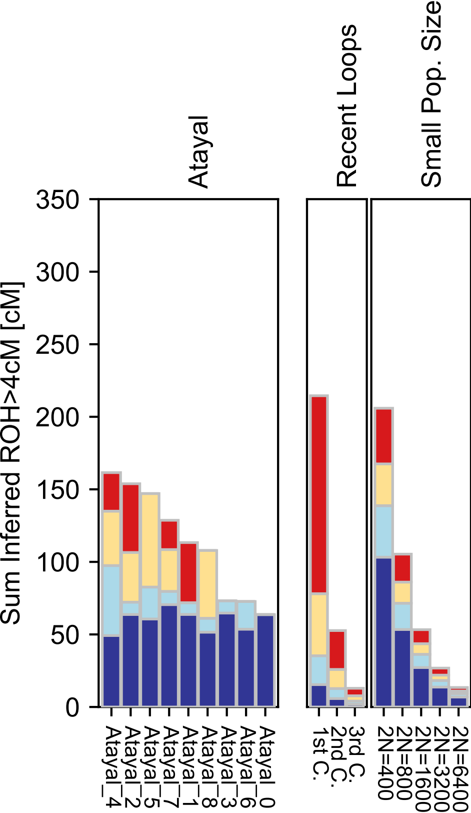

### Australian.pdf

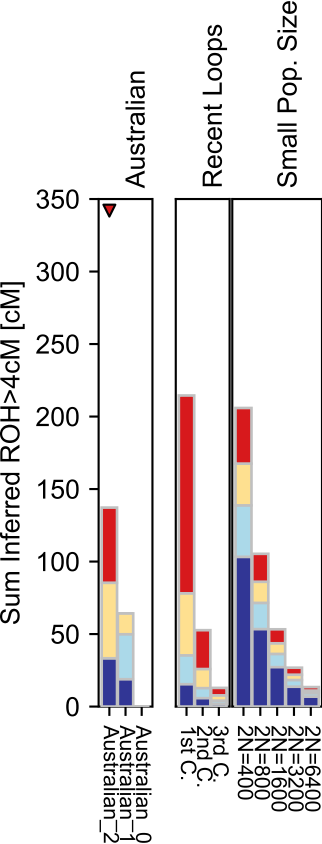

### Balkar.pdf

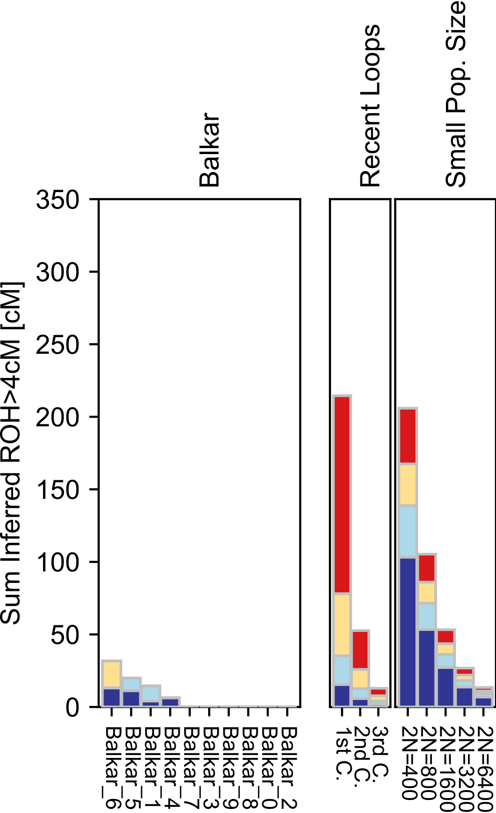

### Balochi.pdf

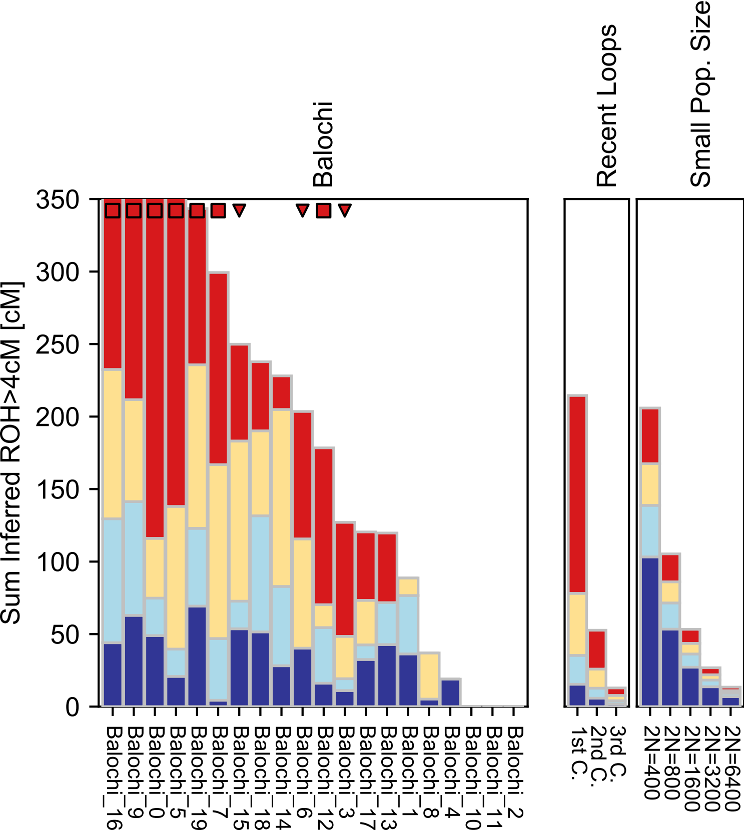

### BantuKenya.pdf

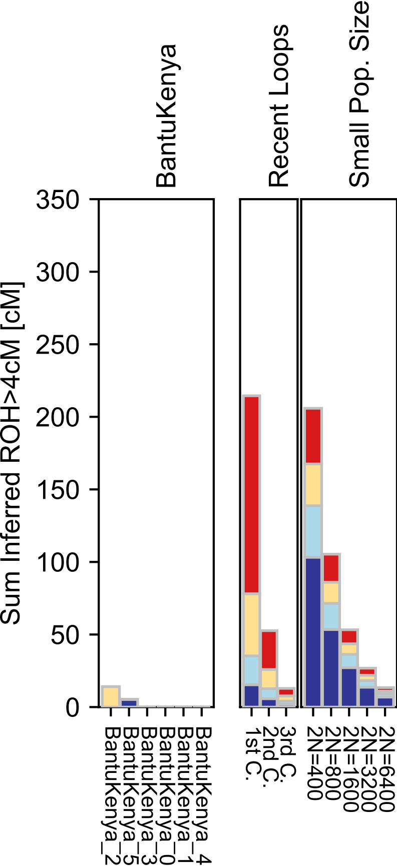

### BantuSA.pdf

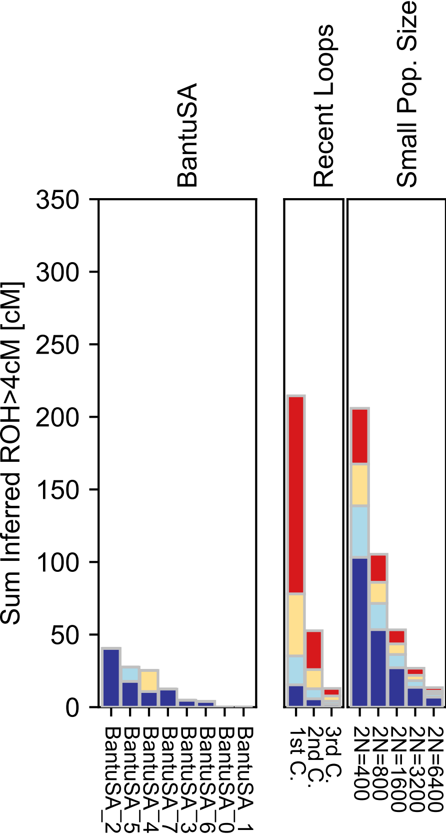

### Basque.pdf

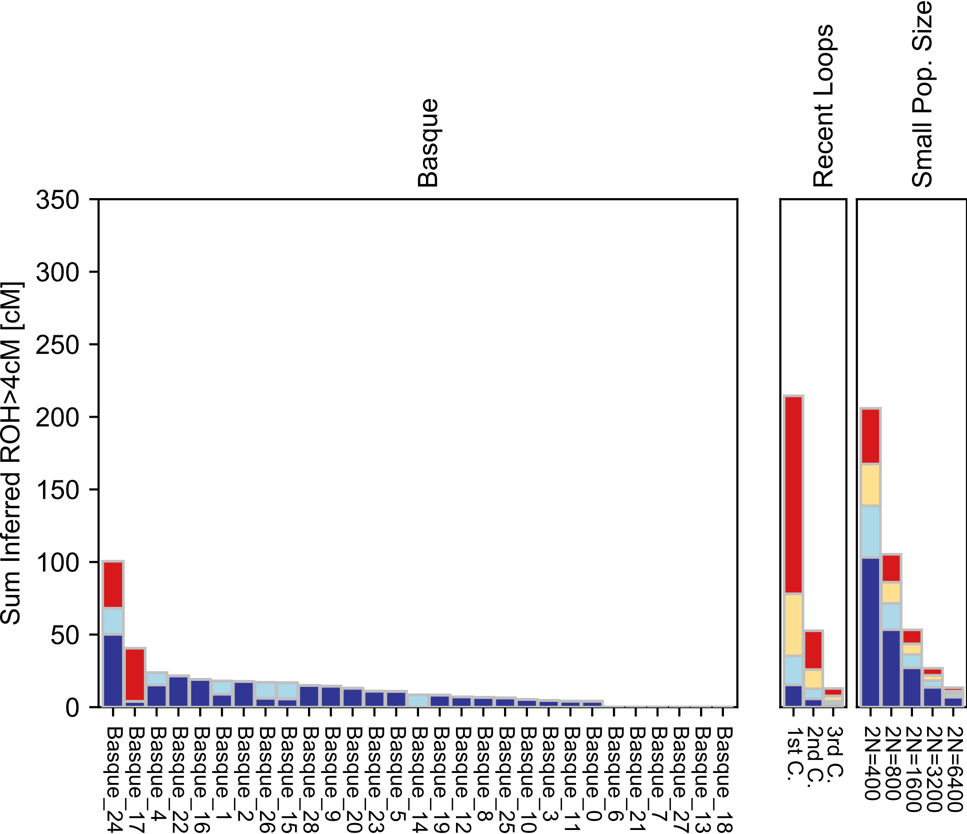

### BedouinA.pdf

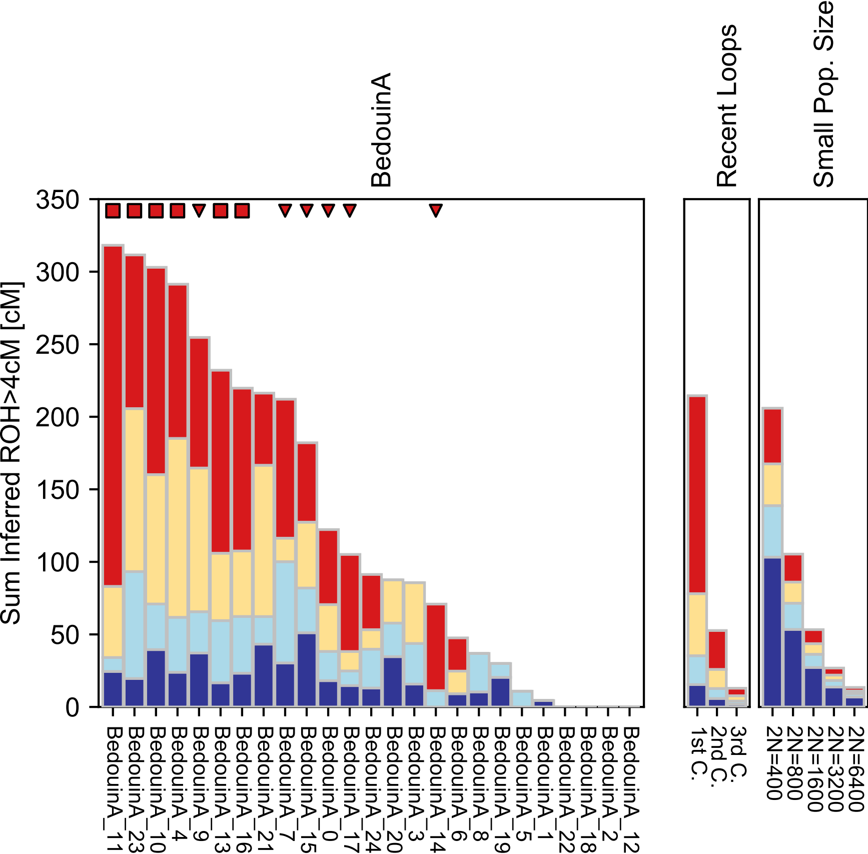

### BedouinB.pdf

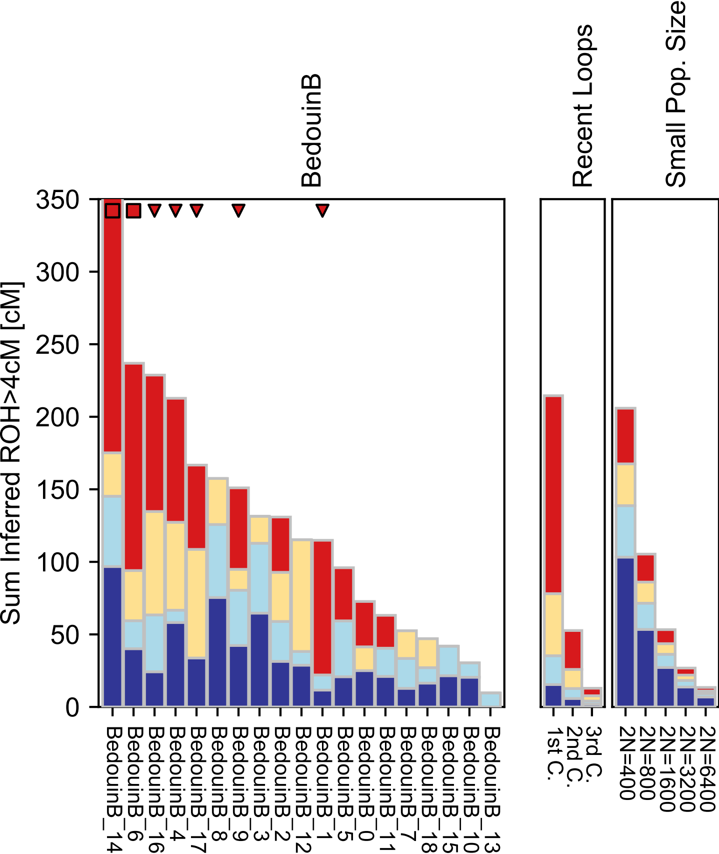

### Belarusian.pdf

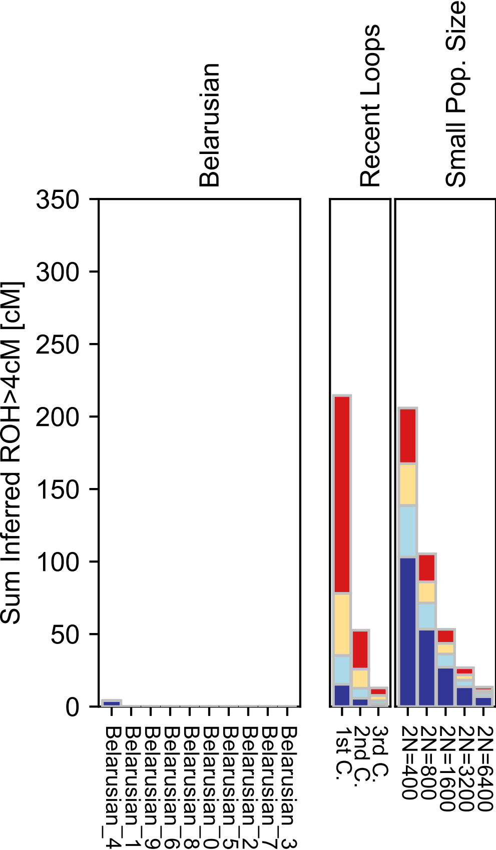

### Bengali.pdf

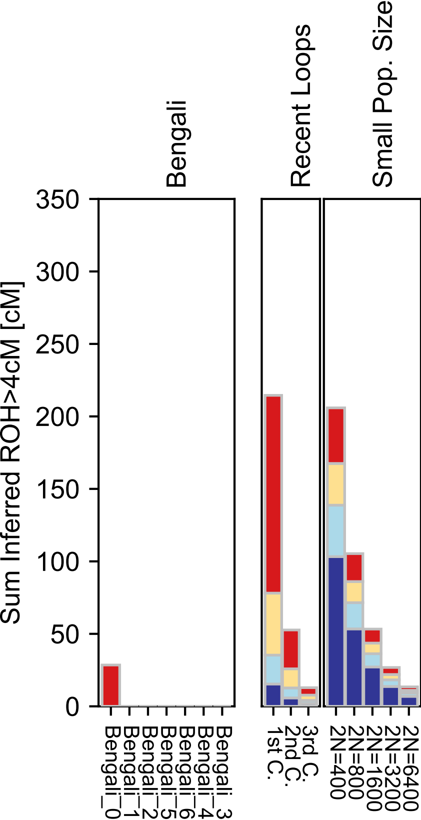

### Bergamo.pdf

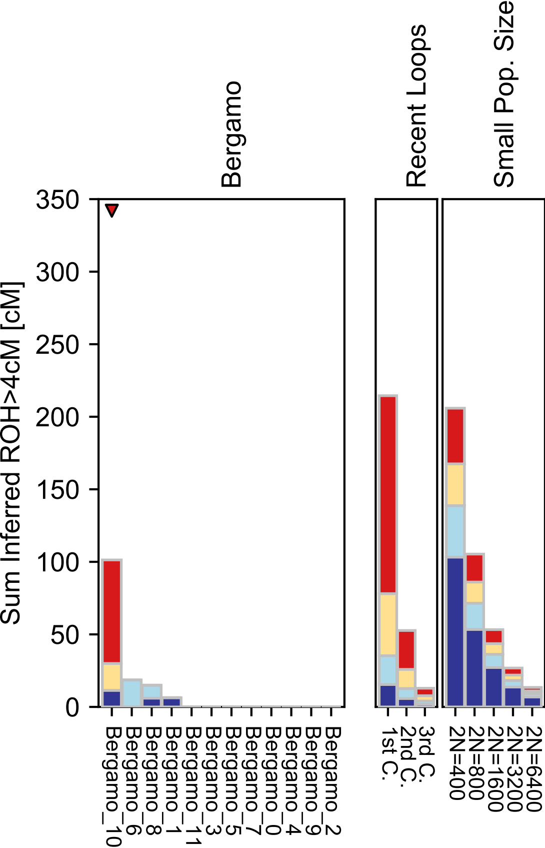

### Biaka.pdf

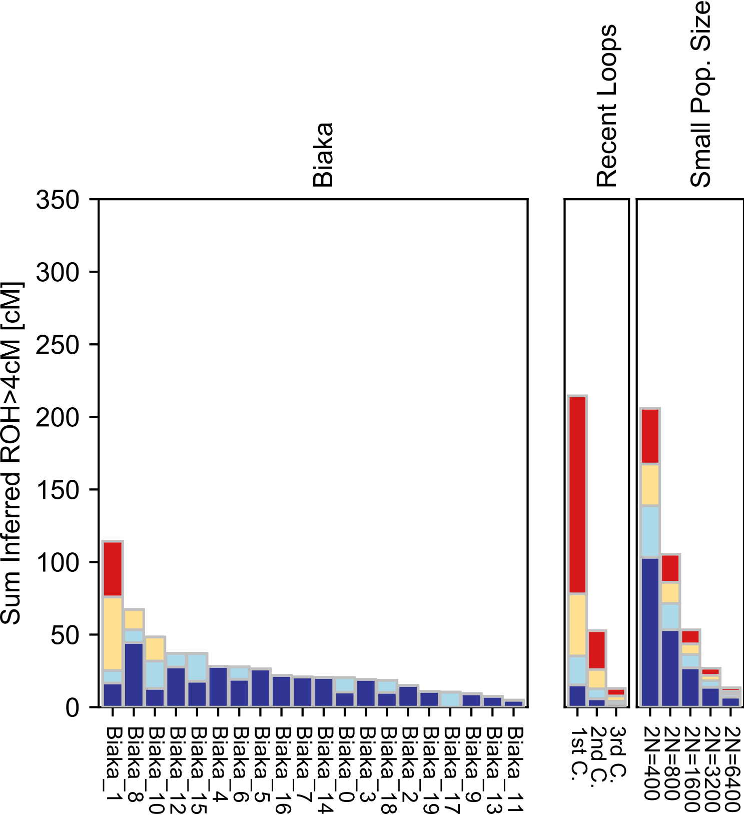

### Bolivian.pdf

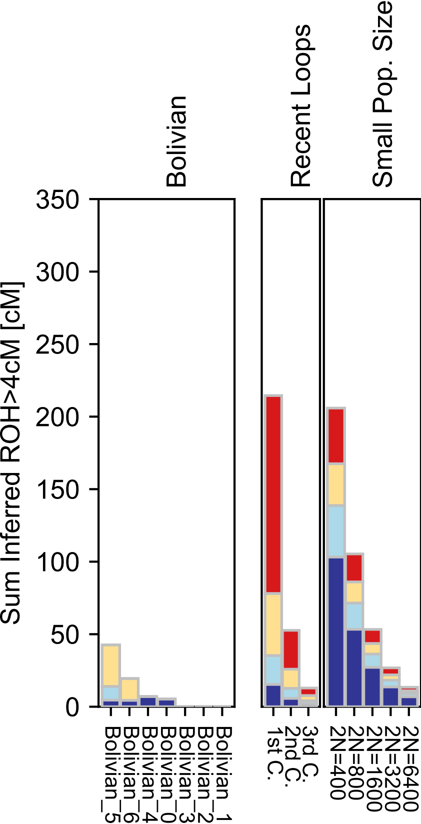

### Bougainville.pdf

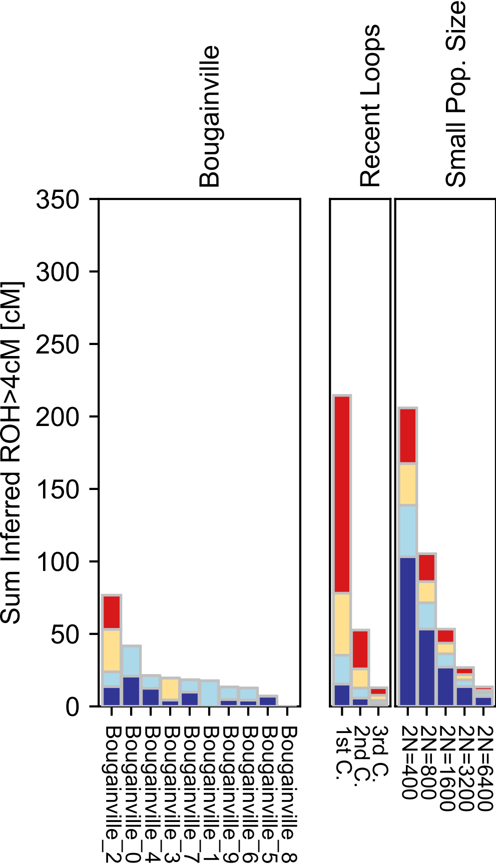

### Brahui.pdf

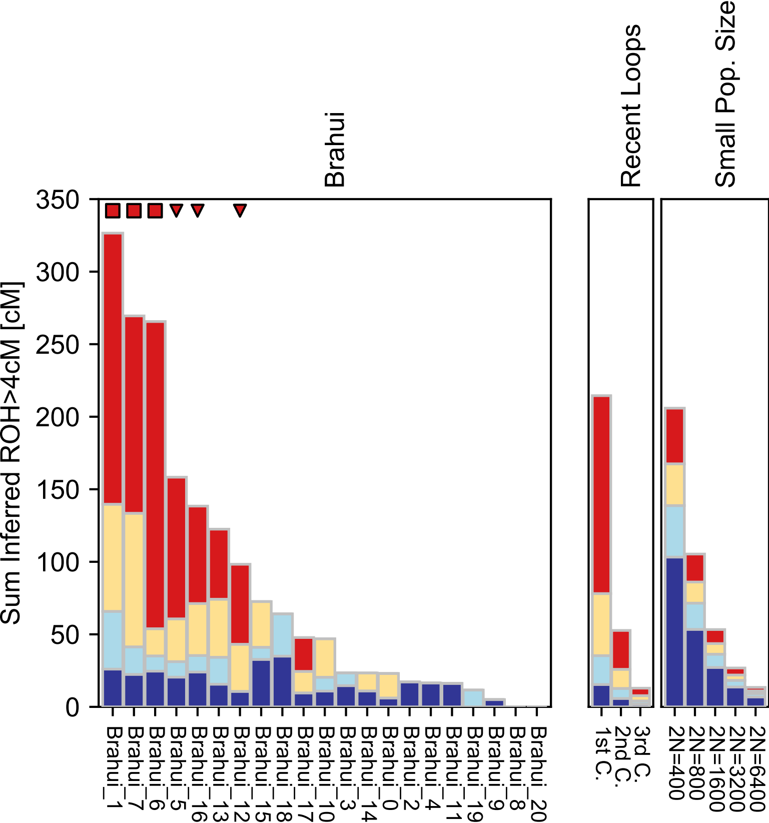

### Bulgarian.pdf

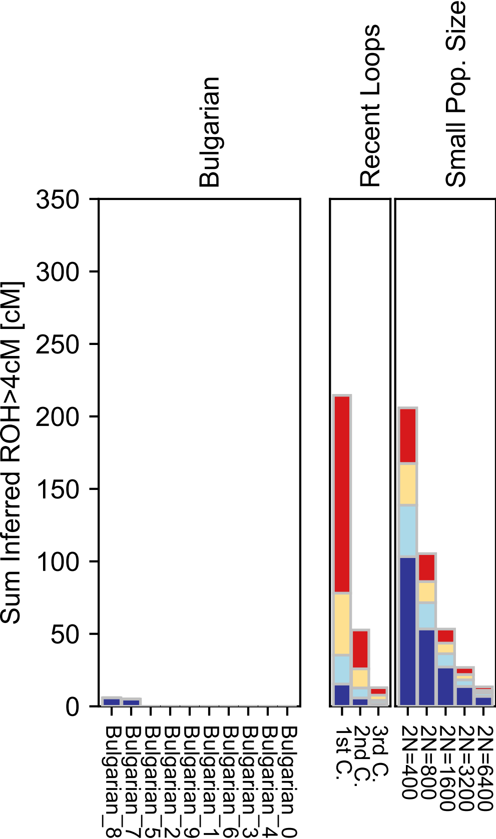

### Burusho.pdf

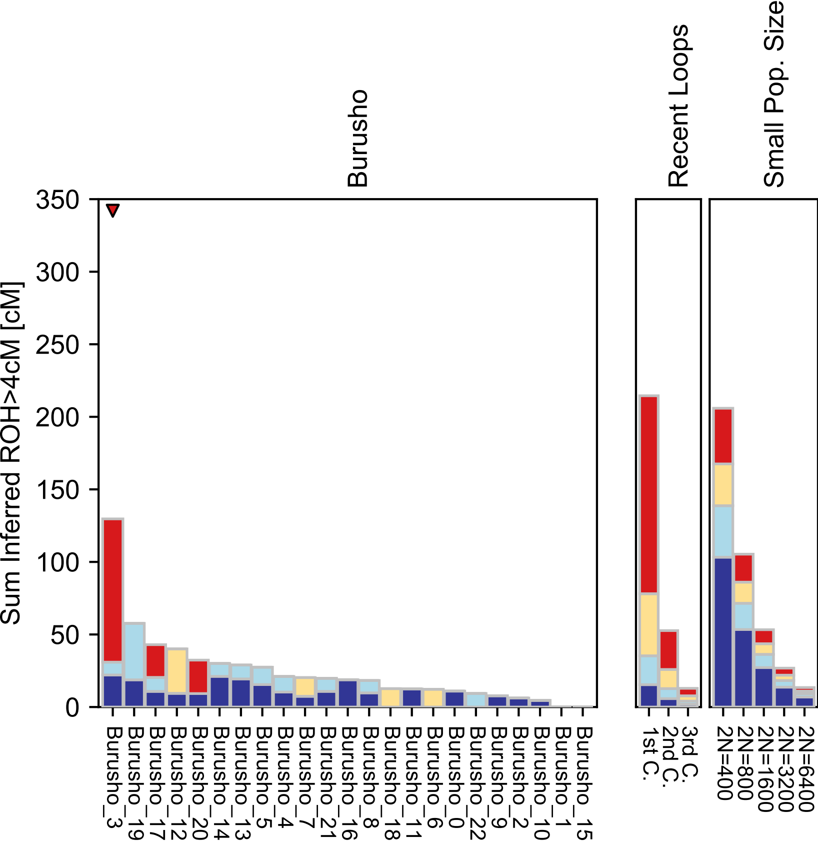

### Cambodian.pdf

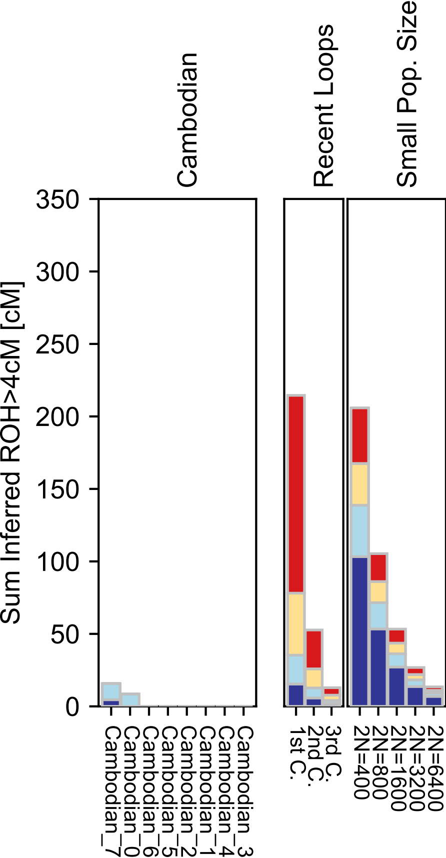

### Canary_Islanders.pdf

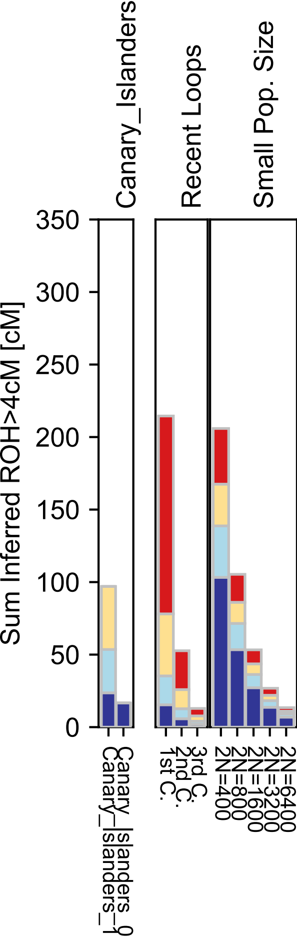

### Chechen.pdf

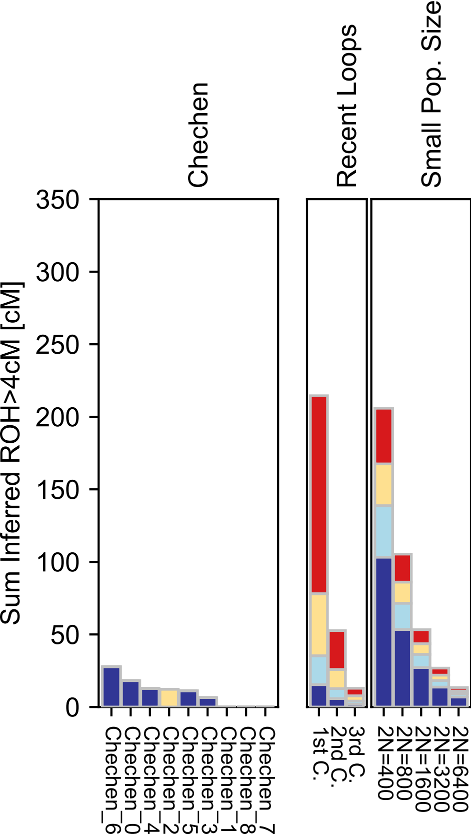

### Chukchi.pdf

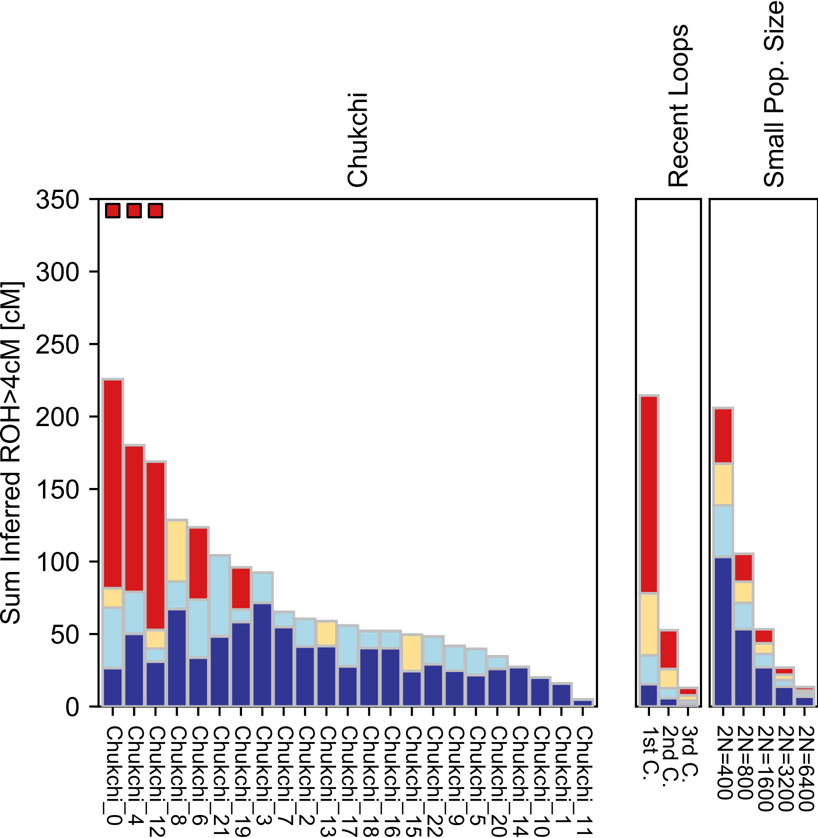

### Chuvash.pdf

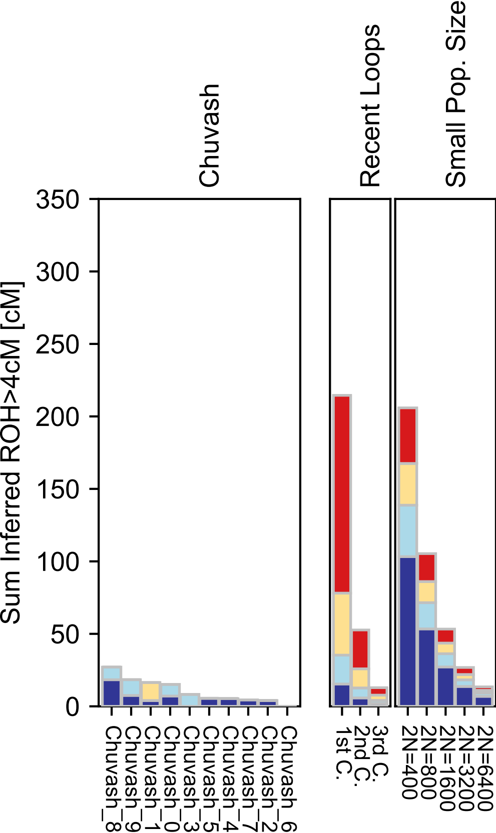

### Cochin_Jew.pdf

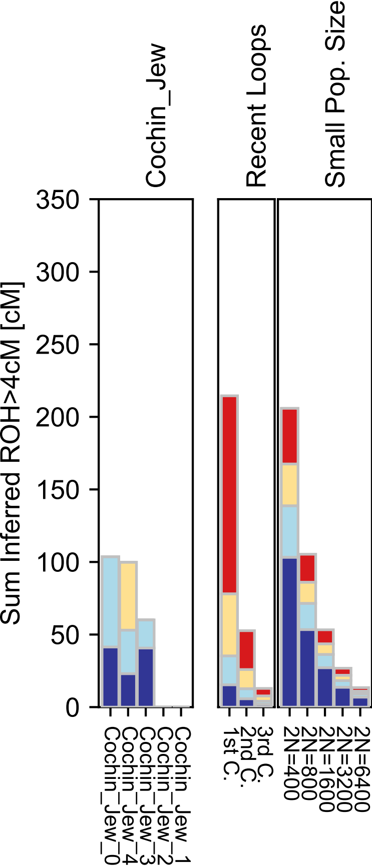

### Croatian.pdf

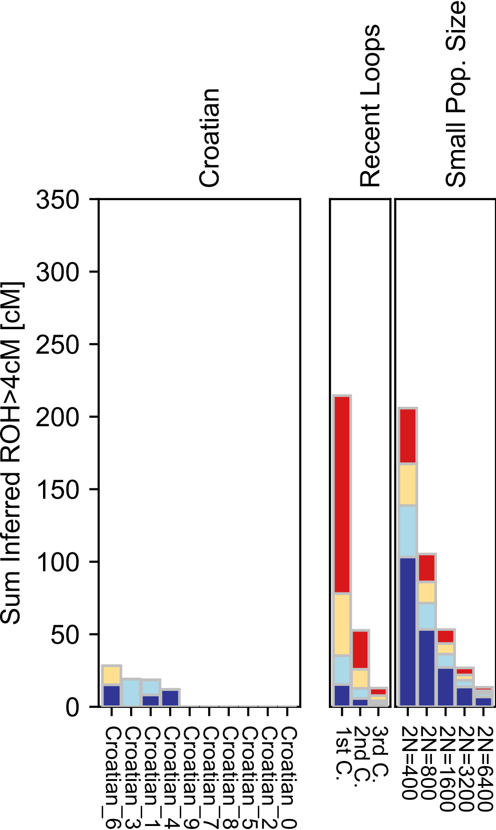

### Cypriot.pdf

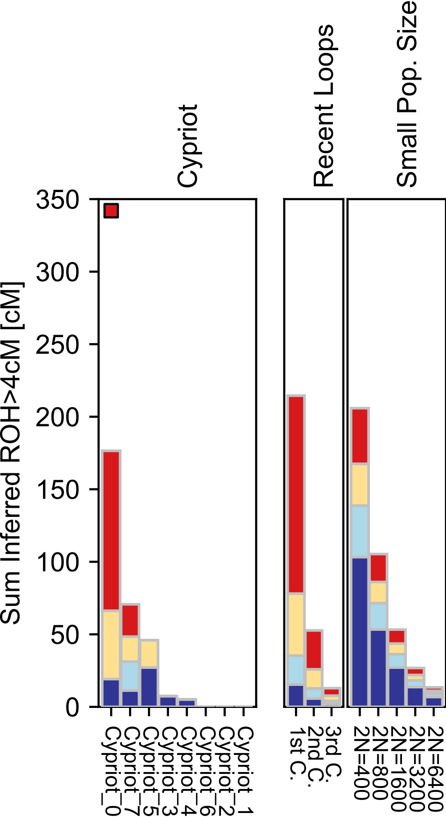

### Czech.pdf

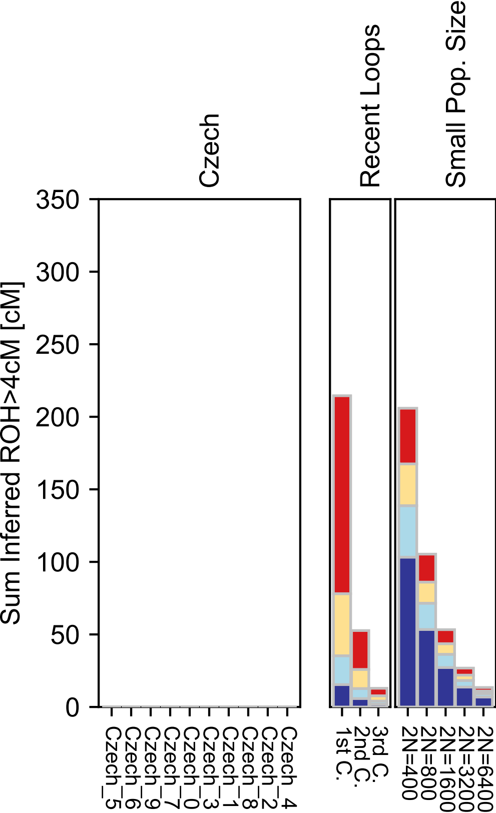

### Dai.pdf

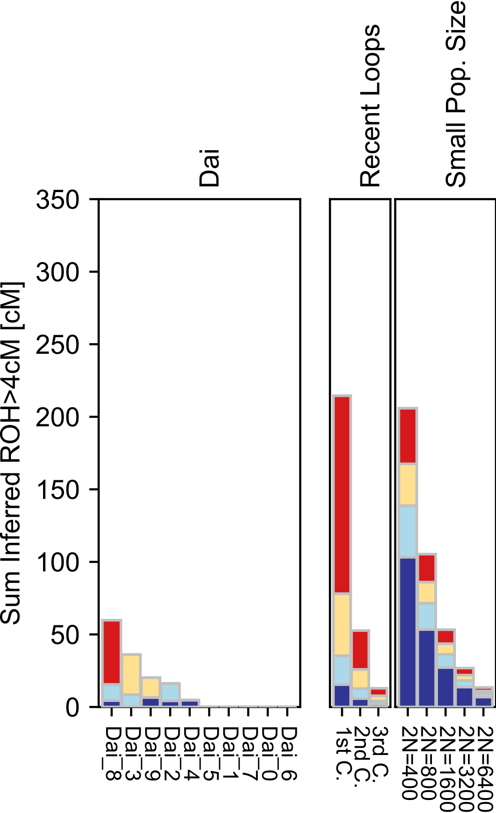

### Datog.pdf

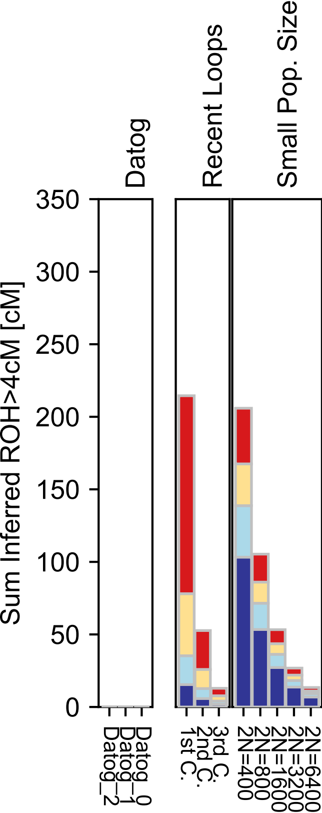

### Daur.pdf

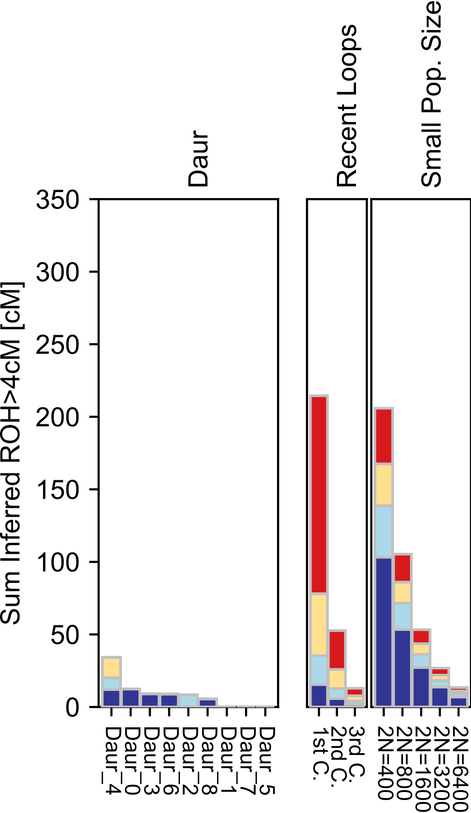

### Dolgan.pdf

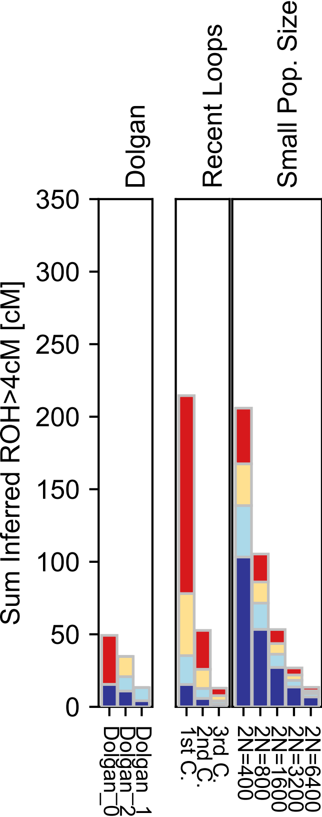

### Druze.pdf

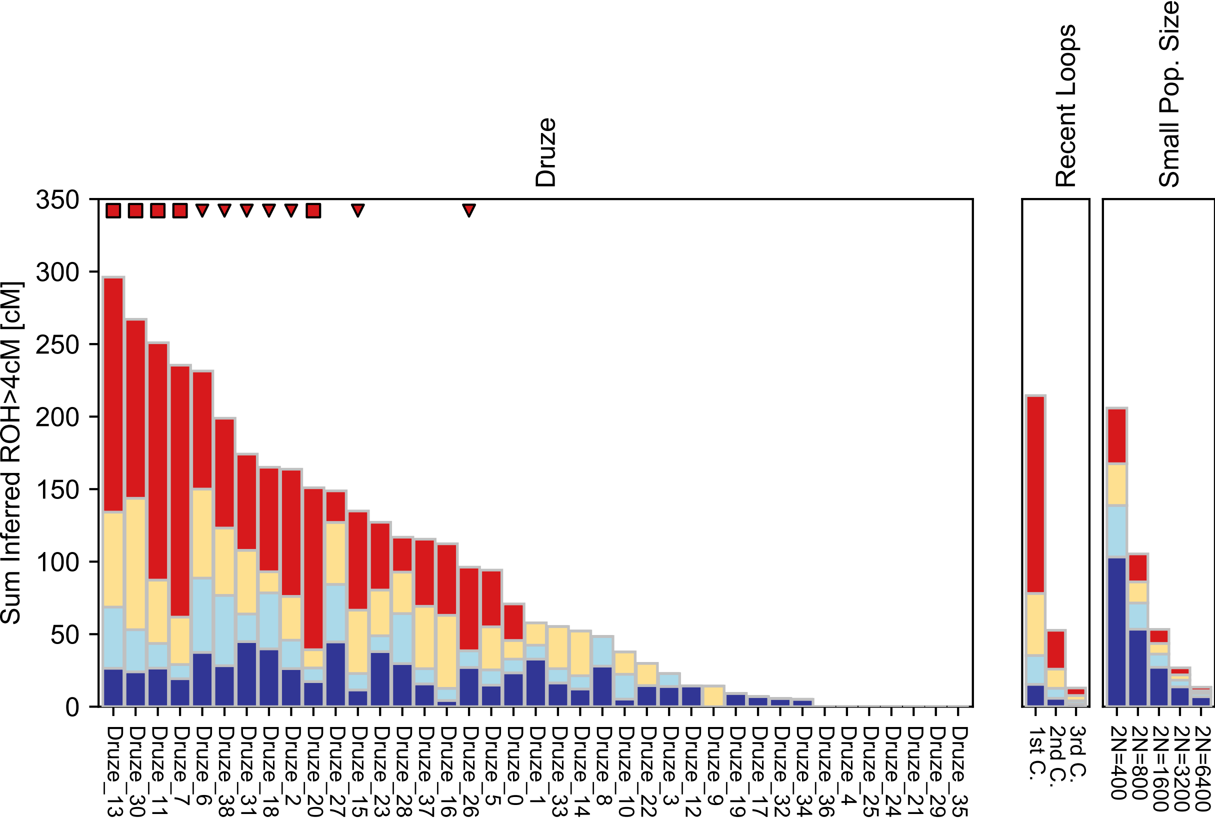

### Egyptian.pdf

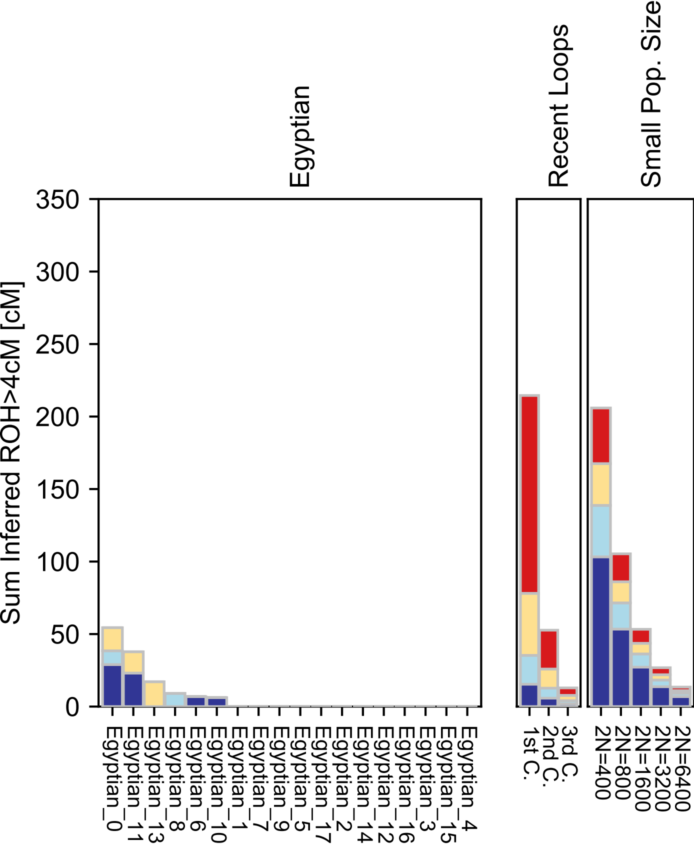

### English.pdf

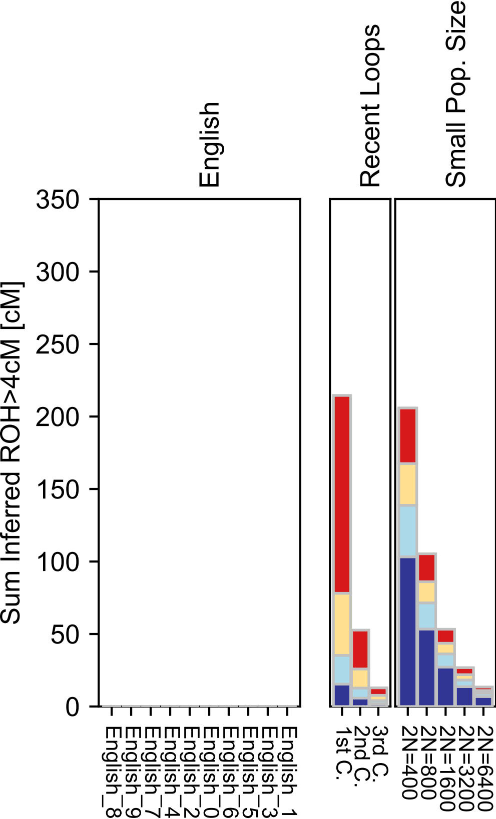

### Esan.pdf

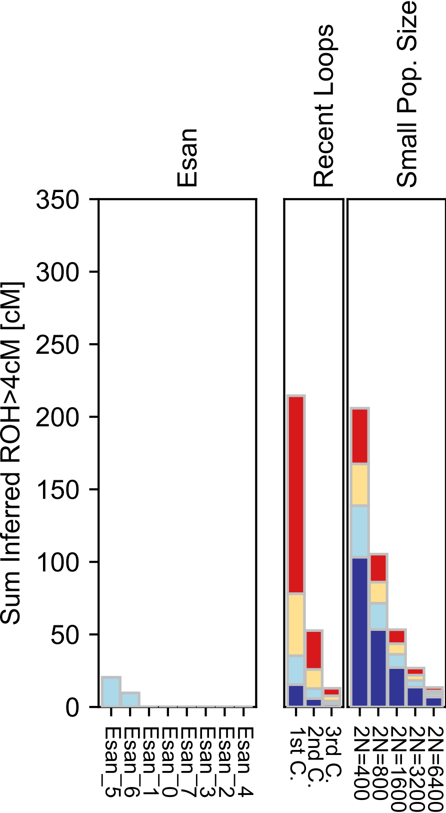

### Eskimo.pdf

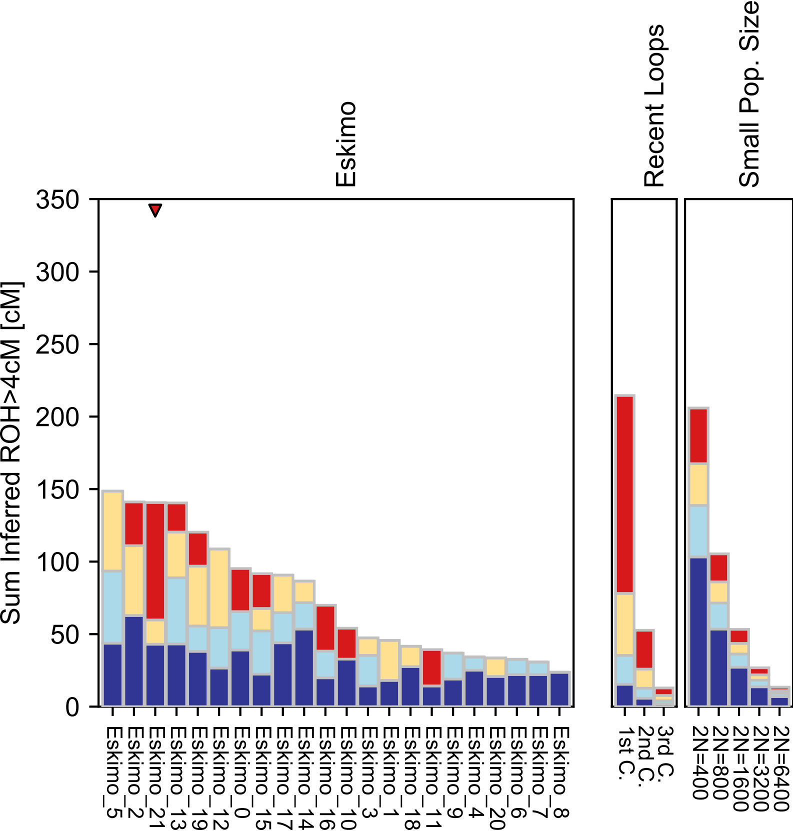

### Estonian.pdf

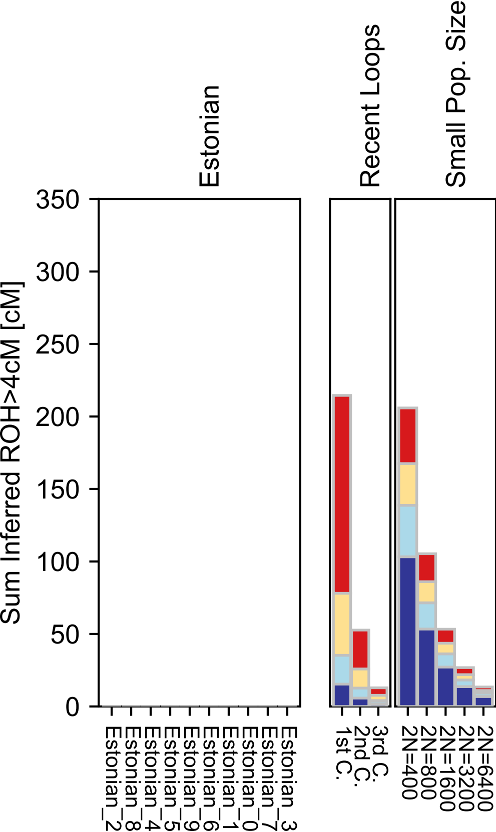

### Ethiopian_Jew.pdf

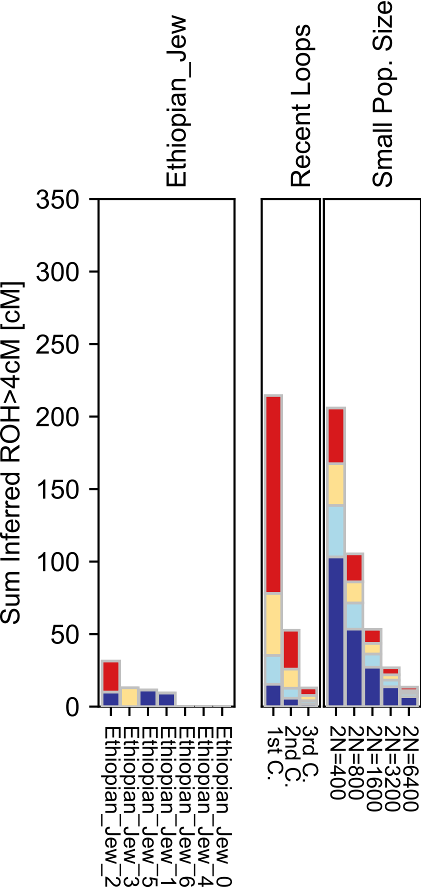

### Even.pdf

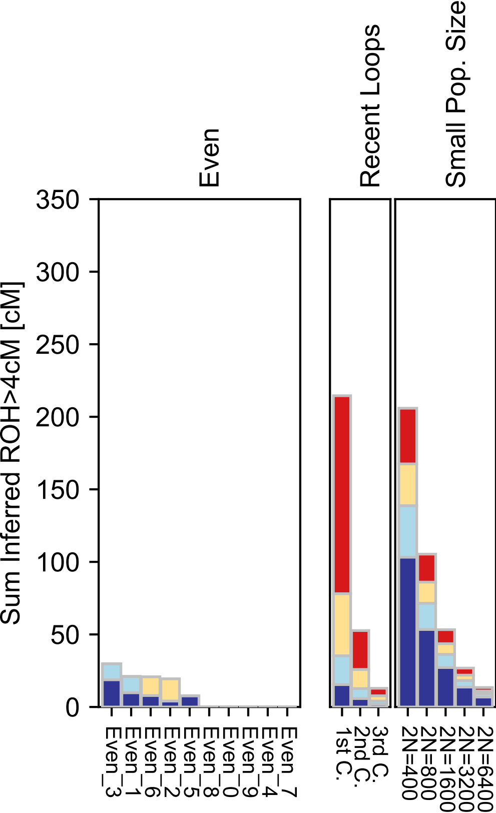

### Finnish.pdf

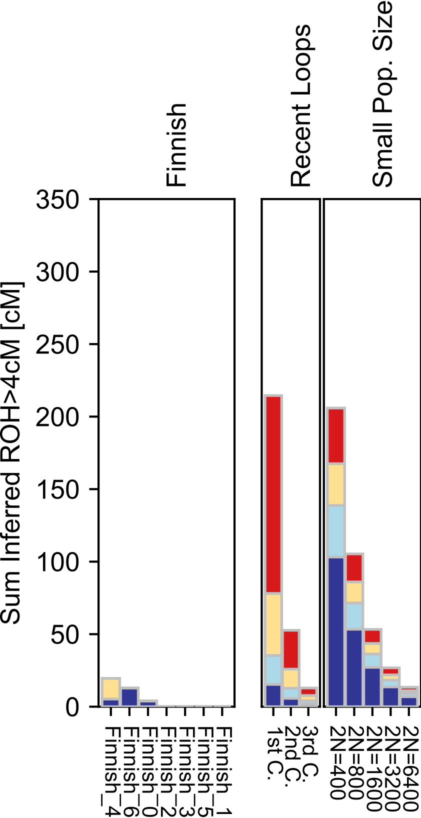

### French.pdf

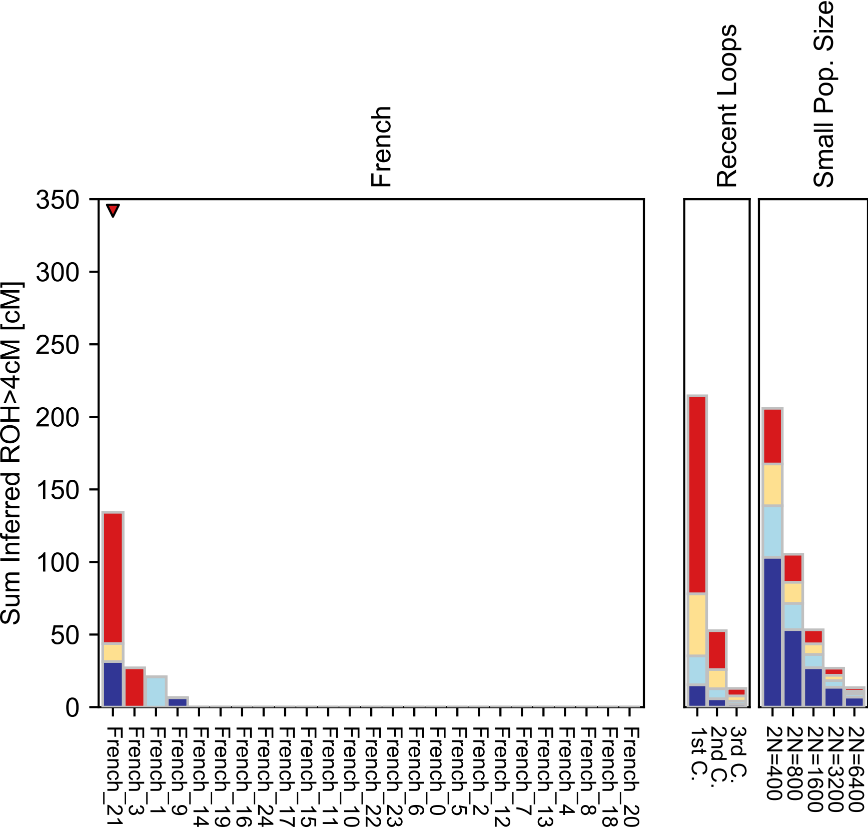

### French_South.pdf

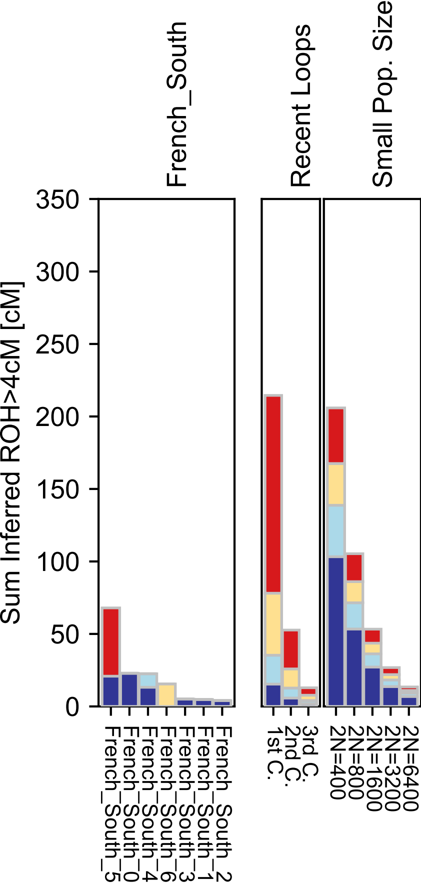

### Gambian.pdf

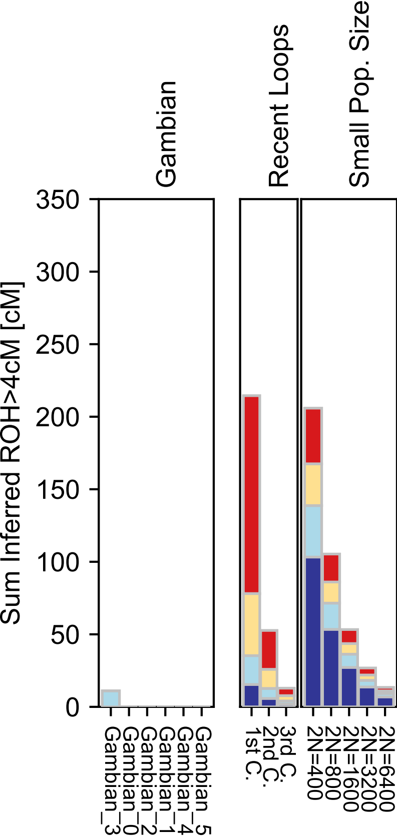

### Georgian.pdf

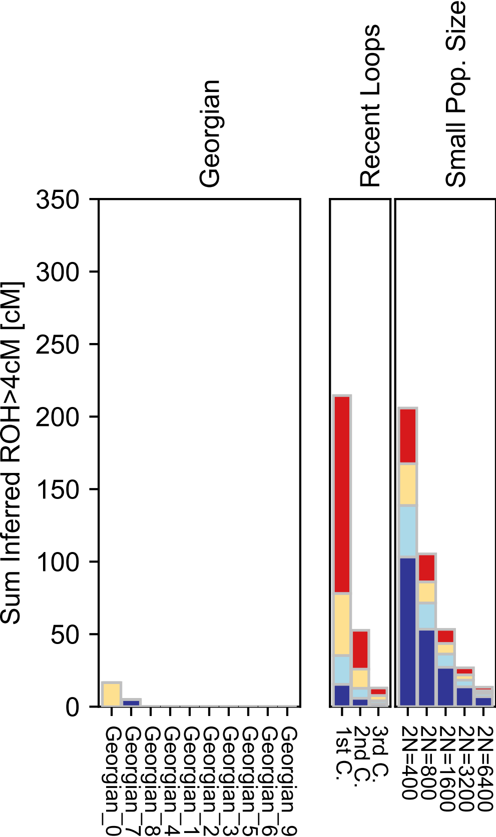

### Georgian_Jew.pdf

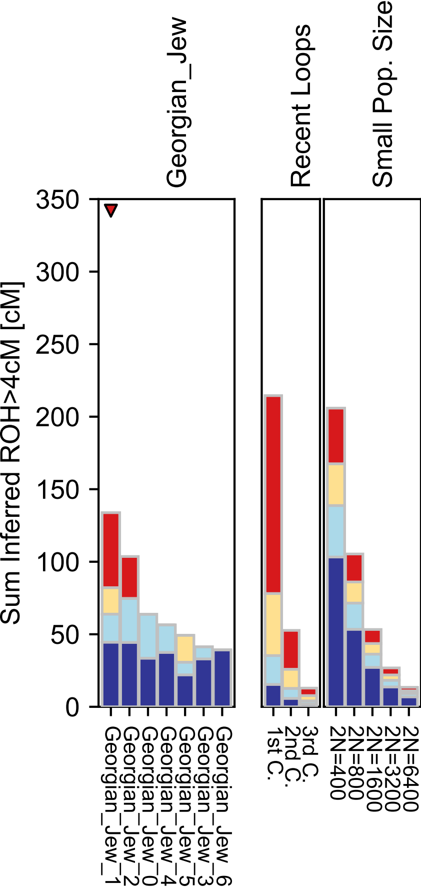

### Greek.pdf

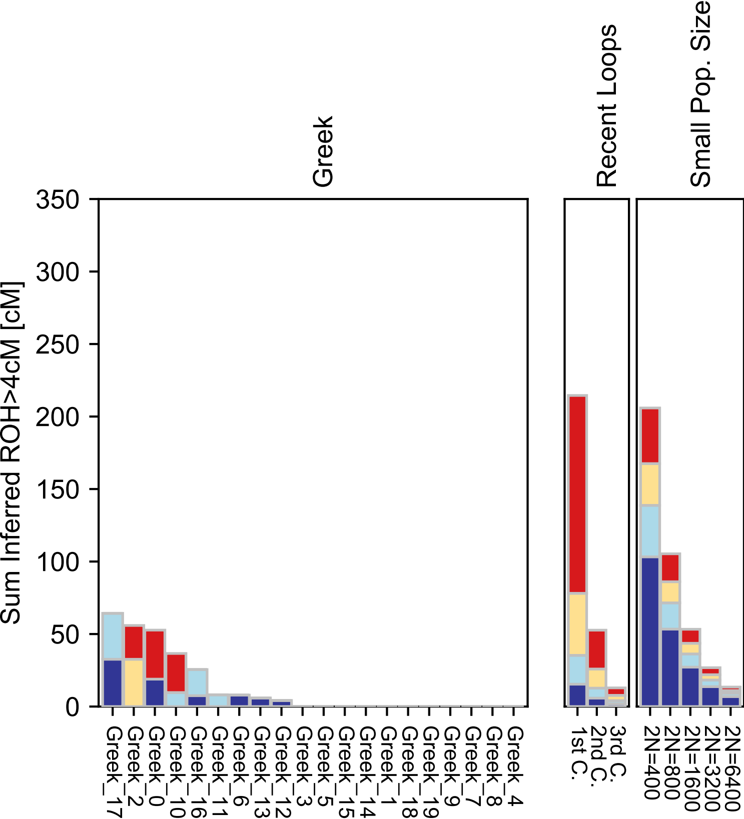

### GujaratiA.pdf

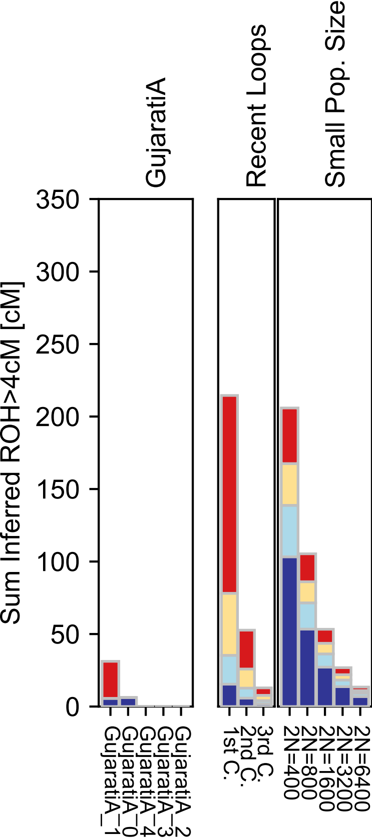

### GujaratiB.pdf

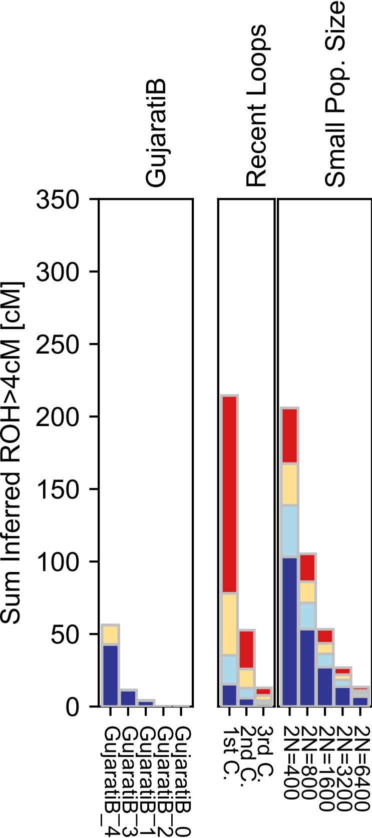

### GujaratiC.pdf

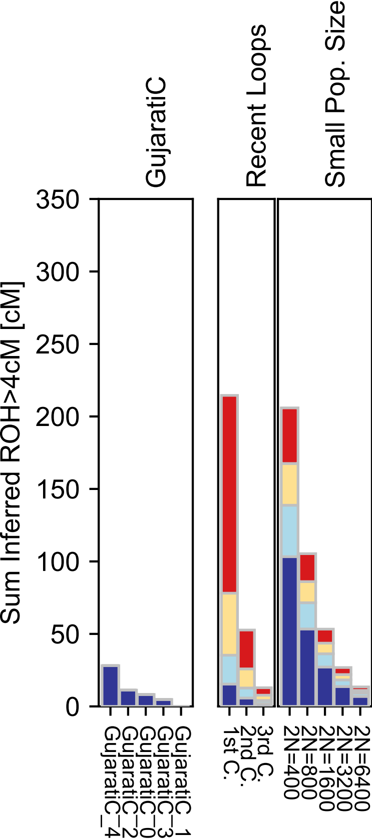

### GujaratiD.pdf

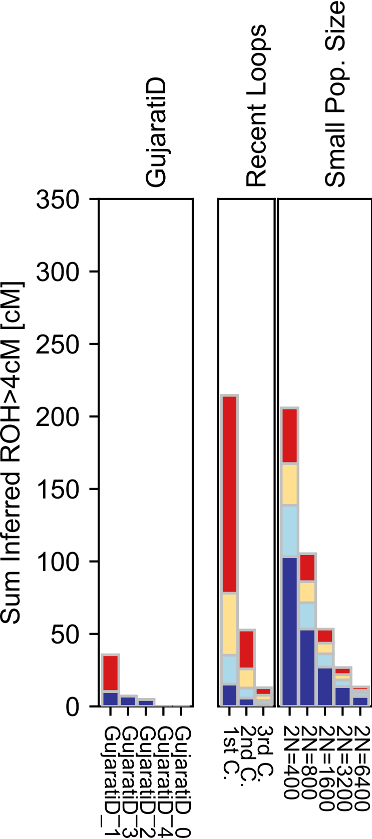

### Hadza.pdf

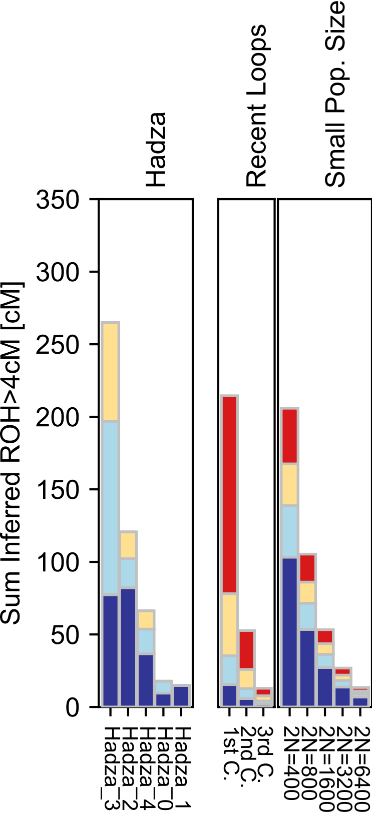

### Han.pdf

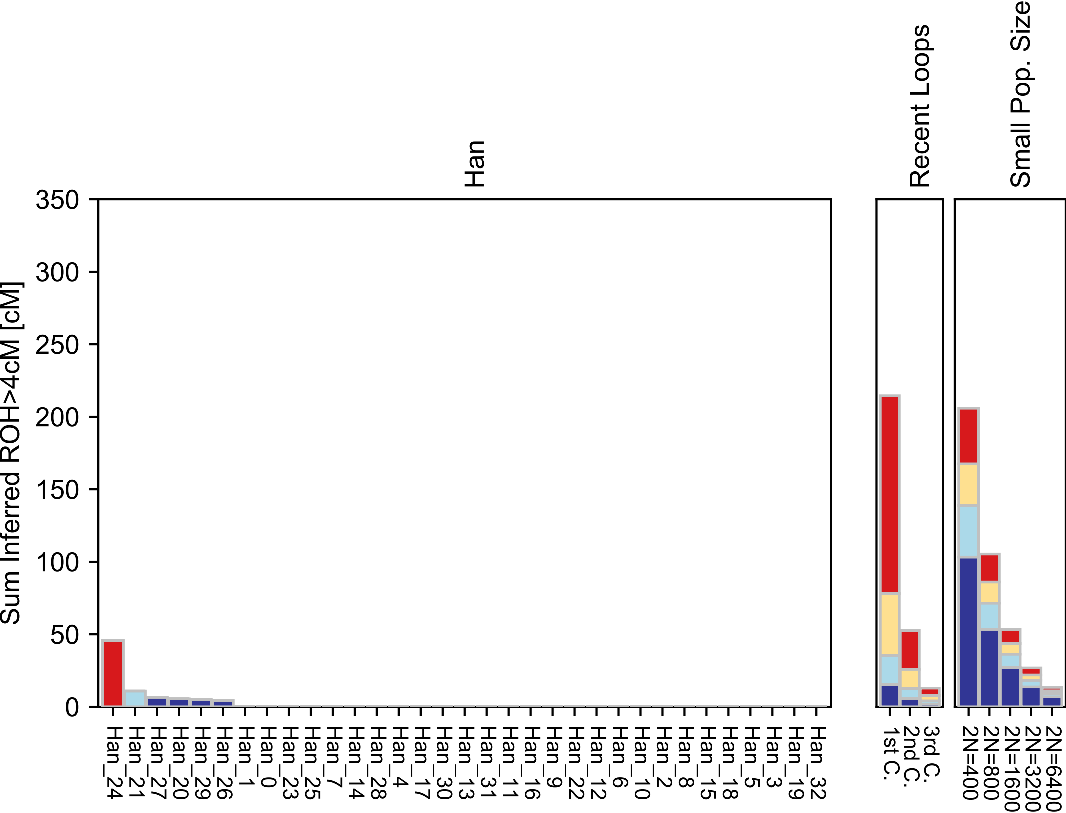

### Han_NChina.pdf

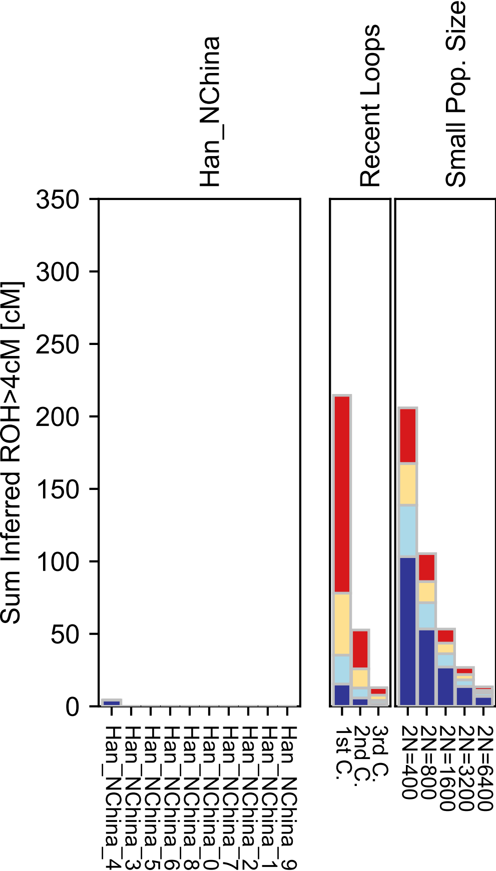

### Hazara.pdf

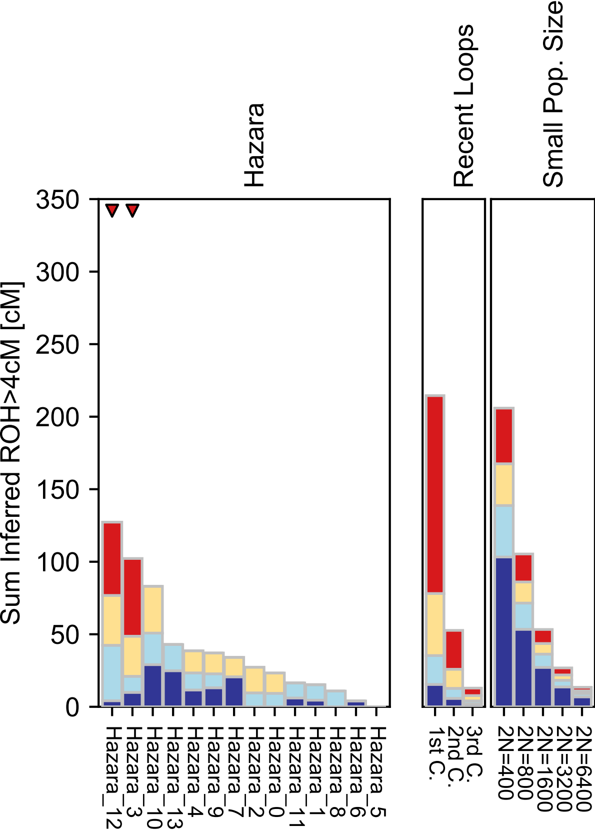

### Hezhen.pdf

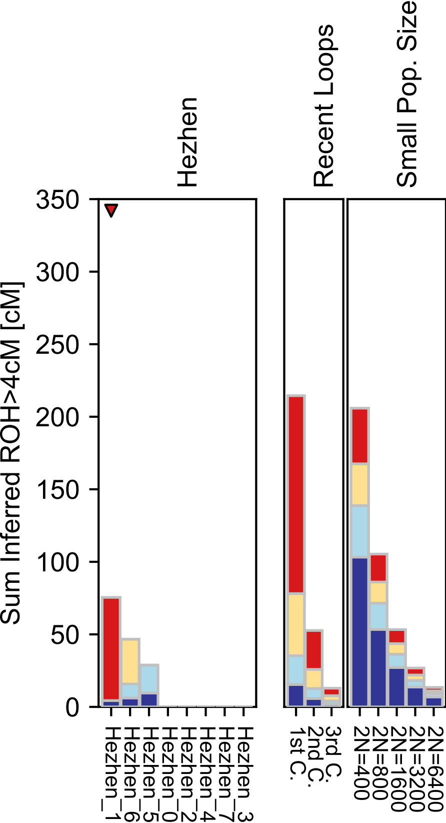

### Hungarian.pdf

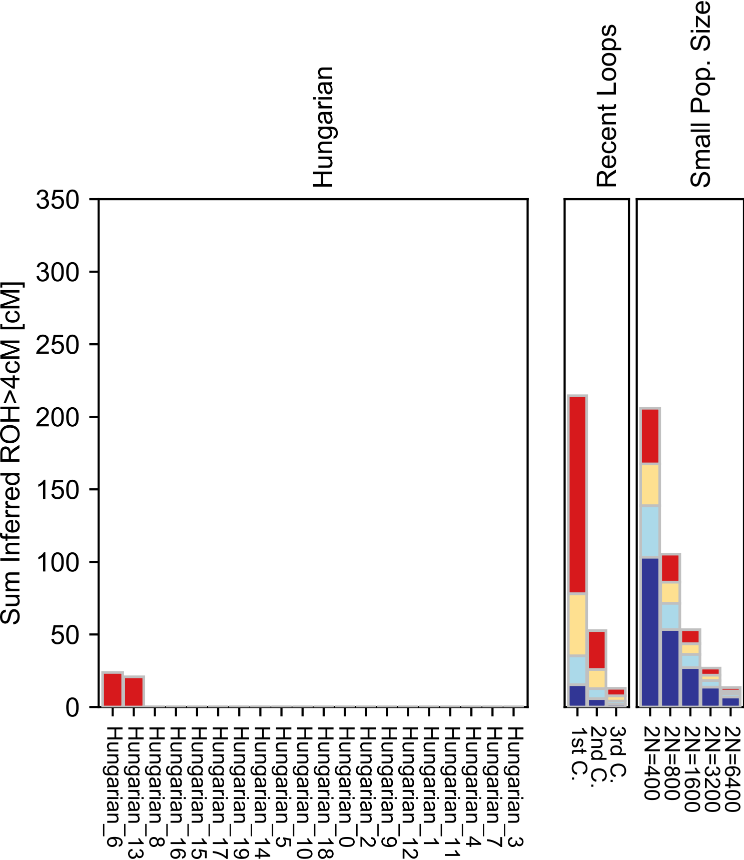

### Icelandic.pdf

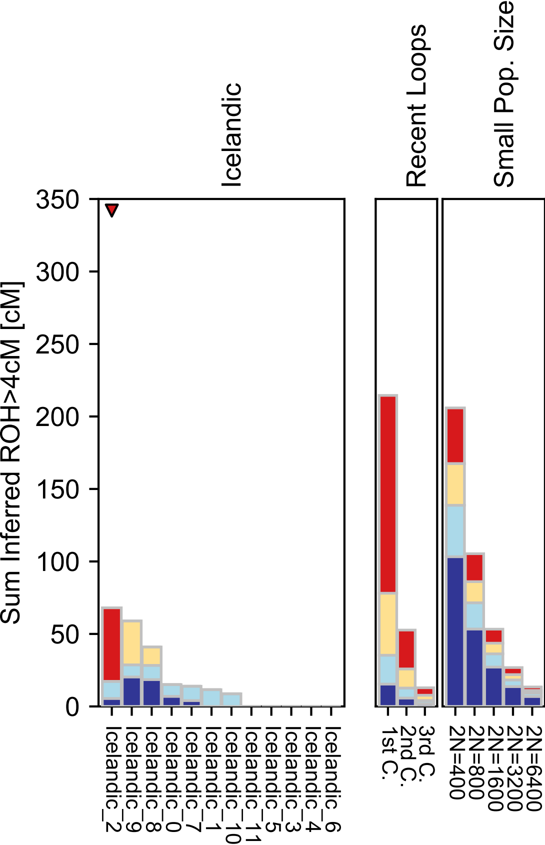

### Iranian.pdf

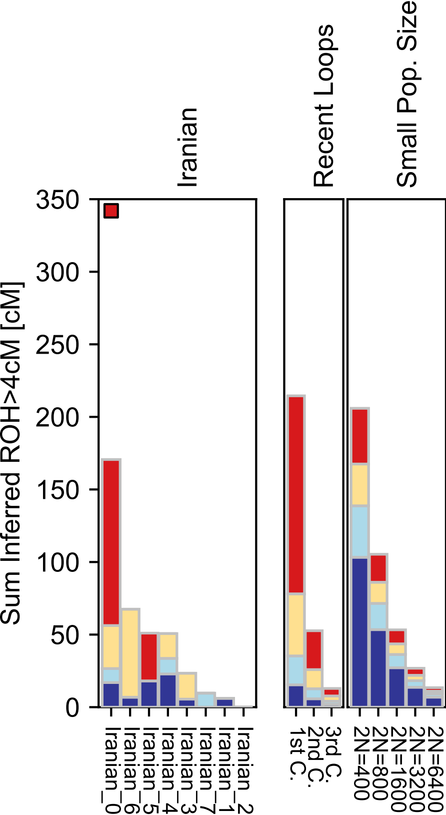

### Iranian_Jew.pdf

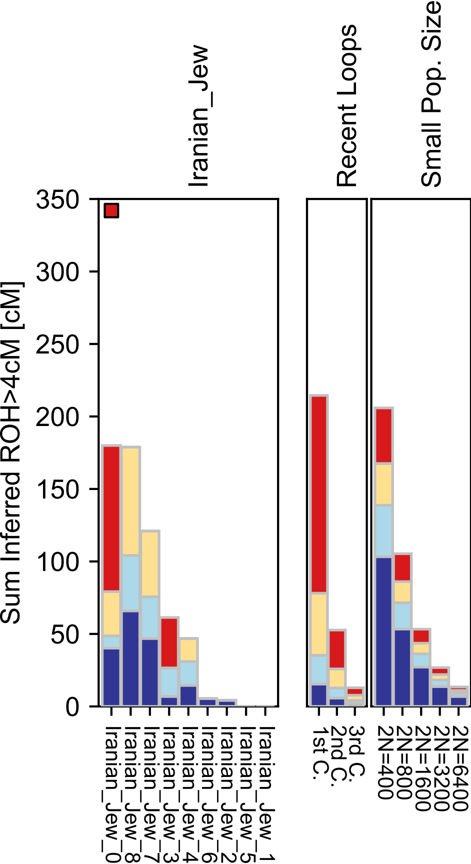

### Iraqi_Jew.pdf

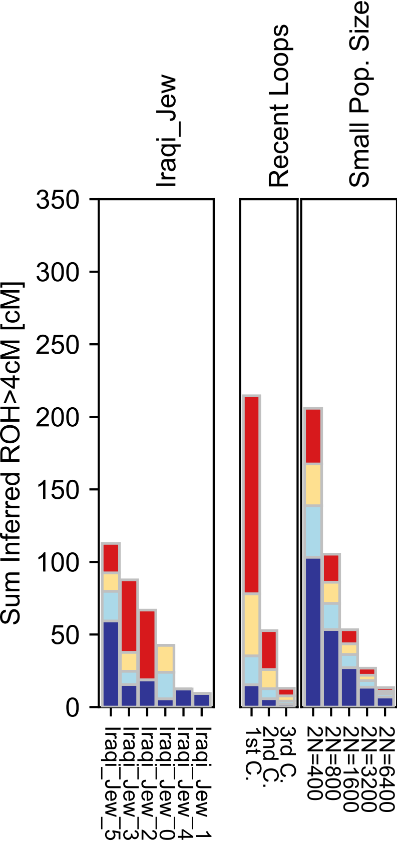

### Italian_South.pdf

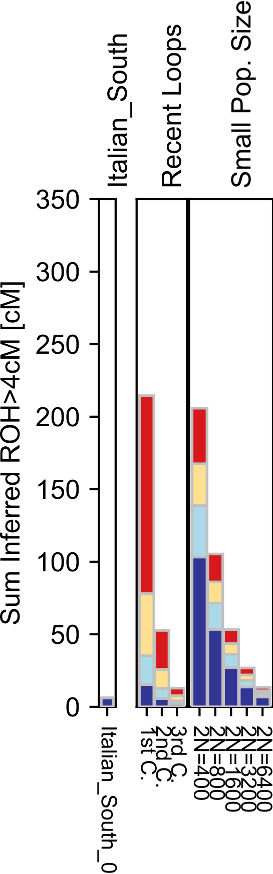

### Itelmen.pdf

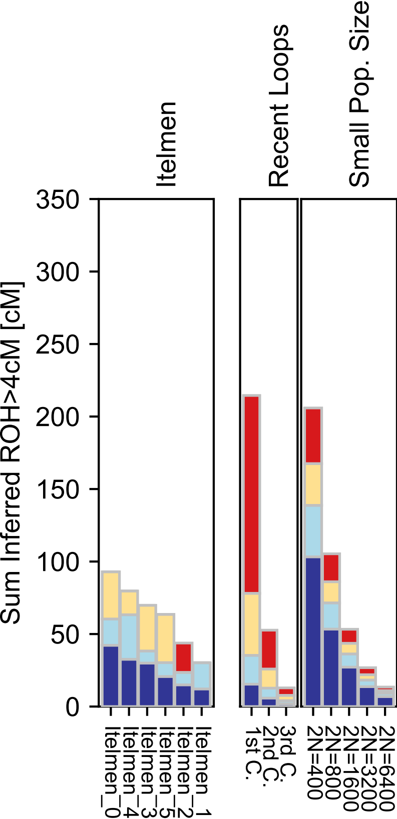

### Japanese.pdf

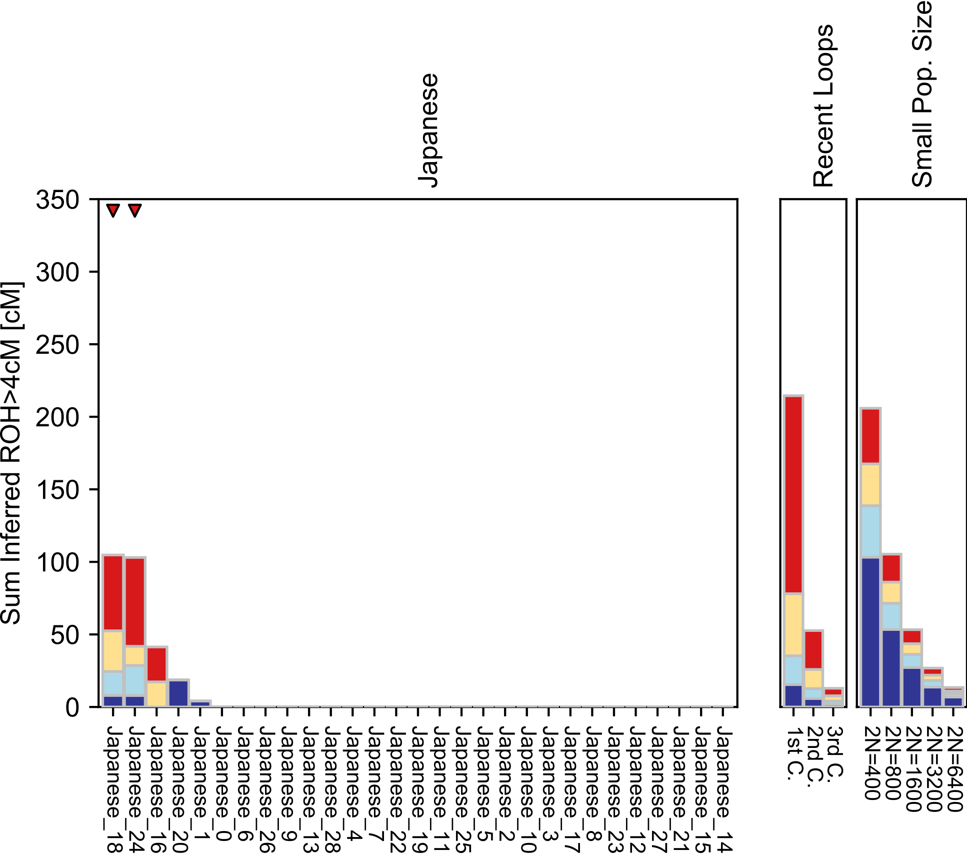

### Jordanian.pdf

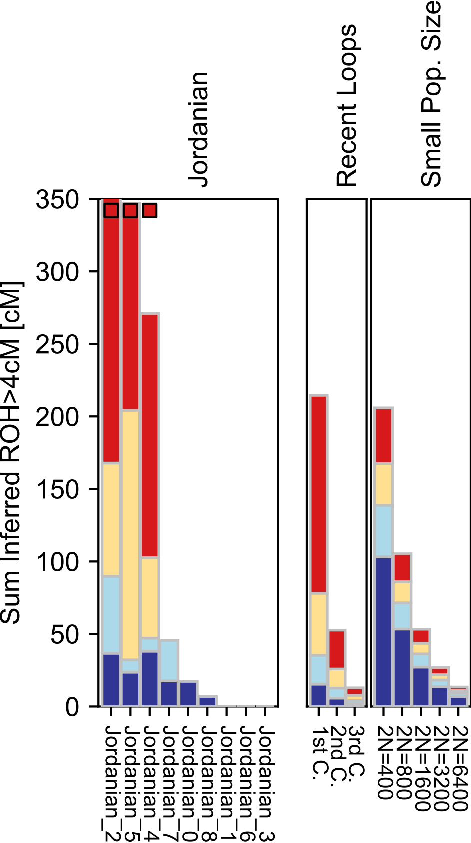

### Ju_hoan_North.pdf

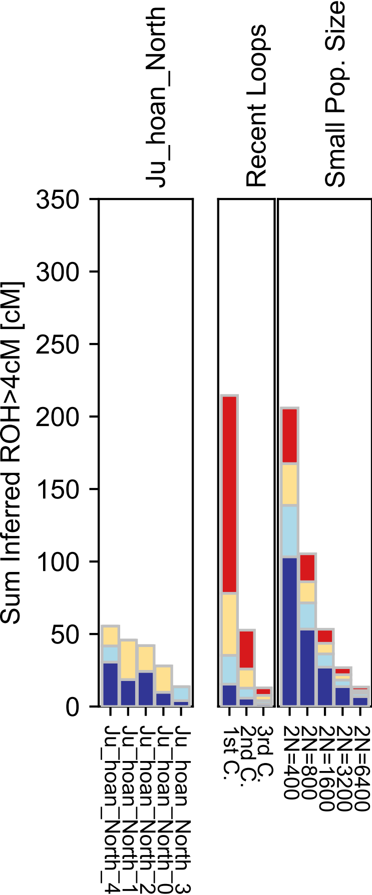

### Kalash.pdf

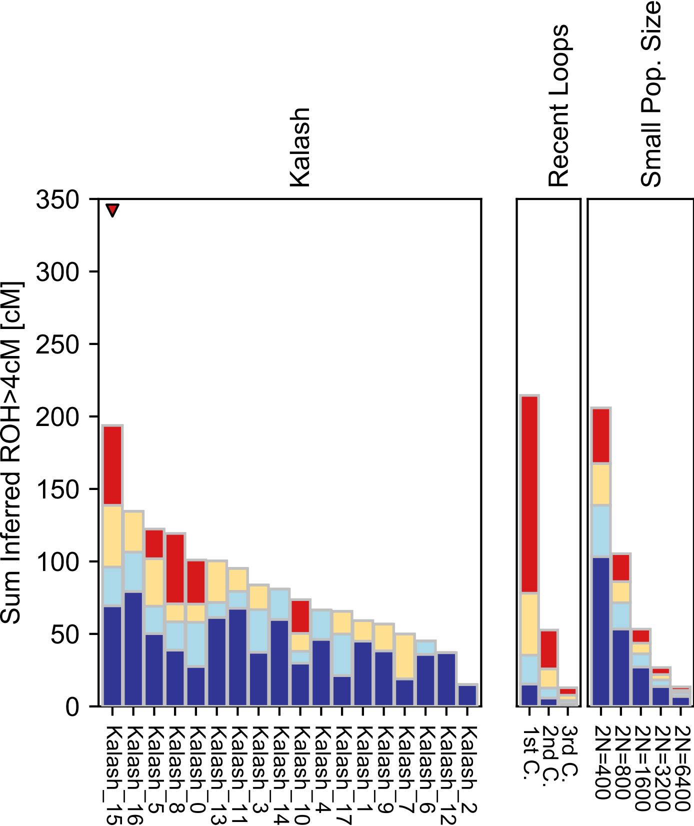

### Kalmyk.pdf

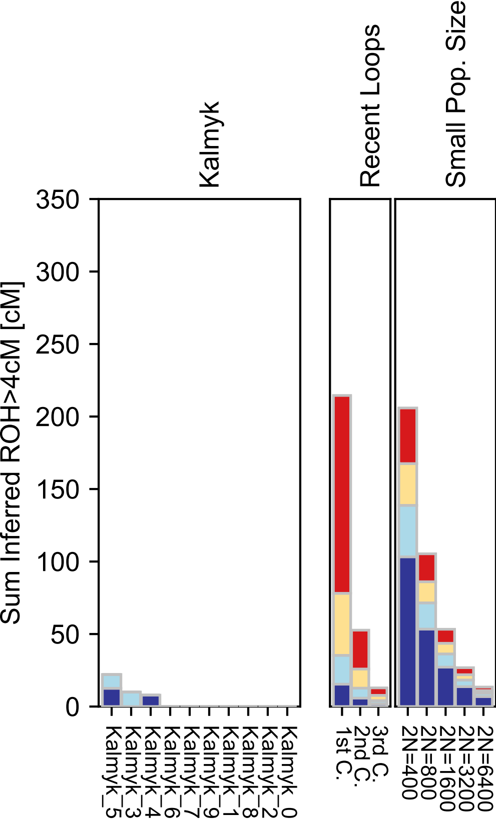

### Karitiana.pdf

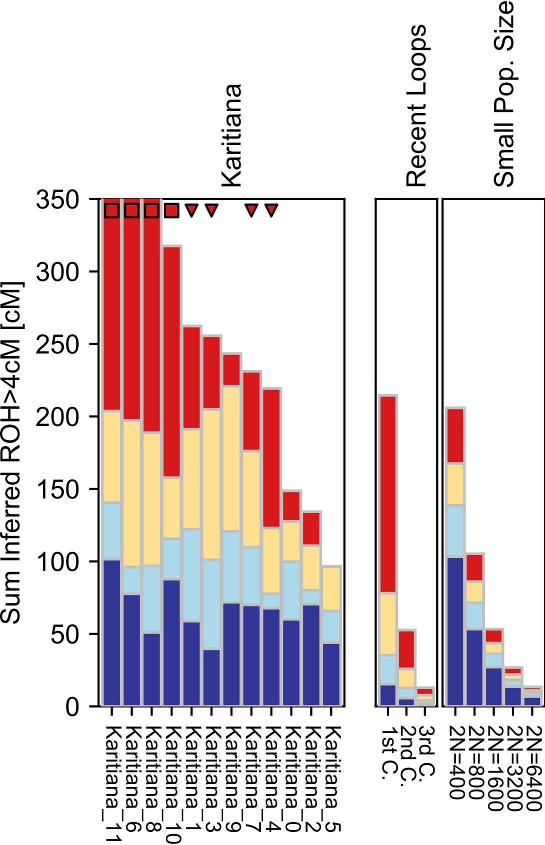

### Khomani.pdf

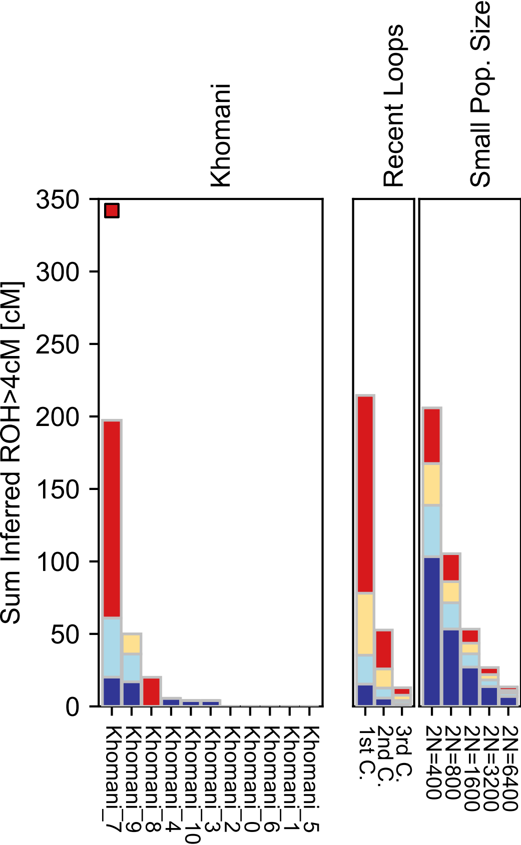

### Kikuyu.pdf

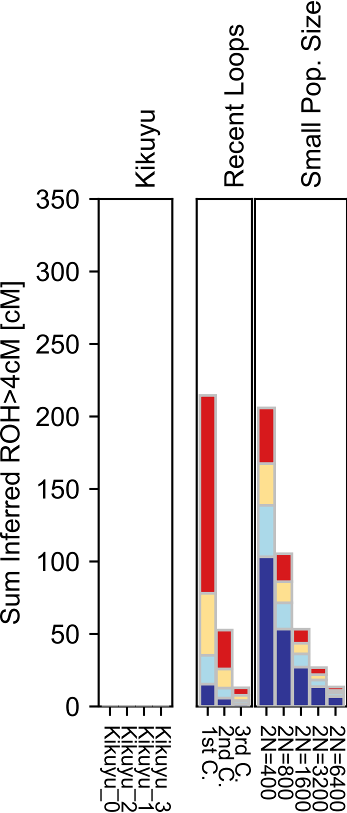

### Kinh.pdf

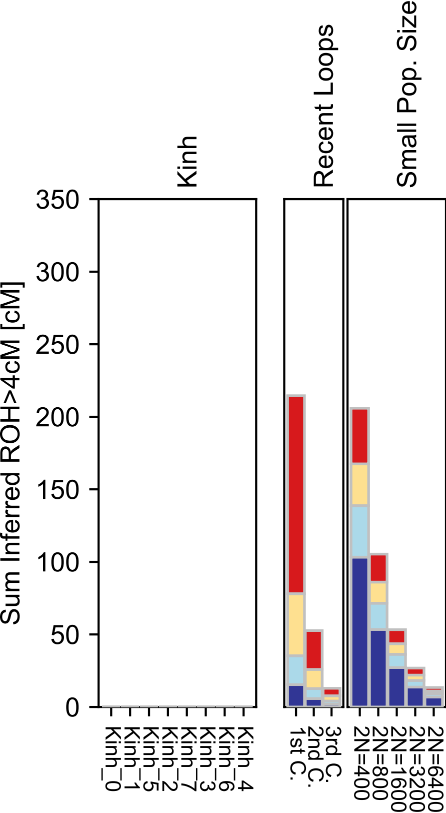

### Korean.pdf

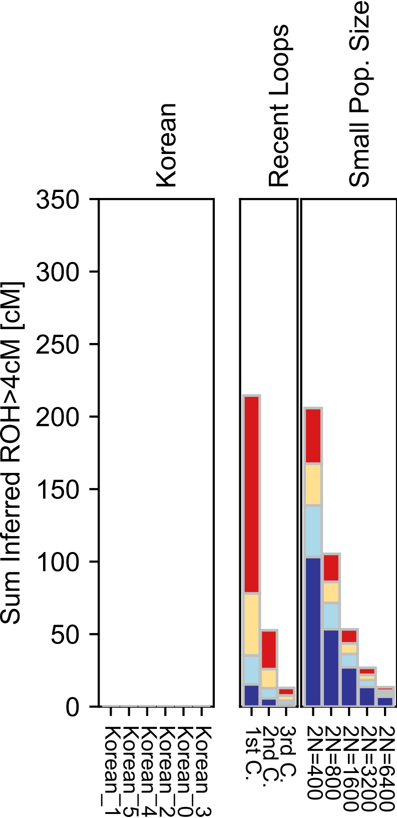

### Koryak.pdf

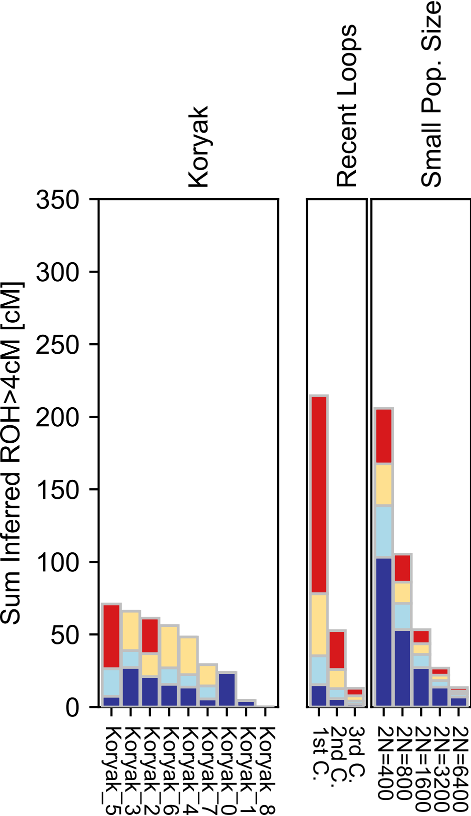

### Kumyk.pdf

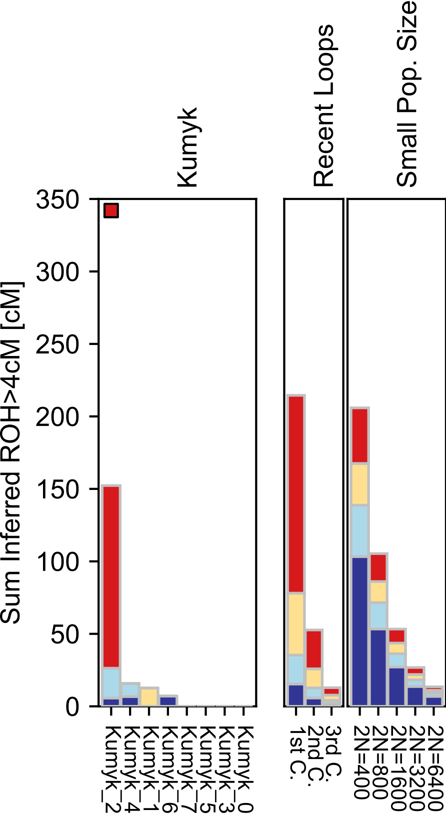

### Kusunda.pdf

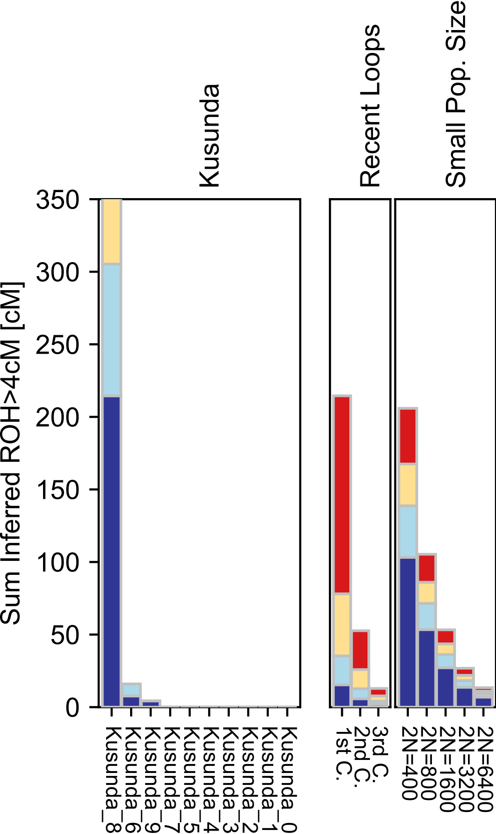

### Kyrgyz.pdf

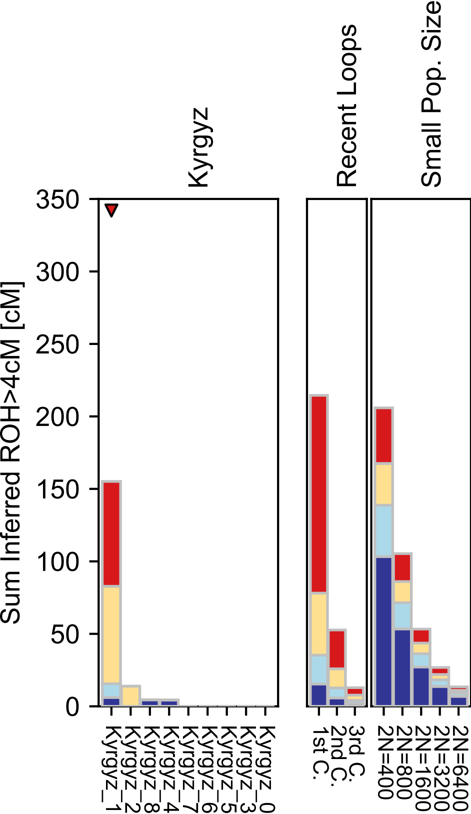

### Lahu.pdf

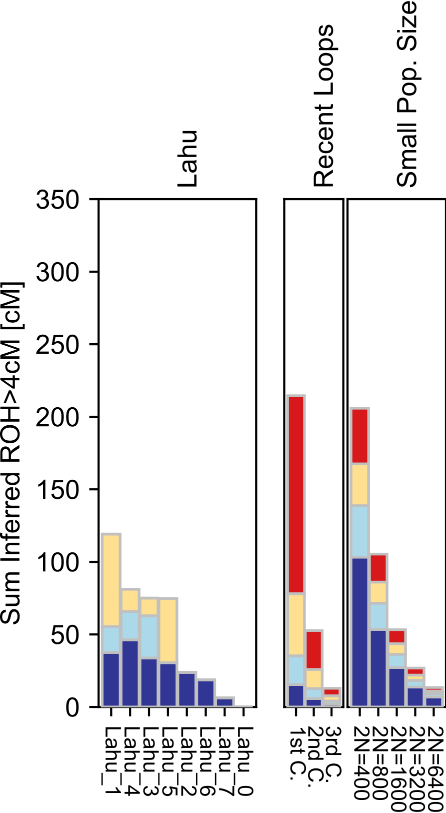

### Lebanese.pdf

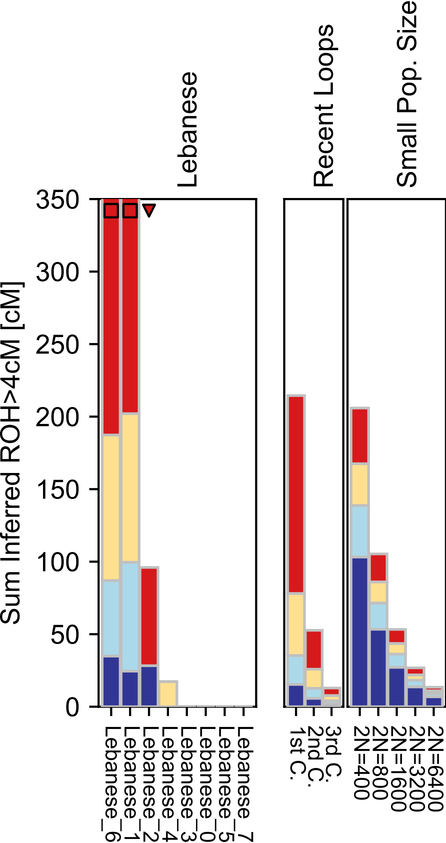

### Lezgin.pdf

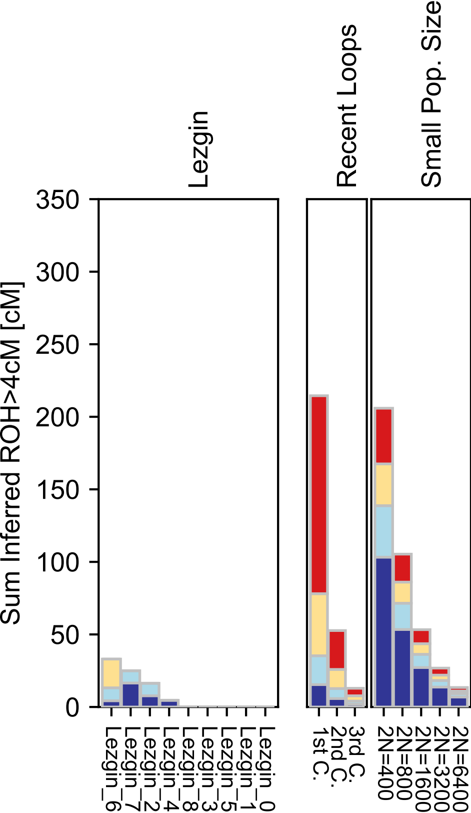

### Libyan_Jew.pdf

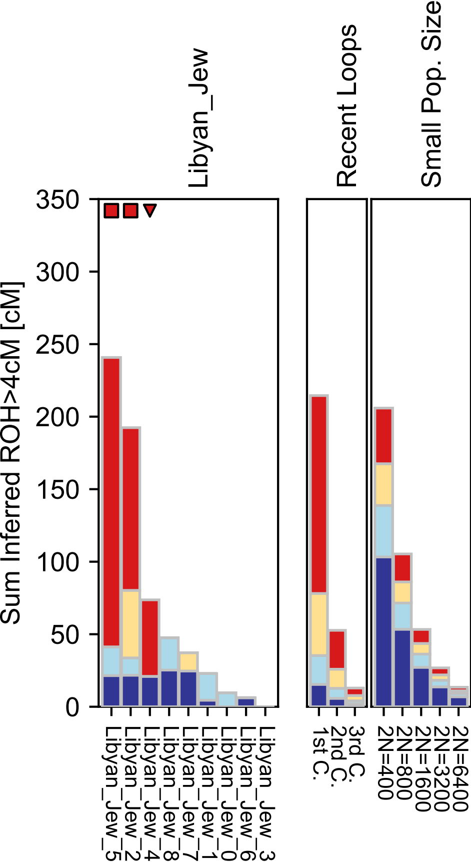

### Lithuanian.pdf

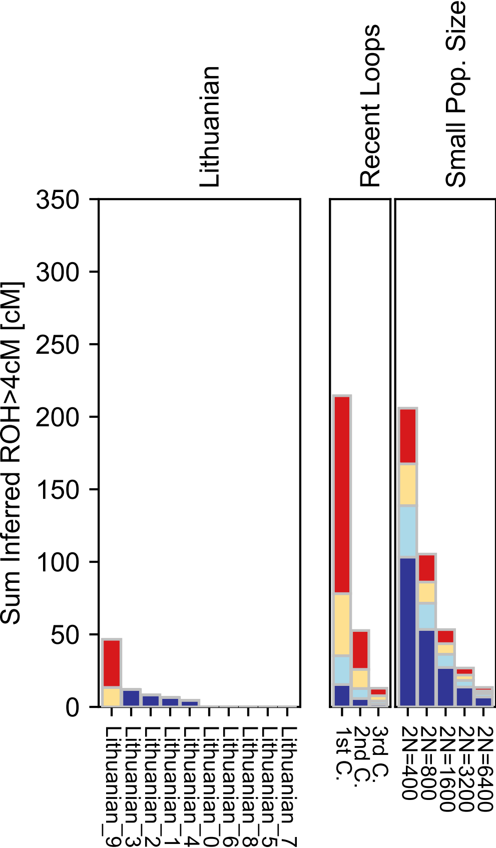

### Luhya.pdf

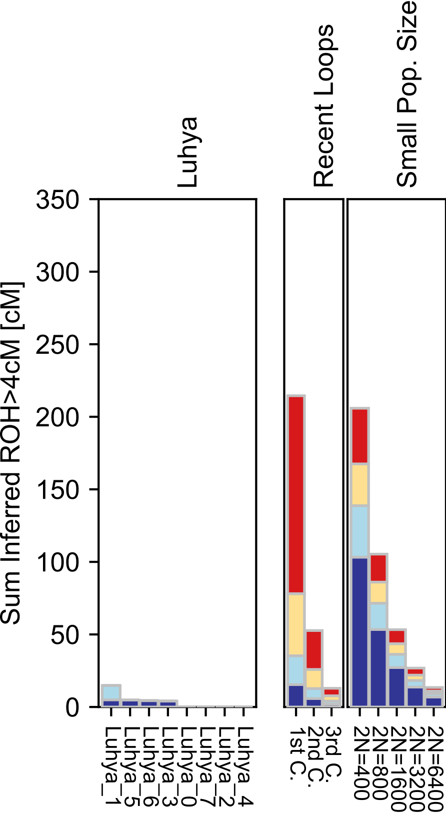

### Luo.pdf

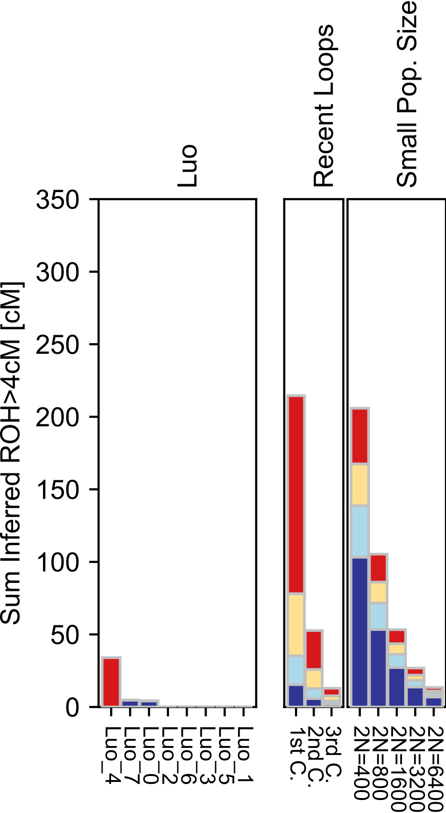

### Makrani.pdf

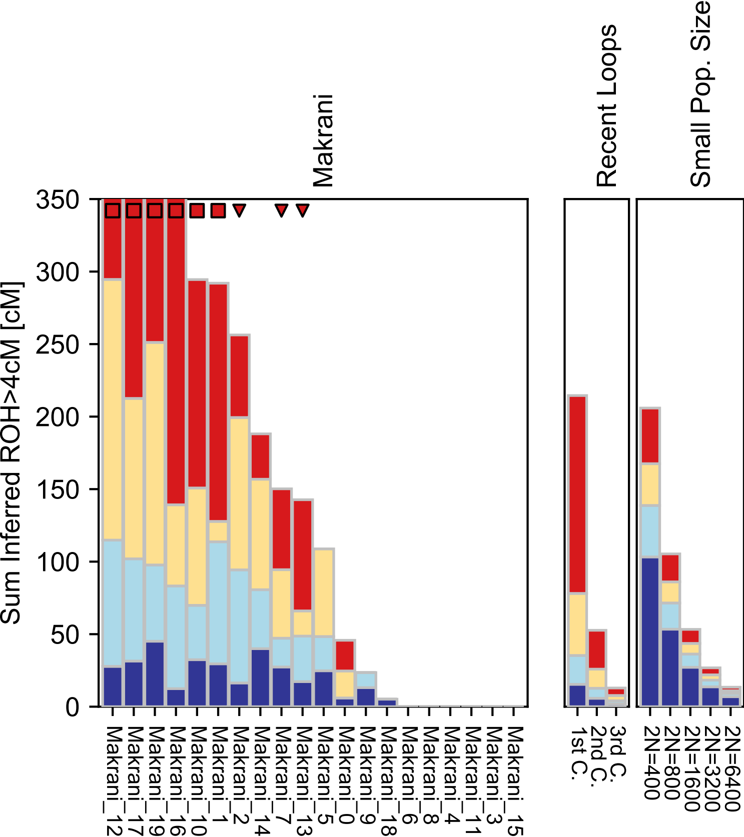

### Maltese.pdf

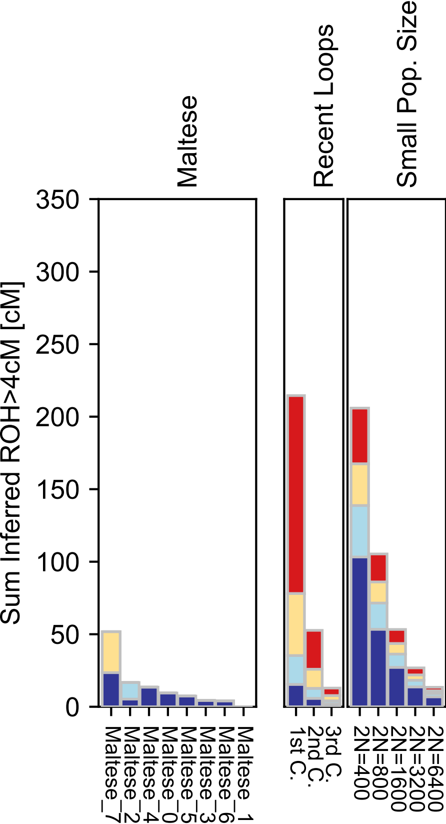

### Mandenka.pdf

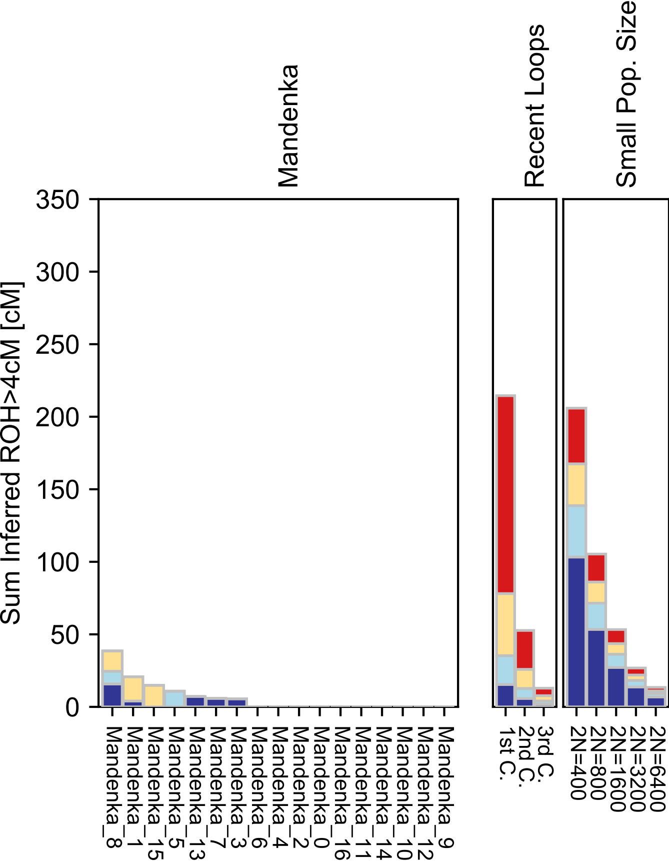

### Mansi.pdf

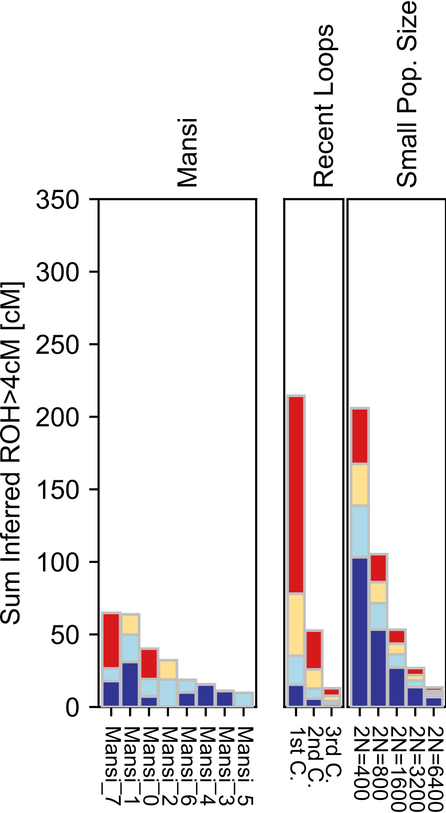

### Masai.pdf

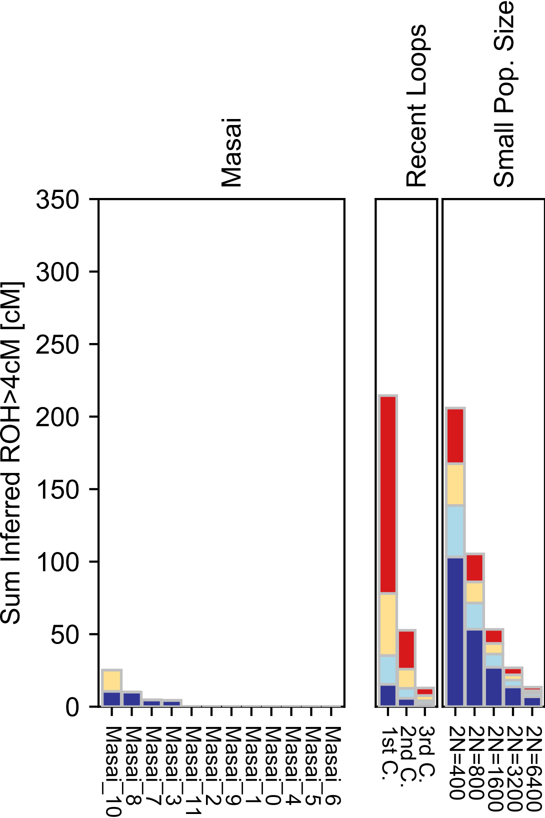

### Mayan.pdf

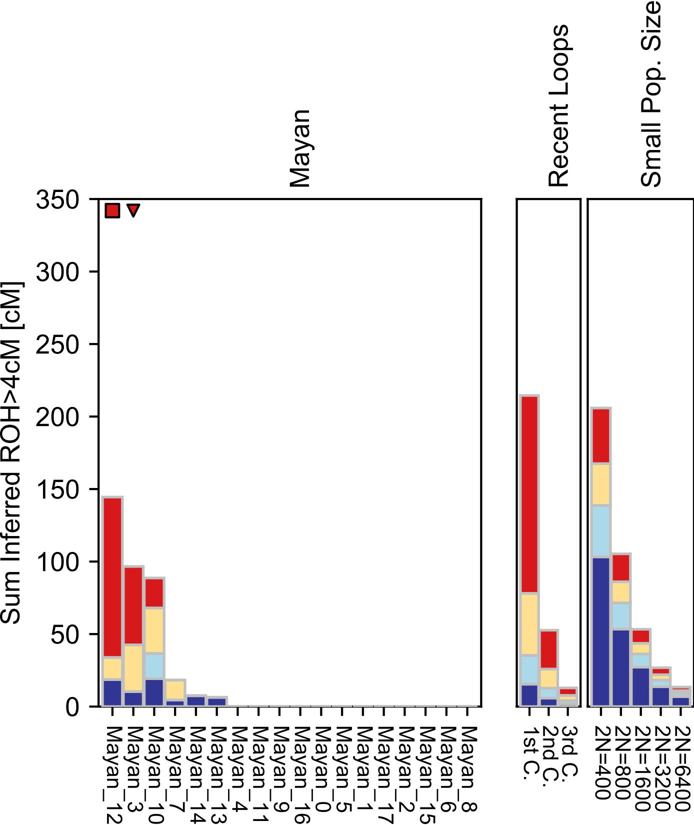

### Mbuti.pdf

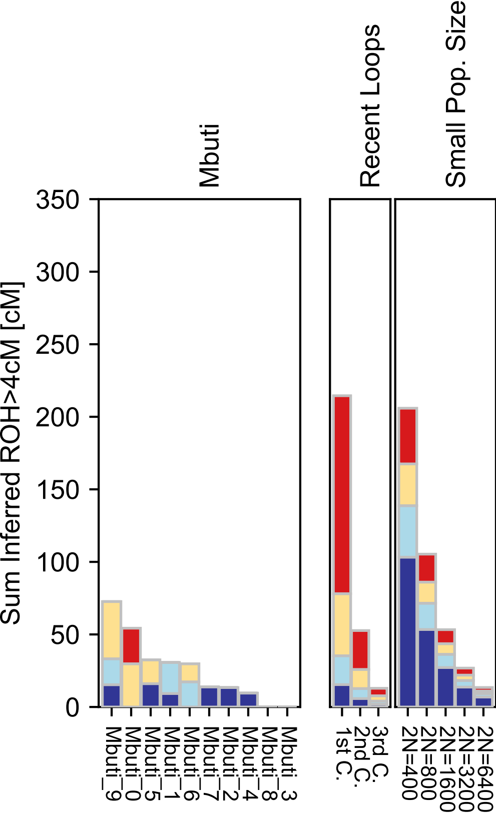
